# *Oryza* genome evolution through a tetraploid lens

**DOI:** 10.1101/2024.05.29.596369

**Authors:** Alice Fornasiero, Tao Feng, Noor Al-Bader, Aseel Alsantely, Saule Mussurova, Nam V. Hoang, Gopal Misra, Yong Zhou, Leonardo Fabbian, Nahed Mohammed, Luis Rivera Serna, Manjula Thimma, Victor Llaca, Praveena Parakkal, David Kudrna, Dario Copetti, Shanmugam Rajasekar, Seunghee Lee, Jayson Talag, Chandler Sobel-Sorenson, Olivier Panaud, Kenneth L. McNally, Jianwei Zhang, Andrea Zuccolo, M. Eric Schranz, Rod A. Wing

**Affiliations:** Biological and Environmental Sciences and Engineering Division (BESE), King Abdullah University of Science and Technology (KAUST), Thuwal, Saudi Arabia; Biosystematics Group, Wageningen University, Droevendaalsesteeg 1, 6708PB Wageningen, The Netherlands; Research and Development, Corteva Agriscience, Johnston, IA, 50131, USA; Arizona Genomics Institute, School of Plant Sciences, University of Arizona, Tucson, AZ 85721, USA; Laboratoire Génome et Développement des Plantes, UMR 5096 UPVD/CNRS, Université de Perpignan Via Domitia, Perpignan, France; International Rice Research Institute (IRRI), Rice Breeding Innovations Department, Los Baños, Laguna, Philippines; National Key Laboratory of Crop Genetic Improvement, Hubei Hongshan Laboratory, Huazhong Agricultural University, Wuhan, 430070, China; Institute of Crop Science, Scuola Superiore Sant’Anna, Pisa, 56127, Italy

## Abstract

*Oryza* is remarkable genus - with two domesticated (i.e. Asian and African rice) and 25 diploid and tetraploid wild species, 11 extant genome types, and a ∼3.4-fold genome size variation - that possesses a virtually untapped reservoir of genes that can be used for crop improvement. Here we unveil and interrogate 11 new chromosome-level assemblies of nine tetraploid and two diploid wild *Oryza* species in the context of ∼15 million years of evolution of the genus. We show that the core *Oryza* (sub)genome across all genome types is only ∼200 Mb and largely syntenic, while the remaining nuclear fractions, spanning ∼80-600 Mb, are intermingled, extremely plastic and rapidly evolving. For the halophyte *O. coarctata*, we show that - despite the detection of gene fractionation in the subgenomes - homoeologous genes are expressed at higher levels in one subgenome over the other in a mosaic form, thereby showing subgenome equivalence. The integration of these 11 new ultra-high quality reference genomes with our previously published genome data sets provide a nearly complete picture of the consequences of natural and artificial selection across the evolutionary history of *Oryza*. This in turn opens the door to unlock their genetic potential for future crop improvement and neodomestication.

## Introduction

The genetic bottleneck imposed by thousands of years of domestication has inevitably impoverished the genetic pool required by the rice crop to adapt to a changing environment^1–3^. New solutions are needed to overcome current and future challenges in rice production and sustainability, and in light of a predicted human population expansion up to 10 billion by 2050^4^. To help reduce this bottleneck, we are exploring and exploiting the standing genetic diversity of the genus *Oryza*, which includes both Asian and African rice, as well as 25 wild species (i.e. 15 diploid genomes with 2n=2x=24 chromosomes and 10 allotetraploid genomes with 2n=4x=48 chromosomes). The wild species, which span 11 extant genome types (i.e. AA, BB, CC, BBCC, CCDD, EE, FF, GG, HHKK, KKLL, HHJJ) and collectively encompass ∼15 million years of evolutionary history^5–7^, represent a crucial resource for tolerance and resistance traits that could be harnessed for crop improvement and/or serve as the raw material for neodomestication^8,9^. Since the late ‘90s, the genome and evolutionary biology of the genus *Oryza* has advanced from single gene trait discovery and cloning^10–12^, to restriction fragment length polymorphism (RFLP)^13^ and physical mapping^14^, to the first reference genome sequences of rice^15–17^, to a set of chromosome-level reference genomes representing the 15 distinct subpopulations of Asian rice^18^. Next steps include the generation of a complete digital GeneBank of cultivated rice as well as a set of ultra high-quality reference genomes of the wild relatives of rice that includes all *Oryza* tetraploid species^19^.

To date, only two tetraploid *Oryza* genomes have been assembled at the chromosome level. Mondal et al. published the first genome draft of the halophytic species *O. coarctata* using Illumina and Nanopore sequencing technology, and identified salinity responsive genes either missing or in low copy number in cultivated rice, showing a loss of the genetic capacity for salt tolerance in cultivated rice^20^. A chromosome-level genome assembly of *O. coarctata* has been published recently^21^. Yu and colleagues released a chromosome-level reference genome of *O. alta* using PacBio data and an optimized protocol for the neodomestication of this species through the editing of key domestication genes, opening to a new era for the improvement of polyploid cereal crops^22^.

Here, we report the generation and interrogation of 11 chromosome-level reference genomes, from nine underutilized wild tetraploid *Oryza* species^23,24^ (i.e. *O. malampuzhaensis* (BBCC), *O. minuta* (BBCC), *O. alta* (CCDD), *O. grandiglumis* (CCDD), *O. latifolia* (CCDD), *O. coarctata* (KKLL), *O. schlechteri* (HHKK), *O. longiglumis* (HHJJ) and *O. ridleyi* (HHJJ)) and two wild diploid species (i.e. *O. australiensis* (EE) and *O. meyeriana* (GG)), using PacBio long-read sequencing technology and Bionano optical validation mapping. These ultra-high quality and near gap-free reference sequences represent a valuable advancement in *Oryza* genome biology, as seven of these genomes were assembled to a chromosome level for the first time. This dataset was used to describe how genome size and composition have evolved across the species in the genus, showing that some species are more malleable than others. The role of transposable elements (TEs) in shaping genome size is particularly evident in the *ridleyi* complex (i.e. *O. ridleyi* and *O. longiglumis*, both HHJJ), where the differential expansion of TEs produced a striking size variation of the homoeologous subgenomes. This mirrors what was observed in the diploid *Oryza* species, i.e. *O. australiensis* (EE) and *O. granulata* (GG), which showed significant increase in genome size due to retrotransposon activity^25,26^. We remodeled the previous phylogenetic tree of the *Oryza* genus^27^ by adding new evidence of the relationships among subgenomes, performed synteny analysis at both macro- and micro-scales to define major chromosomal rearrangements and gene presence-absence variation in the wild species with respect to the AA genomes, explored the extent of gene fractionation in the subgenomes after polyploidization, and investigated subgenome dominance/equivalence in *O. coarctata*. The release of chromosome-level reference genomes of the tetraploid *Oryza* species represents the first step for future research in the fields of evolutionary biology, functional genomics, population genetics, conservation in *Oryza*, and as a robust tool for the neodomestication of climate-adapted rice crops^28–30^.

## Results

### Chromosome-scale genome assemblies of the 11 wild *Oryza* species

Here we report chromosome-level, near gap-free genome assemblies of nine tetraploid and two diploid wild *Oryza* species using a combination of PacBio SMRT/Sequel II sequencing and Bionano Optical Map assembly validation (Figure 1, Table 1, and Supplementary Table 1). We used PacBio CLR and CCS technologies to generate between 37 and 441 Gb of raw reads. The average coverage varied based on the technology used, and ranged between 138- and 450-fold for PacBio CLR sequencing, and between 41- and 46-fold for PacBio CCS sequencing. Average read length, regardless of read type, was reasonably consistent and ranged between a minimum of ∼15 kb for *O. australiensis* (EE) to a maximum of ∼25 kb for *O. alta* (CCDD), *O. ridleyi* (HHJJ) and *O. schlechteri* (HHKK) (Table 1). The wild tetraploid rice species are characterized by a great variation in genome size^8^. The total length of the genomes assembled here have a 2.2-fold variation, varying between ∼556 Mb in *O. coarctata* (KKLL) and ∼1,203 Mb in *O. ridleyi* (HHJJ) (Table 2). Both EE and GG genome type species showed large genome sizes by flow cytometry^31^, and the total length of their assembled genomes was ∼881 and ∼789 Mb for *O. australiensis* (EE) and *O. meyeriana* (GG), respectively (Table 2). We further evaluated the quality of the genome assemblies using BUSCO^32^ with the Poales dataset, and obtained BUSCO scores of “complete genes” ranging between a minimum of 97.9% and a maximum of 99.4%, in *O. australiensis* (EE) and *O. ridleyi* (HHJJ), respectively. The BUSCO score for “complete duplicated genes” changed according to the ploidy, with higher values in the tetraploids due to gene homoeology between subgenomes (Supplementary Table 2 and Supplementary Figure 1).

**Figure 1.**
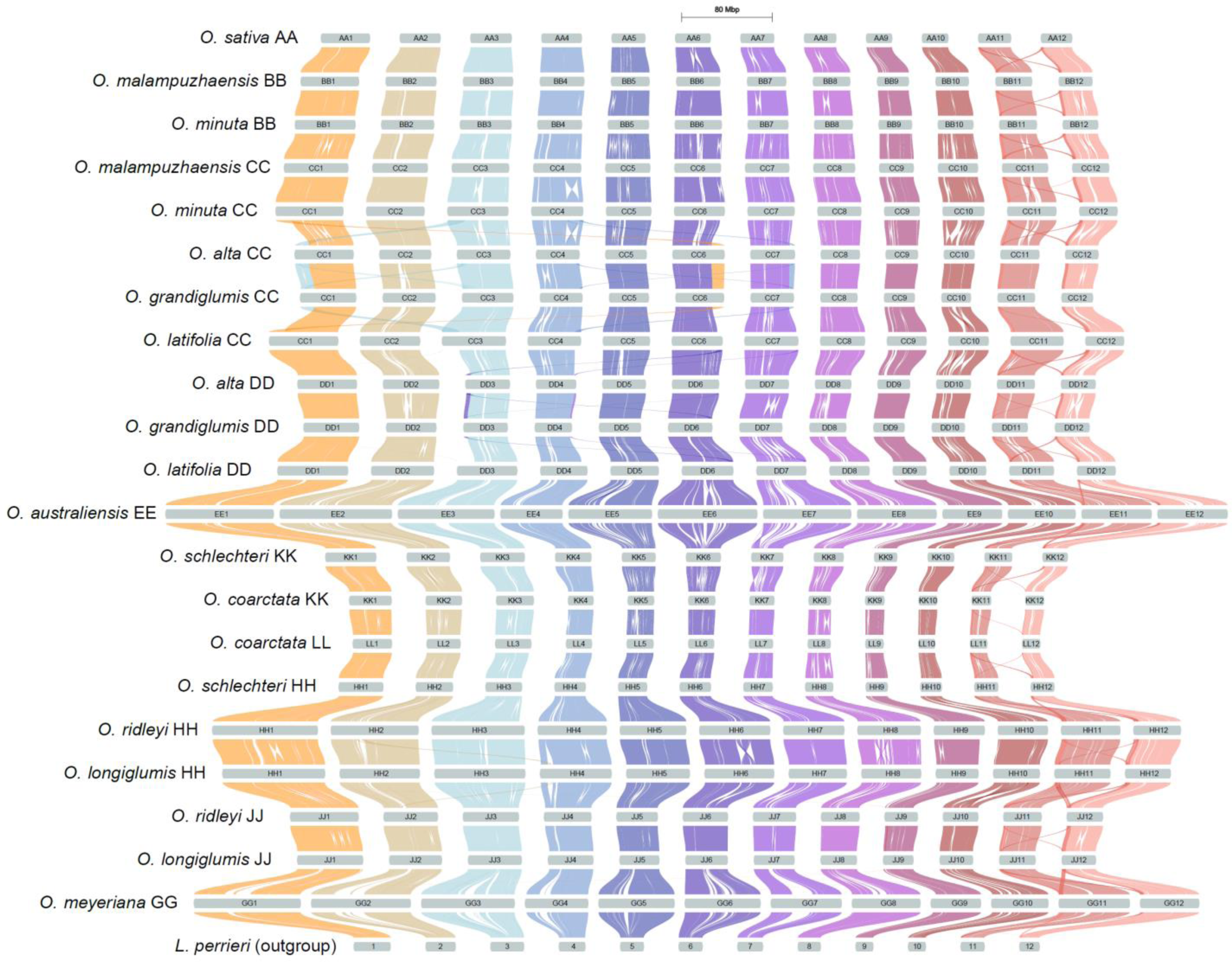
Overview of the syntenic landscape and large-scale structural rearrangements of 12 *Oryza* species (i.e. 21 (sub)genomes) with the outgroup *L. perrieri*. The riparian plot shows macro-syntenic regions and large-scale structural rearrangements (i.e. large duplications and translocations) across the chromosomes of 12 *Oryza* species (i.e. 21 (sub)genomes) and the outgroup species *L. perrieri*. Genome types are shown according to the phylogenetic order in the genus, going from the top (*O. sativa* (AA)) to the bottom (*O. meyeriana* (GG)). Each chromosome is colored as follows: Chr1: orange; Chr2: beige; Chr3: celeste; Chr4: steel blue; Chr5: navy blue; Chr6: deep purple; Chr7: plum; Chr8: magenta; Chr9: raspberry; Chr10: ruby; Chr11: coral; Chr12: salmon. Chromosomes are scaled by assembly length.

**Table 1.** IRGC accessions of *Oryza* species used for the genome assembly and statistics of raw sequencing data.

| Species | IRRI Accession | Country of origin | GRIN-Global link | Sequencing reads | Raw data (Gbp) | Number of subreads | Mean subread length (bp) | Total Assembly length (Mbp) | Coverage (fold) |
| --- | --- | --- | --- | --- | --- | --- | --- | --- | --- |
| *O. alta* | IRGC 105143 | Guyana | https://gringlobal.irri.org/gringlobal/accessiondetail?id=105143 | PacBio CLR | 125 | 4,940,816 | 25,381 | 909.02 | 138 |
| *O. australiensis* | IRGC 100882 | Australia | https://gringlobal.irri.org/gringlobal/accessiondetail?id=100882 | PacBio CLR | 125 | 8,092,633 | 15,446 | 881.42 | 142 |
| *O. coarctata* | IRGC 104502 | Bangladesh | https://gringlobal.irri.org/gringlobal/accessiondetail?id=104502 | PacBio CCS | 107 | 5,458,678 | 19,600 | 555.71 | 193 |
| *O. grandiglumis* | IRGC 105669 | Brazil | https://gringlobal.irri.org/gringlobal/accessiondetail?id=105669 | PacBio CLR | 300 | 16,455,569 | 18,228 | 871.18 | 344 |
| *O. latifolia* | IRGC 100890 | Cuba | https://gringlobal.irri.org/gringlobal/accessiondetail?id=100890 | PacBio CLR | 304 | 15,365,458 | 19,812 | 1058.40 | 288 |
| *O. longiglumis* | IRGC 106525 | Papua New Guinea | https://gringlobal.irri.org/gringlobal/accessiondetail?id=106525 | PacBio CCS | 47 | 2,675,801 | 17,710 | 1146.78 | 41 |
| *O. malampuzhaensis* | IRGC 80765 | India | https://gringlobal.irri.org/gringlobal/accessiondetail?id=80765 | PacBio CLR | 310 | 15,554,278 | 19,960 | 934.73 | 332 |
| *O. meyeriana* | IRGC 106473 | Philippines | https://gringlobal.irri.org/gringlobal/accessiondetail?id=106473 | PacBio CCS | 37 | 2,262,199 | 16,205 | 789.12 | 46 |
| *O. minuta* | IRGC 101141 | Philippines | https://gringlobal.irri.org/gringlobal/accessiondetail?id=101141 | PacBio CLR | 441 | 24,643,407 | 17,899 | 980.38 | 450 |
| *O. ridleyi* | IRGC 100821 | Thailand | https://gringlobal.irri.org/gringlobal/accessiondetail?id=100821 | PacBio CLR | 169 | 6,801,463 | 24,825 | 1202.81 | 140 |
| *O. schlechteri* | IRGC 82047 | Papua New Guinea | https://gringlobal.irri.org/gringlobal/accessiondetail?id=82047 | PacBio CLR | 135 | 5,376,255 | 25,041 | 674.35 | 200 |

**Table 2.** Statistics of the genome assemblies at the *contig* and scaffold level, gene prediction and TE annotation.

|  | *O. malampuzhaensis* | *O. minuta* | *O. alta* | *O. grandiglumis* | *O. latifolia* | *O. australiensis* | *O. coarctata* | *O. schlechteri* | *O. longiglumis* | *O. ridleyi* | *O. meyeriana* |
| --- | --- | --- | --- | --- | --- | --- | --- | --- | --- | --- | --- |
| **Genomic feature** |  |  |  |  |  |  |  |  |  |  |  |
| Genome type | BBCC | BBCC | CCDD | CCDD | CCDD | EE | KKLL | HHKK | HHJJ | HHJJ | GG |
| Chromosomes (#) | 24 | 24 | 24 | 24 | 24 | 12 | 24 | 24 | 24 | 24 | 12 |
| **Assembly** |  |  |  |  |  |  |  |  |  |  |  |
| **Total assembly size (Mbp)** | **934.73** | **980.38** | **909.02** | **871.18** | **1,058.40** | **881.42** | **555.71** | **674.35** | **1,146.78** | **1,202.81** | **789.12** |
| Contig N50 (Mbp) | 19.09 | 20.60 | 18.70 | 19.30 | 35.13 | 44.75 | 22.97 | 22.26 | 51.17 | 53.84 | 66.88 |
| Contig N90 (Mbp) | 8.99 | 11.25 | 10.73 | 12.01 | 15.86 | 7.29 | 16.51 | 9.86 | 31.97 | 33.86 | 49.09 |
| Contig L50 count | 19 | 20 | 19 | 19 | 13 | 7 | 10 | 12 | 9 | 9 | 5 |
| Contig L90 count | 44 | 44 | 42 | 41 | 31 | 24 | 21 | 28 | 20 | 21 | 11 |
| Scaffold N50 (Mbp) | 39.29 | 41.47 | 38.39 | 37.53 | 45.31 | 77.73 | 22.97 | 28.20 | 51.17 | 53.84 | 66.88 |
| Scaffold N90 (Mbp) | 30.49 | 31.40 | 29.35 | 27.76 | 34.13 | 54.84 | 16.51 | 20.81 | 31.97 | 33.86 | 50.49 |
| Scaffold L50 count | 11 | 11 | 11 | 11 | 11 | 5 | 10 | 10 | 9 | 9 | 5 |
| Scaffold L90 count | 21 | 21 | 21 | 21 | 21 | 11 | 21 | 21 | 21 | 21 | 11 |
| GC content (%) | 43.36 | 43.43 | 43.85 | 43.87 | 44.22 | 44.98 | 41.86 | 43.12 | 43.40 | 43.48 | 46.05 |
| **Number of gaps** | **40** | **45** | **34** | **31** | **17** | **67** | **0** | **17** | **2** | **1** | **1** |
| **Complete BUSCO genome assembly (%)** | **99.20** | **99.20** | **99.30** | **99.20** | **99.10** | **97.90** | **98.40** | **99.00** | **99.30** | **99.40** | **98.30** |
| **Annotation** |  |  |  |  |  |  |  |  |  |  |  |
| Transposable Elements (Mbp) | 518.60 | 563.15 | 507.89 | 470.35 | 627.55 | 631.54 | 174.42 | 267.45 | 700.05 | 740.09 | 550.85 |
| Transposable Elements (%) | 55.49 | 57.44 | 55.86 | 53.99 | 59.28 | 71.63 | 31.38 | 39.67 | 61.06 | 61.54 | 69.81 |
| Predicted genes (#) | 77,869 | 74,266 | 66,387 | 66,356 | 73,734 | 39,392 | 55,709 | 70,857 | 73,324 | 82,808 | 45,092 |
| **Complete BUSCO gene prediction (%)** | **94.30** | **89.70** | **93.40** | **93.40** | **93.80** | **88.10** | **86.60** | **81.10** | **90.10** | **95.40** | **90.70** |

### Structural diversity of the *Oryza* (sub)genomes

TE content in the *Oryza* (sub)genomes showed a 7.7-fold variation, ranging from 30.4% of total genome size in LL subgenome of *O. coarctata* (KKLL) to 71.6% in *O. australiensis* (EE). The most abundant group of class-I TEs was the Ty3-*Gypsy* LTR-retroelement (with an average of 24.3% across the (sub)genomes), and the most abundant group of class-II DNA transposons was CACTA (with an average of 6.3% across the (sub)genomes) (Supplementary Table 3; Supplementary Figure 2). As shown in other plant systems^33^, (sub)genome size highly correlates with TE content in the *Oryza* genus (Pearson’s correlation coefficient: 0.9987) (Supplementary Table 4; Figure 2a). The diploid species with the largest genomes, i.e. *O. australiensis* (EE) and *O. meyeriana* (GG), showed the highest TE content (i.e. ∼632 and ∼551 Mb, respectively) and the highest TE content to genome size ratios (i.e. 71.6 and 69.8%, respectively) (Supplementary Table 3). Among the tetraploid species, the HH subgenome of *O. ridleyi* (HHJJ) and *O. longiglumis* (HHJJ) showed the highest TE content (i.e. ∼477 and ∼453 Mb, respectively), the largest sizes (i.e. ∼728 and ∼695 Mb, respectively), and the highest TE content to subgenome size ratios (i.e. 65.5 and 65.2%, respectively). The LL and HH subgenome of *O. coarctata* (KKLL) and *O. schlechteri* (HHKK), respectively, showed the lowest TE abundance (i.e. ∼82 and ∼117 Mb, respectively), the smallest genome sizes (i.e. ∼271 and ∼315 Mb, respectively), and the lowest TE content to genome size ratios (i.e. 30.4% and 37.0%, respectively) (Figure 2; Supplementary Figure 2).

**Figure 2.**
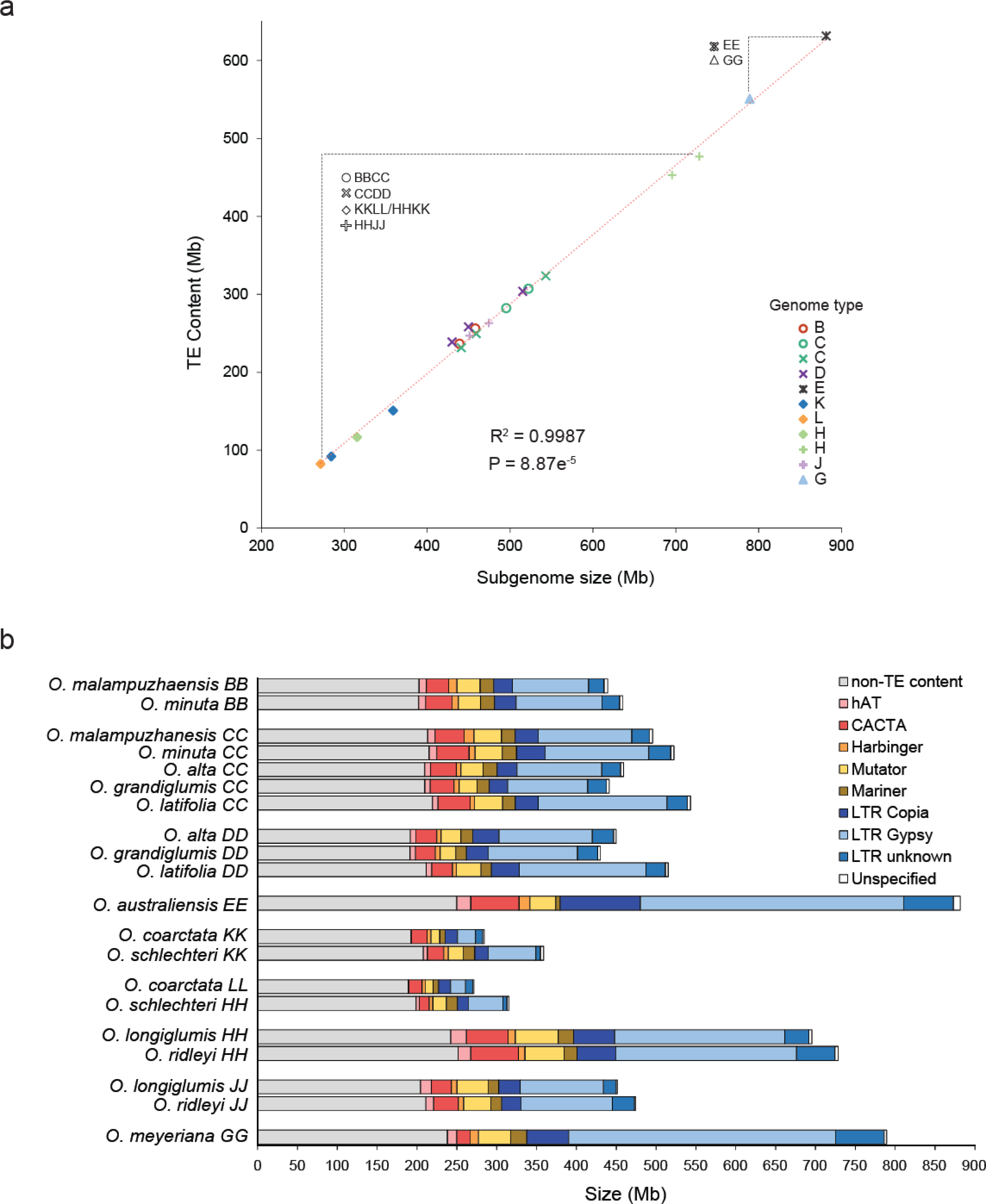
(Sub)genome size and TE content of the *Oryza* species. **a)** Correlation between (sub)genome size (Mb) and TE content (Mb) in the nine tetraploids and two diploid species presented here. Significance of the linear correlation (Pearson′s correlation coefficient, R^2^) was ascertained with a two-sided t-test (p-value, P). **b)** Abundance of the main classes of TEs (Mb) in the nine tetraploids and two diploid species presented here. DNA transposons are shown as: hAT (DTA) - pink; CACTA (DTC) - red; Harbinger (DTH) - orange; Mutator (DTM) - yellow; Mariner (DTT) - ocra. LTR retrotransposons are shown as: LTR Copia - dark blue; LTR Gypsy - light blue; LTR Unknown - steel blue. Unspecified TEs are shown in white; non-TE content is shown in grey. (Sub)genomes of the species are ordered by genome type (i.e. BB, CC, DD, EE, KK, LL, HH, JJ, GG).

Interestingly, the portion of (sub)genomes not populated with TEs (non-TE content in Figure 2 b), despite their ploidy and genome size, ranged between ∼189 and ∼251 Mb (Supplementary Table 4). Gene prediction in the tetraploid species yielded between 55,709 genes in *O. coarctata* (KKLL) to 82,808 in *O. ridleyi* (HHJJ); in the diploid species, 39,392 genes were predicted in *O. australiensis* (EE) and 45,092 in *O. meyeriana* (GG) (Table 2). BUSCO scores were used to evaluate the completeness of gene prediction. The scores for “complete genes” at the whole-genome level ranged between 81.1% in *O. schlechteri* (HHKK) and 95.4% in *O. ridleyi* (HHJJ) (Supplementary Table 2 and Supplementary Figure 1). In the tetraploid species, the sum of BUSCO “missing genes” in each subgenome was higher than the number of BUSCO “missing genes” at the whole-genome level, suggesting gene fractionation in the subgenomes after allopolyploidization.

### Dynamics of differential transposable element amplification in the *ridleyi* complex

Species showing the largest genome sizes in the *Oryza* genus belong to the *ridleyi* complex - i.e. the HHJJ genome species *O. longiglumis* (genome size: 1,147 Mb) and *O. ridleyi* (genome size: 1,203 Mb). Subgenome size was strikingly different in these species, with the HH subgenome showing a ∼1.5-fold variation with respect to the JJ subgenome (Supplementary Table 4). Analysis of TEs showed that size variation in the subgenomes of *O. longiglumis* and *O. ridleyi* was attributed to a difference in TE abundance. The ratio of TE content over non-TE content in HH subgenomes was 1.87 and 1.90 for *O. longiglumis* and *O. ridleyi*, respectively, while the same ratios for the JJ subgenomes were calculated as 1.21 and 1.25 for *O. longiglumis* and *O. ridleyi*, respectively (Figure 2b).

To investigate the preferential expansion of TEs in the HH subgenomes, we investigated the distribution of TEs belonging to six *Oryza*-specific superfamilies (i.e. CACTA, Ty1/Copia, Ty3/Gypsy, MuDR, hAT, LINE) in each subgenome and generated neighbour-joining trees. This analysis did not reveal any evidence for preferential expansion of a TE superfamily over others (Supplementary Figure 8 and Supplementary Figure 9). In these species, the majority of LTR-RTs amplified after polyploidization, estimated ∼2.5-3 Mya^27^ (i.e. 80% of LTR-RT amplification occurred in the last ∼2 Mya in the HH and JJ subgenomes of *O. ridleyi* and in the last ∼2.5 Mya in the HH and JJ subgenomes of *O. longiglumis*, respectively; Supplementary Figure 10). We then determined whether the variation in TE content in the subgenomes was due to a differential rate of either TE accumulation or TE removal in one of the two subgenomes. Unequal recombination and illegitimate recombination serve as mechanisms for LTR-RT elimination. Unequal recombination generates solo-LTRs by recombining LTRs within or between different LTR-RTs, while illegitimate recombination acts on dissimilar DNA sequences removing sections of TE sequences and occasionally leaving incomplete elements^34,35^. To assess TE removal efficacy in *O. ridleyi* and *O. longiglumis*, we calculated the ratio of solo-LTRs to complete LTR-RT elements in each subgenome. We found no relevant difference in the ratios when comparing the HH and JJ subgenomes. In *O. ridleyi*, the ratio of solo-LTRs to complete LTR-RTs in HH and JJ subgenome was 1.2 and 1.5, respectively. In *O. longiglumis*, the ratio was 1.1 and 1.2 in HH and JJ subgenome, respectively. These values are similar to those previously published for in *O. sativa*^35,36^. This evidence confirms previous findings from El Baidouri and Panaud, who showed that the ratio of solo-LTRs to complete LTR-RTs does not depend on genome type^36^.

In summary, our results indicate that the difference in subgenome size in the HHJJ genome species is primarily due to the preferential accumulation of LTR-RT-related sequences in HH subgenome. Distribution of the main *Oryza*-specific TE families revealed no preferential expansion of specific families, and the solo-LTRs to complete LTR-RTs ratios showed no evidence for differential efficiency in TE removal, thereby favoring TE accumulation as the primary mechanism contributing to subgenome size disparity.

### Macro-synteny and large-scale chromosomal rearrangements

To understand and visualize the syntenic relationships across the entire *Oryza* genus, we built the first syntenic map of the *Oryza* genus that includes 21 *Oryza* species (i.e. nine tetraploid and two diploid species from this study, plus 10 publicly available *Oryza* diploid species [including the IRGSP RefSeq]), and the outgroup species *L. perrieri* (Supplementary Table 5). The synteny map (shown as riparian plot) tracks the syntenic orthologous blocks across the ten genome types presented here (i.e. AA, BB, CC, DD, EE, KK, LL, HH, JJ, GG), showing ∼15 million years of evolution in inversions, duplications, and translocations across the genus (Figure 1). The riparian plot in Supplementary Figure 3 shows collinear syntenic blocks inverted in consecutive (sub)genome pairs (i.e. shown as blue ribbons). The small-scale segmental duplication^37^ on chromosome 11 and 12 is shared by the *Oryza* species and is already present in *L. perrieri*. The *O. alta* (CCDD) and *O. grandiglumis* (CCDD) genomes share five unbalanced translocations relative to the *O. sativa* genome (e.g. t(Chr1CC; Chr3), t(Chr6CC; Chr1), t(Chr7CC; Chr4); t(Chr3DD; Chr6); t(Chr4DD; Chr7); Supplementary Figure 4). None of these translocations were found in the CCDD species *O. latifolia*, or in any other species included in this study (Figure 1). Simple sequence repeats (SSR) were found in most of the putative translocation breakpoints when comparing the *O. alta* and *O. grandiglumis* genomes to an *O. sativa* reference sequence – i.e. AT repeats (Supplementary Figure 5 and Supplementary Figure 6).

Reciprocal translocations between homoeologous chromosomes in polyploid genomes can be found by aligning subgenomes to each diploid relative genome species. When aligning the BBCC genome species to their diploid relative genome species (i.e. *O. punctata* (BB) and *O. officinalis* (CC)), a reciprocal translocation between Chr1BB and Chr1CC (i.e. t(Chr1BB; Chr1CC) was found (and confirmed with optical maps) in both *O. minuta* (∼9 Mb translocation size) and *O. malampuzhaensis* (∼8 Mb translocation size) (Supplementary Figure 7).

### The syntenic pangene

To identify core gene sets conserved during *Oryza* evolution, and accessory gene sets that underwent duplication, translocation and/or gene loss, we performed a micro-synteny analysis at the (sub)genome level (Figure 3). Using 832,658 *Oryza* gene sequences, we identified 77,482 syntenic gene clusters in the 30 (sub)genome species dataset used for GENESPACE^38^ analysis (Figure 3a). In *O. alta* and *O. grandiglumis*, as a consequence of chromosomal duplications and unbalanced translocations (described above), underlying genes were also duplicated and translocated, replacing genes on the chromosomal portions that were lost. In Figure 3a, these genes correspond to yellow (i.e. duplicated genes) and grey (i.e. depleted genes) tracks belonging to the same clusters of the dendrogram in either subgenome of *O. alta* and *O. grandiglumis* (Supplementary Table 6). Congruent with the random occurrence of translocations, we could not detect over-represented GO-slim terms when comparing *O. sativa* homologs of either *O. alta* and *O. grandiglumis* genes duplicated in the CC (DD) subgenome and depleted in the DD (CC) subgenome to *O. sativa* homologs of genes in CC (DD) subgenome.

**Figure 3.**
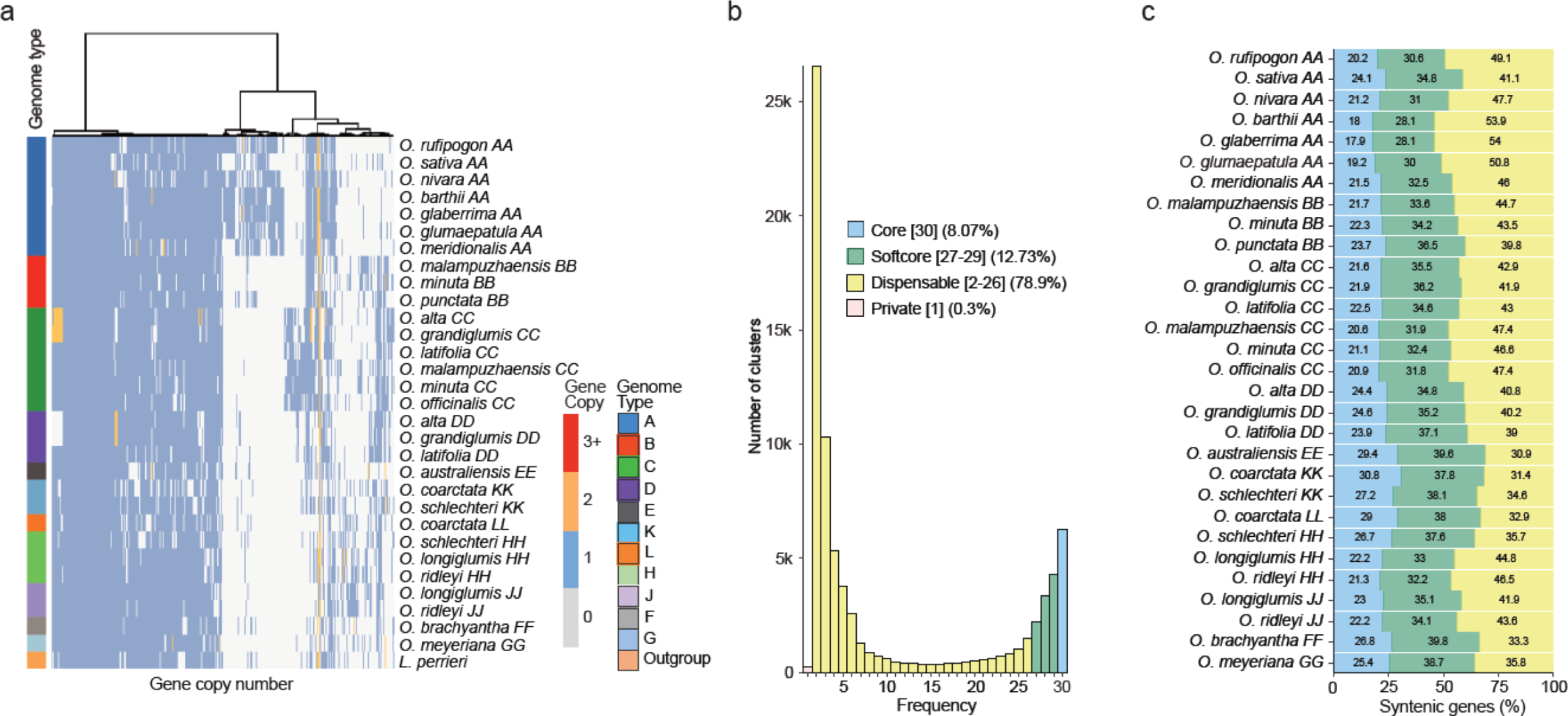
The syntenic pangene. **a)** Phylogenomic synteny profiling (copy-number profiling of micro-synteny gene clusters across a phylogeny) of clusters (size ≥ 2; 0: gene absence; 1: one gene copy; 2: two gene copies; 3+: three or more gene copies) in the *Oryza* genus (i.e. 30 *Oryza* (sub)genomes and the outgroup species). **b)** Histogram shows the frequency distribution of clusters shown in a) shared by different numbers of *Oryza* (sub)genomes. The legend shows the percentage of core cluster (i.e. found in all 30 species), softcore clusters (i.e. found in 27 to 29 species), dispensable clusters (i.e. found in 2 to 26 species), and private clusters. **c)** Percentage of clusters (i.e. core, softcore, and dispensable clusters) in the 30 *Oryza* genomes. Private clusters are not shown.

Syntenic core, softcore, dispensable, and private genes were defined as those present in all 30 (sub)genomes, in 27 to 29 (sub)genomes (≥ 90%), in 2 to 26 (sub)genomes, and in 1 (sub)genome, respectively. We divided clusters into 6,256 (8.1%), 9,865 (12.7%), 61,130 (78.9%), and 231 (0.3%) core, softcore, dispensable, and private syntenic gene families, respectively (Figure 3b). Across (sub)genomes, the number of gene families ranged between 18.0 to 30.8% for core syntenic genes, between 28.1 to 39.8% for softcore syntenic genes, between 31.0 and 54.0% for dispensable syntenic genes, and up to 0.2% for private genes (Figure 3c).

### Reconstruction of the evolutionary history of the *Oryza* chloroplast and nuclear (sub)genomes

To gain insight into the evolutionary history of the genus *Oryza* and their maternal origins, we first reconstructed a chloroplast genome-based phylogenetic tree using the chloroplast sequences of 26 *Oryza* species (i.e. the 10 chloroplast genomes assembled in this study and 16 chloroplast genomes obtained from NCBI), and the outgroup *L. japonica* (Supplementary Table 7). The phylogenetic tree of the *Oryza* genus presented here consists of two main clades and eight distinct monophylies, i.e. AA, BB/BBCC, CC/CCDD, EE, KKLL/HHKK, HHJJ, FF and GG genome types (Figure 4a). This tree is highly consistent with previously reported whole chloroplast genome-based trees^39,40^. Of note, the former study did not include the KKLL and HHKK genomes, while the latter merged the KKLL and HHKK genome types into a single HHKK type. Analogously to the tree shown by Zhang and colleagues^40^, our chloroplast-based tree shows that the HHJJ and HHKK genome types do not form monophyletic clades, suggesting the paternal donor was derived from a HH genome species. Accordingly, the KKLL and HHKK types formed a monophyletic clade in our tree, supporting that the maternal donor was likely a KK genome species. A longer branch length of the KKLL/HHKK clade, with respect to other clades (e.g., BB/BBCC or CC/CCDD), might suggest a different maternal origin of the KK types that hybridized with the LL and HH genome species (Figure 4a).

**Figure 4.**
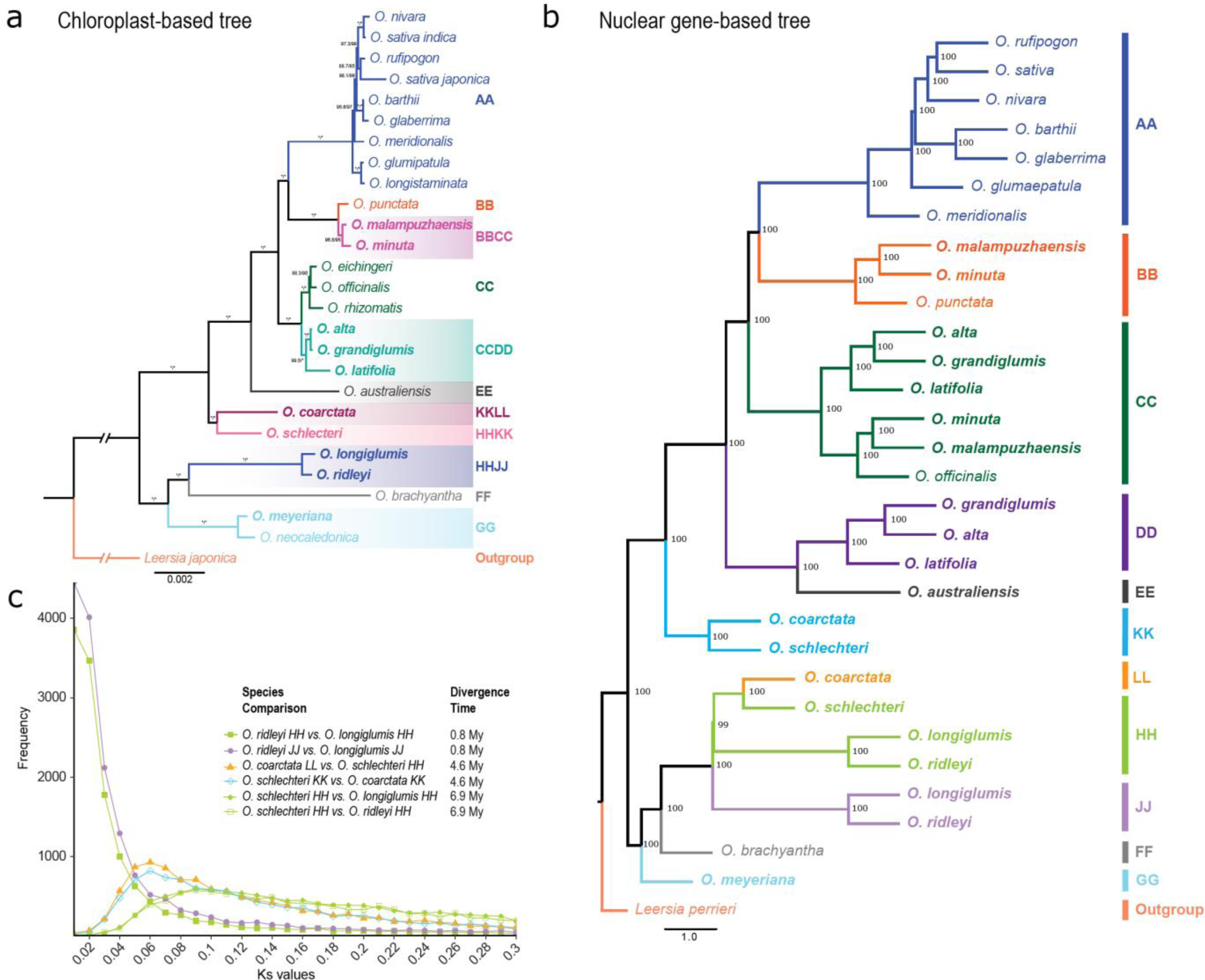
Phylogenetic analysis of the *Oryza* (sub)genomes. **a)** Phylogenetic tree based on chloroplast genome sequences. Maximum Likelihood phylogeny reconstruction was done using IQ-TREE based on the large-single copy (LSC) regions of the chloroplast genomes from 26 *Oryza* species. Supporting values given next to each branch denote are SH-aLRT (Shimodaira-Hasegawa-like approximate likelihood ratio) support (%) / ultrafast bootstrap support (%), respectively. Branch length indicates substitutions per site. Trees were rooted using *Leersia japonica* used as the outgroup. Chloroplast genome sequences from the additional diploid species shown here were downloaded from NCBI, see Supplementary Table 7. **b)** Phylogenetic tree based on nuclear genome sequences. Tetraploid genomes were phased into sub-genomes and nuclear orthologous groups were called by synteny based phylogenomic profiling. The tree is an Astral coalescent tree based on 3,728 nuclear syntenic genes. Branch lengths (in coalescent units) indicate the amount of discordance in the gene trees. Branch support values measure the support for a quadripartition. Proteome sequences from the additional diploid species shown here were downloaded from NCBI, see Supplementary Table 5. **c)** Ks frequency plot for HH, JJ, KK, and LL subgenomes in the *ridleyi* complex and the species in the unclassified group (*O. ridleyi* JJ vs. *O. longiglumis* JJ - blue; *O. ridleyi* HH vs. *O. longiglumis* HH - purple; *O. schlechteri* HH vs. *O. coarctata* LL - light green; *O. schlechteri* KK vs. *O. coarctata* KK - dark green; *O. schlechteri* HH vs. *O. longiglumis* HH - pink; *O. schlechteri* HH vs. *O. ridleyi* HH - magenta). Divergence time in Million years (My) is shown in the legend for each species comparison.

To better understand the evolutionary relationships of 21 of the 27 *Oryza* species for which a chromosome-level assembly was available, we performed coalescent phylogenetic analyses using 3,728 syntenic genes, which grouped the *Oryza* (sub)genomes into eight highly supported monophyletic clades – i.e. AA, BB, CC, DD/EE, KK, LL/HH/JJ, FF and GG (Figure 4b). The diploid CC genome (*O. officinalis*) clustered with the CC subgenomes of *O. minuta* (BBCC) and *O. malampuzhaensis* (BBCC), but not with the CC subgenomes of *O. alta* (CCDD), *O. grandiglumis* (CCDD), and *O. latifolia* (CCDD). This confirms previous studies^41,42^ suggesting that a CC genome species (i.e. likely *O. officinalis* (CC)) is the paternal donor of the BBCC tetraploid species, and the maternal donor is a BB genome species (i.e. likely *O. punctata* (BB), Figure 4a); while the CC genome in the CCDD tetraploid species served as the maternal parent and might be different from *O. officinalis* (Figure 4b).

Analysis of the species in the unclassified group^43^ (i.e. *O. coarctata* (KKLL) and *O. schlechteri* (HHKK)) shows that the LL genome of *O. coarctata* (KKLL) clusters with HH genome of *O. schlechteri* (HHKK), and the latter did not form a monophyletic clade with the HH genomes of HHJJ species (i.e. *O. ridleyi* and *O. longiglumis*) (Figure 4b). This again suggests the species with HHJJ and HHKK genomes had a HH genome as the paternal donor, but also that HH genomes of HHJJ and HHKK species might not be related. To investigate this incongruity, we calculated the synonymous substitution per synonymous site (Ks) values in these genome types (Figure 4c) to estimate the divergence time in pairs of subgenomes with similar Ks distributions using the neutral mutation rate of grasses^44^. The estimated divergence time between the HH genome of *O. schlechteri* (HHKK) and each of the HH genomes in the HHJJ species was ∼6.9 Mya; while the divergence time between the HH genomes of *O. ridleyi* (HHJJ) and *O. longiglumis* (HHJJ) was ∼0.8 Mya. The divergence time between the HH genome of *O. schlechteri* (HHKK) and the LL genome of *O. coarctata* (KKLL) was ∼4.6 Mya, and the same estimation was calculated between the KK genomes in these two species.

Our multi-gene coalescent analysis corroborates results from Ge and colleagues^45^ (i.e. phylogenies of *Adh1* and *Adh2* nuclear genes), and from Zhang and colleagues^40^ (i.e. a phylogeny based on NP78 and R22 nuclear genes), who proposed that *O. coarctata* and *O. schlechteri* belonged to the same genome type. As previously suggested by Lu and colleagues for *O. coarctata*^6^ (previously classified as HHKK), we propose to change genome type designation *O. schlechteri* to KKLL genome type to more accurately reflect the phylogenetic placement of this species in the genus.

### Homoeologous gene fractionation

Following whole genome duplication (i.e. *via* allopolyploidization - the hybridization of two or more distinct species; or *via* autopolyploidization - the multiplication of a complete chromosome set within a species), gene copies can be lost from one homoeologous chromosome or the other(s), a process defined as gene fractionation. Over evolutionary time, gene fractionation leads to reduction of a polyploid genome back to a diploid state where the overall genomic structure has changed substantially^46^. Gene fractionation (Figure 5a) was measured as a percentage of gene retention in the subgenomes of the tetraploid *Oryza* genomes. Red dashed lines represent the average percentage of gene retention calculated genome wide for each species: a lower percentage of gene retention with respect to the genome-wide average indicates over-fractionation (i.e. higher gene loss), while a higher percentage indicates under-fractionation (i.e. higher gene retention). Statistical comparisons (i.e. two-sided Wilcoxon rank sum test) of gene retention between subgenomes showed the most pronounced difference within *O. longiglumis/O. ridleyi* with higher gene retention in the HH subgenome, followed by the species in the *officinalis* complex with higher gene retention in the CC subgenome (p-value < 0.001; Figure 5a).

**Figure 5.**
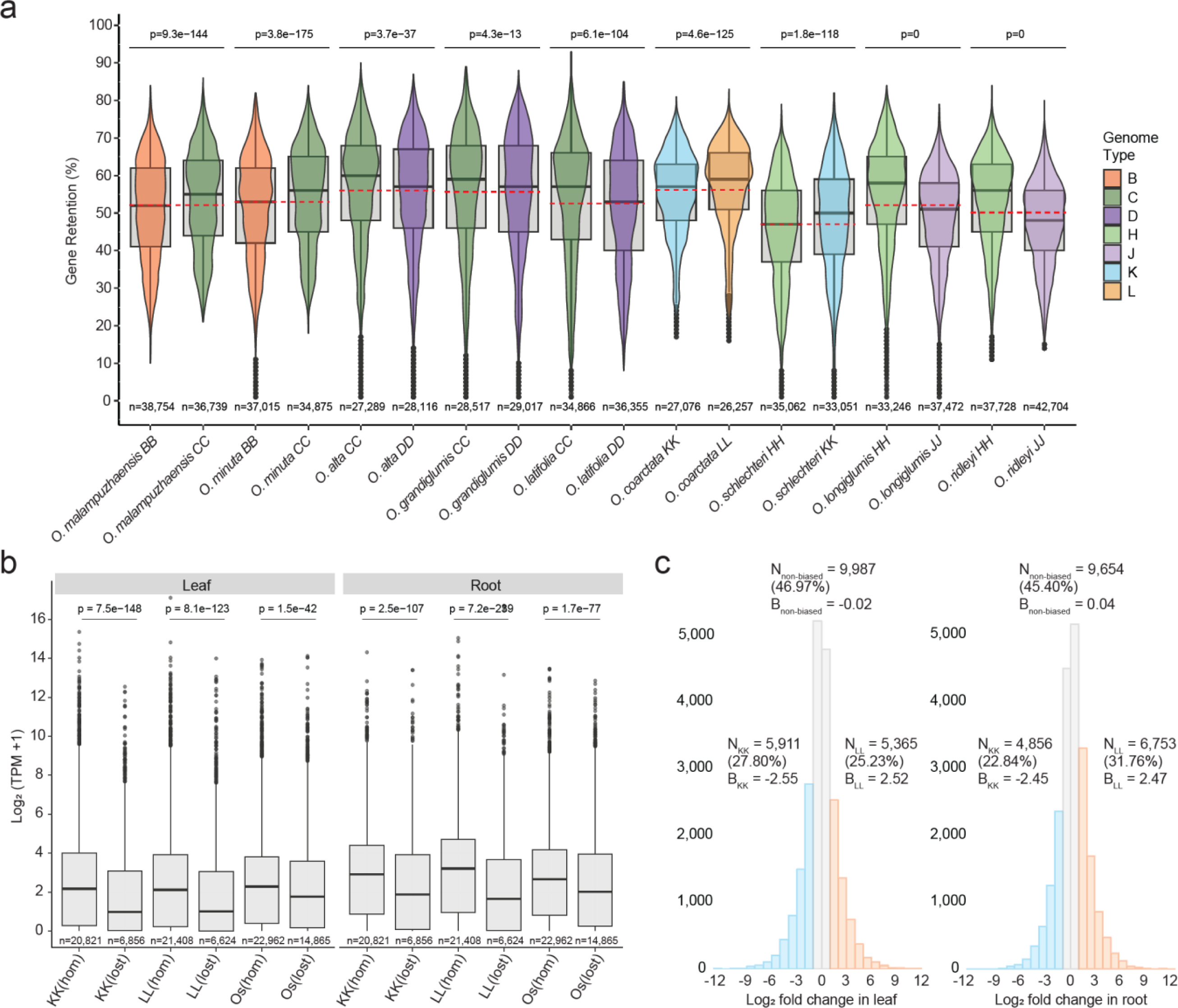
Homoeologous gene retention in *Oryza* and subgenome dominance in *O. coarctata*. **a)** Distribution of gene retention (in percentage, y-axis) in the subgenomes of the tetraploid species (x-axis). Each genome type is colored according to color codes as in Figure 3a and Figure 4b. Red dashed line is the average percentage of gene retention calculated genome-wide in each species. P-values (p) from two-sided Wilcoxon rank-sum test and number of sliding windows (n) are shown. **b)** Transcript abundance of homoeologous genes in *O. coarctata* and their homologs in *O. sativa*. Gene expression as log_2_(TPM + 1) is measured in the leaf and in the root considering all the replicates together. P-values (p) from two-sided Wilcoxon rank-sum test are shown. **c)** Homoeologous gene pair expression bias (B) in the leaf (left) and the root (right) of *O. coarctata*. Blue and orange bars represent the expression of homoeologs biased toward KK (B < -1) and LL (B > 1) subgenomes, respectively. Homoeolog pairs with -1 ≤ B ≤ 1 (grey bars) are defined as non-dominantly expressed. N represents the number of homoeologous gene pairs in the three categories (N_KK_: homoeologous gene dominantly expressed in KK subgenome; N_LL_: homoeologous gene dominantly expressed in LL subgenome; N_non-biased_: homoeologous gene not dominantly expressed). B_KK_, B_LL_ and B_non-biased_ represent average expression bias for the homoelogous pairs in the respective category.

### Gene fractionation and subgenome dominance/equivalence in *O. coarctata*

Subgenome dominance is a phenomenon where, in some polyploid species, genes from one subgenome tend to be expressed at higher levels than those from the homoeologous subgenome. It has been shown that, over evolutionary time, the less expressed subgenome (i.e. the submissive subgenome) tends to lose more homoeolog copies than the higher expressed subgenome, generating biased fractionation^47,46^. Alternatively, subgenome equivalence is where neither genome is “dominant” over the other, as demonstrated in banana and soybean^47^, and, recently, in broomcorn millet^48^.

To investigate these phenomena in the *Oryza* tetraploids, we analyzed gene expression patterns in *O. coarctata* (KKLL), due to its importance as a halophyte species^20^ and the availability of transcriptome data. The average gene retention in *O. coarctata* was 56.1%, meaning that, on average, ∼56% of genes are retained in duplicate and are syntenic in the homoeologous chromosomes. The gene fractionation distribution across homoeologous chromosomes is shown in Supplementary Figure 12. Overall, the KK subgenome showed over fractionation (i.e. a higher gene loss) with respect to the LL subgenome (p-value < 0.001, Wilcoxon rank sum test; Figure 5a; Supplementary Table 8).

We then analyzed transcript abundance of homoeologous gene pairs in *O. coarctata* and compared it to transcript abundance of homologous genes in *O. sativa*, in leaf and root tissues (Figure 5b). The median gene expression in *O. coarctata* was significantly higher (p-value < 0.001, Wilcoxon rank sum test) for homoeologous gene pairs than for genes homologous to *O. sativa* present in single copy (Figure 5b, Supplementary Table 9). In leaf, median values of log_2_(TPM + 1) were 2.17 and 0.98 for KK(hom) and KK(lost), respectively; and 2.11 and 1.01 for LL(hom) and LL(lost), respectively; in root, median values of log_2_(TPM + 1) were 2.91 and 1.88 for KK(hom) and KK(lost), respectively; and 3.20 and 1.66 for LL(hom) and LL(lost), respectively. Os(hom) showed significantly higher expression levels (p-value < 0.001, Wilcoxon rank sum test) than Os(lost) (Figure 5b, Supplementary Table 9): in leaf, median values of log_2_(TPM + 1) were 2.28 and 1.76 for Os(hom) and Os(lost), respectively; in root, 2.67 and 2.02 for Os(hom) and Os(lost), respectively. This evidence suggests that, in *O. coarctata*, genes with two homoeologous copies tend to be expressed at higher levels than genes that lost one homoeologous copy. Moreover, *O. sativa* homologs of KK(hom) and LL(hom) tend to be expressed at higher levels than *O. sativa* genes with no homologs in *O. coarctata*, and *O. sativa* homologs of KK(lost) and LL(lost).

We compared expression in homoeologous gene pairs to investigate whether there is subgenome dominance in the tissue-specific transcriptome data of *O. coarctata*. We quantified expression bias as log_2_-fold change in 21,263 homoeologous gene pairs: in leaf, 5,911 homoeologs (27.8%) were dominantly expressed in KK subgenome and 5,365 (25.2%) in LL subgenome, 9,987 homoeologs (47.0%) were non-dominantly expressed; in root, 4,865 homoeologs (22.8%) were dominantly expressed in KK subgenome and 6,753 (31.8%) in LL subgenome, 9,654 homoeologs (45.4%) were non-dominantly expressed (Figure 5c, Supplementary Table 10). Our analyses showed that, in *O. coarctata*, there is higher gene retention in LL subgenome (suggesting biased gene fractionation) and a fraction of homoeologous genes that are expressed at higher levels in the KK subgenome in the leaf and in the LL subgenome in the root, respectively (Binomial test, p-value < 0.001), suggesting subgenome expression equivalence^47^. Additional tissues need to be analyzed to support evidence of subgenome equivalence in *O. coarctata*.

## Discussion

We generated a comprehensive resource of publicly available wild diploid and tetraploid *Oryza* reference genomes spanning all tetraploid genome types and the EE and GG diploid genome types, using PacBio long-read sequencing and optical map validation. Previous efforts in this direction were made within the framework of the International *Oryza* Map Alignment Project (I*O*MAP^49^), with the establishment of a genus-wide framework of bacterial artificial chromosome (BAC) fingerprint, end-sequenced physical maps, and Illumina short-read sequencing of the *Oryza* species^31,50,51^.

The analysis of our dataset gained noteworthy insights into genome evolution within the genus. We retested the phylogenetic placement of the tetraploid genomes confirming previous findings^45,5,51,39,42^. Our whole-genome scale phylogenetic analysis provided robust confirmation of previous inference (based on the analysis of a few genes) on the origin of the HH, KK, and JJ genome types, where *O. schlechteri* and *O. coarctata* are more closely related to each other and share the same genome type^45^. Subgenome size, analysis of TE content, and proportion of core, softcore and dispensable genes in our high-quality and chromosome-level genomes combined provided strong evidence of the high similarity between the HH subgenome of *O. schlechteri* and LL subgenome of *O. coarctata*, as compared to the HH subgenome of *O. ridleyi*/*O. longiglumis*. This evidence provides further support to previous findings in *O. coarctata*^6^, where the authors proposed to rename this species (previously designated as a HHKK genome type) as a KKLL genome type. We therefore propose to designate *O. schlechteri* to the same genome type as *O. coarctata*, i.e. KKLL genome type^6^. Based on the new designation, the resulting phylogenetic tree is shown in Figure 6.

**Figure 6.**
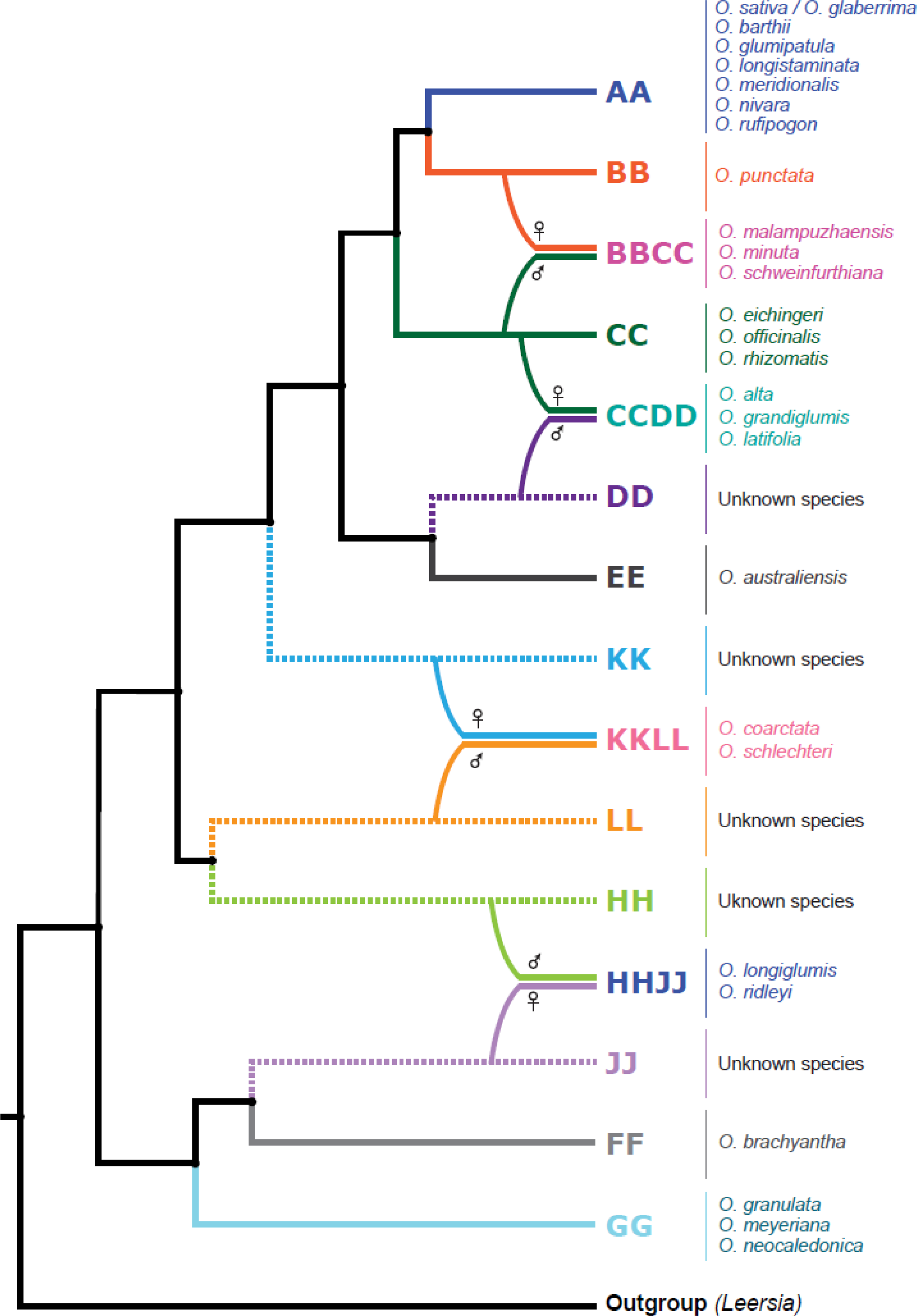
Newly proposed phylogenetic tree illustrating the origins and evolutionary history of diploid and tetraploid *Oryza* species. Tree topology, maternal and parental donors were inferred based on our whole-chloroplast genome tree (Figure 4a) and multi nuclear gene subgenome tree (Figure 4b). *Leersia perrieri* and *L. japonica* (here collectively referred to as *Leersia*) were used as outgroup. Single-line branches denote diploid species, while double-line branches denote tetraploid species. Single dashed lines represent unknown diploid species. Genome types and known representative species are shown next to the terminal nodes. The tree includes the new designation of *O. schlechteri* as KKLL genome type (the same as *O. coarctata*) proposed in this work. Relative times of the four hybridization events were based on previous studies^27^.

We generated the first macro-synteny description of the *Oryza* (sub)genomes, and characterized large chromosomal rearrangements that resulted in our present-day inventory of living *Oryza* species. For example, the description of five large non-reciprocal translocations shared between *O. alta* (CCDD) and *O. grandiglumis* (CCDD) (but not present in *O. latifolia* (CCDD)) added robust evidence to previous hypothesis of their conspecific nature^52,40^. The CCDD genome species were originally classified based on morphological differences^53^; however, intermediate forms between *O. alta* and *O. grandiglumis* were observed^54,55^. Furthermore, *O. alta* is considered a synonym for *O. latifolia* in the Plants of the World Online (POWO) (https://powo.science.kew.org). Population genetics studies are needed to shed light on the history and composition of the CCDD species to resolve the dispute on whether they should be treated as separate species^56^.

We analyzed a quite interesting example of genome size variation between homoeologous genomes in the species of the *ridleyi* complex (i.e *O. ridleyi* and *O. longiglumis*, both HHJJ), and identified TEs as the driving force of genome size change in these species, thereby clarifying the mode of that amplification (i.e. the involvement of the entire TE complement and not just few TE families).

Our investigation of gene fractionation in the tetraploids found variable patterns of gene fractionation among subgenomes, as recently described in other plant systems^46^. Investigation of subgenome dominance/equivalence in leaf and root tissue of *O. coarctata* (KKLL) did not reveal evidence of expression dominance of one subgenome over another, even though gene fractionation was higher in subgenome KK. Additional transcriptome data for *O. coarctata* and the other tetraploid species will be needed to investigate the phenomenon of subgenome dominance/equivalence in the other *Oryza* species.

This dataset provides a valuable resource for future investigations, ranging from the discovery of adaptive genes/traits to improve cultivated rice, to the neo-domestication of the wild *Oryza* species (i.e. now that a successful neo-domestication protocol has been established for *O. alta*^22^), to population genetics of the wild *Oryza* species across their species ranges for conservation and enhancement of their genetic diversity for the planet’s future^56^.

## Methods

### Sample collection

Single seed descent (SSD) germplasm for *O. alta* (IRGC 105143), *O. australiensis* (IRGC 100882), *O. grandiglumis* (IRGC 105669), *O. latifolia* (IRGC 100890), *O. longiglumis* (IRGC 106525), *O. malampuzhaensis* (IRGC 80765), *O. meyeriana* (IRGC 106473), *O. minuta* (IRGC 101141), *O. ridleyi* (IRGC 100821) and *O. schlechteri* (IRGC 82047) were obtained from the International Rice Research Institute (IRRI, Philippines) under the Standard Material Transfer Agreement. Seeds were sown in potting soil and grown under 24-29°C air temperature with 15-25% humidity in the greenhouse. *O. coarctata* (IRGC 104502) leaf tissue was obtained from a vegetative voucher plant imported from IRRI through the USDA and grown under 24-29°C air temperature with 15-25% humidity in the greenhouse at the University of Arizona.

### High molecular weight (HMW) DNA isolation

Approximately 6 grams of young-stage leaf tissue was harvested from 5 to 6-week old seedlings from each accession, rinsed with sterile milli-Q water, immediately frozen in liquid nitrogen, and stored at -80°C prior to the DNA extraction. After grinding leaves, HMW DNA was extracted following Luo and Wing^57^, with slight modifications. Ground leaf tissue was incubated in NIB buffer (10 mM Tris-HCl pH 8.0, 10 mM EDTA pH 8.0, 100 mM KCl, 0.5 M sucrose, 4 mM spermidine, 1 mM spermine) on ice for 15 minutes, and the resulting homogenate was then filtered using Miracloth filters (Merck KGaA, Germany). After adding Triton X-100 to the nuclei-containing solution (1:5 v/v), samples were kept on ice for 15 minutes. Samples were centrifuged, washed, re-suspended in NIB buffer (added with 1% Triton X-100), and centrifuged again. The supernatant fluid was discarded, 2% CTAB was added to the samples, and tubes were gently mixed on a shaker and incubated at 37°C for 30 minutes. After adding proteinase K (Qiagen, Hilden, Germany) to a final concentration of 1 mg/mL, samples were incubated at 50°C for 2 hours. Chloroform: isoamyl alcohol (24:1 v/v) was added to the samples. After centrifugation, the aqueous phase was transferred into a fresh tube, and 2/3 volume of isopropanol was added to precipitate the DNA. After centrifugation, the HMW DNA was washed with 70% ethanol and air-dried in the fume hood for 1-3 minutes. The pellet was gently re-suspended in 1× TE (Tris-HCl 10 mM pH 8.0) buffer overnight at room temperature. DNA concentration and quality were measured using Qubit Fluorometric Quantification (Invitrogen, Thermo Fisher Scientific, MA, US) and Nanodrop (Thermo Fisher Scientific, MA, US), respectively. DNA size and susceptibility to molecular reagents (i.e. *Eco*RI and *Hind*III) were checked on pulsed-field gel electrophoresis.

### HMW DNA library preparation for PacBio sequencing

For Continuous Long Read (CLR) sequencing technology (PacBio, CA, US), 20 µg of HMW DNA was sheared to an appropriate size range (∼25-100 kb) and SMRTbell Express Template Prep Kit 2.0 was used to prepare the library. The final library was size selected on a BluePippin electrophoresis system (Sage Science, MA, US) using U1 markers with size selection >30 kb. The recovered final library was quantified with Qubit Fluorometric Quantification, and fragment size was checked on the Fragment Analyzer System (Agilent Technologies, CA, US). The library was prepared for sequencing with a PacBio Sequel II Sequencing kit 2.0 for CLR libraries, loaded onto 8M SMRT cells, and sequenced in the CLR mode on a Sequel II instrument for 15 hours at the Arizona Genomics Institute (University of Arizona, AZ, USA).

For Circular Consensus Sequencing (CCS) sequencing technology (PacBio, CA, US), HMW DNA was sheared to an appropriate size range (∼10-30 kb) using a Megaruptor 3 (Diagenode, USA). The sequencing library was constructed using the SMRTbell Express Template Prep Kit 2.0 following manufacturer’s protocols. The final library was size selected on a BluePippin electrophoresis system using S1 markers with a 10-25 kb size selection. The recovered size-selected library was quantified by Qubit Fluorometric Quantification, and size checked on a Femto Pulse System (Agilent Technologies, CA, US). The library was prepared for sequencing with a PacBio Sequel II Sequencing kit 2.0 for HiFi libraries, loaded onto 8M SMRT cells, and sequenced in the CCS mode on a Sequel II instrument for 30 hours at the Arizona Genomics Institute.

### Short- and long-read RNA library preparation and sequencing

Transcriptome data was generated from the same *Oryza* accessions that were used for genome sequencing. Illumina short-read sequencing (RNA-Seq) and PacBio long-read sequencing (Iso-Seq) for transcriptome assembly were performed using RNA isolated from leaf tissue of all species, and from panicle tissue from *O. alta*, *O. coarctata*, *O. grandiglumis*, *O. latifolia*, and *O. ridleyi*. Additional RNA-Seq data for *O. coarctata* was generated to measure expression levels of homoeologous genes in leaf and root tissues (see section “Expression of homoeologous genes in *O. coarctata*”). For this purpose, *O. coarctata* cuttings were grown in three separated plant growth tanks with six replicates each, in hydroponic conditions with freshwater irrigation at the KAUST greenhouse facility. The third leaf of each plant was collected for RNA isolation. Upon collection, tissues were immediately frozen in liquid nitrogen and stored at -80°C prior to total RNA isolation with NucleoSpin RNA Plant and Fungi kit (Macherey-Nagel Inc., PA, US). RNA libraries for RNA-Seq were prepared using the Illumina TruSeq Stranded Total RNA Ribo-Zero Plant kit (Illumina, CA, US), and sequenced on the Illumina platform in paired-end mode and 150 bp read-length (Psomagen, MD, US). RNA libraries for Iso-Seq were constructed using PacBio Iso-Seq Express Oligo kits following manufacturer’s instructions, loaded onto 8M SMRT cells, and sequenced in CCS mode on a Sequel II instrument for 24 hours at the Arizona Genomics Institute.

### Optical mapping

Optical mapping was performed by Corteva (Corteva Agriscience, IN, US). Ultra-HMW DNA (uHMW DNA) was isolated from agarose-embedded nuclei following the general procedure by Luo et Wing^57^ with some modifications. Approximately 2 to 4 g frozen leaf tissue was ground in liquid nitrogen and lysed in NIB/Triton-X100 buffer as described in the HMW DNA isolation section above. After centrifugation, the pelleted nuclei were recovered, resuspended, mixed with equal parts of 1% low-melting-point agarose equilibrated at 43°C, and poured into molds. The resulting agarose plugs with embedded intact nuclei were left to solidify at 4°C for 5 minutes and placed in 2.4 mL of lysis buffer (0.5M EDTA, 0.1% SDS and 15% (v/v) proteinase K. The plugs were incubated at 50°C with intermittent shaking at 300 rpm every 10 minutes. The step was repeated with a fresh lysis buffer after approximately 12 hours, for a total combined incubation time of 24-30 hours. After lysis, the buffer containing the plugs was cooled to room temperature, and RNase A (Puregene, Qiagen, Hilden, Germany) was added to a final concentration of 80 µg/mL. The solution was then incubated for 1 hour at 37°C with intermittent shaking at 300 rpm every 10 minutes. After the RNase treatment, the lysis buffer was removed and plugs were rinsed 3 times in T_10_E_10_ (10 mM Tris HCl, pH 7.5, 10 mM EDTA), followed by four T_10_E_10_ 10-minute washes with slow agitation. After an additional five 15-minute TE washes, the plugs were removed from the buffer, and melted at 70°C for 5 minutes. One unit of agarase (Thermo Fisher Scientific, MA, US) was added to each melted plug and incubated for 45 minutes at 43°C to the agarose. The eluted DNA solution was further purified by drop dialysis against TE buffer for 1 hour. HMW DNA was validated by viscosity and Qubit Fluorometric Quantification was used for quantification on 2 µl sonicated aliquots using the three-point method as per Bionano Genomics recommendations (Bionano Genomics, CA, US).

Optical genome mapping for each accession was performed using the Bionano Direct Label and Stain chemistry (DLS) and the Saphyr systems (Bionano Genomics). DNA was labeled and dyed using the DLS Kit and loaded into Bionano Chips (Bionano Genomics). Approximately 800 ng uHMW DNA was incubated for 3.45 hours at 37°C in the presence of the DLE-1 labeling enzyme, DL-Green dye and DLE-1 Buffer, followed by 1-hour incubation with proteinase K at 50°C. Molecule assembly and hybrid scaffolding were performed using the Bionano Access and Tools software v1.4.1, v1.5.2, and v1.6.1. The samples were assembled using the non-haplotype, no ES, no CMPR Cut parameters.

### Assembly and validation of 11 *Oryza* reference genome sequences

For the *de novo* assembly of PacBio CLR reads, Mecat2^58^ v20193.14, Canu^59^ v2.0, and Flye^60^ v2.8.1 software were run using default settings and the following parameter for genome size based on Mondal et al.^61^: *O. coarctata* and O. *schlechteri*: 800 Mb; *O. ridleyi*: 1,200 Mb; *O. alta*, *O. grandiglumis*, *O. latifolia*: 1,008 Mb; *O. minuta* and *O. malampuzhaensis*: 1,124 Mb; *O. longiglumis*: 1,330 Mb. Illumina sequence reads were used to polish contig-level assemblies using the algorithm *arrow* implemented in GCPP (Generate Highly Accurate Reference Contigs, https://github.com/PacificBiosciences/gcpp), and the software tool Pilon^62^ v1.23. To assign contigs to chromosomes, contigs belonging to the best assembly output were aligned to the chromosome sequence of publicly available diploid *Oryza* assemblies - i.e. the reference genome sequence of *O. sativa* v.g. japonica cv. Nipponbare (a.k.a. IRGSP RefSeq^63^, <u>GCF_001433935.1</u>), *O. punctata*^64^ (GenBank assembly accession: <u>GCA_000573905.2</u>), *O. officinalis*^42^ (GenBank assembly accession: <u>GCA_008326285.1</u>) and *O. brachyantha*^65^ (GenBank assembly accession: <u>GCF_000231095.2</u>) - using the software Mashmap^66^ v2.0. Contigs showing overlapping alignment on the reference sequences were considered homoeologous (i.e. orthologous contigs between two subgenomes). Once the two sets of homoeologous contigs were determined, a phylogenetic analysis was used to identify which contigs belonged to their respective subgenome. Homology Aware K-mers (HAKMER^67^) and Phylogenetic Analysis using PAUP^68^ v4.0a software were used to build phylogenetic trees using the already available genome sequences of *O. sativa* (AA), *O. punctata* (BB), *O. officinalis* (CC), and *O. brachyantha* (FF), and the *de novo* genome assembly of *O. australiensis* (EE). The rationale was that contigs belonging to the same genome type are expected to form a clade in the expected evolutionary relationship with the diploid *Oryza* species (e.g. contigs belonging to BB subgenome in *O. malampuzhanesis* (BBCC) and *O. minuta* (BBCC) are expected to cluster in the same clade with *O. punctata* (BB)). The ordering and orientation of contigs on a chromosome were accomplished by aligning their sequence to reference genome sequence(s) with Genome Puzzle Master^69^, and/or RagTag^70^ v2.1.0. The assembly with the highest assembly metrics was referred to as the primary assembly. Contigs from secondary assemblies overlapping two or more contigs in the primary assembly were used to close gaps through GPM or RagTag *patch* function. For the *de novo* assembly of PacBio HiFi reads, the software Hifiasm^71^ v0.19.5 and HiCanu^72^ were used with default settings. Our strategy for scaffolding, contig assignment to chromosomes and subgenomes, contig ordering and orientation on the chromosome, and gap fixing was the same as described above.

Validation of the scaffolding was performed by anchoring each assembly to its respective Bionano optical maps (Corteva, IN, US). Optical map anchors spanning gaps in the primary assembly allowed us to estimate gap sizes between scaffolds. We evaluated the completeness of each nuclear genome assembly by Benchmarking Universal Single-Copy Orthologs (BUSCO^32^ v5.1.2), using the *poales_odb10* database (4,896 genes) and the assessment mode *genome*. For chloroplast genome assembly, contigs from each raw assembly were aligned against the chloroplast sequence of *O. sativa*^73^ using Blastn^74^ for local alignment of conserved sequences, and Minimap2^75^ v2.19 for global alignment of contigs to find those covering the entire reference sequence.

### Transposable element (TE) annotation

Extensive De-novo TE Annotator (EDTA^76^ v1.9.0) was used to generate a *de novo* non-redundant TE library for each genome, using default settings. *De novo* TE libraries were then used to annotate TEs in each respective genome using EDTA, and to soft-mask genome sequences using RepeatMasker^77^ v4.1.0 for the following step of gene prediction.

### Gene prediction

After evaluating read quality with FastQC (v0.11.8) and removing adapters with Trimmomatic^78^, RNA-Seq reads were aligned to their respective genomes using STAR^79^ aligner embedded in the tool OmicsBox v3.2 (Bioinformatics Made Easy, BioBam Bioinformatics, https://www.biobam.com/omicsbox, 2019). GMAP^80^ aligner implemented in OmicsBox was used to align high-quality transcripts from Iso-Seq reads. *Ab-initio* gene prediction was carried out on soft-masked genomes using Augustus^81^ software embedded in Omicsbox using a model training set derived for *O. sativa*, for all species except *O. australiensis*.

For *O. australiensis*, MAKER-P^82^ v3.10.02 was used to carry out gene prediction using RNA-Seq from leaf tissue. MAKER-P was run on the soft-masked genome of *O. australiensis* with Augustus^81^, SNAP^83^ and Fgenesh+^84^ gene predictors. Genes and transcripts were retained if the annotation edit distance (AED) was lower than 1.

For the macro-synteny and the phylogenetic analysis, gene models in each genome species were filtered to retain the longest isoform using AGAT toolsuite (https://agat.readthedocs.io/en/latest/index.html).

### Functional gene annotation

Gene function annotation was achieved by retaining the best match for each predicted protein sequence aligned against the NCBI^85^ non-redundant (nr) and NCBI Reference Sequence (RefSeq) protein databases v5 (http://www.ncbi.nlm.nih.gov/RefSeq/) using Diamond Blastp^86^ v0.9. Protein motifs and domains were annotated using InterProScan^87^ v5.39 with default settings. Functional information obtained from InterProScan and Blast was then translated into Gene Ontology (GO) terms using the Blast2GO^88^ suite. Gene function annotation in *O. australiensis* was carried out using MAKER-P^82^ v3.10.02 and a publicly available non-redundant database of protein and mRNA sequences of *Oryza*^51^. Lastly, resulting gene sets in each genome species were checked for completeness using the *poales_odb10* database in BUSCO^32^ v5.1.2 and the assessment mode *proteins*.

### Gene Ontology (GO) Enrichment analysis

GO enrichment analysis was performed using PANTHER v18.0 (https://www.pantherdb.org/). PANTHER GO-Slim annotations for each ontology (i.e. Molecular Function, Biological Process, and Cellular Component) were assigned to all genes using the Fisher’s exact test and *fdr* corrected p-values.

### TE amplification and insertion time of long terminal repeat-retrotransposons (LTR-RTs) in the subgenomes of *O. ridleyi* and *O. longiglumis*

Conserved tracts of 100 amino acids in length of transposase (TP) and reverse transcriptase (RT) enzymes were used as queries in tBlastn^74^ to identify DNA TEs and retroelements, respectively, in the subgenomes of *O. ridleyi* and *O. longiglumis*. Five hundred paralogs covering at least 80% of the query length were randomly selected among the tBlastn output hits for six TE superfamilies (i.e. Ty1/Copia and Ty3/Gypsy LTR retroelements, LINE and CACTA DNA TEs, MuDR and hAT) and aligned on their respective subgenomes using MUSCLE^89^ v3.8.425. The multiple sequence alignments of each TE superfamily were then used to build a Neighbor-Joining tree using MegaX v10.1^90^. Bootstrap values were calculated in MegaX for 1,000 replicates using the pairwise deletion option and shown on the tree when greater than 50. Evolutionary distances were estimated using the Poisson correction distance^91^. We estimated the insertion time of the complete LTR-RTs using the method proposed by SanMiguel and colleagues^92^, which is based on the nucleotide distance between the two flanking LTR sequences of a complete LTR-RT. For each genome species, LTR sequences flanking complete LTR-RTs were aligned pairwise using the global aligner STRETCHER^93^. The nucleotide distance between the two LTR sequences was quantified using the Kimura 2p method^94^ as implemented in the DISTMAT software^93^. The nucleotide distance (D) was then converted to insertion time (T) in Million years (My) using the formula: 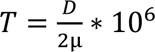; where μ 2μ is a substitution rate of 1.3×10^−8^ per site per year^95^.

### Macro- and micro-synteny analysis

To track the genomic dynamics across the genus *Oryza*, we performed a macro-synteny analysis using GENESPACE v1.3^38^. This software implements and improves features from OrthoFinder^96^ and MCScanX_h^97^ to link gene sequence homology with gene coordinates, and traces the process of genome polyploidization, reduction, rearrangement, and translocation across a set of genomes. The protein sequences of the nine tetraploid and the two diploid genomes reported in this study, as well as the publicly available protein coding genes of 10 diploid *Oryza* species and the outgroup *Leersia perrieri* (Supplementary Table 5) were analyzed in GENESPACE using default settings. The macro-synteny analysis was run separating each tetraploid genome into two individual subgenomes (i.e. the analysis was performed considering each genome type separately). The macro-synteny results were visualized as a riparian plot using the embedded *plot_riparian* function, and the information of homology and collinearity at the subgenome level was used to build a synteny-constrained phylogenomic framework of the *Oryza* (sub)genomes. Gene micro-synteny analysis across the *Oryza* genomes was performed using a previously developed pipeline^98^ with the modification that pairwise synteny was inferred by GENESPACE, and syntenet clusters were generated using custom developed scripts (https://github.com/xiaoyezao/SynNet-Py). Copy-number variation of syntenic orthologs (i.e. a group of genes derived from a single common ancestor and retained syntenic relationships) was profiled using the SYNTENET^99^ package, and visualized as heat maps. The syntenic orthologs were then used for multi-loci phylogenomic analysis to infer the phylogeny of the (sub)genomes as described in the next section.

### Phylogenetic relationship analysis of the chloroplast and nuclear genome across the *Oryza* genus

We performed a phylogenomic analysis using chloroplast-based and multi nuclear gene-based approach to infer the evolutionary history of the *Oryza* species/genomes.

For chloroplast-based phylogeny, we used the chloroplast genome sequences of 10 *Oryza* species (i.e. *O. malampuzhaensis*, *O. minuta*, *O. alta*, *O. grandiglumis*, *O. latifolia*, *O. coarctata*, *O. schlecteri*, *O. ridleyi*, *O. longiglumis*, and *O. meyeriana*) assembled from whole-genome PacBio sequencing data (i.e. this study), plus 17 publicly available chloroplast genome sequences from 16 diploid *Oryza* species and the outgroup *Leersia japonica* (Supplementary Table 7). To construct the maximum likelihood (ML) phylogenetic tree, the large-single copy (LSC) regions of the chloroplast genomes were aligned by MAFFT^100^ v7.480. Poorly aligned regions were trimmed in trimAL^101^ v1.4 with the option “*-automated1*”. The alignment files were subjected to IQ-TREE^102^ v1.6.12 with default settings (1,000 bootstrap iterations) and with the best-fit substitution model identified by ModelFinder^103^. The resulting tree was visualized in FigTree v1.4.3 (http://tree.bio.ed.ac.uk/software/figtree/) and rooted using *L. japonica* as the outgroup. Sequence alignments and machine-readable phylogenetic trees are provided in the Supplementary Dataset.

For nuclear gene-base phylogeny, the nuclear gene dataset was obtained from the GENESPACE^38^ output as described above. A total of 3,728 syntenic ortholog groups containing at least 30 genes were used for the phylogenomic analysis. Syntenic ortholog groups were aligned using MAFFT^100^ v7.520 (--*genafpair*; --*maxiterate* 1,000), and the alignments were cleaned using trimAl^101^ v1.4.1 (-*gt* 0.6; -*st* 0.001). The gene trees were inferred using RAxML-NG^104^ using the model *Q.plant*s^105^. All gene trees were then subjected to a coalescent algorithm in Astral-Pro2^106^ to infer the phylogeny at the (sub)genome level. Branch supports of the Astral tree were estimated as local posterior probabilities^107^.

### Ks analysis

SynMap^108^ and CodeML^109^ tools, implemented in the Comparative Genomics (CoGe^110^) platform v7 (https://genomevolution.org/coge), were used to identify collinear blocks of homologous genes between subgenomes of the same type (e.g. between the KK subgenomes of *O. coarctata* and *O. schlechteri*) and calculate the fraction of synonymous substitutions per synonymous site (Ks). The SynMap analysis was run using default paramenters (i.e. comparison algorithm: Last; window size: 100 genes; minimum number of aligned pairs: 5 genes; maximum distance between two matches: 20 genes). The distribution of Ks values ≤ 0.3 for each pair of subgenomes was plotted using a bin width of 0.01. The divergence time (T) of subgenomes in each pair was estimated using the Ks value at the peak of the distribution and the neutral mutation rate of grasses^44^ (i.e. μ = 6.5 x 10^-9^ mutations per site per year) using the formula T = Ks / 2μ.

### Gene fractionation

Gene fractionation was run on each tetraploid genome using SynMap^108^ and FractBias^111^, both implemented in CoGe^110^. SynMap was used to define gene homoeology and collinearity between subgenomes in each tetraploid genome as described above. FractBias was used to calculate and plot gene retention by setting a quota align ratio of 1 to 1.

### Expression of homoeologous genes in *O. coarctata*

To identify pairs of homoeologous genes (i.e. orthologous genes between KK and LL subgenomes) among the predicted genes in the *O. coarctata* genome, we used the reciprocal best hit (RBH) approach, where protein sequence of two genes residing on different subgenomes find each other as the best hit in the opposite subgenome. To generate RBHs of each gene, we used Blastp^86^ and an e-value cutoff of 1e-05. The same method was used to identify genes in *O. sativa* homologous to genes in *O. coarctata*. In *O. coarctata*, genes were divided in the following categories: KK(hom): genes in KK subgenome with a homoeologous copy on LL subgenome; KK(lost): genes in LL subgenome homologous to *O. sativa* but with no homoeologous copy in KK subgenome; LL(hom): genes in LL subgenome with a homoeologous copy on KK subgenome; LL(lost): genes in KK subgenome homologous to *O. sativa* but with no homoeologous in LL subgenome. We classified genes in *O. sativa* that are homologous to homoeologous gene pairs in *O. coarctata* as Osat(hom). *O. sativa* genes with no homologs in *O. coarctata*, and *O. sativa* genes homologous of homoeologs that lost one copy in *O. coarctata* were classified as Osat(lost).

As density distribution of log_2_-transformed raw read counts was homogenous in all samples across the three plant growth tanks (Supplementary Figure 11), we considered the 18 individual plants as technical replicates. Paired-end reads from all the plants were mapped on each individual subgenome sequence of *O. coarctata* using Tophat2^112^. For reads that mapped to both subgenome sequences, we used Explicit Alternative Genome Likelihood Evaluator (Eagle)-RC^113^ to determine the likelihood of read alignment against each subgenome without knowing the genotype differences explicitly (--*ngi*), and chose the best alignment. Expression level was determined using transcripts per kilobase of exon model per million mapped reads (TPM) with TPMCalculator^114^. To compare expression of genes in *O. coarctata* homologous to genes in *O. sativa*, we obtained publicly available datasets of Illumina paired-end RNA-Seq of flag leaf (i.e. <u>SAMN22452874</u>, <u>SRR16526865</u>, <u>SRR16526866</u>), leaf (i.e. <u>SRR4017523</u>, <u>SRR4017527</u>) and root (i.e. <u>SRR25078452</u>, <u>SRR25078455</u>, <u>SRR25078456</u>) of *O. sativa*. Sequencing data were mapped to the *O. sativa* IRGSP RefSeq using Tophat2^112^. TPM values were calculated as described above. Difference in expression levels in the six gene categories was visualized as log_2_(TPM+1), and p-values between “hom” and “lost” categories were obtained using a two-sided Wilcoxon rank-sum test.

Expression bias (B) was quantified in homoeologous gene pairs over leaf and root tissue using the log_2_-fold change:

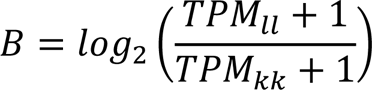

where *kk* and *ll* indicate the expression level (as TPM) in KK and LL subgenomes, respectively. Expression bias values greater than 1 (B > 1) indicates that the homoeologous copy on LL subgenome is dominantly expressed, while expression bias values lower than -1 (B < -1) indicates the homoeologous copy on KK subgenome is dominantly expressed^115^. The homoeologous gene pairs showing less than a two-fold change (-1 ≤ B ≤ 1) were classified as non-dominantly expressed. Binomial test was used to assess p-value of significance.

## Supplementary Figures

**Supplementary Figure 1.**
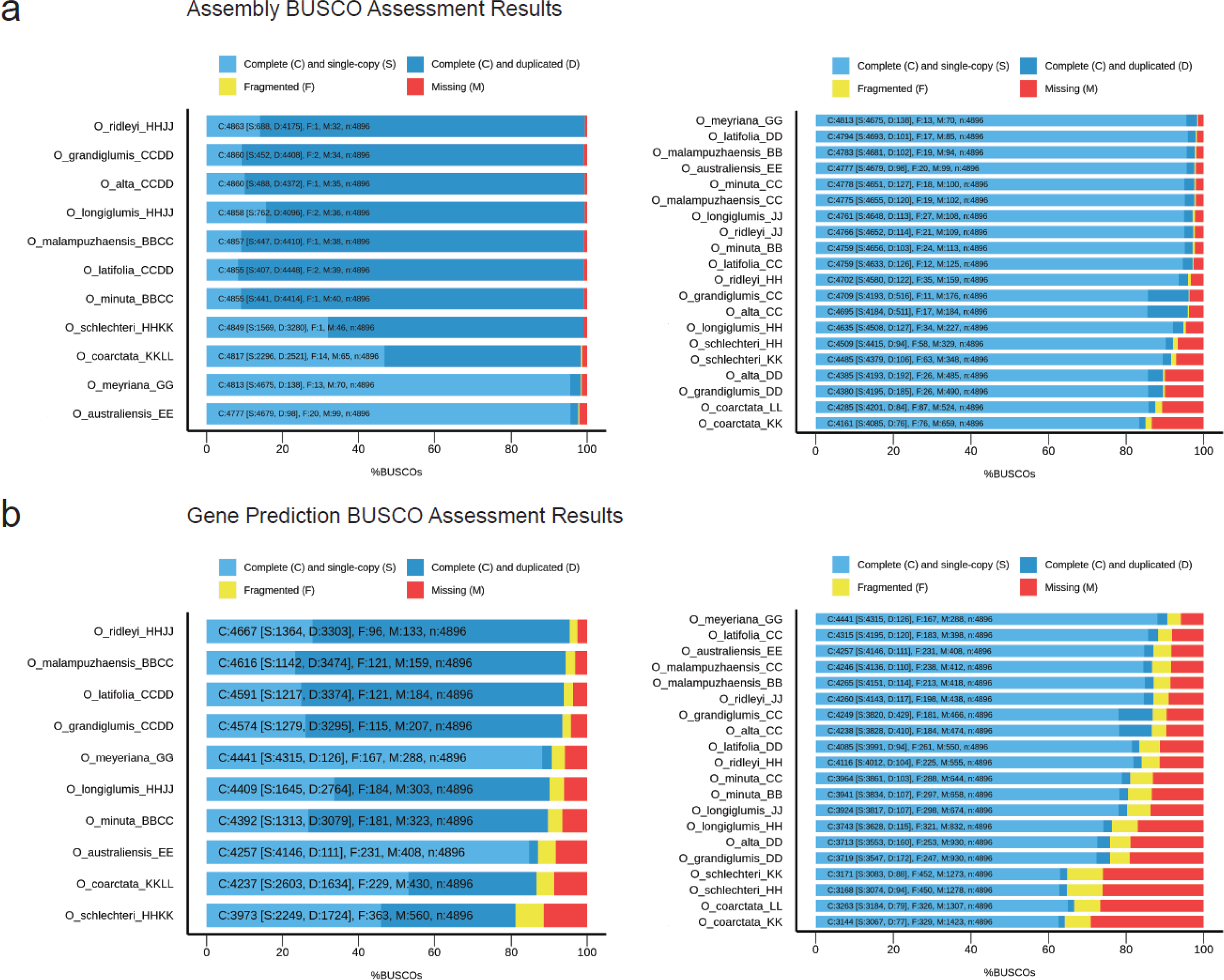
BUSCO assessment results. BUSCO assembly results for “Assembly” (A) and “Gene Prediction” (B) are shown at the genome level (left) and at the subgenome level (right) using the Poales database (4,896 genes). For each (sub)genome, barplots show the number of genes falling in the following categories: complete (C) and single copy (S) genes – light blue; complete (C) and duplicated (D) genes – dark blue; fragmented (F) genes - yellow; missing (M) genes - red. In each panel, (sub)genomes are ordered based on the percentage of missing genes, from the lowest (top) to the highest (bottom).

**Supplementary Figure 2.**
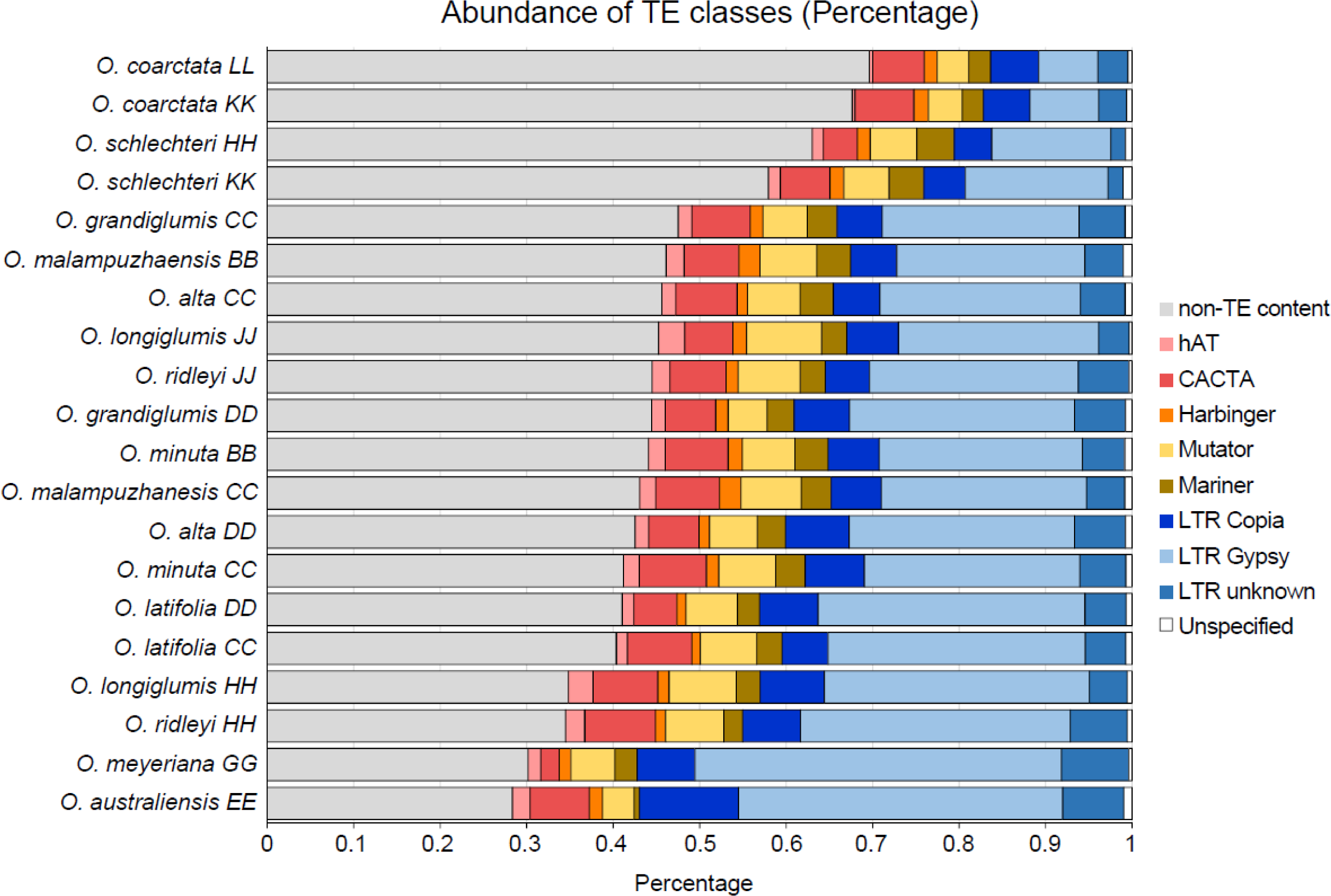
Percentage of the main classes of TEs in the wild *Oryza* (sub)genomes. (Sub)genomes are ordered based on the cumulative percentage of TE-related content, from the lowest (top row, LL subgenome in *O. coarctata*) to the highest (bottom row, *O. australiensis* EE). DNA transposons are shown as: hAT (DTA) - pink; CACTA (DTC) - red; Harbinger (DTH) - orange; Mutator (DTM) - yellow; Mariner (DTT) - ocra. LTR retrotransposons are shown as: LTR Copia – dark blue; LTR Gypsy – light blue; LTR Unknown – steel blue). Unspecified TEs are shown in white; non-TE content is shown in grey.

**Supplementary Figure 3.**
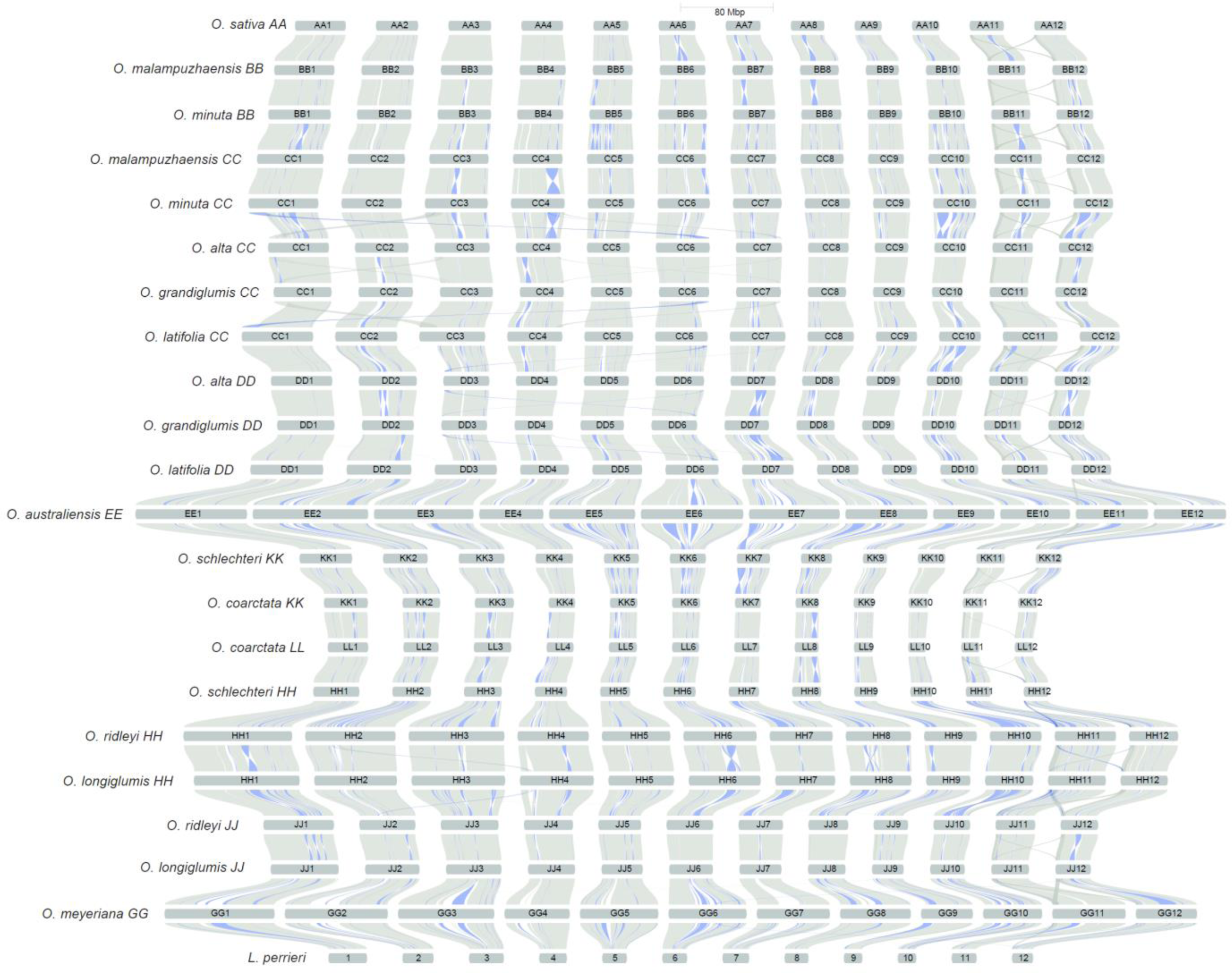
Inverted synteny blocks in the *Oryza* (sub)genomes. The riparian plot shows macro-synteny and collinearity across chromosomes in the *Oryza* (sub)genomes (grey). Blue ribbons highlight inverted collinear syntenic blocks in each pair of (sub)genomes. Genome types are shown according to the phylogenetic order in the genus, going from the top (*O. sativa* (AA)) to the bottom (*O. meyeriana* (GG)). *L. perrieri* is shown as an outgroup. Chromosomes are scaled by assembly length.

**Supplementary Figure 4.**
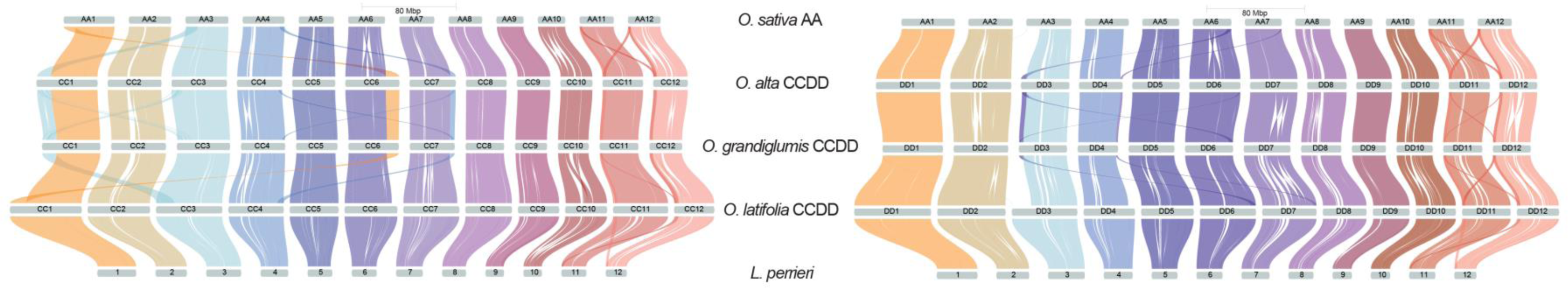
Non-reciprocal translocations in the subgenomes of *O. alta* and *O. grandiglumis*. Riparian plots show synteny between the subgenomes of *O. alta* (CCDD), *O. grandiglumis* (CCDD), *O. latifolia* (CCDD) and the genome of *O. sativa* (AA) revealing five non-reciprocal chromosomal translocations in the two former tetraploid species relative to *O. sativa*, which are absent in the latter. Left: non-reciprocal translocation in the CC subgenome (i.e. t(Chr1CC; Chr3), t(Chr6CC; Chr1), t(Chr7CC; Chr4)); right: non-reciprocal translocation in the DD subgenome (i.e. t(Chr3DD; Chr6); t(Chr4DD; Chr7)) in the genomes of *O. alta* and *O. grandiglumis*. *L. perrieri* is shown as the outgroup. A description of each translocation event follows: **a)** A segment of ∼10 Mb on Chr3 in *O. sativa* (0-10 Mb) was duplicated and translocated to the beginning of Chr1CC in *O. grandiglumis* (0-10 Mb) and *O. alta* (0-12 Mb), replacing the ∼10 Mb of Chr1 translocated on Chr6CC; **b)** A segment of ∼10 Mb on Chr1 in *O. sativa* (0-10 Mb) translocated to the end of Chr6CC in *O. grandiglumis* (33-42 Mb) and *O. alta* (35-45 Mb), replacing the telomeric portion of Chr6 (the last ∼3.5 Mb of Chr6 of *O. sativa* does not have any homologous sequence on Chr6CC in *O. grandiglumis* and *O. alta*); **c)** A segment of ∼3 Mb on Chr4 in *O. sativa* (32-36 Mb) was duplicated and translocated to the end of Chr7CC in *O. grandiglumis* (34-38 Mb) and *O. alta* (35-38 Mb), replacing the telomeric portion of Chr7 (the last ∼1.3 and ∼1.8 Mb on Chr7 on *O. sativa* do not have any homologous sequence on Chr7CC in *O. grandiglumis* and *O. alta*, respectively); **d)** A segment of ∼4 Mb on Chr6 in *O. sativa* (28-31 Mb) was duplicated and translocated to the beginning of Chr3DD in *O. grandiglumis* (0-3 Mb) and *O. alta* (0-4 Mb), replacing the telomeric portion of Chr3 in *O. sativa* (the first ∼11 Mb on Chr3 of *O. sativa* do not have any homologous sequence on Chr3DD in *O. grandiglumis* and *O. alta*); **e**) A segment of ∼3 Mb of Chr7 in *O. sativa* (28-30 Mb) translocated to the end of Chr4DD in *O. grandiglumi*s (31-33 Mb) and *O. alta* (34-35 Mb), replacing the telomeric portion of Chr4 (the last ∼4 Mb on Chr4 of *O. sativa* do not have any homologous sequence on Chr4DD in *O. grandiglumis* and *O. alta*).

**Supplementary Figure 5.**
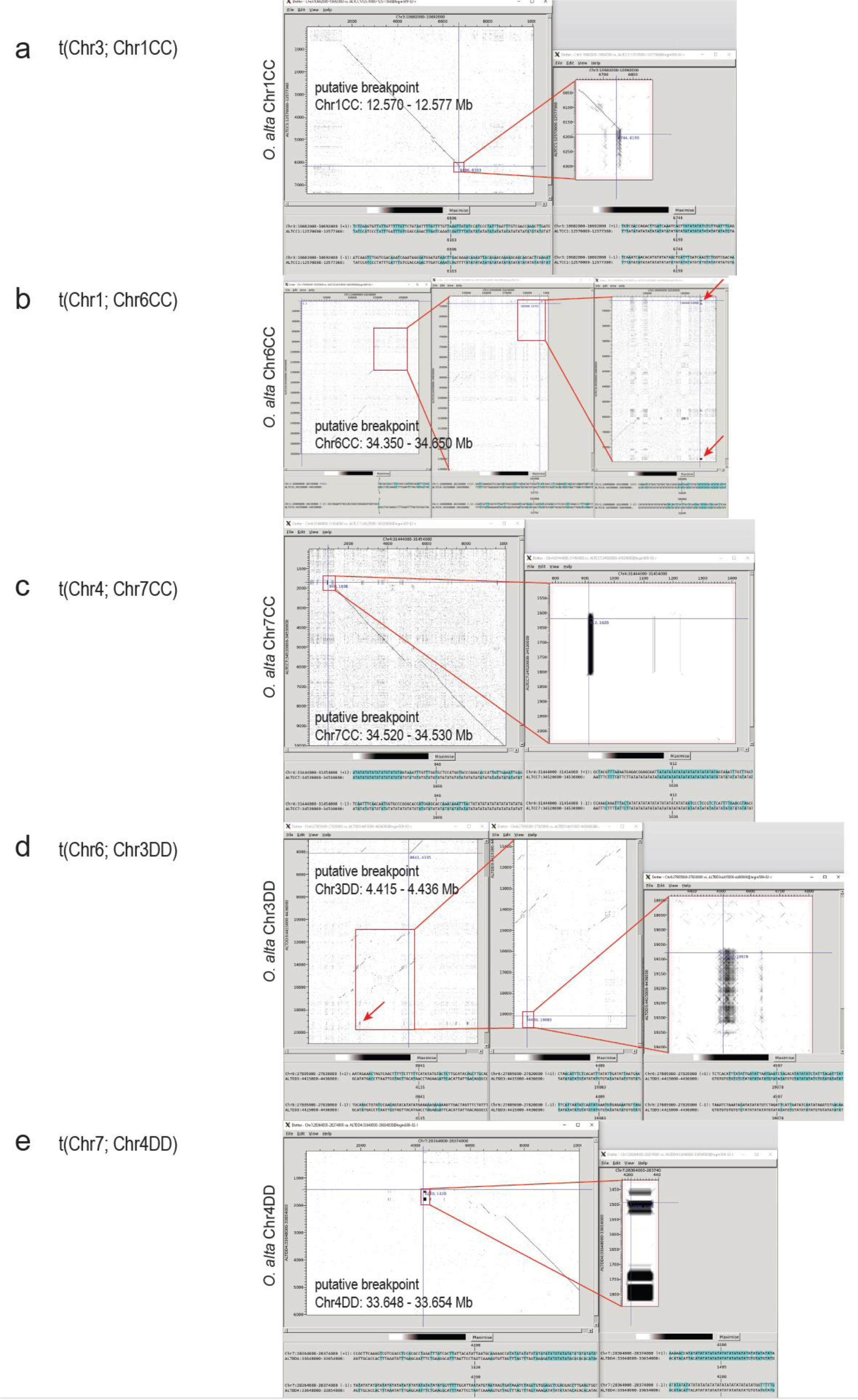
Dotplots of SRR sequences found at the putative translocation breakpoints in *O. alta*. For each translocation event, named from a) to e) as described in Supplementary Figure 4, *O. alta* (CCDD) (on the y-axis) is compared to *O. sativa* (AA) (on the x-axis) and the range for the putative translocation breakpoint position is defined in each panel. Two or three dotplots show increasing magnification of each putative translocation breakpoint to highlight SRR signal and sequence (AT repeats). Sequence comparisons were obtained using DOTTER^116^ in the SeqTools package (Genome Research Ltd; https://www.sanger.ac.uk/tool/seqtools/).

**Supplementary Figure 6.**
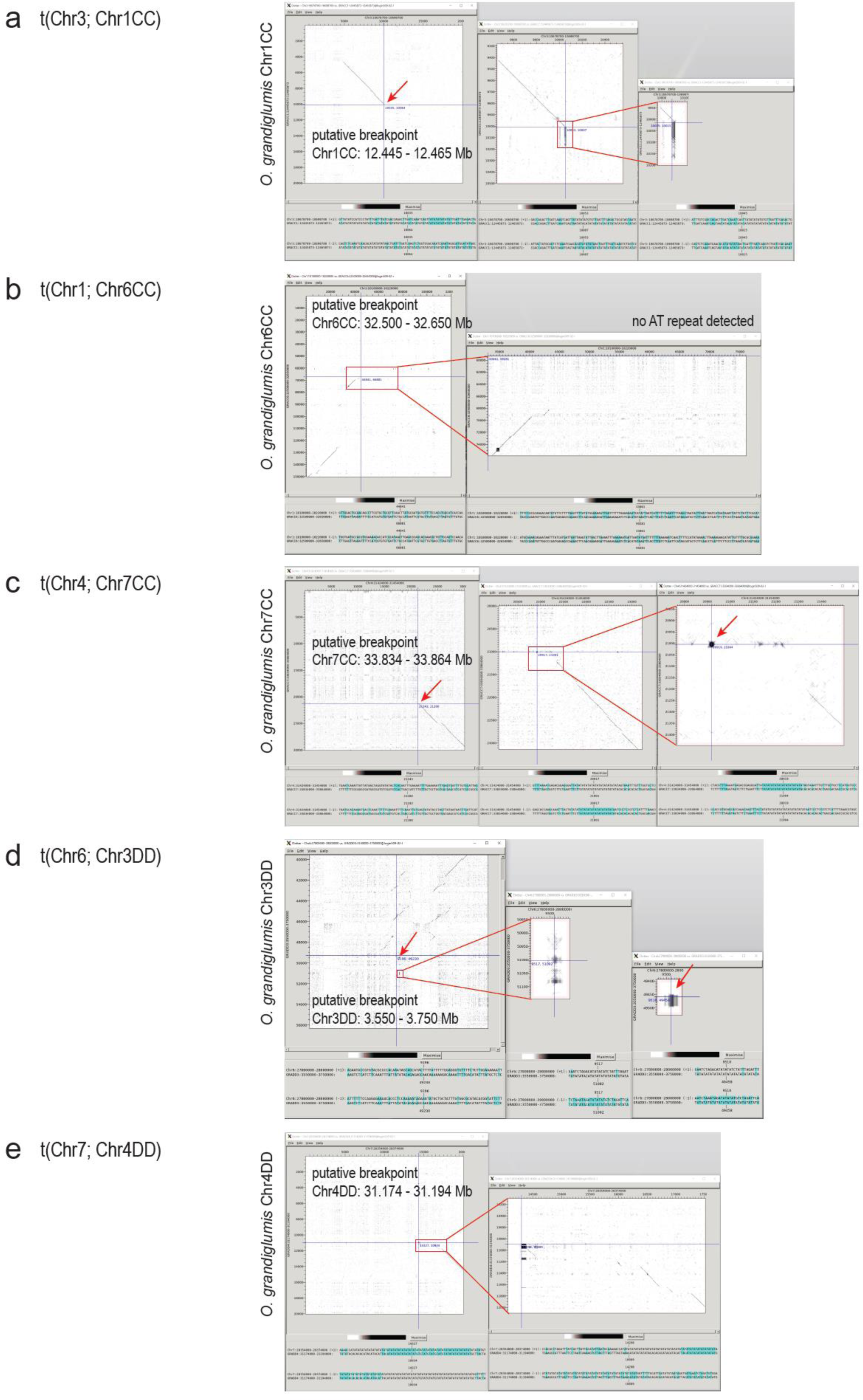
Dotplots of SRR sequences found at the putative translocation breakpoints in *O. grandiglumis*. For each translocation event, named from a) to e) as described in Supplementary Figure 4, *O. grandiglumis* (CCDD) (on the y-axis) is compared to *O. sativa* (AA) (on the x-axis) and the range for the putative translocation breakpoint position is defined in each panel. Two or three dotplots show increasing magnification of each putative translocation breakpoint to highlight SRR signal and sequence (AT repeats). Sequence comparisons were obtained using DOTTER^116^ in the SeqTools package (Genome Research Ltd; https://www.sanger.ac.uk/tool/seqtools/).

**Supplementary Figure 7.**
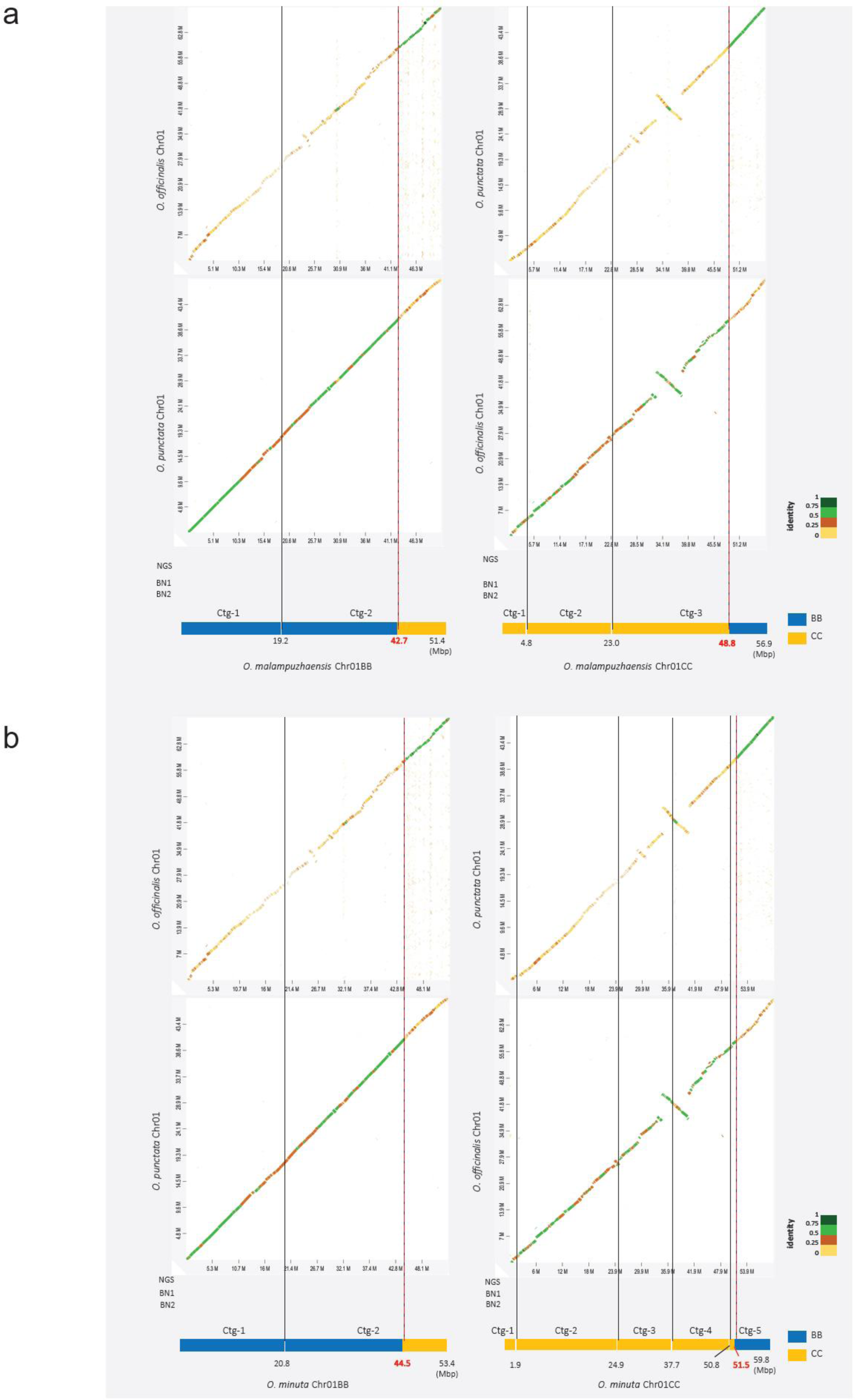
Reciprocal translocation between Chr1BB and Chr1CC in *O. malampuzhaensis* and *O. minuta*. **a)** In the top panel, dotplots show the percentage of sequence identity when aligning Chr1BB (left) and Chr1CC (right) of *O. malampuzhaensis* (BBCC) against chromosome 1 of *O. punctata* (BB) and *O. officinalis* (CC), respectively (percent identity < 0.25 - yellow; percent identity > 0.25 and < 0.5 - orange; percent identity > 0.5 and < 0.75 - light green; percent identity > 0.75 - dark green). Alignments were obtained using Minimap2^75^ implemented in D-Genies^117^, using the parameter *many repeats*. In the middle panel, optical maps (BN1 and BN2 tracks) support the scaffolding of the assembled chromosome (NGS track), i.e. Chr1BB (left) and Chr1CC (right). In the bottom panel, models representing contigs of the scaffolded chromosomes (i.e. Chr1BB (left) and Chr1CC (right)) are shown: black dashed lines represent subdivision into contigs; red dashed lines represent the putative breakpoint of the translocation; blue tracks represent contigs belonging to BB genome type; yellow tracks represent contigs belonging to CC genome type. **b)** In the top panel, dotplots show the percentage of sequence identity when aligning Chr1BB (left) and Chr1CC (right) of *O. minuta* (BBCC) against chromosome 1 of *O. punctata* (BB) and *O. officinalis* (CC), respectively (percent identity < 0.25 - yellow; percent identity > 0.25 and < 0.5 - orange; percent identity > 0.5 and < 0.75 - light green; percent identity > 0.75 - dark green). In the middle panel, optical maps (BN1 and BN2 tracks) support the scaffolding of the assembled chromosome (NGS track), i.e. Chr1BB (left) and Chr1CC (right). In the bottom panel, models representing contigs of the scaffolded chromosomes (i.e. Chr1BB (left) and Chr1CC (right)) are shown: black dashed lines represent subdivision into contigs; red dashed lines represent the putative breakpoint of the translocation; blue tracks represent contigs belonging to BB genome type; yellow tracks represent contigs belonging to CC genome type.

**Supplementary Figure 8.**
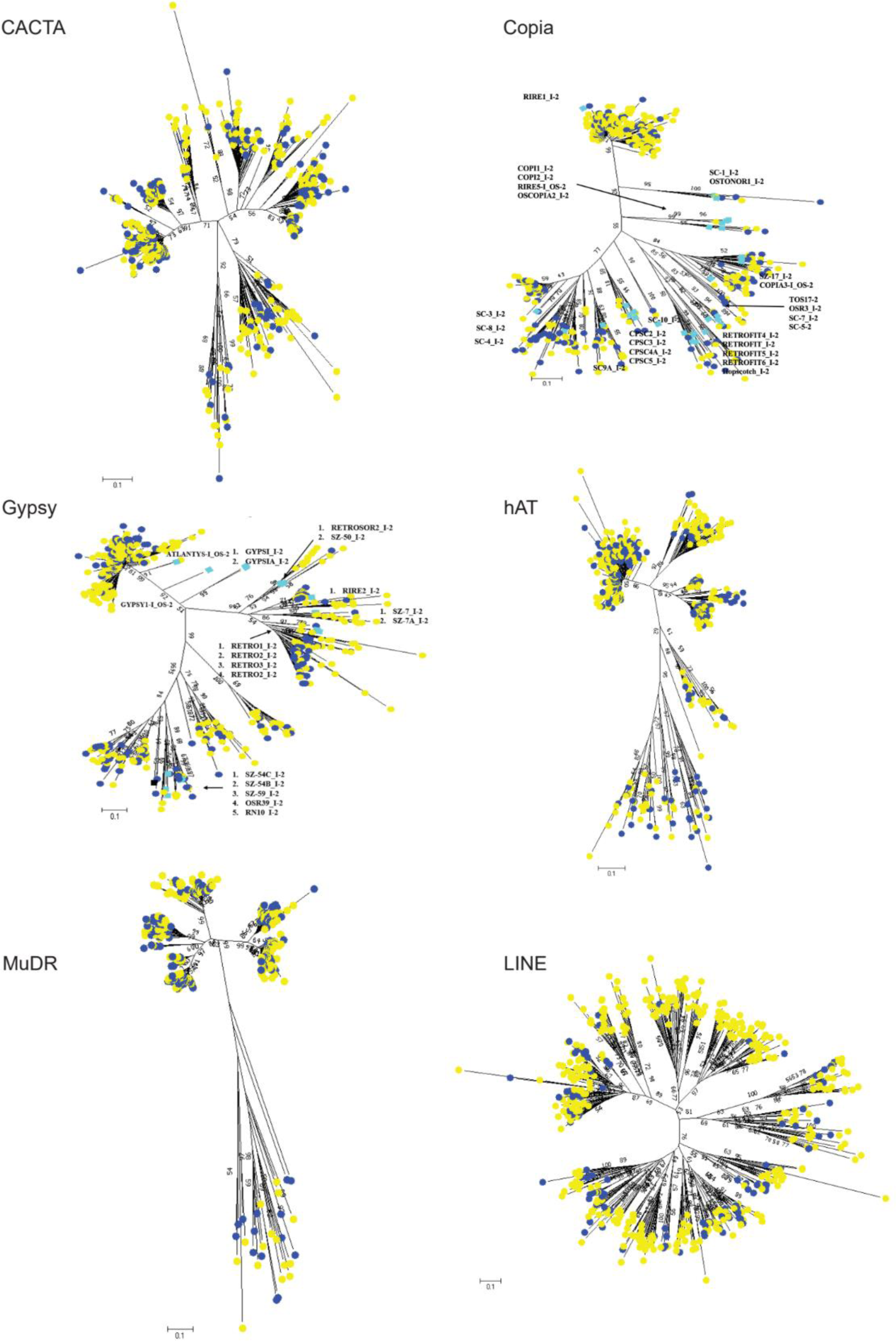
Distribution of TEs belonging to six *Oryza*-specific superfamilies in the subgenomes of *O. ridleyi* (HHJJ). Neighbor-Joining trees were built for six TE superfamilies (i.e. CACTA, Ty1/Copia, Ty3/Gypsy, MuDR, hAT, LINE) using 500 TEs randomly chosen from each subgenome. Bootstrap values were calculated for 1,000 replicates and shown when greater than 50. Paralogs from HH subgenome are indicated by yellow dots and those from JJ subgenome by blue dots. The representative sequences of major Ty3-Gypsy families identified in *O. sativa* (AA) are indicated by turquoise squares.

**Supplementary Figure 9.**
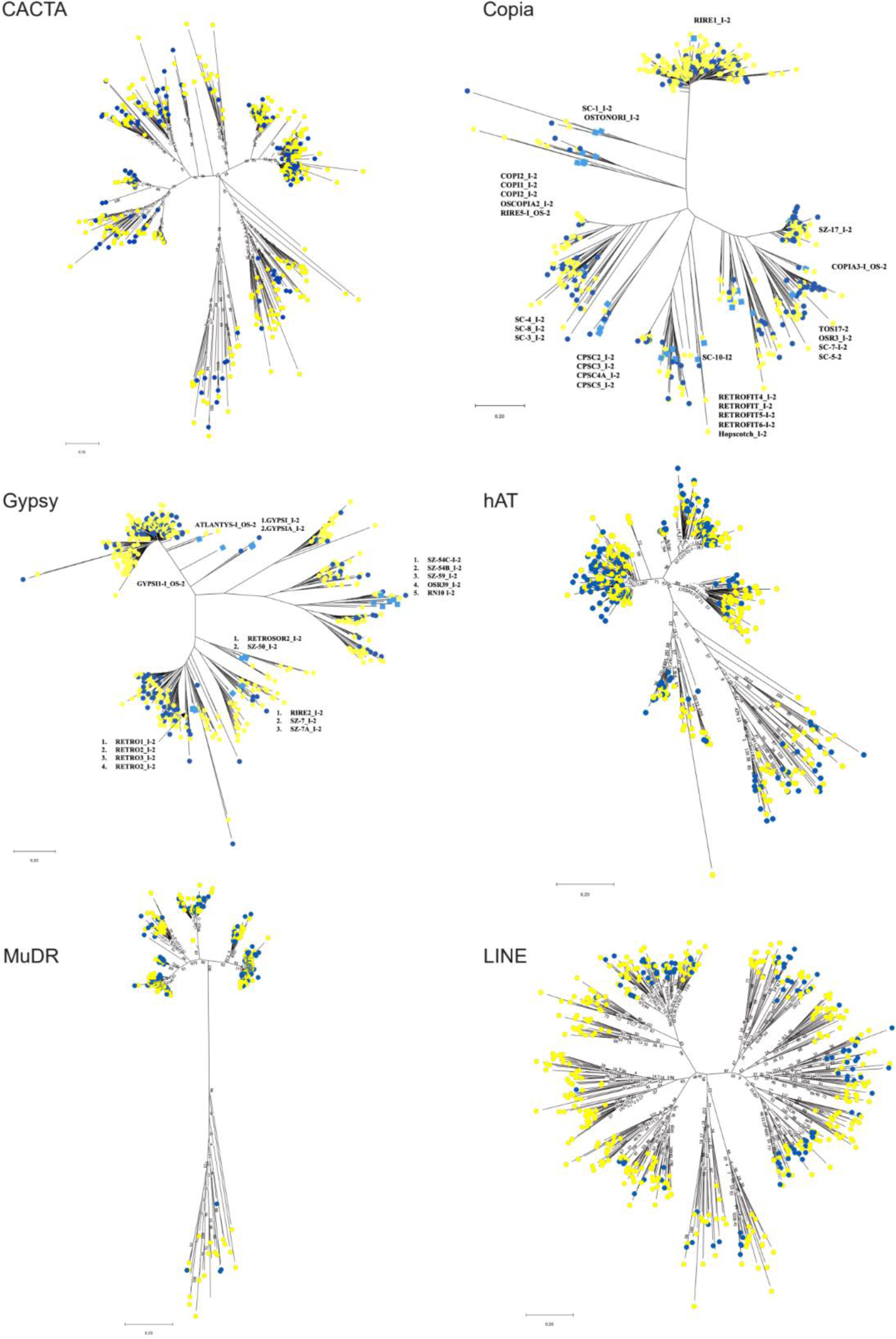
Distribution of TEs belonging to six *Oryza*-specific superfamilies in the subgenomes of *O. longiglumis* (HHJJ). Neighbor-Joining trees were built for six TE superfamilies (i.e. CACTA, Ty1/Copia, Ty3/Gypsy, MuDR, hAT, LINE) using 500 TEs randomly chosen from each subgenome. Bootstrap values were calculated for 1,000 replicates and shown when greater than 50. Paralogs from HH subgenome are indicated by yellow dots and those from JJ subgenome by blue dots. The representative sequences of major Ty3-Gypsy families identified in *O. sativa* (AA) are indicated by turquoise squares.

**Supplementary Figure 10.**
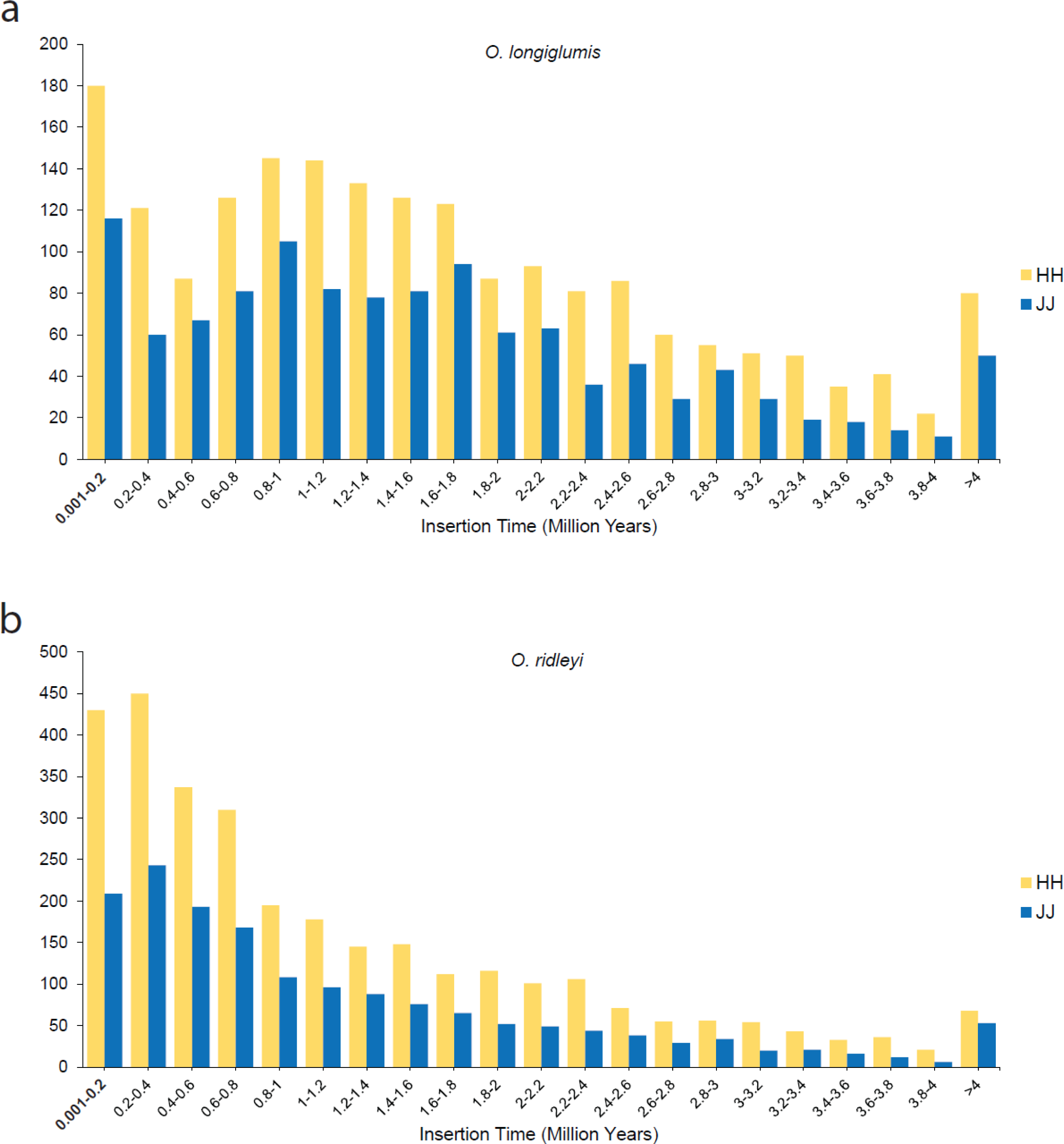
LTR insertion time in the subgenome of *O. longiglumis* and *O. ridleyi*. Histograms show insertion time (T) in Million years (My) measured as nucleotide distance (D) between LTR sequences flanking LTR-RTs in HH (yellow bars) and JJ (blue bars) subgenomes of **a)** *O. longiglumis* (HHJJ) and **b)** *O. ridleyi* (HHJJ), using the formula 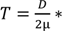 2μ 10^6^, where μ is a substitution rate of 1.3×10^−8^ per site per year.

**Supplementary Figure 11.**
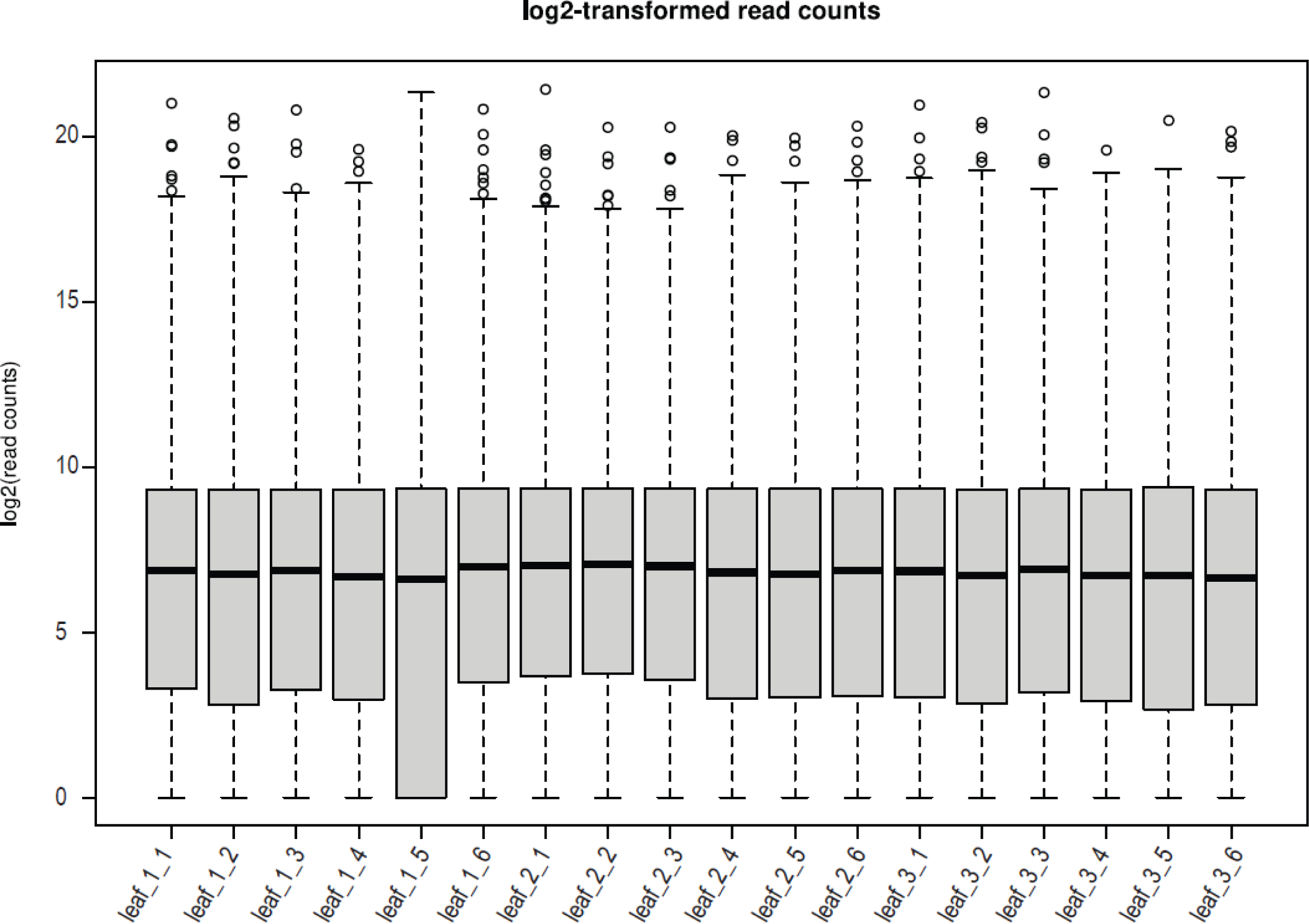
Distribution of log_2_-transformed read count of genes expressed in the leaf of *O. coarctata* (KKLL) per technical replicate.

**Supplementary Figure 12.**
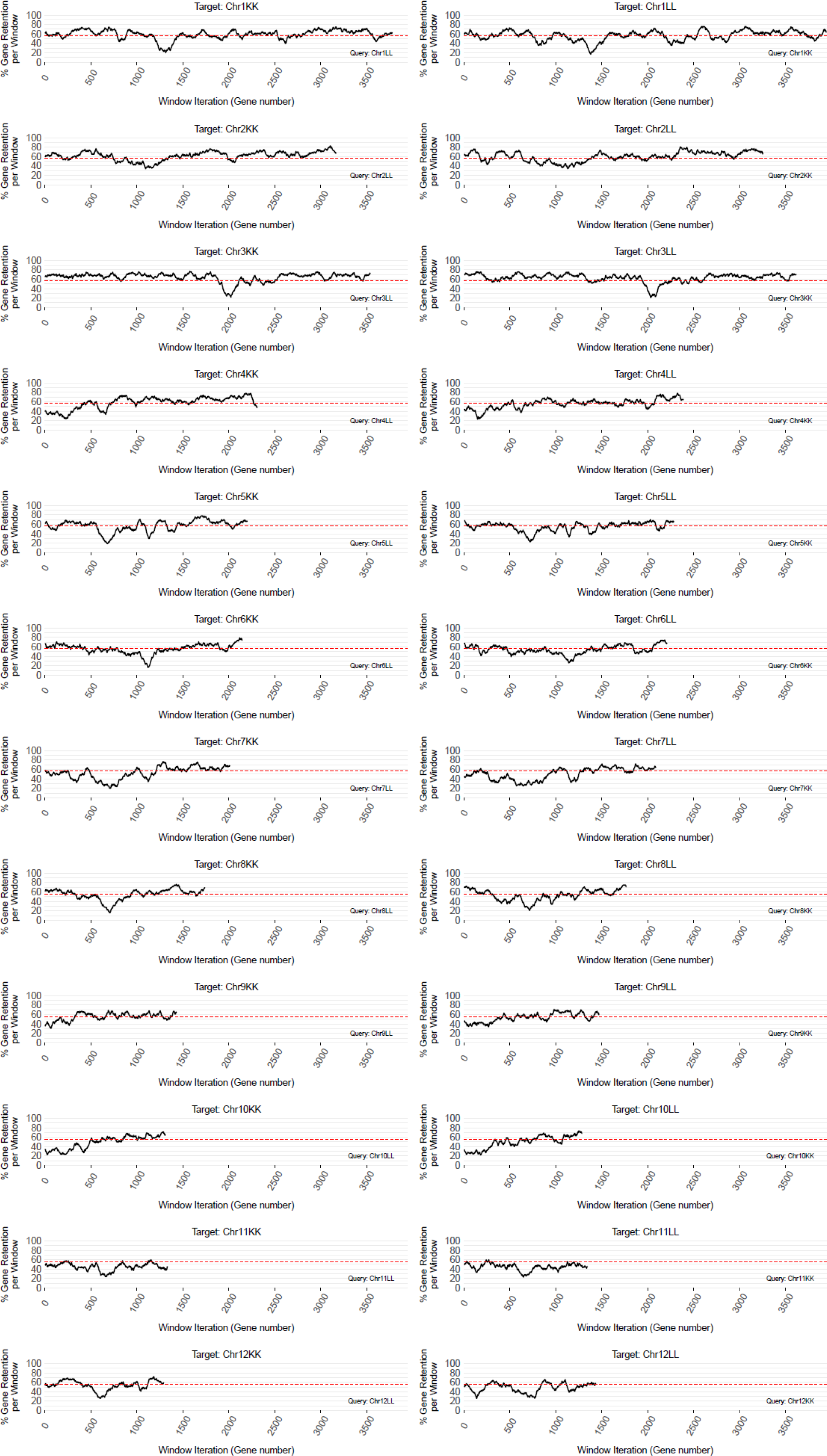
Distribution of gene fractionation in the homoeologous chromosomes of *O. coarctata*. The percentage of gene retention (y-axis) in each query chromosome of *O. coarctata* (KKLL) is shown in sliding windows of 100 genes along the corresponding homoeologous chromosome (i.e. the target chromosomes on the x-axis) using FractBias and default parameters. Lower percentage of gene retention indicates over-fractionation (i.e. syntenic regions with more gene loss), whereas higher percentage of gene retention indicates under-fractionation (i.e. syntenic regions with less gene loss). Red dashed line represents the average percentage of gene retention calculated genome-wide.

## Table and Supplementary Table legends

**Supplementary Table 1.**

Statistics of the optical genome molecules and maps of the *Oryza* species.

**Supplementary Table 2.**

BUSCO assessment of the genome assemblies (“Assembly”) and the assembled transcriptomes (“Gene Prediction”) of the wild *Oryza* species. Number of genes and percentages of each category in BUSCO are shown at the genome and subgenome level. The *poales_odb10* was used (4,896 genes in total).

**Supplementary Table 3.**

Abundance of the main classes of TEs (DNA transposons: DNA TE; Long Terminal Repeat retrotransposons: LTR Copia, LTR Gypsy, LTR Unknown) is shown in basepairs (bp) and in percentage (%). The class of DNA transposons includes: hAT (DTA), CACTA (DTC), Harbinger (DTH), Mutator (DTM), and Mariner (DTT).

**Supplementary Table 4.**

Chromosome and subgenome length bp of the *Oryza* species, total amount of TE content and non-TE content in bp.

**Supplementary Table 5.**

GenBank accessions of proteome sequences and number of the longest isoforms used in GENESPACE analysis. Proteome sequences of the species under study are shown in bold.

**Supplementary Table 6.**

List of genes in *O. alta* and *O. grandiglumis* that were duplicated and translocated as a consequence of chromosomal duplications and unbalanced translocations in these species, and replaced genes on the chromosomal portions that were lost. The homologous *sativa* genes are also reported.

**Supplementary Table 7.**

GenBank accessions of chloroplast genomes used in the chloroplast-based phylogenetic tree. Chloroplast genome sequences of the species under study are shown in bold.

**Supplementary Table 8.**

Average percentage of gene retention at the chromosome, subgenome, and genome level.

**Supplementary Table 9.**

Number of genes and median transcript abundance as log_2_(TPM + 1) in the four categories of homoeologous genes of *O. coarctata* and the two categories of homologous genes of *O. sativa*, in the leaf and in the root.

**Supplementary Table 10.**

Number of genes expressed at least two-fold higher compared to their homoeologs in the other subgenome, and genes not dominantly expressed, in the leaf and in the root. Binomial test p-value is provided for each tissue.

**Supplementary Data Table**. BioSample, BioProject, and SRA identification numbers. TBD: to be determined.

## Acknowledgements

This work was supported by King Abdullah University of Science and Technology (KAUST) with Baseline Funding to R.A.W. The authors thank Mara Miculan and Vanessa Melino for their contributions to the experimental design of transcriptome analysis and subgenome dominance in *O. coarctata*.

## Author Contributions

R.A.W. conceived and designed the project. R.A.W., A.Z. and A.F. coordinated the project. N.M. prepared the tissue samples for genome and transcriptome sequencing with assistance from D.K., D.C., S.R., L.S., and J.T. (at the Arizona Genomics Institute). C.S-S. provided computational support for data generation and transferring. N.A-B. grew and prepared the tissue samples for subgenome dominance analysis in *O. coarctata*. P.P. and V.L. generated optical maps for genome assembly validation. A.F., N.A-B., A.A., S.M., G.M., L.F. and A.Z. performed genome assembly and validation, gene and TE annotation of the *Oryza* species with significant contributions from Y.Z., L.R.S., V.L., and M.T. S.M. and A.Z. performed TE analysis in *O. ridleyi* and *O. longiglumis*. T.F. and N.V.H. performed phylogenomic analysis. N.A-B. performed Ks analysis. A.F. and T.F. performed macro- and micro-synteny analysis. A.F. performed gene fractionation analysis with significant contributions from M.E.S. G.M. performed analysis of subgenome dominance in *O. coarctata*. O.P. contributed transcriptome data for *O. australiensis*. K.L.M. grew, maintained and provided the germplasm collection of the *Oryza* tetraploids. M.E.S. and J.Z. contributed valuable suggestions to analyses and interpretation of results. A.F. wrote the paper with significant contributions from T.F., N.A-B., N.V.H., A.Z., M.E.S. and R.A.W.

## Data availability

All PacBio and Illumina DNA-Seq and RNA-Seq data have been deposited in GenBank. GenBank BioProject codes: *O. alta*, PRJNA1039467; *O. australiensis*, PRJNA591699; *O. coarctata*, PRJNA439330; *O. grandiglumis*, PRJNA737282; *O. latifolia*, PRJNA737486; *O. longiglumis*, PRJNA1016142; *O. malampuzhaensis*, PRJNA757598*; O. meyeriana*, PRJNA1039468; *O. minuta*, PRJNA757599; *O. ridleyi*, PRJNA687623; *O. schlechteri*, PRJNA732115

