## Supplementary_Tables for "*Oryza* genome evolution through a tetraploid lens"

**Supplementary Table 1.** Statistics of the optical genome molecules and maps of the *Oryza* species.

| | Molecules | | | | | Assembled Maps | | | |
|---|---|---|---|---|---|---|---|---|---|
| | Total number of molecules | Total length (Mbp) | Average length (kbp) | Molecule N50 (kbp) | Label density (/100kb) | Genome map count | Total genome map length (Mbp) | Genome map N50 (Mbp) | Max length (Mbp) |
| *O. malampuzhaensis* (IRGC 80765) | 1,046,725 | 353,159.69 | 337.395 | 329.307 | 11.467 | 61 | 1,002.19 | 21.811 | 38.798 |
| *O. minuta* (IRGC 101141) | 719,828 | 268,855.58 | 373.5 | 363.104 | 11.401 | 71 | 1,087.06 | 22.536 | 54.839 |
| *O. alta* (IRGC 105143) | 580,349 | 261,319.72 | 450.28 | 435.504 | 12.956 | 90 | 1,573.25 | 24.179 | |
| *O. grandiglumis* (IRGC 105669) | 577,227 | 251,654.11 | 435.971 | 423.16 | 13.222 | 72 | 961.573 | 20.43 | 46.744 |
| *O. latifolia* (IRGC 100890) | 584,460 | 227,600.63 | 389.42 | 377.874 | 14.853 | 63 | 1,646.26 | 39.163 | 65.773 |
| *O. australiensis* (IRGC 100882) | 1,000,780 | 308,768.55 | 308.528 | 310.712 | 13.455 | 182 | 1,413.81 | 64.958 | 119.363 |
| *O. coarctata* (IRGC 104502) | 278,430 | 146,400.84 | 525.808 | 509.356 | 15.007 | 30 | 592.678 | 24.182 | 39.376 |
| *O. schlechteri* (IRGC 82047) | 411,865 | 186,063.96 | 451.76 | 438.657 | 12.611 | 42 | 746.635 | 25.147 | 48.426 |
| *O. ridleyi* (IRGC 100821) | 551,560 | 298,170.73 | 540.595 | 527.45 | 14.614 | 27 | 1,256.72 | 56.084 | 97.177 |

**Supplementary Table 2.** BUSCO assessment of the genome assemblies ("Assembly") and the assembled transcriptomes ("Gene Predicti
Number of genes and percentages of each category in BUSCO are shown at the genome and subgenome level.
The Poales lineage version 10 was used (4,896 genes in total).

| Genome | Class | Assembly | Percentage | Gene Prediction | Percentage |
|---|---|---|---|---|---|
| *O. malampuzhaensis BBCC* | Complete BUSCOs | 4,857 | 99.20% | 4,616 | 94.30% |
| | Complete Single-Copy BUSCOs | 447 | 9.10% | 1,142 | 23.30% |
| | Complete Duplicated BUSCOs | 4,410 | 90.10% | 3,474 | 71.00% |
| | Fragmented BUSCOs | 1 | 0.00% | 121 | 2.50% |
| | Missing BUSCOs | 38 | 0.80% | 159 | 3.20% |
| | Total BUSCOs | 4,896 | 100% | 4,896 | 100% |
| *O. malampuzhaensis* BB | Complete BUSCOs | 4,783 | 97.70% | 4,265 | 87.10% |
| | Complete Single-Copy BUSCOs | 4,681 | 95.60% | 4,151 | 84.80% |
| | Complete Duplicated BUSCOs | 102 | 2.10% | 114 | 2.30% |
| | Fragmented BUSCOs | 19 | 0.40% | 213 | 4.40% |
| | Missing BUSCOs | 94 | 1.90% | 418 | 8.50% |
| | Total BUSCOs | 4,896 | 100% | 4,896 | 100% |
| *O. malampuzhaensis CC* | Complete BUSCOs | 4,775 | 97.60% | 4,246 | 86.70% |
| | Complete Single-Copy BUSCOs | 4,655 | 95.10% | 4,136 | 84.50% |
| | Complete Duplicated BUSCOs | 120 | 2.50% | 110 | 2.20% |
| | Fragmented BUSCOs | 19 | 0.40% | 238 | 4.90% |
| | Missing BUSCOs | 102 | 2.00% | 412 | 8.40% |
| | Total BUSCOs | 4,896 | 100% | 4,896 | 100% |
| *O. minuta BBCC* | Complete BUSCOs | 4,855 | 99.20% | 4,392 | 89.70% |
| | Complete Single-Copy BUSCOs | 441 | 9.00% | 1,313 | 26.80% |
| | Complete Duplicated BUSCOs | 4,414 | 90.20% | 3,079 | 62.90% |
| | Fragmented BUSCOs | 1 | 0.00% | 181 | 3.70% |
| | Missing BUSCOs | 40 | 0.80% | 323 | 6.60% |
| | Total BUSCOs | 4,896 | 100% | 4,896 | 100% |
| *O. minuta* BB | Complete BUSCOs | 4,759 | 97.20% | 3,941 | 80.50% |
| | Complete Single-Copy BUSCOs | 4,656 | 95.10% | 3,834 | 78.30% |
| | Complete Duplicated BUSCOs | 103 | 2.10% | 107 | 2.20% |
| | Fragmented BUSCOs | 24 | 0.50% | 297 | 6.10% |
| | Missing BUSCOs | 113 | 2.30% | 658 | 13.40% |
| | Total BUSCOs | 4,896 | 100% | 4,896 | 100% |
| *O. minuta* CC | Complete BUSCOs | 4,778 | 97.60% | 3,964 | 81.00% |
| | Complete Single-Copy BUSCOs | 4,651 | 95.00% | 3,861 | 78.90% |
| | Complete Duplicated BUSCOs | 127 | 2.60% | 103 | 2.10% |
| | Fragmented BUSCOs | 18 | 0.40% | 288 | 5.90% |
| | Missing BUSCOs | 100 | 2.00% | 644 | 13.10% |
| | Total BUSCOs | 4,896 | 100% | 4,896 | 100% |

| | | | | | |
|---|---|---|---|---|---|
| *O. alta CCDD* | Complete BUSCOs | 4,860 | 99.30% | 4,571 | 93.40% |
| | Complete Single-Copy BUSCOs | 488 | 10.00% | 1,325 | 27.10% |
| | Complete Duplicated BUSCOs | 4,372 | 89.30% | 3,246 | 66.30% |
| | Fragmented BUSCOs | 1 | 0.00% | 128 | 2.60% |
| | Missing BUSCOs | 35 | 0.70% | 197 | 4.00% |
| | Total BUSCOs | 4,896 | 100% | 4,896 | 100% |
| *O. alta* CC | Complete BUSCOs | 4,695 | 95.90% | 4,238 | 86.60% |
| | Complete Single-Copy BUSCOs | 4,184 | 85.50% | 3,828 | 78.20% |
| | Complete Duplicated BUSCOs | 511 | 10.40% | 410 | 8.40% |
| | Fragmented BUSCOs | 17 | 0.30% | 184 | 3.80% |
| | Missing BUSCOs | 184 | 3.80% | 474 | 9.60% |
| | Total BUSCOs | 4,896 | 100% | 4,896 | 100% |
| *O. alta* DD | Complete BUSCOs | 4,385 | 89.50% | 3,713 | 75.90% |
| | Complete Single-Copy BUSCOs | 4,193 | 85.60% | 3,553 | 72.60% |
| | Complete Duplicated BUSCOs | 192 | 3.90% | 160 | 3.30% |
| | Fragmented BUSCOs | 26 | 0.50% | 253 | 5.20% |
| | Missing BUSCOs | 485 | 10.00% | 930 | 18.90% |
| | Total BUSCOs | 4,896 | 100% | 4,896 | 100% |
| *O. grandiglumis CCDD* | Complete BUSCOs | 4,860 | 99.20% | 4,574 | 93.40% |
| | Complete Single-Copy BUSCOs | 452 | 9.20% | 1,279 | 26.10% |
| | Complete Duplicated BUSCOs | 4,408 | 90.00% | 3,295 | 67.30% |
| | Fragmented BUSCOs | 2 | 0.00% | 115 | 2.30% |
| | Missing BUSCOs | 34 | 0.80% | 207 | 4.30% |
| | Total BUSCOs | 4,896 | 100% | 4,896 | 100% |
| *O. grandiglumis* CC | Complete BUSCOs | 4,709 | 96.10% | 4,249 | 86.80% |
| | Complete Single-Copy BUSCOs | 4,193 | 85.60% | 3,820 | 78.00% |
| | Complete Duplicated BUSCOs | 516 | 10.50% | 429 | 8.80% |
| | Fragmented BUSCOs | 11 | 0.20% | 181 | 3.70% |
| | Missing BUSCOs | 176 | 3.70% | 466 | 9.50% |
| | Total BUSCOs | 4,896 | 100% | 4,896 | 100% |
| *O. grandiglumis* DD | Complete BUSCOs | 4,380 | 89.50% | 3,719 | 75.90% |
| | Complete Single-Copy BUSCOs | 4,195 | 85.70% | 3,547 | 72.40% |
| | Complete Duplicated BUSCOs | 185 | 3.80% | 172 | 3.50% |
| | Fragmented BUSCOs | 26 | 0.50% | 247 | 5.00% |
| | Missing BUSCOs | 490 | 10.00% | 930 | 19.10% |
| | Total BUSCOs | 4,896 | 100% | 4,896 | 100% |
| *O. latifolia CCDD* | Complete BUSCOs | 4,855 | 99.10% | 4,591 | 93.80% |
| | Complete Single-Copy BUSCOs | 407 | 8.30% | 1,217 | 24.90% |
| | Complete Duplicated BUSCOs | 4,448 | 90.80% | 3,374 | 68.90% |
| | Fragmented BUSCOs | 2 | 0.00% | 121 | 2.50% |

| | | | | | |
|---|---|---|---|---|---|
| | Missing BUSCOs | 39 | 0.90% | 184 | 3.70% |
| | Total BUSCOs | 4,896 | 100% | 4,896 | 100% |
| *O. latifolia* CC | Complete BUSCOs | 4,759 | 97.20% | 4,315 | 88.20% |
| | Complete Single-Copy BUSCOs | 4,633 | 94.60% | 4,195 | 85.70% |
| | Complete Duplicated BUSCOs | 126 | 2.60% | 120 | 2.50% |
| | Fragmented BUSCOs | 12 | 0.20% | 183 | 3.70% |
| | Missing BUSCOs | 125 | 2.60% | 398 | 8.10% |
| | Total BUSCOs | 4,896 | 100% | 4,896 | 100% |
| *O. latifolia* DD | Complete BUSCOs | 4,794 | 98.00% | 4,085 | 83.40% |
| | Complete Single-Copy BUSCOs | 4,693 | 95.90% | 3,991 | 81.50% |
| | Complete Duplicated BUSCOs | 101 | 2.10% | 94 | 1.90% |
| | Fragmented BUSCOs | 17 | 0.30% | 261 | 5.30% |
| | Missing BUSCOs | 85 | 1.70% | 550 | 11.30% |
| | Total BUSCOs | 4,896 | 100% | 4,896 | 100% |
| *O. australiensis EE* | Complete BUSCOs | 4,777 | 97.60% | 4,257 | 87.00% |
| | Complete Single-Copy BUSCOs | 4,679 | 95.60% | 4,146 | 84.70% |
| | Complete Duplicated BUSCOs | 98 | 2.00% | 111 | 2.30% |
| | Fragmented BUSCOs | 20 | 0.40% | 231 | 4.70% |
| | Missing BUSCOs | 99 | 2.00% | 408 | 8.30% |
| | Total BUSCOs | 4,896 | 100% | 4,896 | 100% |
| *O. coarctata KKLL* | Complete BUSCOs | 4,817 | 98.40% | 4,237 | 86.60% |
| | Complete Single-Copy BUSCOs | 2,296 | 46.90% | 2,603 | 53.20% |
| | Complete Duplicated BUSCOs | 2,521 | 51.50% | 1,634 | 33.40% |
| | Fragmented BUSCOs | 14 | 0.30% | 229 | 4.70% |
| | Missing BUSCOs | 65 | 1.30% | 430 | 8.70% |
| | Total BUSCOs | 4,896 | 100% | 4,896 | 100% |
| *O. coarctata* KK | Complete BUSCOs | 4,161 | 85.00% | 3,144 | 64.20% |
| | Complete Single-Copy BUSCOs | 4,085 | 83.40% | 3,067 | 62.60% |
| | Complete Duplicated BUSCOs | 76 | 1.60% | 77 | 1.60% |
| | Fragmented BUSCOs | 76 | 1.60% | 329 | 6.70% |
| | Missing BUSCOs | 659 | 13.40% | 1,423 | 29.10% |
| | Total BUSCOs | 4,896 | 100% | 4,896 | 100% |
| *O. coarctata* LL | Complete BUSCOs | 4,285 | 87.50% | 3,263 | 66.60% |
| | Complete Single-Copy BUSCOs | 4,201 | 85.80% | 3,184 | 65.00% |
| | Complete Duplicated BUSCOs | 84 | 1.70% | 79 | 1.60% |
| | Fragmented BUSCOs | 87 | 1.80% | 326 | 6.70% |
| | Missing BUSCOs | 524 | 10.70% | 1,307 | 26.70% |
| | Total BUSCOs | 4,896 | 100% | 4,896 | 100% |
| | Complete BUSCOs | 4,849 | 99.00% | 3,973 | 81.10% |
| | Complete Single-Copy BUSCOs | 1,569 | 32.00% | 2,249 | 45.90% |

| | | | | | |
|---|---|---|---|---|---|
| *O. schlechteri HHKK* | Complete Duplicated BUSCOs | 3,280 | 67.00% | 1,724 | 35.20% |
| | Fragmented BUSCOs | 1 | 0.00% | 363 | 7.40% |
| | Missing BUSCOs | 46 | 1.00% | 560 | 11.50% |
| | Total BUSCOs | 4,896 | 100% | 4,896 | 100% |
| *O. schlechteri* HH | Complete BUSCOs | 4,509 | 92.10% | 3,168 | 64.70% |
| | Complete Single-Copy BUSCOs | 4,415 | 90.20% | 3,074 | 62.80% |
| | Complete Duplicated BUSCOs | 94 | 1.90% | 94 | 1.90% |
| | Fragmented BUSCOs | 58 | 1.20% | 450 | 9.20% |
| | Missing BUSCOs | 329 | 6.70% | 1,278 | 26.10% |
| | Total BUSCOs | 4,896 | 100% | 4,896 | 100% |
| *O. schlechteri* KK | Complete BUSCOs | 4,485 | 91.60% | 3,171 | 64.80% |
| | Complete Single-Copy BUSCOs | 4,379 | 89.40% | 3,083 | 63.00% |
| | Complete Duplicated BUSCOs | 106 | 2.20% | 88 | 1.80% |
| | Fragmented BUSCOs | 63 | 1.30% | 452 | 9.20% |
| | Missing BUSCOs | 348 | 7.10% | 1,273 | 26.00% |
| | Total BUSCOs | 4,896 | 100% | 4,896 | 100% |
| *O. longiglumis HHJJ* | Complete BUSCOs | 4,858 | 99.30% | 4,409 | 90.10% |
| | Complete Single-Copy BUSCOs | 762 | 15.60% | 1,645 | 33.60% |
| | Complete Duplicated BUSCOs | 4,096 | 83.70% | 2,764 | 56.50% |
| | Fragmented BUSCOs | 2 | 0.00% | 184 | 3.80% |
| | Missing BUSCOs | 36 | 0.70% | 303 | 6.10% |
| | Total BUSCO | 4,896 | 100% | 4,896 | 100% |
| *O. longiglumis* HH | Complete BUSCOs | 4,635 | 94.70% | 3,743 | 76.40% |
| | Complete Single-Copy BUSCOs | 4,508 | 92.10% | 3,628 | 74.10% |
| | Complete Duplicated BUSCOs | 127 | 2.60% | 115 | 2.30% |
| | Fragmented BUSCOs | 34 | 0.70% | 321 | 6.60% |
| | Missing BUSCOs | 227 | 4.60% | 832 | 17.00% |
| | Total BUSCOss | 4896 | 100% | 4,896 | 100% |
| *O. longiglumis* JJ | Complete BUSCOs | 4,761 | 97.20% | 3,924 | 80.20% |
| | Complete Single-Copy BUSCOs | 4,648 | 94.90% | 3,817 | 78.00% |
| | Complete Duplicated BUSCOs | 113 | 2.30% | 107 | 2.20% |
| | Fragmented BUSCOs | 27 | 0.60% | 298 | 6.10% |
| | Missing BUSCOs | 108 | 2.20% | 674 | 13.70% |
| | Total BUSCOs | 4,896 | 100% | 4,896 | 100% |
| *O. ridleyi HHJJ* | Complete BUSCOs | 4,863 | 99.40% | 4,667 | 95.40% |
| | Complete Single-Copy BUSCOs | 688 | 14.10% | 1,364 | 27.90% |
| | Complete Duplicated BUSCOs | 4,175 | 85.30% | 3,303 | 67.50% |
| | Fragmented BUSCOs | 1 | 0.00% | 96 | 2.00% |
| | Missing BUSCOs | 32 | 0.60% | 133 | 2.60% |
| | Total BUSCOs | 4,896 | 100% | 4,896 | 100% |

| | | | | | |
|---|---|---|---|---|---|
| *O. ridleyi* HH | Complete BUSCOs | 4,702 | 96.00% | 4,116 | 84.00% |
| | Complete Single-Copy BUSCOs | 4,580 | 93.50% | 4,012 | 81.90% |
| | Complete Duplicated BUSCOs | 122 | 2.50% | 104 | 2.10% |
| | Fragmented BUSCOs | 35 | 0.70% | 225 | 4.60% |
| | Missing BUSCOs | 159 | 3.30% | 555 | 11.40% |
| | Total BUSCOs | 4,896 | 100% | 4,896 | 100% |
| *O. ridleyi* JJ | Complete BUSCOs | 4,766 | 97.30% | 4,260 | 87.00% |
| | Complete Single-Copy BUSCOs | 4,652 | 95.00% | 4,143 | 84.60% |
| | Complete Duplicated BUSCOs | 114 | 2.30% | 117 | 2.40% |
| | Fragmented BUSCOs | 21 | 0.40% | 198 | 4.00% |
| | Missing BUSCOs | 109 | 2.30% | 438 | 9.00% |
| | Total BUSCOs | 4,896 | 100% | 4,896 | 100% |
| *O. meyeriana GG* | Complete BUSCOs | 4,813 | 98.30% | 4,441 | 90.70% |
| | Complete Single-Copy BUSCOs | 4,675 | 95.50% | 4,315 | 88.10% |
| | Complete Duplicated BUSCOs | 138 | 2.80% | 126 | 2.60% |
| | Fragmented BUSCOs | 13 | 0.30% | 167 | 3.40% |
| | Missing BUSCOs | 70 | 1.40% | 288 | 5.90% |
| | Total BUSCOs | 4,896 | 100% | 4,896 | 100% |

**Supplementary Table 3.** Abundance of the main classes of TEs (DNA transposons: DNA TE; Long Terminal Repeat retrotransposons
The class of DNA transposons includes: hAT (DTA), CACTA (DTC), Harbinger (DTH), Mutator (DTM), and Mariner (DTT).

| Species | Type | Genome | | Subgenome 1 | | Subgenome 2 | |
|---|---|---|---|---|---|---|---|
| | | Length (bp) | Percentage | Length (bp) | Percentage | Length (bp) | Percentage |
| | DNA TE | 203,289,589 | 21.75 | 93,632,859 | 21.34 | 109,656,730 | 22.11 |
| | LTR Gypsy | 213,092,549 | 22.80 | 95,495,109 | 21.75 | 117,597,440 | 23.73 |
| ***O. malampuzhaensis*** | LTR Copia | 52,214,202 | 5.59 | 23,376,179 | 5.32 | 28,838,023 | 5.82 |
| **BBCC** | LTR Unknown | 41,445,528 | 4.43 | 19,539,529 | 4.45 | 21,905,999 | 4.42 |
| | Unspecified | 8,556,538 | 0.92 | 4,448,194 | 1.01 | 4,108,344 | 0.83 |
| | **Total** | **518,598,406** | **55.49** | **236,491,870** | **53.87** | **282,106,536** | **56.91** |
| | DNA TE | 204,863,603 | 20.90 | 95,120,202 | 20.76 | 109,743,401 | 21.03 |
| | LTR Gypsy | 237,658,165 | 24.24 | 107,536,144 | 23.47 | 130,122,021 | 24.92 |
| ***O. minuta*** | LTR Copia | 62,757,020 | 6.40 | 27,078,149 | 5.91 | 35,678,871 | 6.83 |
| **BBCC** | LTR Unknown | 50,200,992 | 5.12 | 22,404,786 | 4.89 | 27,796,206 | 5.32 |
| | Unspecified | 7,668,992 | 0.78 | 3,962,965 | 0.86 | 3,706,027 | 0.71 |
| | **Total** | **563,148,772** | **57.44** | **256,102,246** | **55.89** | **307,046,526** | **58.81** |
| | DNA TE | 169,025,320 | 18.59 | 90,934,129 | 19.80 | 78,091,191 | 17.37 |
| | LTR Gypsy | 223,622,239 | 24.60 | 106,441,199 | 23.18 | 117,181,040 | 26.05 |
| ***O. alta*** | LTR Copia | 57,928,011 | 6.37 | 24,792,391 | 5.40 | 33,135,620 | 7.37 |
| **CCDD** | LTR Unknown | 50,179,778 | 5.52 | 23,715,195 | 5.17 | 26,464,583 | 5.88 |
| | Unspecified | 7,134,872 | 0.78 | 3,690,505 | 0.80 | 3,444,367 | 0.77 |
| | **Total** | **507,890,220** | **55.86** | **249,573,419** | **54.35** | **258,316,801** | **57.44** |
| | DNA TE | 151,844,217 | 17.43 | 80,955,071 | 18.35 | 70,889,146 | 16.48 |
| | LTR Gypsy | 212,134,359 | 24.35 | 100,269,989 | 22.73 | 111,864,370 | 26.02 |
| ***O. grandiglumis*** | LTR Copia | 50,752,194 | 5.83 | 23,228,145 | 5.26 | 27,524,049 | 6.40 |
| **CCDD** | LTR Unknown | 48,826,187 | 5.60 | 23,503,687 | 5.33 | 25,322,500 | 5.89 |
| | Unspecified | 6,795,376 | 0.78 | 3,546,108 | 0.80 | 3,249,268 | 0.76 |
| | **Total** | **470,352,333** | **53.99** | **231,503,000** | **52.47** | **238,849,333** | **55.55** |
| | DNA TE | 185,820,267 | 17.55 | 103,948,068 | 19.14 | 81,872,199 | 15.89 |
| | LTR Gypsy | 320,604,719 | 30.29 | 161,596,227 | 29.75 | 159,008,492 | 30.86 |
| ***O. latifolia*** | LTR Copia | 63,712,718 | 6.02 | 28,815,923 | 5.31 | 34,896,795 | 6.77 |
| **CCDD** | LTR Unknown | 49,849,017 | 4.71 | 25,431,388 | 4.68 | 24,417,629 | 4.74 |
| | Unspecified | 7,566,845 | 0.71 | 3,968,412 | 0.73 | 3,598,433 | 0.70 |
| | **Total** | **627,553,566** | **59.28** | **323,760,018** | **59.61** | **303,793,548** | **58.96** |
| | DNA TE | 129,667,742 | 14.70 | - | - | - | - |
| | LTR Gypsy | 330,566,969 | 37.50 | - | - | - | - |
| ***O. australiensis*** | LTR Copia | 100,604,566 | 11.41 | - | - | - | - |
| **EE** | LTR Unknown | 62,398,218 | 7.08 | - | - | - | - |
| | Unspecified | 8,297,780 | 0.94 | - | - | - | - |
| | **Total** | **631,535,275** | **71.63** | - | - | - | - |
| | DNA TE | 81,233,584 | 14.61 | 43,201,323 | 15.19 | 38,032,261 | 14.02 |
| | LTR Gypsy | 41,215,650 | 7.42 | 22,638,000 | 7.96 | 18,577,650 | 6.85 |
| ***O. coarctata*** | LTR Copia | 30,319,231 | 5.46 | 15,282,702 | 5.37 | 15,036,529 | 5.54 |
| **KKLL** | LTR Unknown | 18,569,896 | 3.34 | 9,102,633 | 3.20 | 9,467,263 | 3.49 |
| | Unspecified | 3,083,956 | 0.55 | 1,828,539 | 0.64 | 1,255,417 | 0.46 |
| | **Total** | **174,422,317** | **31.38** | **92,053,197** | **32.36** | **82,369,120** | **30.36** |
| | DNA TE | 116,170,212 | 17.23 | 51,791,822 | 16.43 | 64,378,390 | 17.94 |
| | LTR Gypsy | 102,624,488 | 15.22 | 43,426,346 | 13.77 | 59,198,142 | 16.50 |
| ***O. schlechteri*** | LTR Copia | 30,966,683 | 4.59 | 13,736,938 | 4.35 | 17,229,745 | 4.80 |
| **HHKK** | LTR Unknown | 11,444,316 | 1.70 | 5,258,457 | 1.67 | 6,185,859 | 1.72 |
| | Unspecified | 6,239,855 | 0.93 | 2,426,556 | 0.77 | 3,813,299 | 1.06 |
| | **Total** | **267,445,554** | **39.67** | **116,640,119** | **36.99** | **150,805,435** | **42.02** |
| | DNA TE | 252,201,434 | 22.01 | 154,046,261 | 22.15 | 98,155,173 | 21.76 |
| | LTR Gypsy | 317,756,766 | 27.71 | 213,202,044 | 30.66 | 104,554,722 | 23.17 |
| ***O. longiglumis*** | LTR Copia | 78,496,860 | 6.84 | 51,604,619 | 7.42 | 26,892,241 | 5.96 |
| **HHJJ** | LTR Unknown | 46,324,188 | 4.04 | 30,371,387 | 4.37 | 15,952,801 | 3.53 |
| | Unspecified | 5,271,368 | 0.46 | 3,838,046 | 0.55 | 1,433,322 | 0.32 |
| | **Total** | **700,050,616** | **61.06** | **453,062,357** | **65.15** | **246,988,259** | **54.74** |
| | DNA TE | 244,207,408 | 20.31 | 149,236,913 | 20.48 | 94,970,495 | 20.01 |
| | LTR Gypsy | 341,595,940 | 28.40 | 227,156,462 | 31.19 | 114,439,478 | 24.12 |
| ***O. ridleyi*** | LTR Copia | 72,679,843 | 6.04 | 48,408,536 | 6.65 | 24,271,307 | 5.12 |
| **HHJJ** | LTR Unknown | 75,997,097 | 6.32 | 48,082,141 | 6.60 | 27,914,956 | 5.88 |
| | Unspecified | 5,605,382 | 0.47 | 4,000,373 | 0.55 | 1,605,009 | 0.34 |
| | **Total** | **740,085,670** | **61.54** | **476,884,425** | **65.47** | **263,201,245** | **55.47** |
| | DNA TE | 99,409,809 | 12.59 | - | - | - | - |
| | LTR Gypsy | 334,559,036 | 42.40 | - | - | - | - |
| ***O. meyeriana*** | LTR Copia | 52,717,819 | 6.68 | - | - | - | - |
| **GG** | LTR Unknown | 61,279,226 | 7.77 | - | - | - | - |
| | Unspecified | 2,884,473 | 0.37 | - | - | - | - |
| | **Total** | **550,850,363** | **69.81** | - | - | - | - |

**Supplementary Table 4.** Chromosome and subgenome length bp of the *Oryza* species, total amount of TE content and no

| Species | Chromosome | Chromosome length (bp) | (Sub)genome length (bp) | TE content (bp) | non-TE content (bp) |
|---|---|---|---|---|---|
| *O. malampuzhaensis BBCC* | Chr01BB | 51,433,510 | 439,146,677 | 236,491,870 | 202,654,807 |
| | Chr02BB | 44,723,947 | | | |
| | Chr03BB | 44,121,184 | | | |
| | Chr04BB | 39,289,042 | | | |
| | Chr05BB | 33,942,194 | | | |
| | Chr06BB | 38,945,939 | | | |
| | Chr07BB | 34,467,025 | | | |
| | Chr08BB | 32,924,554 | | | |
| | Chr09BB | 28,087,883 | | | |
| | Chr10BB | 29,239,299 | | | |
| | Chr11BB | 32,265,118 | | | |
| | Chr12BB | 29,706,982 | | | |
| | Chr01CC | 56,907,141 | 495,583,665 | 282,106,536 | 213,477,129 |
| | Chr02CC | 48,783,732 | | | |
| | Chr03CC | 51,132,304 | | | |
| | Chr04CC | 42,346,194 | | | |
| | Chr05CC | 39,648,836 | | | |
| | Chr06CC | 41,902,448 | | | |
| | Chr07CC | 38,919,005 | | | |
| | Chr08CC | 36,954,940 | | | |
| | Chr09CC | 30,487,973 | | | |
| | Chr10CC | 35,350,819 | | | |
| | Chr11CC | 40,556,046 | | | |
| | Chr12CC | 32,594,227 | | | |
| *O. minuta BBCC* | Chr01BB | 53,443,419 | 458,169,213 | 256,102,246 | 202,066,967 |
| | Chr02BB | 46,650,487 | | | |
| | Chr03BB | 45,609,606 | | | |
| | Chr04BB | 39,305,625 | | | |
| | Chr05BB | 36,340,438 | | | |
| | Chr06BB | 41,471,532 | | | |
| | Chr07BB | 35,946,503 | | | |
| | Chr08BB | 34,030,533 | | | |
| | Chr09BB | 29,423,328 | | | |
| | Chr10BB | 31,347,020 | | | |
| | Chr11BB | 33,211,271 | | | |
| | Chr12BB | 31,389,451 | | | |
| | Chr01CC | 59,841,146 | 522,212,561 | 307,046,526 | 215,166,035 |
| | Chr02CC | 51,549,145 | | | |
| | Chr03CC | 53,814,567 | | | |
| | Chr04CC | 45,899,044 | | | |
| | Chr05CC | 39,902,222 | | | |
| | Chr06CC | 45,343,885 | | | |
| | Chr07CC | 41,513,474 | | | |
| | Chr08CC | 38,954,463 | | | |
| | Chr09CC | 31,399,422 | | | |
| | Chr10CC | 36,818,097 | | | |

| | | | | | |
|---|---|---|---|---|---|
| | Chr11CC | 43,410,324 | | | |
| | Chr12CC | 33,766,772 | | | |
| *O. alta CCDD* | Chr01CC | 51,980,881 | 459,126,675 | 249,573,419 | 209,553,256 |
| | Chr02CC | 45,715,131 | | | |
| | Chr03CC | 47,032,460 | | | |
| | Chr04CC | 38,387,894 | | | |
| | Chr05CC | 36,472,194 | | | |
| | Chr06CC | 46,146,386 | | | |
| | Chr07CC | 39,314,619 | | | |
| | Chr08CC | 34,567,762 | | | |
| | Chr09CC | 28,560,818 | | | |
| | Chr10CC | 27,160,941 | | | |
| | Chr11CC | 34,433,237 | | | |
| | Chr12CC | 29,354,352 | | | |
| | Chr01DD | 52,500,904 | 449,890,086 | 258,316,801 | 191,573,285 |
| | Chr02DD | 49,550,911 | | | |
| | Chr03DD | 39,270,264 | | | |
| | Chr04DD | 35,312,625 | | | |
| | Chr05DD | 38,358,089 | | | |
| | Chr06DD | 41,451,446 | | | |
| | Chr07DD | 38,419,420 | | | |
| | Chr08DD | 31,952,016 | | | |
| | Chr09DD | 28,725,978 | | | |
| | Chr10DD | 30,454,639 | | | |
| | Chr11DD | 33,316,366 | | | |
| | Chr12DD | 30,577,428 | | | |
| *O. grandiglumis CCDD* | Chr01CC | 49,488,190 | 441,222,684 | 231,503,000 | 209,719,684 |
| | Chr02CC | 46,466,481 | | | |
| | Chr03CC | 45,158,914 | | | |
| | Chr04CC | 37,534,706 | | | |
| | Chr05CC | 35,136,762 | | | |
| | Chr06CC | 42,838,117 | | | |
| | Chr07CC | 38,119,252 | | | |
| | Chr08CC | 32,600,130 | | | |
| | Chr09CC | 27,190,549 | | | |
| | Chr10CC | 26,011,040 | | | |
| | Chr11CC | 32,922,160 | | | |
| | Chr12CC | 27,756,383 | | | |
| | Chr01DD | 49,069,858 | 429,956,128 | 238,849,333 | 191,106,795 |
| | Chr02DD | 44,215,309 | | | |
| | Chr03DD | 39,006,132 | | | |
| | Chr04DD | 32,685,806 | | | |
| | Chr05DD | 37,029,269 | | | |
| | Chr06DD | 38,908,841 | | | |
| | Chr07DD | 37,820,362 | | | |
| | Chr08DD | 32,511,845 | | | |
| | Chr09DD | 27,460,625 | | | |
| | Chr10DD | 29,196,643 | | | |
| | Chr11DD | 31,514,035 | | | |
| | Chr12DD | 30,537,403 | | | |

| | | | | | |
|---|---|---|---|---|---|
| *O. latifolia CCDD* | Chr01CC | 61,078,200 | 543,110,172 | 323,760,018 | 219,350,154 |
| | Chr02CC | 53,068,387 | | | |
| | Chr03CC | 56,361,507 | | | |
| | Chr04CC | 47,064,164 | | | |
| | Chr05CC | 41,517,217 | | | |
| | Chr06CC | 45,305,489 | | | |
| | Chr07CC | 47,547,828 | | | |
| | Chr08CC | 39,549,717 | | | |
| | Chr09CC | 35,131,561 | | | |
| | Chr10CC | 35,547,642 | | | |
| | Chr11CC | 46,822,374 | | | |
| | Chr12CC | 34,116,086 | | | |
| | Chr01DD | 62,437,566 | 515,284,943 | 303,793,548 | 211,491,395 |
| | Chr02DD | 55,375,680 | | | |
| | Chr03DD | 53,092,564 | | | |
| | Chr04DD | 41,536,542 | | | |
| | Chr05DD | 42,603,481 | | | |
| | Chr06DD | 45,332,387 | | | |
| | Chr07DD | 44,070,911 | | | |
| | Chr08DD | 35,362,949 | | | |
| | Chr09DD | 29,709,973 | | | |
| | Chr10DD | 32,329,327 | | | |
| | Chr11DD | 39,301,476 | | | |
| | Chr12DD | 34,132,087 | | | |
| *O. australiensis EE* | Chr01EE | 95,704,982 | 881,424,134 | 631,535,275 | 249,888,859 |
| | Chr02EE | 99,411,906 | | | |
| | Chr03EE | 85,719,274 | | | |
| | Chr04EE | 54,842,428 | | | |
| | Chr05EE | 73,882,514 | | | |
| | Chr06EE | 87,850,232 | | | |
| | Chr07EE | 77,725,253 | | | |
| | Chr08EE | 70,210,424 | | | |
| | Chr09EE | 52,528,534 | | | |
| | Chr10EE | 59,776,721 | | | |
| | Chr11EE | 61,883,842 | | | |
| | Chr12EE | 61,888,024 | | | |
| *O. coarctata KKLL* | Chr01KK | 37,403,316 | 284,342,476 | 92,053,197 | 192,289,279 |
| | Chr02KK | 32,025,281 | | | |
| | Chr03KK | 33,301,484 | | | |
| | Chr04KK | 22,579,943 | | | |
| | Chr05KK | 23,299,831 | | | |
| | Chr06KK | 23,978,878 | | | |
| | Chr07KK | 22,057,888 | | | |
| | Chr08KK | 19,697,393 | | | |
| | Chr09KK | 17,708,968 | | | |
| | Chr10KK | 16,507,214 | | | |
| | Chr11KK | 18,132,009 | | | |
| | Chr12KK | 17,650,271 | | | |
| | Chr01LL | 34,559,743 | | | |
| | Chr02LL | 30,037,761 | | | |

| | | | | | |
|---|---|---|---|---|---|
| | Chr03LL | 31,871,245 | 271,370,639 | 82,369,120 | 189,001,519 |
| | Chr04LL | 22,966,623 | | | |
| | Chr05LL | 23,723,593 | | | |
| | Chr06LL | 22,879,443 | | | |
| | Chr07LL | 21,351,436 | | | |
| | Chr08LL | 20,218,890 | | | |
| | Chr09LL | 16,154,592 | | | |
| | Chr10LL | 15,049,904 | | | |
| | Chr11LL | 15,286,362 | | | |
| | Chr12LL | 17,271,047 | | | |
| *O. schlechteri HHKK* | Chr01HH | 39,288,230 | 315,466,884 | 116,640,119 | 198,826,765 |
| | Chr02HH | 32,772,114 | | | |
| | Chr03HH | 32,192,824 | | | |
| | Chr04HH | 27,451,360 | | | |
| | Chr05HH | 25,457,751 | | | |
| | Chr06HH | 27,519,638 | | | |
| | Chr07HH | 25,910,478 | | | |
| | Chr08HH | 24,702,924 | | | |
| | Chr09HH | 19,395,129 | | | |
| | Chr10HH | 18,931,908 | | | |
| | Chr11HH | 21,033,303 | | | |
| | Chr12HH | 20,811,225 | | | |
| | Chr01KK | 44,874,505 | 358,884,113 | 150,805,435 | 208,078,678 |
| | Chr02KK | 38,093,980 | | | |
| | Chr03KK | 39,114,924 | | | |
| | Chr04KK | 31,920,337 | | | |
| | Chr05KK | 29,681,401 | | | |
| | Chr06KK | 30,695,947 | | | |
| | Chr07KK | 28,204,038 | | | |
| | Chr08KK | 26,285,975 | | | |
| | Chr09KK | 20,321,454 | | | |
| | Chr10KK | 23,334,132 | | | |
| | Chr11KK | 24,102,792 | | | |
| | Chr12KK | 22,254,628 | | | |
| *O. longiglumis HHJJ* | Chr01HH | 90,717,158 | 695,455,420 | 453,062,357 | 242,393,063 |
| | Chr02HH | 70,843,932 | | | |
| | Chr03HH | 80,212,505 | | | |
| | Chr04HH | 63,285,260 | | | |
| | Chr05HH | 56,117,850 | | | |
| | Chr06HH | 61,490,705 | | | |
| | Chr07HH | 51,173,885 | | | |
| | Chr08HH | 52,702,290 | | | |
| | Chr09HH | 37,741,570 | | | |
| | Chr10HH | 41,175,777 | | | |
| | Chr11HH | 49,389,483 | | | |
| | Chr12HH | 40,605,005 | | | |
| | Chr01JJ | 58,173,238 | | | |
| | Chr02JJ | 46,449,883 | | | |
| | Chr03JJ | 47,392,436 | | | |
| | Chr04JJ | 40,113,981 | | | |

| | | | | | |
|---|---|---|---|---|---|
| | Chr05JJ | 35,087,737 | 451,321,844 | 246,988,259 | 204,333,585 |
| | Chr06JJ | 37,213,842 | | | |
| | Chr07JJ | 35,818,993 | | | |
| | Chr08JJ | 31,971,099 | | | |
| | Chr09JJ | 26,815,775 | | | |
| | Chr10JJ | 29,453,885 | | | |
| | Chr11JJ | 32,701,311 | | | |
| | Chr12JJ | 30,129,664 | | | |
| *O. ridleyi HHJJ* | Chr01HH | 93,211,884 | 728,355,013 | 476,884,425 | 251,470,588 |
| | Chr02HH | 77,422,302 | | | |
| | Chr03HH | 82,065,610 | | | |
| | Chr04HH | 61,031,230 | | | |
| | Chr05HH | 58,542,548 | | | |
| | Chr06HH | 62,970,374 | | | |
| | Chr07HH | 53,837,027 | | | |
| | Chr08HH | 55,831,168 | | | |
| | Chr09HH | 44,960,517 | | | |
| | Chr10HH | 44,217,715 | | | |
| | Chr11HH | 52,360,653 | | | |
| | Chr12HH | 41,903,985 | | | |
| | Chr01JJ | 60,565,028 | 474,450,680 | 263,201,245 | 211,249,435 |
| | Chr02JJ | 47,917,918 | | | |
| | Chr03JJ | 49,342,092 | | | |
| | Chr04JJ | 41,873,402 | | | |
| | Chr05JJ | 36,411,554 | | | |
| | Chr06JJ | 39,763,614 | | | |
| | Chr07JJ | 37,888,564 | | | |
| | Chr08JJ | 33,859,925 | | | |
| | Chr09JJ | 29,331,480 | | | |
| | Chr10JJ | 31,175,818 | | | |
| | Chr11JJ | 34,359,842 | | | |
| | Chr12JJ | 31,961,443 | | | |
| *O. meyeriana GG* | Chr01GG | 94,529,614 | 789,124,293 | 550,850,363 | 238,273,930 |
| | Chr02GG | 88,317,990 | | | |
| | Chr03GG | 82,639,008 | | | |
| | Chr04GG | 55,838,731 | | | |
| | Chr05GG | 67,122,205 | | | |
| | Chr06GG | 66,879,100 | | | |
| | Chr07GG | 62,512,216 | | | |
| | Chr08GG | 60,543,954 | | | |
| | Chr09GG | 44,311,931 | | | |
| | Chr10GG | 50,489,843 | | | |
| | Chr11GG | 63,604,183 | | | |
| | Chr12GG | 52,335,518 | | | |

**Supplementary Figure 5.** GenBank accessions of proteome sequences and number of the longest isoforms used in GENESPACE analysis. Proteome sequences of the species under study are shown in bold.

| | Species | Genome Type | NCBI GenBank assembly | NCBI GenBank BioProject | Source | # Proteins (longest isoform) | Reference (DOI or GenBank Link) |
|---|---|---|---|---|---|---|---|
| 1 | *O. sativa japonica* Nipponbare (IRGSP v1.0) | AA | GCA_001433935.1 | PRJDB1747 | https://ftp.gramene.org/oryza/release-7/fasta/oryza_sativa/pep | 35,126 | 10.1186/1939-8433-6-4 |
| 2 | *O. glaberrima* | AA | GCA_000147395.3 | PRJNA13765 | https://ftp.gramene.org/oryza/release-7/fasta/oryza_glaberrima/pep | 40,508 | GCA_000147395 |
| 3 | *O. barthii* | AA | GCA_000182155.4 | PRJNA30379 | https://ftp.gramene.org/oryza/release-7/fasta/oryza_barthii/pep | 40,477 | GCA_000182155 |
| 4 | *O. glumipatula* | AA | GCA_000576495.2 | PRJNA48429 | https://ftp.gramene.org/oryza/release-7/fasta/oryza_glumaepatula/pep | 41,093 | GCA_000576495 |
| 5 | *O. meridionalis* | AA | GCA_000338895.3 | PRJNA48433 | https://ftp.gramene.org/oryza/release-7/fasta/oryza_meridionalis/pep | 41,736 | GCA_000338895 |
| 6 | *O. nivara* | AA | GCA_000576065.2 | PRJNA48107 | https://ftp.ensemblgenomes.ebi.ac.uk/pub/plants/release-57/fasta/oryza_nivara/pep | 36,313 | 10.1101/gr.3766306 |
| 7 | *O. rufipogon* | AA | GCA_000817225.2 | PRJEB4137 | https://ftp.gramene.org/oryza/release-7/fasta/oryza_rufipogon/pep | 37,071 | GCA_000817225 |
| 8 | *O. punctata* | BB | GCA_000573905.2 | PRJNA13770 | https://ftp.gramene.org/oryza/release-7/fasta/oryza_punctata/pep | 35,021 | GCA_000573905 |
| 9 | *O. officinalis* | CC | GCA_008326285.1 | PRJDB2223 | gene prediction made using Augustus on softmasked assembly from Shenton et al., 2019 https://www.ncbi.nlm.nih.gov/datasets/genome/GCA_008326285.1/ | 52,630 | 10.1093/gbe/evaa037 |
| **10** | ***O. malampuzhaensis*** | **BBCC** | **This study** | PRJNA757598 | gene prediction made using Augustus on in-house softmasked assembly | **77,869** | - |
| **11** | ***O. minuta*** | **BBCC** | **This study** | PRJNA757599 | gene prediction made using Augustus on in-house softmasked assembly | **74,265** | - |
| **12** | ***O. alta*** | **CCDD** | **This study** | PRJNA1039467 | gene prediction made using Augustus on in-house softmasked assembly | **66,387** | - |
| **13** | ***O. grandiglumis*** | **CCDD** | **This study** | PRJNA737282 | gene prediction made using Augustus on in-house softmasked assembly | **66,356** | - |
| **14** | ***O. latifolia*** | **CCDD** | **This study** | PRJNA737486 | gene prediction made using Augustus on in-house softmasked assembly | **73,734** | - |
| **15** | ***O. australiensis*** | **EE** | **This study** | PRJNA591699 | gene prediction made using Augustus, SNAP, and fgenesh+ on in-house softmasked genome assembl | **39,392** | - |
| **16** | ***O. coarctata*** | **KKLL** | **This study** | PRJNA439330 | gene prediction made using Augustus on in-house softmasked assembly | **55,709** | - |
| **17** | ***O. schlechteri*** | **HHKK** | **This study** | PRJNA732115 | gene prediction made using Augustus on in-house softmasked assembly | **70,852** | - |
| **18** | ***O. longiglumis*** | **HHJJ** | **This study** | PRJNA1016142 | gene prediction made using Augustus on in-house softmasked assembly | **73,324** | - |
| **19** | ***O. ridleyi*** | **HHJJ** | **This study** | PRJNA687623 | gene prediction made using Augustus on in-house softmasked assembly | **82,806** | - |
| 20 | *O. brachyantha* | FF | GCA_000231095.3 | PRJNA70533 | https://ftp.gramene.org/oryza/release-7/fasta/oryza_brachyantha/pep | 29,766 | 10.1038/ncomms2596 |
| **21** | ***O. meyeriana*** | **GG** | **This study** | PRJNA1039468 | gene prediction made using Augustus on in-house softmasked assembly | **45,092** | - |
| 22 | *L. perrieri* | outgroup | GCA_000325765.3 | PRJNA163065 | https://ftp.ensemblgenomes.ebi.ac.uk/pub/plants/release-57/fasta/leersia_perrieri/pep | 29,078 | GCA_000325765 |

**Supplementary Table 6.** List of genes in *O. alta* and *O. grandiglumis* tha
The homologous *O. sativa* genes are also reported.

| Species | gene ID | Chromosome | Gene Start | Gene End |
|---|---|---|---|---|
| *O. alta* | g2 | Chr1CC | 58591 | 59808 |
| *O. alta* | g16857 | Chr3CC | 384015 | 385070 |
| *O. sativa* | Os03g0105300 | Chr3 | 321593 | 324469 |
| *O. alta* | g9 | Chr1CC | 483701 | 484107 |
| *O. alta* | g16940 | Chr3CC | 1110965 | 1112198 |
| *O. sativa* | Os03g0116316 | Chr3 | 925852 | 926283 |
| *O. alta* | g16941 | Chr3CC | 1113261 | 1115329 |
| *O. alta* | g10 | Chr1CC | 486275 | 488369 |
| *O. sativa* | Os03g0116300 | Chr3 | 922684 | 928190 |
| *O. alta* | g11 | Chr1CC | 491645 | 494111 |
| *O. alta* | g16942 | Chr3CC | 1123469 | 1125634 |
| *O. sativa* | Os03g0116400 | Chr3 | 932241 | 934775 |
| *O. alta* | g12 | Chr1CC | 494860 | 499560 |
| *O. alta* | g16943 | Chr3CC | 1126379 | 1130508 |
| *O. sativa* | Os03g0116500 | Chr3 | 935129 | 939195 |
| *O. alta* | g13 | Chr1CC | 504232 | 505717 |
| *O. alta* | g16944 | Chr3CC | 1155209 | 1156830 |
| *O. sativa* | Os03g0116700 | Chr3 | 942736 | 944993 |
| *O. alta* | g14 | Chr1CC | 507133 | 510179 |
| *O. alta* | g16945 | Chr3CC | 1158279 | 1161145 |
| *O. sativa* | Os03g0116800 | Chr3 | 946032 | 948981 |
| *O. alta* | g15 | Chr1CC | 517340 | 519028 |
| *O. alta* | g16947 | Chr3CC | 1167831 | 1169519 |
| *O. sativa* | Os03g0116900 | Chr3 | 949797 | 957750 |
| *O. alta* | g16 | Chr1CC | 519897 | 522153 |
| *O. alta* | g16948 | Chr3CC | 1170603 | 1172311 |
| *O. sativa* | Os03g0117100 | Chr3 | 958129 | 960101 |
| *O. alta* | g22 | Chr1CC | 551770 | 552339 |
| *O. alta* | g16950 | Chr3CC | 1182294 | 1182863 |
| *O. alta* | g24 | Chr1CC | 557630 | 561614 |
| *O. alta* | g16952 | Chr3CC | 1187148 | 1190881 |

| | | | | |
|---|---|---|---|---|
| *O. sativa* | Os03g0118100 | Chr3 | 985771 | 989760 |
| *O. alta* | g25 | Chr1CC | 569249 | 572485 |
| *O. alta* | g16953 | Chr3CC | 1192854 | 1195666 |
| *O. sativa* | Os03g0118200 | Chr3 | 991125 | 994700 |
| *O. alta* | g26 | Chr1CC | 577045 | 580112 |
| *O. alta* | g16954 | Chr3CC | 1201199 | 1204318 |
| *O. alta* | g28 | Chr1CC | 584077 | 586338 |
| *O. alta* | g16959 | Chr3CC | 1221586 | 1226005 |
| *O. sativa* | Os03g0118600 | Chr3 | 1004328 | 1007269 |
| *O. alta* | g27 | Chr1CC | 581692 | 582060 |
| *O. alta* | g16955 | Chr3CC | 1207632 | 1207988 |
| *O. alta* | g32 | Chr1CC | 633722 | 634754 |
| *O. alta* | g16960 | Chr3CC | 1229224 | 1231382 |
| *O. sativa* | Os03g0118700 | Chr3 | 1008493 | 1010511 |
| *O. alta* | g35 | Chr1CC | 647978 | 650317 |
| *O. alta* | g16962 | Chr3CC | 1239438 | 1241739 |
| *O. sativa* | Os03g0118900 | Chr3 | 1021458 | 1024531 |
| *O. alta* | g36 | Chr1CC | 651325 | 652916 |
| *O. alta* | g16963 | Chr3CC | 1242751 | 1244316 |
| *O. sativa* | Os03g0119000 | Chr3 | 1024732 | 1026761 |
| *O. alta* | g37 | Chr1CC | 653684 | 657734 |
| *O. alta* | g16964 | Chr3CC | 1245075 | 1248751 |
| *O. sativa* | Os03g0119100 | Chr3 | 1027590 | 1031440 |
| *O. alta* | g38 | Chr1CC | 660361 | 662622 |
| *O. alta* | g16965 | Chr3CC | 1251396 | 1253645 |
| *O. sativa* | Os03g0119300 | Chr3 | 1034068 | 1036671 |
| *O. alta* | g48 | Chr1CC | 710055 | 729677 |
| *O. alta* | g16966 | Chr3CC | 1288584 | 1325285 |
| *O. sativa* | Os03g0119500 | Chr3 | 1049042 | 1075622 |
| *O. alta* | g49 | Chr1CC | 796247 | 797527 |
| *O. alta* | g16967 | Chr3CC | 1352111 | 1353376 |
| *O. sativa* | Os03g0119700 | Chr3 | 1076133 | 1079259 |
| *O. alta* | g60 | Chr1CC | 879079 | 883845 |
| *O. alta* | g16969 | Chr3CC | 1366552 | 1372290 |

| | | | | |
|---|---|---|---|---|
| *O. alta* | g61 | Chr1CC | 886754 | 890770 |
| *O. alta* | g16970 | Chr3CC | 1375203 | 1379412 |
| *O. sativa* | Os03g0119966 | Chr3 | 1089453 | 1093410 |
| *O. alta* | g62 | Chr1CC | 895963 | 898866 |
| *O. alta* | g16972 | Chr3CC | 1382063 | 1385566 |
| *O. sativa* | Os03g0120000 | Chr3 | 1099335 | 1102869 |
| *O. alta* | g63 | Chr1CC | 899578 | 901839 |
| *O. alta* | g16973 | Chr3CC | 1388268 | 1390523 |
| *O. sativa* | Os03g0120100 | Chr3 | 1102854 | 1106636 |
| *O. alta* | g64 | Chr1CC | 904694 | 904951 |
| *O. alta* | g16974 | Chr3CC | 1391726 | 1391983 |
| *O. alta* | g65 | Chr1CC | 905663 | 909129 |
| *O. alta* | g16975 | Chr3CC | 1392671 | 1396589 |
| *O. sativa* | Os03g0120200 | Chr3 | 1109242 | 1113045 |
| *O. alta* | g66 | Chr1CC | 915438 | 918851 |
| *O. alta* | g16976 | Chr3CC | 1404335 | 1406735 |
| *O. sativa* | Os03g0120300 | Chr3 | 1119802 | 1123751 |
| *O. alta* | g67 | Chr1CC | 928169 | 929387 |
| *O. alta* | g16977 | Chr3CC | 1408793 | 1410040 |
| *O. sativa* | Os03g0120400 | Chr3 | 1124982 | 1127056 |
| *O. alta* | g68 | Chr1CC | 946754 | 948473 |
| *O. alta* | g16978 | Chr3CC | 1421434 | 1423876 |
| *O. sativa* | Os03g0120800 | Chr3 | 1137129 | 1141062 |
| *O. alta* | g69 | Chr1CC | 961334 | 962245 |
| *O. alta* | g16979 | Chr3CC | 1443439 | 1444383 |
| *O. sativa* | Os03g0120900 | Chr3 | 1152918 | 1154697 |
| *O. alta* | g70 | Chr1CC | 1021670 | 1025765 |
| *O. alta* | g16980 | Chr3CC | 1472844 | 1476263 |
| *O. sativa* | Os03g0121200 | Chr3 | 1172250 | 1176178 |
| *O. alta* | g73 | Chr1CC | 1044741 | 1045830 |
| *O. alta* | g16983 | Chr3CC | 1484089 | 1485182 |
| *O. sativa* | Os03g0121300 | Chr3 | 1178482 | 1180131 |
| *O. alta* | g72 | Chr1CC | 1042769 | 1043959 |
| *O. alta* | g16982 | Chr3CC | 1481909 | 1483336 |
| *O. alta* | g75 | Chr1CC | 1057351 | 1066200 |

| | | | | |
|---|---|---|---|---|
| *O. alta* | g16985 | Chr3CC | 1496500 | 1505583 |
| *O. sativa* | Os03g0121800 | Chr3 | 1195075 | 1204839 |
| *O. alta* | g74 | Chr1CC | 1055793 | 1056774 |
| *O. alta* | g16984 | Chr3CC | 1494811 | 1495807 |
| *O. sativa* | Os03g0121700 | Chr3 | 1194347 | 1195744 |
| *O. alta* | g77 | Chr1CC | 1093087 | 1095846 |
| *O. alta* | g16989 | Chr3CC | 1565380 | 1568142 |
| *O. sativa* | Os03g0122100 | Chr3 | 1226308 | 1229550 |
| *O. alta* | g78 | Chr1CC | 1101917 | 1103833 |
| *O. alta* | g16990 | Chr3CC | 1571783 | 1573860 |
| *O. sativa* | Os03g0122200 | Chr3 | 1232247 | 1234438 |
| *O. alta* | g79 | Chr1CC | 1104473 | 1109763 |
| *O. alta* | g16991 | Chr3CC | 1574559 | 1579008 |
| *O. sativa* | Os03g0122300 | Chr3 | 1235460 | 1244932 |
| *O. alta* | g80 | Chr1CC | 1134314 | 1135307 |
| *O. alta* | g16992 | Chr3CC | 1599397 | 1599810 |
| *O. sativa* | Os03g0122600 | Chr3 | 1270320 | 1300273 |
| *O. alta* | g82 | Chr1CC | 1194347 | 1200073 |
| *O. alta* | g16996 | Chr3CC | 1724698 | 1726089 |
| *O. sativa* | Os03g0123200 | Chr3 | 1322549 | 1324314 |
| *O. alta* | g83 | Chr1CC | 1202383 | 1205527 |
| *O. alta* | g16997 | Chr3CC | 1728412 | 1731575 |
| *O. sativa* | Os03g0123300 | Chr3 | 1327450 | 1331022 |
| *O. alta* | g85 | Chr1CC | 1217399 | 1221702 |
| *O. alta* | g16998 | Chr3CC | 1747375 | 1753425 |
| *O. sativa* | Os03g0123500 | Chr3 | 1340067 | 1345240 |
| *O. alta* | g17000 | Chr3CC | 1762436 | 1764197 |
| *O. alta* | g89 | Chr1CC | 1246977 | 1247793 |
| *O. sativa* | Os03g0123800 | Chr3 | 1355516 | 1358037 |
| *O. alta* | g17001 | Chr3CC | 1789895 | 1791554 |
| *O. alta* | g90 | Chr1CC | 1270084 | 1274443 |
| *O. sativa* | Os03g0124000 | Chr3 | 1394429 | 1400133 |
| *O. alta* | g17004 | Chr3CC | 1806384 | 1807669 |
| *O. alta* | g91 | Chr1CC | 1280101 | 1281754 |
| *O. sativa* | Os03g0124100 | Chr3 | 1402828 | 1404996 |

| | | | | |
|---|---|---|---|---|
| *O. alta* | g17006 | Chr3CC | 1812704 | 1815262 |
| *O. alta* | g92 | Chr1CC | 1292503 | 1295055 |
| *O. sativa* | Os03g0124200 | Chr3 | 1406065 | 1408803 |
| *O. alta* | g17007 | Chr3CC | 1818674 | 1820095 |
| *O. alta* | g94 | Chr1CC | 1301690 | 1303105 |
| *O. sativa* | Os03g0124300 | Chr3 | 1410020 | 1411723 |
| *O. alta* | g17008 | Chr3CC | 1826789 | 1827589 |
| *O. alta* | g95 | Chr1CC | 1310239 | 1311134 |
| *O. sativa* | Os03g0124500 | Chr3 | 1419786 | 1420939 |
| *O. alta* | g17010 | Chr3CC | 1836672 | 1839406 |
| *O. alta* | g96 | Chr1CC | 1324700 | 1327413 |
| *O. sativa* | Os03g0124900 | Chr3 | 1430583 | 1433728 |
| *O. alta* | g17011 | Chr3CC | 1839749 | 1841124 |
| *O. alta* | g97 | Chr1CC | 1327736 | 1329102 |
| *O. sativa* | Os03g0125000 | Chr3 | 1433749 | 1435315 |
| *O. alta* | g17012 | Chr3CC | 1842568 | 1844645 |
| *O. alta* | g98 | Chr1CC | 1330622 | 1332706 |
| *O. sativa* | Os03g0125100 | Chr3 | 1436356 | 1439055 |
| *O. alta* | g17013 | Chr3CC | 1850925 | 1854174 |
| *O. alta* | g99 | Chr1CC | 1338383 | 1341838 |
| *O. sativa* | Os03g0125300 | Chr3 | 1444996 | 1448651 |
| *O. alta* | g17014 | Chr3CC | 1854857 | 1857512 |
| *O. alta* | g100 | Chr1CC | 1342534 | 1345194 |
| *O. sativa* | Os03g0125400 | Chr3 | 1449071 | 1451951 |
| *O. alta* | g101 | Chr1CC | 1352042 | 1355507 |
| *O. alta* | g17015 | Chr3CC | 1866025 | 1869513 |
| *O. sativa* | Os03g0125600 | Chr3 | 1457515 | 1461362 |
| *O. alta* | g103 | Chr1CC | 1368359 | 1373810 |
| *O. alta* | g17016 | Chr3CC | 1880938 | 1885883 |
| *O. sativa* | Os03g0125800 | Chr3 | 1476335 | 1483349 |
| *O. alta* | g104 | Chr1CC | 1375125 | 1376515 |
| *O. alta* | g17017 | Chr3CC | 1887089 | 1888489 |
| *O. sativa* | Os03g0125900 | Chr3 | 1484212 | 1486097 |
| *O. alta* | g105 | Chr1CC | 1377147 | 1381403 |
| *O. alta* | g17018 | Chr3CC | 1889191 | 1893426 |
| *O. sativa* | Os03g0126000 | Chr3 | 1486030 | 1490594 |

| | | | | |
|---|---|---|---|---|
| *O. alta* | g106 | Chr1CC | 1385246 | 1390080 |
| *O. alta* | g17019 | Chr3CC | 1897527 | 1902165 |
| *O. sativa* | Os03g0126100 | Chr3 | 1494308 | 1497245 |
| *O. alta* | g107 | Chr1CC | 1391030 | 1395080 |
| *O. alta* | g17020 | Chr3CC | 1903288 | 1907094 |
| *O. sativa* | Os03g0126300 | Chr3 | 1499619 | 1504235 |
| *O. alta* | g108 | Chr1CC | 1395851 | 1396564 |
| *O. alta* | g17021 | Chr3CC | 1908520 | 1909248 |
| *O. sativa* | Os03g0126450 | Chr3 | 1505340 | 1508525 |
| *O. alta* | g110 | Chr1CC | 1405850 | 1409586 |
| *O. alta* | g17023 | Chr3CC | 1943399 | 1947200 |
| *O. alta* | g112 | Chr1CC | 1422554 | 1422928 |
| *O. alta* | g17024 | Chr3CC | 1951496 | 1951840 |
| *O. sativa* | Os03g0126900 | Chr3 | 1522404 | 1523048 |
| *O. alta* | g113 | Chr1CC | 1425403 | 1426732 |
| *O. alta* | g17025 | Chr3CC | 1954266 | 1955966 |
| *O. sativa* | Os03g0127000 | Chr3 | 1525222 | 1527076 |
| *O. alta* | g114 | Chr1CC | 1428712 | 1431455 |
| *O. alta* | g17026 | Chr3CC | 1958529 | 1960587 |
| *O. alta* | g115 | Chr1CC | 1458829 | 1461211 |
| *O. alta* | g17027 | Chr3CC | 1978082 | 1980681 |
| *O. sativa* | Os03g0127500 | Chr3 | 1551291 | 1554059 |
| *O. alta* | g116 | Chr1CC | 1461919 | 1466735 |
| *O. alta* | g17028 | Chr3CC | 1981286 | 1986024 |
| *O. sativa* | Os03g0127600 | Chr3 | 1554059 | 1559153 |
| *O. alta* | g117 | Chr1CC | 1471595 | 1474370 |
| *O. alta* | g17029 | Chr3CC | 1989223 | 1991983 |
| *O. sativa* | Os03g0127700 | Chr3 | 1562044 | 1566735 |
| *O. alta* | g118 | Chr1CC | 1474855 | 1475601 |
| *O. alta* | g17030 | Chr3CC | 1992473 | 1993713 |
| *O. alta* | g119 | Chr1CC | 1480410 | 1484437 |
| *O. alta* | g17031 | Chr3CC | 1995650 | 2003341 |
| *O. sativa* | Os03g0127900 | Chr3 | 1567984 | 1571080 |
| *O. alta* | g120 | Chr1CC | 1488127 | 1489326 |

| | | | | |
|---|---|---|---|---|
| *O. alta* | g17032 | Chr3CC | 2006701 | 2007915 |
| *O. sativa* | Os03g0128000 | Chr3 | 1579382 | 1580910 |
| *O. alta* | g122 | Chr1CC | 1537289 | 1537863 |
| *O. alta* | g17036 | Chr3CC | 2033166 | 2033754 |
| *O. sativa* | Os03g0128300 | Chr3 | 1608031 | 1609003 |
| *O. alta* | g123 | Chr1CC | 1541178 | 1546842 |
| *O. alta* | g17037 | Chr3CC | 2037171 | 2043508 |
| *O. sativa* | Os03g0128500 | Chr3 | 1612327 | 1616443 |
| *O. alta* | g124 | Chr1CC | 1556527 | 1559736 |
| *O. alta* | g17038 | Chr3CC | 2052305 | 2055360 |
| *O. sativa* | Os03g0128700 | Chr3 | 1625098 | 1629527 |
| *O. alta* | g125 | Chr1CC | 1561078 | 1562071 |
| *O. alta* | g17039 | Chr3CC | 2055631 | 2060132 |
| *O. sativa* | Os03g0128800 | Chr3 | 1630177 | 1631745 |
| *O. alta* | g126 | Chr1CC | 1584692 | 1586983 |
| *O. alta* | g17041 | Chr3CC | 2089488 | 2091715 |
| *O. alta* | g127 | Chr1CC | 1587454 | 1590590 |
| *O. alta* | g17042 | Chr3CC | 2092170 | 2095448 |
| *O. sativa* | Os03g0129100 | Chr3 | 1645164 | 1648764 |
| *O. alta* | g128 | Chr1CC | 1598142 | 1606289 |
| *O. alta* | g17043 | Chr3CC | 2099543 | 2110197 |
| *O. sativa* | Os03g0129200 | Chr3 | 1653394 | 1661497 |
| *O. alta* | g129 | Chr1CC | 1607149 | 1609914 |
| *O. alta* | g17044 | Chr3CC | 2111034 | 2113892 |
| *O. sativa* | Os03g0129300 | Chr3 | 1662306 | 1665491 |
| *O. alta* | g130 | Chr1CC | 1614811 | 1616496 |
| *O. alta* | g17045 | Chr3CC | 2125081 | 2130896 |
| *O. sativa* | Os03g0129800 | Chr3 | 1668201 | 1673176 |
| *O. alta* | g132 | Chr1CC | 1621629 | 1625639 |
| *O. alta* | g17046 | Chr3CC | 2146631 | 2150848 |
| *O. alta* | g133 | Chr1CC | 1637283 | 1637615 |
| *O. alta* | g17047 | Chr3CC | 2163178 | 2163510 |
| *O. sativa* | Os03g0129900 | Chr3 | 1685564 | 1689540 |
| *O. alta* | g134 | Chr1CC | 1674677 | 1676516 |
| *O. alta* | g17048 | Chr3CC | 2183609 | 2185529 |

| | | | | |
|---|---|---|---|---|
| *O. sativa* | Os03g0130100 | Chr3 | 1706053 | 1708135 |
| *O. alta* | g135 | Chr1CC | 1688495 | 1688822 |
| *O. alta* | g17049 | Chr3CC | 2192243 | 2192587 |
| *O. sativa* | Os03g0130300 | Chr3 | 1714575 | 1715192 |
| *O. alta* | g136 | Chr1CC | 1690395 | 1693101 |
| *O. alta* | g17050 | Chr3CC | 2194358 | 2197084 |
| *O. sativa* | Os03g0130400 | Chr3 | 1716495 | 1719628 |
| *O. alta* | g137 | Chr1CC | 1694074 | 1701622 |
| *O. alta* | g17051 | Chr3CC | 2198130 | 2206406 |
| *O. sativa* | Os03g0130500 | Chr3 | 1720072 | 1727907 |
| *O. alta* | g138 | Chr1CC | 1712992 | 1713733 |
| *O. alta* | g17052 | Chr3CC | 2218193 | 2219008 |
| *O. alta* | g139 | Chr1CC | 1738445 | 1749208 |
| *O. alta* | g17053 | Chr3CC | 2220547 | 2231330 |
| *O. sativa* | Os03g0130700 | Chr3 | 1737302 | 1742634 |
| *O. alta* | g140 | Chr1CC | 1752806 | 1758257 |
| *O. alta* | g17054 | Chr3CC | 2238397 | 2243834 |
| *O. sativa* | Os03g0130800 | Chr3 | 1752160 | 1758211 |
| *O. alta* | g141 | Chr1CC | 1778931 | 1782527 |
| *O. alta* | g17056 | Chr3CC | 2254853 | 2258599 |
| *O. sativa* | Os03g0130900 | Chr3 | 1765350 | 1770210 |
| *O. alta* | g142 | Chr1CC | 1784049 | 1786319 |
| *O. alta* | g17057 | Chr3CC | 2260168 | 2262446 |
| *O. sativa* | Os03g0131000 | Chr3 | 1774911 | 1775816 |
| *O. alta* | g143 | Chr1CC | 1787794 | 1791306 |
| *O. alta* | g17058 | Chr3CC | 2263699 | 2267610 |
| *O. sativa* | Os03g0131100 | Chr3 | 1776356 | 1780897 |
| *O. alta* | g144 | Chr1CC | 1801344 | 1804055 |
| *O. alta* | g17059 | Chr3CC | 2273575 | 2275968 |
| *O. sativa* | Os03g0131200 | Chr3 | 1787679 | 1790806 |
| *O. alta* | g145 | Chr1CC | 1805651 | 1810160 |
| *O. alta* | g17060 | Chr3CC | 2277223 | 2281486 |
| *O. sativa* | Os03g0131300 | Chr3 | 1791608 | 1796372 |
| *O. alta* | g146 | Chr1CC | 1810787 | 1815975 |
| *O. alta* | g17061 | Chr3CC | 2282107 | 2287404 |

| | | | | |
|---|---|---|---|---|
| *O. sativa* | Os03g0131400 | Chr3 | 1798724 | 1804320 |
| *O. alta* | g147 | Chr1CC | 1816819 | 1817076 |
| *O. alta* | g17062 | Chr3CC | 2291795 | 2294336 |
| *O. sativa* | Os03g0131500 | Chr3 | 1804488 | 1813229 |
| *O. alta* | g148 | Chr1CC | 1819920 | 1820940 |
| *O. alta* | g17063 | Chr3CC | 2295599 | 2297079 |
| *O. sativa* | Os03g0131950 | Chr3 | 1817620 | 1818990 |
| *O. alta* | g149 | Chr1CC | 1822128 | 1823300 |
| *O. alta* | g17064 | Chr3CC | 2297798 | 2298970 |
| *O. sativa* | Os03g0131900 | Chr3 | 1817533 | 1818855 |
| *O. alta* | g150 | Chr1CC | 1830958 | 1834362 |
| *O. alta* | g17065 | Chr3CC | 2304210 | 2307303 |
| *O. sativa* | Os03g0132000 | Chr3 | 1822255 | 1825976 |
| *O. alta* | g151 | Chr1CC | 1853186 | 1854621 |
| *O. alta* | g17066 | Chr3CC | 2313464 | 2314902 |
| *O. sativa* | Os03g0132200 | Chr3 | 1831332 | 1833035 |
| *O. alta* | g152 | Chr1CC | 1858537 | 1861139 |
| *O. alta* | g17067 | Chr3CC | 2318842 | 2320461 |
| *O. sativa* | Os03g0132300 | Chr3 | 1836607 | 1839772 |
| *O. alta* | g153 | Chr1CC | 1866912 | 1867901 |
| *O. alta* | g17068 | Chr3CC | 2325362 | 2326348 |
| *O. alta* | g155 | Chr1CC | 1885440 | 1889542 |
| *O. alta* | g17069 | Chr3CC | 2337652 | 2343849 |
| *O. sativa* | Os03g0132800 | Chr3 | 1852910 | 1857693 |
| *O. alta* | g157 | Chr1CC | 1898023 | 1898289 |
| *O. alta* | g17070 | Chr3CC | 2346475 | 2347501 |
| *O. sativa* | Os03g0132900 | Chr3 | 1860429 | 1861772 |
| *O. alta* | g158 | Chr1CC | 1905329 | 1906833 |
| *O. alta* | g17071 | Chr3CC | 2363247 | 2364778 |
| *O. sativa* | Os03g0133000 | Chr3 | 1862381 | 1864245 |
| *O. alta* | g159 | Chr1CC | 1916046 | 1916393 |
| *O. alta* | g17073 | Chr3CC | 2378455 | 2378802 |
| *O. sativa* | Os03g0133100 | Chr3 | 1870804 | 1871499 |
| *O. alta* | g160 | Chr1CC | 1919582 | 1923765 |
| *O. alta* | g17074 | Chr3CC | 2381267 | 2385448 |

| | | | | |
|---|---|---|---|---|
| *O. alta* | g162 | Chr1CC | 1927109 | 1928963 |
| *O. alta* | g17077 | Chr3CC | 2395721 | 2401190 |
| *O. sativa* | Os03g0133800 | Chr3 | 1885977 | 1891066 |
| *O. alta* | g164 | Chr1CC | 1936747 | 1947123 |
| *O. alta* | g17078 | Chr3CC | 2404239 | 2408166 |
| *O. sativa* | Os03g0133900 | Chr3 | 1892829 | 1897309 |
| *O. alta* | g166 | Chr1CC | 1949317 | 1953468 |
| *O. alta* | g17079 | Chr3CC | 2410152 | 2414450 |
| *O. sativa* | Os03g0134300 | Chr3 | 1909201 | 1919394 |
| *O. alta* | g167 | Chr1CC | 1954718 | 1956590 |
| *O. alta* | g17080 | Chr3CC | 2415537 | 2417409 |
| *O. sativa* | Os03g0134500 | Chr3 | 1919768 | 1922021 |
| *O. alta* | g168 | Chr1CC | 1988601 | 1989596 |
| *O. alta* | g17081 | Chr3CC | 2429270 | 2430247 |
| *O. sativa* | Os03g0134800 | Chr3 | 1936269 | 1937270 |
| *O. alta* | g169 | Chr1CC | 1990587 | 1991739 |
| *O. alta* | g17082 | Chr3CC | 2430988 | 2432118 |
| *O. sativa* | Os03g0134900 | Chr3 | 1938240 | 1939691 |
| *O. alta* | g170 | Chr1CC | 1994359 | 1995270 |
| *O. alta* | g17083 | Chr3CC | 2435099 | 2436006 |
| *O. sativa* | Os03g0135100 | Chr3 | 1948386 | 1949543 |
| *O. alta* | g172 | Chr1CC | 2000926 | 2001893 |
| *O. alta* | g17085 | Chr3CC | 2440705 | 2441667 |
| *O. sativa* | Os03g0135300 | Chr3 | 1954717 | 1955959 |
| *O. alta* | g173 | Chr1CC | 2002974 | 2005534 |
| *O. alta* | g17086 | Chr3CC | 2442765 | 2445291 |
| *O. sativa* | Os03g0135400 | Chr3 | 1956859 | 1959763 |
| *O. alta* | g174 | Chr1CC | 2074198 | 2076969 |
| *O. alta* | g17087 | Chr3CC | 2455656 | 2458538 |
| *O. sativa* | Os03g0135600 | Chr3 | 1970948 | 1979567 |
| *O. alta* | g175 | Chr1CC | 2082146 | 2083665 |
| *O. alta* | g17090 | Chr3CC | 2468597 | 2470151 |
| *O. sativa* | Os03g0135700 | Chr3 | 1983566 | 1985563 |
| *O. alta* | g177 | Chr1CC | 2100051 | 2101142 |
| *O. alta* | g17092 | Chr3CC | 2489579 | 2490670 |

| | | | | |
|---|---|---|---|---|
| *O. sativa* | Os03g0136200 | Chr3 | 2003246 | 2004830 |
| *O. alta* | g178 | Chr1CC | 2109002 | 2110582 |
| *O. alta* | g17093 | Chr3CC | 2508039 | 2509741 |
| *O. sativa* | Os03g0136400 | Chr3 | 2010988 | 2013532 |
| *O. alta* | g179 | Chr1CC | 2111437 | 2111847 |
| *O. alta* | g17094 | Chr3CC | 2510599 | 2511009 |
| *O. sativa* | Os03g0136500 | Chr3 | 2014048 | 2014743 |
| *O. alta* | g180 | Chr1CC | 2112196 | 2112678 |
| *O. alta* | g17095 | Chr3CC | 2511577 | 2512059 |
| *O. sativa* | Os03g0136600 | Chr3 | 2014957 | 2015602 |
| *O. alta* | g181 | Chr1CC | 2113259 | 2115622 |
| *O. alta* | g17096 | Chr3CC | 2512648 | 2515011 |
| *O. alta* | g182 | Chr1CC | 2116163 | 2118817 |
| *O. alta* | g17097 | Chr3CC | 2515518 | 2518174 |
| *O. sativa* | Os03g0136800 | Chr3 | 2018736 | 2021964 |
| *O. alta* | g183 | Chr1CC | 2120545 | 2126950 |
| *O. alta* | g17098 | Chr3CC | 2519657 | 2526080 |
| *O. sativa* | Os03g0136900 | Chr3 | 2026817 | 2033506 |
| *O. alta* | g184 | Chr1CC | 2141715 | 2144188 |
| *O. alta* | g17100 | Chr3CC | 2541687 | 2544111 |
| *O. sativa* | Os03g0137200 | Chr3 | 2055599 | 2058939 |
| *O. alta* | g185 | Chr1CC | 2145544 | 2148599 |
| *O. alta* | g17101 | Chr3CC | 2545464 | 2548477 |
| *O. sativa* | Os03g0137300 | Chr3 | 2059793 | 2062965 |
| *O. alta* | g186 | Chr1CC | 2153555 | 2155933 |
| *O. alta* | g17102 | Chr3CC | 2550510 | 2552888 |
| *O. sativa* | Os03g0137400 | Chr3 | 2065049 | 2067737 |
| *O. alta* | g188 | Chr1CC | 2160333 | 2162937 |
| *O. alta* | g17103 | Chr3CC | 2554931 | 2558133 |
| *O. sativa* | Os03g0137500 | Chr3 | 2069458 | 2073230 |
| *O. alta* | g189 | Chr1CC | 2164202 | 2165315 |
| *O. alta* | g17104 | Chr3CC | 2559176 | 2560678 |
| *O. sativa* | Os03g0137600 | Chr3 | 2073420 | 2074942 |
| *O. alta* | g190 | Chr1CC | 2167308 | 2168194 |
| *O. alta* | g17105 | Chr3CC | 2563047 | 2563820 |

| | | | | |
|---|---|---|---|---|
| *O. sativa* | Os03g0137700 | Chr3 | 2075322 | 2077636 |
| *O. alta* | g191 | Chr1CC | 2180700 | 2181062 |
| *O. alta* | g17107 | Chr3CC | 2584539 | 2587094 |
| *O. sativa* | Os03g0137800 | Chr3 | 2088673 | 2091654 |
| *O. alta* | g192 | Chr1CC | 2191836 | 2195016 |
| *O. alta* | g17108 | Chr3CC | 2597695 | 2602850 |
| *O. sativa* | Os03g0138000 | Chr3 | 2097607 | 2101999 |
| *O. alta* | g194 | Chr1CC | 2196914 | 2204210 |
| *O. alta* | g17110 | Chr3CC | 2608178 | 2615469 |
| *O. sativa* | Os03g0138100 | Chr3 | 2103191 | 2112148 |
| *O. alta* | g196 | Chr1CC | 2210227 | 2211810 |
| *O. alta* | g17112 | Chr3CC | 2621937 | 2623505 |
| *O. sativa* | Os03g0138200 | Chr3 | 2115174 | 2116896 |
| *O. alta* | g197 | Chr1CC | 2215858 | 2216130 |
| *O. alta* | g17113 | Chr3CC | 2625683 | 2626192 |
| *O. sativa* | Os03g0138400 | Chr3 | 2118865 | 2123085 |
| *O. alta* | g198 | Chr1CC | 2221523 | 2223061 |
| *O. alta* | g17114 | Chr3CC | 2630024 | 2631547 |
| *O. sativa* | Os03g0138500 | Chr3 | 2123692 | 2125645 |
| *O. alta* | g199 | Chr1CC | 2223681 | 2227174 |
| *O. alta* | g17115 | Chr3CC | 2632235 | 2635746 |
| *O. sativa* | Os03g0138600 | Chr3 | 2125787 | 2129505 |
| *O. alta* | g200 | Chr1CC | 2230181 | 2234027 |
| *O. alta* | g17116 | Chr3CC | 2640871 | 2645150 |
| *O. sativa* | Os03g0138700 | Chr3 | 2131982 | 2136725 |
| *O. alta* | g202 | Chr1CC | 2241648 | 2242595 |
| *O. alta* | g17117 | Chr3CC | 2647775 | 2648776 |
| *O. sativa* | Os03g0138900 | Chr3 | 2139916 | 2141093 |
| *O. alta* | g204 | Chr1CC | 2247570 | 2249434 |
| *O. alta* | g17119 | Chr3CC | 2653785 | 2655862 |
| *O. sativa* | Os03g0139200 | Chr3 | 2145765 | 2150519 |
| *O. alta* | g206 | Chr1CC | 2258875 | 2259639 |
| *O. alta* | g17121 | Chr3CC | 2662818 | 2663585 |
| *O. sativa* | Os03g0139400 | Chr3 | 2154645 | 2155581 |
| *O. alta* | g207 | Chr1CC | 2261279 | 2263066 |

| | | | | |
|---|---|---|---|---|
| *O. alta* | g17122 | Chr3CC | 2665217 | 2668700 |
| *O. sativa* | Os03g0139500 | Chr3 | 2159337 | 2161733 |
| *O. alta* | g210 | Chr1CC | 2299687 | 2301213 |
| *O. alta* | g17127 | Chr3CC | 2728565 | 2730091 |
| *O. sativa* | Os03g0140100 | Chr3 | 2199772 | 2201456 |
| *O. alta* | g208 | Chr1CC | 2279810 | 2281363 |
| *O. alta* | g17124 | Chr3CC | 2703319 | 2704593 |
| *O. alta* | g211 | Chr1CC | 2308448 | 2309980 |
| *O. alta* | g17128 | Chr3CC | 2735365 | 2744148 |
| *O. alta* | g214 | Chr1CC | 2329435 | 2331045 |
| *O. alta* | g17129 | Chr3CC | 2752231 | 2753835 |
| *O. sativa* | Os03g0140400 | Chr3 | 2223309 | 2225202 |
| *O. alta* | g215 | Chr1CC | 2332176 | 2334764 |
| *O. alta* | g17130 | Chr3CC | 2754616 | 2757279 |
| *O. sativa* | Os03g0140500 | Chr3 | 2225971 | 2228510 |
| *O. alta* | g216 | Chr1CC | 2345812 | 2347047 |
| *O. alta* | g17131 | Chr3CC | 2767141 | 2768306 |
| *O. sativa* | Os03g0140700 | Chr3 | 2248907 | 2250627 |
| *O. alta* | g217 | Chr1CC | 2355906 | 2358496 |
| *O. alta* | g17132 | Chr3CC | 2775042 | 2777617 |
| *O. sativa* | Os03g0140900 | Chr3 | 2260797 | 2263995 |
| *O. alta* | g218 | Chr1CC | 2359453 | 2361369 |
| *O. alta* | g17133 | Chr3CC | 2778426 | 2779959 |
| *O. sativa* | Os03g0141000 | Chr3 | 2264925 | 2266928 |
| *O. alta* | g219 | Chr1CC | 2364276 | 2367123 |
| *O. alta* | g17134 | Chr3CC | 2790015 | 2792861 |
| *O. sativa* | Os03g0141200 | Chr3 | 2272966 | 2276161 |
| *O. alta* | g220 | Chr1CC | 2384653 | 2389547 |
| *O. alta* | g17136 | Chr3CC | 2810065 | 2810718 |
| *O. sativa* | Os03g0141800 | Chr3 | 2302598 | 2306807 |
| *O. alta* | g221 | Chr1CC | 2392903 | 2396437 |
| *O. alta* | g17138 | Chr3CC | 2819725 | 2822993 |
| *O. sativa* | Os03g0142500 | Chr3 | 2336789 | 2341656 |
| *O. alta* | g222 | Chr1CC | 2397374 | 2398302 |
| *O. alta* | g17139 | Chr3CC | 2823966 | 2824898 |

| | | | | |
|---|---|---|---|---|
| *O. sativa* | Os03g0142600 | Chr3 | 2341955 | 2343111 |
| *O. alta* | g223 | Chr1CC | 2424304 | 2430043 |
| *O. alta* | g17140 | Chr3CC | 2841587 | 2847324 |
| *O. sativa* | Os03g0142800 | Chr3 | 2367856 | 2374437 |
| *O. alta* | g224 | Chr1CC | 2431794 | 2434148 |
| *O. alta* | g17141 | Chr3CC | 2849019 | 2851733 |
| *O. sativa* | Os03g0142900 | Chr3 | 2375216 | 2378035 |
| *O. alta* | g225 | Chr1CC | 2436938 | 2442109 |
| *O. alta* | g17142 | Chr3CC | 2856164 | 2870711 |
| *O. sativa* | Os03g0143000 | Chr3 | 2380055 | 2384722 |
| *O. alta* | g227 | Chr1CC | 2453941 | 2456292 |
| *O. alta* | g17144 | Chr3CC | 2876137 | 2878479 |
| *O. alta* | g228 | Chr1CC | 2469539 | 2472840 |
| *O. alta* | g17146 | Chr3CC | 2886793 | 2890158 |
| *O. alta* | g229 | Chr1CC | 2475958 | 2480914 |
| *O. alta* | g17147 | Chr3CC | 2891028 | 2896079 |
| *O. sativa* | Os03g0143400 | Chr3 | 2401014 | 2406713 |
| *O. alta* | g230 | Chr1CC | 2485629 | 2489318 |
| *O. alta* | g17148 | Chr3CC | 2901792 | 2905574 |
| *O. sativa* | Os03g0143600 | Chr3 | 2409889 | 2413431 |
| *O. alta* | g231 | Chr1CC | 2495502 | 2498746 |
| *O. alta* | g17149 | Chr3CC | 2910614 | 2913881 |
| *O. sativa* | Os03g0143700 | Chr3 | 2415606 | 2419564 |
| *O. alta* | g232 | Chr1CC | 2503693 | 2520113 |
| *O. alta* | g17150 | Chr3CC | 2917474 | 2942378 |
| *O. sativa* | Os03g0143800 | Chr3 | 2432922 | 2448897 |
| *O. alta* | g233 | Chr1CC | 2520648 | 2521472 |
| *O. alta* | g17151 | Chr3CC | 2942903 | 2943733 |
| *O. sativa* | Os03g0143900 | Chr3 | 2449426 | 2450253 |
| *O. alta* | g235 | Chr1CC | 2526568 | 2528439 |
| *O. alta* | g17153 | Chr3CC | 2954088 | 2955003 |
| *O. alta* | g236 | Chr1CC | 2530093 | 2531616 |
| *O. alta* | g17154 | Chr3CC | 2955921 | 2957435 |
| *O. sativa* | Os03g0144300 | Chr3 | 2466460 | 2468340 |

| | | | | |
|---|---|---|---|---|
| *O. alta* | g237 | Chr1CC | 2534633 | 2536180 |
| *O. alta* | g17156 | Chr3CC | 2960315 | 2962066 |
| *O. sativa* | Os03g0144500 | Chr3 | 2471481 | 2473021 |
| *O. alta* | g240 | Chr1CC | 2586073 | 2591099 |
| *O. alta* | g17157 | Chr3CC | 2967541 | 2969355 |
| *O. sativa* | Os03g0144800 | Chr3 | 2486920 | 2489480 |
| *O. alta* | g242 | Chr1CC | 2602358 | 2605459 |
| *O. alta* | g17159 | Chr3CC | 2974830 | 2977940 |
| *O. sativa* | Os03g0145102 | Chr3 | 2494796 | 2498898 |
| *O. alta* | g245 | Chr1CC | 2612865 | 2613625 |
| *O. alta* | g17160 | Chr3CC | 2984290 | 2985027 |
| *O. sativa* | Os03g0145200 | Chr3 | 2506064 | 2507170 |
| *O. alta* | g246 | Chr1CC | 2616892 | 2617518 |
| *O. alta* | g17161 | Chr3CC | 2987526 | 2988149 |
| *O. sativa* | Os03g0145300 | Chr3 | 2512872 | 2516234 |
| *O. alta* | g247 | Chr1CC | 2620406 | 2622559 |
| *O. alta* | g17163 | Chr3CC | 2991001 | 2993154 |
| *O. alta* | g248 | Chr1CC | 2623964 | 2631733 |
| *O. alta* | g17164 | Chr3CC | 2994627 | 3002362 |
| *O. sativa* | Os03g0145500 | Chr3 | 2516859 | 2525960 |
| *O. alta* | g252 | Chr1CC | 2637319 | 2639740 |
| *O. alta* | g17167 | Chr3CC | 3014857 | 3016937 |
| *O. sativa* | Os03g0145800 | Chr3 | 2543680 | 2546338 |
| *O. alta* | g253 | Chr1CC | 2648431 | 2653188 |
| *O. alta* | g17170 | Chr3CC | 3045498 | 3050896 |
| *O. sativa* | Os03g0146000 | Chr3 | 2553889 | 2559108 |
| *O. alta* | g254 | Chr1CC | 2656912 | 2658441 |
| *O. alta* | g17171 | Chr3CC | 3062360 | 3063858 |
| *O. sativa* | Os03g0146100 | Chr3 | 2562812 | 2564646 |
| *O. alta* | g256 | Chr1CC | 2667576 | 2670849 |
| *O. alta* | g17173 | Chr3CC | 3076125 | 3079600 |
| *O. sativa* | Os03g0146400 | Chr3 | 2576609 | 2580404 |
| *O. alta* | g257 | Chr1CC | 2698838 | 2700721 |
| *O. alta* | g17174 | Chr3CC | 3080983 | 3082365 |
| *O. sativa* | Os03g0146500 | Chr3 | 2581186 | 2583486 |

| | | | | |
|---|---|---|---|---|
| *O. alta* | g258 | Chr1CC | 2701995 | 2722911 |
| *O. alta* | g17176 | Chr3CC | 3093460 | 3105574 |
| *O. sativa* | Os03g0146600 | Chr3 | 2605032 | 2605475 |
| *O. alta* | g260 | Chr1CC | 2737894 | 2738641 |
| *O. alta* | g17177 | Chr3CC | 3109470 | 3111237 |
| *O. sativa* | Os03g0147001 | Chr3 | 2613684 | 2615443 |
| *O. alta* | g267 | Chr1CC | 2814158 | 2814929 |
| *O. alta* | g17178 | Chr3CC | 3112961 | 3113719 |
| *O. alta* | g261 | Chr1CC | 2754776 | 2756699 |
| *O. alta* | g17179 | Chr3CC | 3116872 | 3118748 |
| *O. sativa* | Os03g0147400 | Chr3 | 2651951 | 2661701 |
| *O. alta* | g263 | Chr1CC | 2775070 | 2778535 |
| *O. alta* | g17181 | Chr3CC | 3136778 | 3142297 |
| *O. sativa* | Os03g0147700 | Chr3 | 2677259 | 2682146 |
| *O. alta* | g265 | Chr1CC | 2787409 | 2789190 |
| *O. alta* | g17182 | Chr3CC | 3147237 | 3148500 |
| *O. sativa* | Os03g0148000 | Chr3 | 2685582 | 2688291 |
| *O. alta* | g266 | Chr1CC | 2801899 | 2803169 |
| *O. alta* | g17183 | Chr3CC | 3153588 | 3154502 |
| *O. sativa* | Os03g0148300 | Chr3 | 2695582 | 2696627 |
| *O. alta* | g268 | Chr1CC | 2818428 | 2819723 |
| *O. alta* | g17184 | Chr3CC | 3156214 | 3157506 |
| *O. sativa* | Os03g0148400 | Chr3 | 2698774 | 2700402 |
| *O. alta* | g270 | Chr1CC | 2832496 | 2839447 |
| *O. alta* | g17186 | Chr3CC | 3180897 | 3183113 |
| *O. sativa* | Os03g0148800 | Chr3 | 2713317 | 2715918 |
| *O. alta* | g271 | Chr1CC | 2843992 | 2844650 |
| *O. alta* | g17188 | Chr3CC | 3194658 | 3195316 |
| *O. sativa* | Os03g0149000 | Chr3 | 2723037 | 2723835 |
| *O. alta* | g272 | Chr1CC | 2847460 | 2848975 |
| *O. alta* | g17189 | Chr3CC | 3198209 | 3199069 |
| *O. sativa* | Os03g0149100 | Chr3 | 2726696 | 2727559 |
| *O. alta* | g273 | Chr1CC | 2858602 | 2861688 |
| *O. alta* | g17190 | Chr3CC | 3209383 | 3212462 |
| *O. sativa* | Os03g0149200 | Chr3 | 2739653 | 2743118 |

| | | | | |
|---|---|---|---|---|
| *O. alta* | g274 | Chr1CC | 2869550 | 2871630 |
| *O. alta* | g17191 | Chr3CC | 3218296 | 3220359 |
| *O. sativa* | Os03g0149300 | Chr3 | 2748162 | 2753327 |
| *O. alta* | g277 | Chr1CC | 2890525 | 2891412 |
| *O. alta* | g17193 | Chr3CC | 3235984 | 3236883 |
| *O. sativa* | Os03g0149700 | Chr3 | 2771187 | 2772433 |
| *O. alta* | g279 | Chr1CC | 2897657 | 2898568 |
| *O. alta* | g17194 | Chr3CC | 3241349 | 3242266 |
| *O. sativa* | Os03g0149800 | Chr3 | 2776626 | 2777769 |
| *O. alta* | g282 | Chr1CC | 2929459 | 2931039 |
| *O. alta* | g17196 | Chr3CC | 3264196 | 3265779 |
| *O. sativa* | Os03g0150600 | Chr3 | 2807616 | 2809540 |
| *O. alta* | g286 | Chr1CC | 2970801 | 2972372 |
| *O. alta* | g17197 | Chr3CC | 3271102 | 3272673 |
| *O. sativa* | Os03g0150800 | Chr3 | 2815502 | 2817322 |
| *O. alta* | g287 | Chr1CC | 2977313 | 2982176 |
| *O. alta* | g17198 | Chr3CC | 3279929 | 3282541 |
| *O. sativa* | Os03g0151100 | Chr3 | 2823077 | 2828037 |
| *O. alta* | g288 | Chr1CC | 2995609 | 3003524 |
| *O. alta* | g17201 | Chr3CC | 3316470 | 3324260 |
| *O. alta* | g289 | Chr1CC | 3004257 | 3004571 |
| *O. alta* | g17202 | Chr3CC | 3325005 | 3325316 |
| *O. sativa* | Os03g0151500 | Chr3 | 2841790 | 2842391 |
| *O. alta* | g291 | Chr1CC | 3010652 | 3016786 |
| *O. alta* | g17204 | Chr3CC | 3329585 | 3335027 |
| *O. sativa* | Os03g0151700 | Chr3 | 2845714 | 2851680 |
| *O. alta* | g292 | Chr1CC | 3017878 | 3021809 |
| *O. alta* | g17205 | Chr3CC | 3336063 | 3340023 |
| *O. sativa* | Os03g0151800 | Chr3 | 2852295 | 2856681 |
| *O. alta* | g293 | Chr1CC | 3034865 | 3037461 |
| *O. alta* | g17206 | Chr3CC | 3341316 | 3343921 |
| *O. sativa* | Os03g0151900 | Chr3 | 2857179 | 2860218 |
| *O. alta* | g294 | Chr1CC | 3042630 | 3044038 |
| *O. alta* | g17207 | Chr3CC | 3354959 | 3356365 |
| *O. sativa* | Os03g0152000 | Chr3 | 2866379 | 2868145 |

| | | | | |
|---|---|---|---|---|
| *O. alta* | g296 | Chr1CC | 3055210 | 3057182 |
| *O. alta* | g17208 | Chr3CC | 3365216 | 3369973 |
| *O. sativa* | Os03g0152100 | Chr3 | 2870376 | 2875830 |
| *O. alta* | g297 | Chr1CC | 3062863 | 3064363 |
| *O. alta* | g17209 | Chr3CC | 3374093 | 3375352 |
| *O. sativa* | Os03g0152300 | Chr3 | 2878831 | 2880830 |
| *O. alta* | g298 | Chr1CC | 3066122 | 3076305 |
| *O. alta* | g17210 | Chr3CC | 3378291 | 3382725 |
| *O. sativa* | Os03g0152400 | Chr3 | 2883005 | 2887887 |
| *O. alta* | g300 | Chr1CC | 3089532 | 3092502 |
| *O. alta* | g17211 | Chr3CC | 3384238 | 3387088 |
| *O. sativa* | Os03g0152600 | Chr3 | 2897477 | 2900285 |
| *O. alta* | g302 | Chr1CC | 3103453 | 3107785 |
| *O. alta* | g17213 | Chr3CC | 3401582 | 3405942 |
| *O. sativa* | Os03g0152800 | Chr3 | 2907115 | 2912658 |
| *O. alta* | g303 | Chr1CC | 3109516 | 3113478 |
| *O. alta* | g17214 | Chr3CC | 3420271 | 3429422 |
| *O. sativa* | Os03g0152900 | Chr3 | 2913921 | 2922883 |
| *O. alta* | g304 | Chr1CC | 3119741 | 3127020 |
| *O. alta* | g17215 | Chr3CC | 3429958 | 3437129 |
| *O. alta* | g307 | Chr1CC | 3149017 | 3151059 |
| *O. alta* | g17217 | Chr3CC | 3453976 | 3454885 |
| *O. sativa* | Os03g0153900 | Chr3 | 2958312 | 2960398 |
| *O. alta* | g309 | Chr1CC | 3160790 | 3163007 |
| *O. alta* | g17219 | Chr3CC | 3456772 | 3459203 |
| *O. sativa* | Os03g0154100 | Chr3 | 2964716 | 2967286 |
| *O. alta* | g310 | Chr1CC | 3166362 | 3167316 |
| *O. alta* | g17220 | Chr3CC | 3462010 | 3462972 |
| *O. sativa* | Os03g0155300 | Chr3 | 2998855 | 2999999 |
| *O. alta* | g314 | Chr1CC | 3189579 | 3190420 |
| *O. alta* | g17222 | Chr3CC | 3475238 | 3476203 |
| *O. sativa* | Os03g0155700 | Chr3 | 3006419 | 3007309 |
| *O. alta* | g315 | Chr1CC | 3192603 | 3193774 |
| *O. alta* | g17221 | Chr3CC | 3472496 | 3473487 |
| *O. sativa* | Os03g0155500 | Chr3 | 3003413 | 3004537 |

| | | | | |
|---|---|---|---|---|
| *O. alta* | g312 | Chr1CC | 3177426 | 3178421 |
| *O. alta* | g17224 | Chr3CC | 3486879 | 3488106 |
| *O. sativa* | Os03g0156000 | Chr3 | 3026522 | 3027521 |
| *O. alta* | g317 | Chr1CC | 3214258 | 3214919 |
| *O. alta* | g17227 | Chr3CC | 3507521 | 3508179 |
| *O. sativa* | Os03g0156600 | Chr3 | 3046791 | 3047782 |
| *O. alta* | g318 | Chr1CC | 3215788 | 3218537 |
| *O. alta* | g17228 | Chr3CC | 3509204 | 3511897 |
| *O. sativa* | Os03g0156700 | Chr3 | 3048617 | 3051742 |
| *O. alta* | g319 | Chr1CC | 3223092 | 3224892 |
| *O. alta* | g17230 | Chr3CC | 3537253 | 3540917 |
| *O. sativa* | Os03g0157300 | Chr3 | 3068969 | 3074025 |
| *O. alta* | g320 | Chr1CC | 3229127 | 3233963 |
| *O. alta* | g17231 | Chr3CC | 3548552 | 3553422 |
| *O. sativa* | Os03g0157400 | Chr3 | 3076026 | 3085989 |
| *O. alta* | g321 | Chr1CC | 3243794 | 3244382 |
| *O. alta* | g17232 | Chr3CC | 3571762 | 3575624 |
| *O. sativa* | Os03g0157500 | Chr3 | 3091858 | 3095644 |
| *O. alta* | g322 | Chr1CC | 3248050 | 3248658 |
| *O. alta* | g17233 | Chr3CC | 3576613 | 3577194 |
| *O. sativa* | Os03g0157600 | Chr3 | 3096030 | 3096880 |
| *O. alta* | g327 | Chr1CC | 3282351 | 3287485 |
| *O. alta* | g17235 | Chr3CC | 3591967 | 3597082 |
| *O. sativa* | Os03g0157800 | Chr3 | 3103834 | 3109737 |
| *O. alta* | g328 | Chr1CC | 3289304 | 3290572 |
| *O. alta* | g17236 | Chr3CC | 3598783 | 3600063 |
| *O. sativa* | Os03g0157900 | Chr3 | 3110725 | 3112384 |
| *O. alta* | g329 | Chr1CC | 3293101 | 3296837 |
| *O. alta* | g17237 | Chr3CC | 3602215 | 3605971 |
| *O. sativa* | Os03g0158200 | Chr3 | 3115986 | 3120267 |
| *O. alta* | g330 | Chr1CC | 3297425 | 3298969 |
| *O. alta* | g17238 | Chr3CC | 3606642 | 3608414 |
| *O. sativa* | Os03g0158300 | Chr3 | 3120322 | 3123758 |
| *O. alta* | g331 | Chr1CC | 3304408 | 3307988 |
| *O. alta* | g17239 | Chr3CC | 3624717 | 3628277 |
| *O. sativa* | Os03g0158500 | Chr3 | 3126216 | 3130478 |

| | | | | |
|---|---|---|---|---|
| *O. alta* | g332 | Chr1CC | 3311753 | 3312094 |
| *O. alta* | g17240 | Chr3CC | 3635575 | 3636171 |
| *O. sativa* | Os03g0158600 | Chr3 | 3132124 | 3132741 |
| *O. alta* | g333 | Chr1CC | 3323891 | 3329953 |
| *O. alta* | g17241 | Chr3CC | 3639576 | 3646379 |
| *O. sativa* | Os03g0158700 | Chr3 | 3151514 | 3165238 |
| *O. alta* | g334 | Chr1CC | 3335323 | 3337991 |
| *O. alta* | g17242 | Chr3CC | 3650557 | 3653218 |
| *O. sativa* | Os03g0159100 | Chr3 | 3165801 | 3168806 |
| *O. alta* | g335 | Chr1CC | 3339049 | 3344873 |
| *O. alta* | g17243 | Chr3CC | 3655759 | 3662008 |
| *O. sativa* | Os03g0159200 | Chr3 | 3170103 | 3176292 |
| *O. alta* | g336 | Chr1CC | 3347670 | 3348335 |
| *O. alta* | g17244 | Chr3CC | 3664649 | 3665296 |
| *O. sativa* | Os03g0159400 | Chr3 | 3178572 | 3179590 |
| *O. alta* | g338 | Chr1CC | 3370537 | 3371508 |
| *O. alta* | g17245 | Chr3CC | 3678734 | 3679741 |
| *O. sativa* | Os03g0159600 | Chr3 | 3184496 | 3185936 |
| *O. alta* | g339 | Chr1CC | 3374015 | 3375609 |
| *O. alta* | g17246 | Chr3CC | 3680539 | 3680841 |
| *O. sativa* | Os03g0159700 | Chr3 | 3186782 | 3189325 |
| *O. alta* | g341 | Chr1CC | 3378780 | 3379262 |
| *O. alta* | g17247 | Chr3CC | 3693480 | 3693980 |
| *O. sativa* | Os03g0159900 | Chr3 | 3195766 | 3196656 |
| *O. alta* | g342 | Chr1CC | 3385324 | 3391050 |
| *O. alta* | g17248 | Chr3CC | 3698103 | 3705606 |
| *O. sativa* | Os03g0160100 | Chr3 | 3201345 | 3208276 |
| *O. alta* | g343 | Chr1CC | 3392882 | 3393451 |
| *O. alta* | g17249 | Chr3CC | 3705928 | 3706497 |
| *O. sativa* | Os03g0160200 | Chr3 | 3208862 | 3211539 |
| *O. alta* | g345 | Chr1CC | 3401210 | 3404375 |
| *O. alta* | g17251 | Chr3CC | 3712476 | 3715476 |
| *O. sativa* | Os03g0160400 | Chr3 | 3215854 | 3220186 |
| *O. alta* | g349 | Chr1CC | 3451770 | 3460807 |
| *O. alta* | g17256 | Chr3CC | 3790861 | 3802989 |

| | | | | |
|---|---|---|---|---|
| *O. sativa* | Os03g0161100 | Chr3 | 3260107 | 3270386 |
| *O. alta* | g350 | Chr1CC | 3461832 | 3466525 |
| *O. alta* | g17257 | Chr3CC | 3804001 | 3809151 |
| *O. sativa* | Os03g0161200 | Chr3 | 3271150 | 3276878 |
| *O. alta* | g352 | Chr1CC | 3477321 | 3478140 |
| *O. alta* | g17259 | Chr3CC | 3846842 | 3847568 |
| *O. sativa* | Os03g0161400 | Chr3 | 3310338 | 3312548 |
| *O. alta* | g354 | Chr1CC | 3481070 | 3483374 |
| *O. alta* | g17261 | Chr3CC | 3850483 | 3853060 |
| *O. sativa* | Os03g0161500 | Chr3 | 3314782 | 3317408 |
| *O. alta* | g356 | Chr1CC | 3506477 | 3508809 |
| *O. alta* | g17264 | Chr3CC | 3871644 | 3873488 |
| *O. sativa* | Os03g0161900 | Chr3 | 3342254 | 3344547 |
| *O. alta* | g357 | Chr1CC | 3525305 | 3527756 |
| *O. alta* | g17266 | Chr3CC | 3888405 | 3891061 |
| *O. sativa* | Os03g0162000 | Chr3 | 3355041 | 3360448 |
| *O. alta* | g358 | Chr1CC | 3543823 | 3545660 |
| *O. alta* | g17267 | Chr3CC | 3909582 | 3910148 |
| *O. sativa* | Os03g0162200 | Chr3 | 3371375 | 3372344 |
| *O. alta* | g359 | Chr1CC | 3559055 | 3560082 |
| *O. alta* | g17268 | Chr3CC | 3922956 | 3923970 |
| *O. sativa* | Os03g0162500 | Chr3 | 3380200 | 3381695 |
| *O. alta* | g367 | Chr1CC | 3657275 | 3657604 |
| *O. alta* | g17274 | Chr3CC | 3991456 | 3992369 |
| *O. sativa* | Os03g0163400 | Chr3 | 3433024 | 3434082 |
| *O. alta* | g361 | Chr1CC | 3606668 | 3609067 |
| *O. alta* | g17270 | Chr3CC | 3961513 | 3963978 |
| *O. sativa* | Os03g0162900 | Chr3 | 3407989 | 3413958 |
| *O. alta* | g362 | Chr1CC | 3616684 | 3623542 |
| *O. alta* | g17271 | Chr3CC | 3970834 | 3976688 |
| *O. sativa* | Os03g0163100 | Chr3 | 3417333 | 3424868 |
| *O. alta* | g363 | Chr1CC | 3624554 | 3628176 |
| *O. alta* | g17273 | Chr3CC | 3979487 | 3983488 |
| *O. sativa* | Os03g0163300 | Chr3 | 3427553 | 3432223 |
| *O. alta* | g368 | Chr1CC | 3660923 | 3663289 |

| | | | | |
|---|---|---|---|---|
| *O. alta* | g17275 | Chr3CC | 3998952 | 4001318 |
| *O. sativa* | Os03g0163500 | Chr3 | 3435024 | 3439334 |
| *O. alta* | g370 | Chr1CC | 3678970 | 3679206 |
| *O. alta* | g17277 | Chr3CC | 4011945 | 4012181 |
| *O. sativa* | Os03g0163900 | Chr3 | 3452742 | 3453008 |
| *O. alta* | g372 | Chr1CC | 3684676 | 3684918 |
| *O. alta* | g17279 | Chr3CC | 4023162 | 4023437 |
| *O. alta* | g374 | Chr1CC | 3714095 | 3716583 |
| *O. alta* | g17282 | Chr3CC | 4034891 | 4037377 |
| *O. sativa* | Os03g0164700 | Chr3 | 3479411 | 3482159 |
| *O. alta* | g375 | Chr1CC | 3717488 | 3719956 |
| *O. alta* | g17283 | Chr3CC | 4038758 | 4041114 |
| *O. sativa* | Os03g0164800 | Chr3 | 3483353 | 3485708 |
| *O. alta* | g376 | Chr1CC | 3724540 | 3738856 |
| *O. alta* | g17284 | Chr3CC | 4061629 | 4064428 |
| *O. alta* | g377 | Chr1CC | 3741890 | 3744792 |
| *O. alta* | g17285 | Chr3CC | 4067109 | 4069049 |
| *O. sativa* | Os03g0165100 | Chr3 | 3501294 | 3504478 |
| *O. alta* | g380 | Chr1CC | 3749841 | 3757775 |
| *O. alta* | g17286 | Chr3CC | 4069307 | 4076496 |
| *O. sativa* | Os03g0165266 | Chr3 | 3506365 | 3512883 |
| *O. alta* | g381 | Chr1CC | 3758545 | 3760569 |
| *O. alta* | g17287 | Chr3CC | 4077332 | 4079370 |
| *O. alta* | g382 | Chr1CC | 3778069 | 3783515 |
| *O. alta* | g17288 | Chr3CC | 4094178 | 4099645 |
| *O. sativa* | Os03g0165400 | Chr3 | 3525115 | 3531350 |
| *O. alta* | g384 | Chr1CC | 3797647 | 3808218 |
| *O. alta* | g17289 | Chr3CC | 4109006 | 4119039 |
| *O. sativa* | Os03g0165600 | Chr3 | 3539613 | 3544723 |
| *O. alta* | g385 | Chr1CC | 3811880 | 3813898 |
| *O. alta* | g17290 | Chr3CC | 4121241 | 4123308 |
| *O. sativa* | Os03g0165800 | Chr3 | 3551775 | 3554415 |
| *O. alta* | g386 | Chr1CC | 3815093 | 3816922 |
| *O. alta* | g17291 | Chr3CC | 4124605 | 4126826 |
| *O. sativa* | Os03g0165900 | Chr3 | 3555490 | 3558352 |

| | | | | |
|---|---|---|---|---|
| *O. alta* | g387 | Chr1CC | 3819634 | 3823045 |
| *O. alta* | g17292 | Chr3CC | 4128748 | 4132127 |
| *O. sativa* | Os03g0166000 | Chr3 | 3561596 | 3565365 |
| *O. alta* | g388 | Chr1CC | 3833129 | 3835124 |
| *O. alta* | g17293 | Chr3CC | 4137331 | 4139191 |
| *O. sativa* | Os03g0166200 | Chr3 | 3570194 | 3571828 |
| *O. alta* | g391 | Chr1CC | 3860561 | 3862072 |
| *O. alta* | g17296 | Chr3CC | 4155479 | 4156946 |
| *O. sativa* | Os03g0167000 | Chr3 | 3626392 | 3628578 |
| *O. alta* | g393 | Chr1CC | 3866289 | 3870712 |
| *O. alta* | g17299 | Chr3CC | 4167543 | 4171982 |
| *O. sativa* | Os03g0167200 | Chr3 | 3631183 | 3635525 |
| *O. alta* | g392 | Chr1CC | 3864667 | 3866093 |
| *O. alta* | g17298 | Chr3CC | 4165121 | 4167323 |
| *O. sativa* | Os10g0411700 | Chr10 | 14323590 | 14324864 |
| *O. alta* | g396 | Chr1CC | 3895180 | 3897528 |
| *O. alta* | g17303 | Chr3CC | 4211998 | 4214383 |
| *O. sativa* | Os03g0167600 | Chr3 | 3653709 | 3657069 |
| *O. alta* | g397 | Chr1CC | 3897599 | 3897853 |
| *O. alta* | g17304 | Chr3CC | 4214447 | 4214716 |
| *O. alta* | g399 | Chr1CC | 3904905 | 3908891 |
| *O. alta* | g17306 | Chr3CC | 4222766 | 4226744 |
| *O. sativa* | Os03g0167800 | Chr3 | 3664640 | 3669746 |
| *O. alta* | g400 | Chr1CC | 3913694 | 3916500 |
| *O. alta* | g17307 | Chr3CC | 4233453 | 4236326 |
| *O. sativa* | Os03g0168000 | Chr3 | 3674285 | 3678314 |
| *O. alta* | g401 | Chr1CC | 3917042 | 3918257 |
| *O. alta* | g17308 | Chr3CC | 4236875 | 4238183 |
| *O. sativa* | Os03g0168100 | Chr3 | 3678292 | 3679969 |
| *O. alta* | g402 | Chr1CC | 3919458 | 3921243 |
| *O. alta* | g17309 | Chr3CC | 4246241 | 4248039 |
| *O. sativa* | Os03g0168200 | Chr3 | 3681100 | 3683235 |
| *O. alta* | g403 | Chr1CC | 3921677 | 3923137 |
| *O. alta* | g17310 | Chr3CC | 4248467 | 4250084 |
| *O. sativa* | Os03g0168300 | Chr3 | 3683172 | 3685295 |

| | | | | |
|---|---|---|---|---|
| *O. alta* | g404 | Chr1CC | 3926821 | 3931188 |
| *O. alta* | g17312 | Chr3CC | 4257247 | 4261687 |
| *O. alta* | g405 | Chr1CC | 3931456 | 3932108 |
| *O. alta* | g17313 | Chr3CC | 4261957 | 4263649 |
| *O. sativa* | Os03g0168500 | Chr3 | 3694742 | 3696580 |
| *O. alta* | g406 | Chr1CC | 3936808 | 3938454 |
| *O. alta* | g17314 | Chr3CC | 4267892 | 4269526 |
| *O. sativa* | Os03g0168550 | Chr3 | 3700039 | 3701677 |
| *O. alta* | g407 | Chr1CC | 3938734 | 3940653 |
| *O. alta* | g17315 | Chr3CC | 4269964 | 4271888 |
| *O. sativa* | Os03g0168600 | Chr3 | 3702125 | 3704085 |
| *O. alta* | g408 | Chr1CC | 3941665 | 3943296 |
| *O. alta* | g17316 | Chr3CC | 4272858 | 4274468 |
| *O. sativa* | Os03g0168700 | Chr3 | 3705167 | 3706777 |
| *O. alta* | g409 | Chr1CC | 3944355 | 3946380 |
| *O. alta* | g17317 | Chr3CC | 4275727 | 4277507 |
| *O. sativa* | Os03g0168900 | Chr3 | 3711147 | 3713253 |
| *O. alta* | g410 | Chr1CC | 3947326 | 3950461 |
| *O. alta* | g17318 | Chr3CC | 4278446 | 4281610 |
| *O. sativa* | Os03g0169100 | Chr3 | 3713990 | 3717346 |
| *O. alta* | g411 | Chr1CC | 3952067 | 3955301 |
| *O. alta* | g17319 | Chr3CC | 4282278 | 4285582 |
| *O. sativa* | Os03g0169000 | Chr3 | 3713492 | 3721105 |
| *O. alta* | g412 | Chr1CC | 3956931 | 3958557 |
| *O. alta* | g17320 | Chr3CC | 4288924 | 4290462 |
| *O. sativa* | Os03g0169300 | Chr3 | 3724584 | 3726496 |
| *O. alta* | g413 | Chr1CC | 3960254 | 3961029 |
| *O. alta* | g17321 | Chr3CC | 4292267 | 4293056 |
| *O. sativa* | Os03g0169400 | Chr3 | 3727764 | 3728922 |
| *O. alta* | g414 | Chr1CC | 3961453 | 3966600 |
| *O. alta* | g17322 | Chr3CC | 4293477 | 4297830 |
| *O. sativa* | Os03g0169500 | Chr3 | 3729174 | 3733685 |
| *O. alta* | g415 | Chr1CC | 3974960 | 3977127 |
| *O. alta* | g17323 | Chr3CC | 4307808 | 4309996 |
| *O. sativa* | Os03g0169600 | Chr3 | 3738868 | 3741611 |

| | | | | |
|---|---|---|---|---|
| *O. alta* | g416 | Chr1CC | 3981938 | 3983211 |
| *O. alta* | g17324 | Chr3CC | 4313172 | 4314704 |
| *O. sativa* | Os03g0169800 | Chr3 | 3744487 | 3746086 |
| *O. alta* | g419 | Chr1CC | 4005800 | 4006309 |
| *O. alta* | g17338 | Chr3CC | 4596048 | 4596407 |
| *O. sativa* | Os03g0170100 | Chr3 | 3757543 | 3758135 |
| *O. alta* | g420 | Chr1CC | 4008359 | 4011495 |
| *O. alta* | g17340 | Chr3CC | 4603622 | 4607466 |
| *O. sativa* | Os03g0170300 | Chr3 | 3763827 | 3767153 |
| *O. alta* | g421 | Chr1CC | 4012253 | 4014166 |
| *O. alta* | g17341 | Chr3CC | 4608231 | 4610996 |
| *O. alta* | g423 | Chr1CC | 4025370 | 4025930 |
| *O. alta* | g17342 | Chr3CC | 4617081 | 4617647 |
| *O. sativa* | Os03g0170500 | Chr3 | 3776809 | 3777621 |
| *O. alta* | g424 | Chr1CC | 4031602 | 4033550 |
| *O. alta* | g17343 | Chr3CC | 4630486 | 4632511 |
| *O. sativa* | Os03g0170600 | Chr3 | 3784341 | 3785140 |
| *O. alta* | g425 | Chr1CC | 4038076 | 4039104 |
| *O. alta* | g17344 | Chr3CC | 4639418 | 4640446 |
| *O. sativa* | Os03g0170800 | Chr3 | 3794582 | 3795566 |
| *O. alta* | g426 | Chr1CC | 4041313 | 4046737 |
| *O. alta* | g17345 | Chr3CC | 4641913 | 4647184 |
| *O. sativa* | Os03g0170900 | Chr3 | 3797698 | 3804132 |
| *O. alta* | g429 | Chr1CC | 4077481 | 4078773 |
| *O. alta* | g17347 | Chr3CC | 4683481 | 4684776 |
| *O. sativa* | Os03g0171600 | Chr3 | 3834150 | 3836735 |
| *O. alta* | g431 | Chr1CC | 4096008 | 4099780 |
| *O. alta* | g17349 | Chr3CC | 4708091 | 4710928 |
| *O. sativa* | Os03g0171900 | Chr3 | 3854304 | 3858338 |
| *O. alta* | g432 | Chr1CC | 4105335 | 4109209 |
| *O. alta* | g17350 | Chr3CC | 4714256 | 4718133 |
| *O. sativa* | Os03g0172000 | Chr3 | 3859447 | 3864548 |
| *O. alta* | g433 | Chr1CC | 4111802 | 4112044 |
| *O. alta* | g17351 | Chr3CC | 4722699 | 4722941 |
| *O. sativa* | Os03g0172100 | Chr3 | 3866613 | 3867121 |

| | | | | |
|---|---|---|---|---|
| *O. alta* | g434 | Chr1CC | 4112623 | 4115936 |
| *O. alta* | g17352 | Chr3CC | 4723698 | 4727228 |
| *O. sativa* | Os03g0172200 | Chr3 | 3867253 | 3871354 |
| *O. alta* | g436 | Chr1CC | 4118477 | 4118869 |
| *O. alta* | g17353 | Chr3CC | 4729335 | 4729733 |
| *O. sativa* | Os03g0172400 | Chr3 | 3874153 | 3874548 |
| *O. alta* | g437 | Chr1CC | 4129355 | 4129975 |
| *O. alta* | g17354 | Chr3CC | 4741358 | 4742026 |
| *O. alta* | g439 | Chr1CC | 4138204 | 4138581 |
| *O. alta* | g17355 | Chr3CC | 4750027 | 4750404 |
| *O. sativa* | Os03g0172700 | Chr3 | 3889565 | 3890307 |
| *O. alta* | g440 | Chr1CC | 4141924 | 4142289 |
| *O. alta* | g17356 | Chr3CC | 4762200 | 4762565 |
| *O. sativa* | Os03g0172850 | Chr3 | 3893027 | 3893580 |
| *O. alta* | g442 | Chr1CC | 4146838 | 4147203 |
| *O. alta* | g17357 | Chr3CC | 4769215 | 4769571 |
| *O. alta* | g445 | Chr1CC | 4161730 | 4165758 |
| *O. alta* | g17362 | Chr3CC | 4840917 | 4845747 |
| *O. sativa* | Os03g0173900 | Chr3 | 3961041 | 3966305 |
| *O. alta* | g444 | Chr1CC | 4156360 | 4160444 |
| *O. alta* | g17363 | Chr3CC | 4847427 | 4851516 |
| *O. sativa* | Os03g0174100 | Chr3 | 3970216 | 3974508 |
| *O. alta* | g443 | Chr1CC | 4153522 | 4155586 |
| *O. alta* | g17364 | Chr3CC | 4852463 | 4854555 |
| *O. sativa* | Os03g0174200 | Chr3 | 3974957 | 3977413 |
| *O. alta* | g447 | Chr1CC | 4200573 | 4204636 |
| *O. alta* | g17365 | Chr3CC | 4858416 | 4862470 |
| *O. sativa* | Os03g0174300 | Chr3 | 3978668 | 3983475 |
| *O. alta* | g448 | Chr1CC | 4206267 | 4207202 |
| *O. alta* | g17366 | Chr3CC | 4864077 | 4865009 |
| *O. alta* | g449 | Chr1CC | 4208338 | 4211205 |
| *O. alta* | g17367 | Chr3CC | 4866318 | 4869172 |
| *O. sativa* | Os03g0174500 | Chr3 | 3986852 | 3989994 |

| | | | | |
|---|---|---|---|---|
| *O. alta* | g451 | Chr1CC | 4219246 | 4219446 |
| *O. alta* | g17369 | Chr3CC | 4876023 | 4876223 |
| *O. sativa* | Os03g0174800 | Chr3 | 3999092 | 4000029 |
| *O. alta* | g452 | Chr1CC | 4243805 | 4246729 |
| *O. alta* | g17370 | Chr3CC | 4880636 | 4881930 |
| *O. alta* | g453 | Chr1CC | 4260122 | 4262428 |
| *O. alta* | g17372 | Chr3CC | 4893311 | 4894953 |
| *O. alta* | g454 | Chr1CC | 4268828 | 4272901 |
| *O. alta* | g17374 | Chr3CC | 4904762 | 4908683 |
| *O. sativa* | Os03g0175600 | Chr3 | 4024383 | 4028509 |
| *O. alta* | g455 | Chr1CC | 4291119 | 4292054 |
| *O. alta* | g17375 | Chr3CC | 4918528 | 4919475 |
| *O. sativa* | Os03g0175800 | Chr3 | 4038341 | 4039509 |
| *O. alta* | g457 | Chr1CC | 4315584 | 4319264 |
| *O. alta* | g17377 | Chr3CC | 4941825 | 4945879 |
| *O. sativa* | Os03g0176300 | Chr3 | 4057561 | 4059890 |
| *O. alta* | g458 | Chr1CC | 4325605 | 4327661 |
| *O. alta* | g17378 | Chr3CC | 4951005 | 4952884 |
| *O. sativa* | Os03g0176700 | Chr3 | 4069860 | 4072360 |
| *O. alta* | g459 | Chr1CC | 4355587 | 4355904 |
| *O. alta* | g17380 | Chr3CC | 4970489 | 4970806 |
| *O. sativa* | Os03g0176900 | Chr3 | 4079447 | 4080084 |
| *O. alta* | g462 | Chr1CC | 4386716 | 4388504 |
| *O. alta* | g17383 | Chr3CC | 4989139 | 4992646 |
| *O. sativa* | Os03g0178000 | Chr3 | 4116932 | 4119755 |
| *O. alta* | g460 | Chr1CC | 4365795 | 4369265 |
| *O. alta* | g17382 | Chr3CC | 4978354 | 4986989 |
| *O. sativa* | Os03g0177300 | Chr3 | 4090005 | 4093914 |
| *O. alta* | g465 | Chr1CC | 4415320 | 4420168 |
| *O. alta* | g17387 | Chr3CC | 5021851 | 5028969 |
| *O. sativa* | Os03g0178100 | Chr3 | 4120367 | 4126939 |
| *O. alta* | g467 | Chr1CC | 4427048 | 4428363 |
| *O. alta* | g17388 | Chr3CC | 5030638 | 5031798 |
| *O. sativa* | Os03g0178200 | Chr3 | 4129653 | 4131131 |
| *O. alta* | g468 | Chr1CC | 4428891 | 4430115 |

| | | | | |
|---|---|---|---|---|
| *O. alta* | g17389 | Chr3CC | 5032297 | 5033498 |
| *O. sativa* | Os03g0178300 | Chr3 | 4131128 | 4132544 |
| *O. alta* | g469 | Chr1CC | 4430767 | 4432646 |
| *O. alta* | g17390 | Chr3CC | 5034113 | 5036309 |
| *O. sativa* | Os03g0178400 | Chr3 | 4133458 | 4136490 |
| *O. alta* | g470 | Chr1CC | 4433419 | 4438810 |
| *O. alta* | g17391 | Chr3CC | 5037397 | 5042067 |
| *O. sativa* | Os03g0178500 | Chr3 | 4136960 | 4142715 |
| *O. alta* | g472 | Chr1CC | 4447010 | 4453391 |
| *O. alta* | g17392 | Chr3CC | 5043605 | 5050009 |
| *O. alta* | g473 | Chr1CC | 4453804 | 4454499 |
| *O. alta* | g17393 | Chr3CC | 5050432 | 5051127 |
| *O. sativa* | Os03g0179100 | Chr3 | 4157591 | 4158512 |
| *O. alta* | g474 | Chr1CC | 4469204 | 4473980 |
| *O. alta* | g17396 | Chr3CC | 5072884 | 5078151 |
| *O. sativa* | Os03g0179400 | Chr3 | 4173178 | 4179068 |
| *O. alta* | g477 | Chr1CC | 4487458 | 4488337 |
| *O. alta* | g17397 | Chr3CC | 5084954 | 5085832 |
| *O. sativa* | Os03g0179700 | Chr3 | 4181381 | 4182557 |
| *O. alta* | g478 | Chr1CC | 4491647 | 4497447 |
| *O. alta* | g17399 | Chr3CC | 5093497 | 5099125 |
| *O. sativa* | Os03g0179900 | Chr3 | 4187158 | 4193200 |
| *O. alta* | g479 | Chr1CC | 4499095 | 4501023 |
| *O. alta* | g17400 | Chr3CC | 5100087 | 5102184 |
| *O. sativa* | Os03g0180000 | Chr3 | 4194372 | 4196475 |
| *O. alta* | g480 | Chr1CC | 4506505 | 4506921 |
| *O. alta* | g17401 | Chr3CC | 5105375 | 5105788 |
| *O. sativa* | Os03g0180100 | Chr3 | 4199851 | 4200735 |
| *O. alta* | g481 | Chr1CC | 4511270 | 4515574 |
| *O. alta* | g17402 | Chr3CC | 5110724 | 5115084 |
| *O. sativa* | Os03g0180300 | Chr3 | 4203524 | 4210236 |
| *O. alta* | g482 | Chr1CC | 4516291 | 4518478 |
| *O. alta* | g17403 | Chr3CC | 5115800 | 5118089 |
| *O. sativa* | Os03g0180400 | Chr3 | 4210533 | 4212985 |
| *O. alta* | g483 | Chr1CC | 4521211 | 4522173 |

| | | | | |
|---|---|---|---|---|
| *O. alta* | g17404 | Chr3CC | 5126841 | 5128084 |
| *O. sativa* | Os03g0180600 | Chr3 | 4222085 | 4223728 |
| *O. alta* | g485 | Chr1CC | 4543202 | 4543723 |
| *O. alta* | g17406 | Chr3CC | 5137823 | 5138335 |
| *O. sativa* | Os03g0180800 | Chr3 | 4232004 | 4233038 |
| *O. alta* | g486 | Chr1CC | 4548851 | 4549759 |
| *O. alta* | g17407 | Chr3CC | 5154507 | 5155395 |
| *O. sativa* | Os03g0180900 | Chr3 | 4236724 | 4238058 |
| *O. alta* | g487 | Chr1CC | 4559040 | 4559600 |
| *O. alta* | g17408 | Chr3CC | 5162944 | 5163504 |
| *O. sativa* | Os03g0181100 | Chr3 | 4248884 | 4249820 |
| *O. alta* | g489 | Chr1CC | 4590944 | 4594578 |
| *O. alta* | g17410 | Chr3CC | 5196001 | 5199551 |
| *O. sativa* | Os03g0181500 | Chr3 | 4266978 | 4271781 |
| *O. alta* | g490 | Chr1CC | 4606308 | 4611829 |
| *O. alta* | g17412 | Chr3CC | 5208012 | 5213948 |
| *O. sativa* | Os03g0181750 | Chr3 | 4281418 | 4284885 |
| *O. alta* | g491 | Chr1CC | 4616581 | 4619047 |
| *O. alta* | g17413 | Chr3CC | 5217922 | 5220952 |
| *O. sativa* | Os03g0181800 | Chr3 | 4287354 | 4291799 |
| *O. alta* | g492 | Chr1CC | 4630656 | 4633677 |
| *O. alta* | g17414 | Chr3CC | 5230509 | 5233495 |
| *O. sativa* | Os03g0182000 | Chr3 | 4311292 | 4314363 |
| *O. alta* | g493 | Chr1CC | 4644659 | 4651437 |
| *O. alta* | g17415 | Chr3CC | 5248171 | 5255254 |
| *O. sativa* | Os03g0182400 | Chr3 | 4327773 | 4336225 |
| *O. alta* | g494 | Chr1CC | 4653218 | 4655002 |
| *O. alta* | g17416 | Chr3CC | 5257214 | 5259035 |
| *O. sativa* | Os03g0182600 | Chr3 | 4336742 | 4338901 |
| *O. alta* | g495 | Chr1CC | 4655767 | 4658651 |
| *O. alta* | g17417 | Chr3CC | 5259932 | 5262842 |
| *O. sativa* | Os03g0182700 | Chr3 | 4339485 | 4342714 |
| *O. alta* | g496 | Chr1CC | 4661082 | 4662306 |
| *O. alta* | g17418 | Chr3CC | 5263968 | 5265202 |
| *O. sativa* | Os03g0182800 | Chr3 | 4343807 | 4345689 |

| | | | | |
|---|---|---|---|---|
| *O. alta* | g497 | Chr1CC | 4666176 | 4667298 |
| *O. alta* | g17419 | Chr3CC | 5268939 | 5270067 |
| *O. sativa* | Os03g0183000 | Chr3 | 4348736 | 4350531 |
| *O. alta* | g499 | Chr1CC | 4695781 | 4696740 |
| *O. alta* | g17420 | Chr3CC | 5279466 | 5280446 |
| *O. alta* | g501 | Chr1CC | 4705922 | 4706917 |
| *O. alta* | g17421 | Chr3CC | 5285235 | 5286218 |
| *O. alta* | g502 | Chr1CC | 4712073 | 4712699 |
| *O. alta* | g17422 | Chr3CC | 5292605 | 5293234 |
| *O. sativa* | Os03g0183500 | Chr3 | 4379668 | 4380736 |
| *O. alta* | g503 | Chr1CC | 4714379 | 4718560 |
| *O. alta* | g17423 | Chr3CC | 5298848 | 5304358 |
| *O. sativa* | Os03g0183600 | Chr3 | 4384585 | 4389310 |
| *O. alta* | g505 | Chr1CC | 4724879 | 4730076 |
| *O. alta* | g17424 | Chr3CC | 5308346 | 5313544 |
| *O. sativa* | Os03g0183800 | Chr3 | 4397245 | 4402938 |
| *O. alta* | g506 | Chr1CC | 4730587 | 4735583 |
| *O. alta* | g17425 | Chr3CC | 5314059 | 5319083 |
| *O. sativa* | Os03g0183900 | Chr3 | 4403063 | 4405969 |
| *O. alta* | g507 | Chr1CC | 4739972 | 4744487 |
| *O. alta* | g17426 | Chr3CC | 5321388 | 5325802 |
| *O. alta* | g508 | Chr1CC | 4747098 | 4748819 |
| *O. alta* | g17427 | Chr3CC | 5328122 | 5329743 |
| *O. sativa* | Os03g0184100 | Chr3 | 4416654 | 4419091 |
| *O. alta* | g509 | Chr1CC | 4756597 | 4761381 |
| *O. alta* | g17428 | Chr3CC | 5340763 | 5345790 |
| *O. sativa* | Os03g0184300 | Chr3 | 4428838 | 4434161 |
| *O. alta* | g510 | Chr1CC | 4762728 | 4764956 |
| *O. alta* | g17429 | Chr3CC | 5346655 | 5349006 |
| *O. sativa* | Os03g0184400 | Chr3 | 4434560 | 4437813 |
| *O. alta* | g511 | Chr1CC | 4765569 | 4767172 |
| *O. alta* | g17430 | Chr3CC | 5349591 | 5351178 |
| *O. sativa* | Os03g0184500 | Chr3 | 4437782 | 4439874 |

| | | | | |
|---|---|---|---|---|
| *O. alta* | g512 | Chr1CC | 4773953 | 4779842 |
| *O. alta* | g17431 | Chr3CC | 5352559 | 5354239 |
| *O. sativa* | Os03g0184550 | Chr3 | 4441960 | 4443828 |
| *O. alta* | g513 | Chr1CC | 4786285 | 4788369 |
| *O. alta* | g17432 | Chr3CC | 5355139 | 5357190 |
| *O. alta* | g514 | Chr1CC | 4788988 | 4791022 |
| *O. alta* | g17433 | Chr3CC | 5357800 | 5359846 |
| *O. sativa* | Os03g0184700 | Chr3 | 4446728 | 4449163 |
| *O. alta* | g515 | Chr1CC | 4803756 | 4807264 |
| *O. alta* | g17435 | Chr3CC | 5412115 | 5413530 |
| *O. sativa* | Os03g0185200 | Chr3 | 4470567 | 4476128 |
| *O. alta* | g516 | Chr1CC | 4808631 | 4814673 |
| *O. alta* | g17436 | Chr3CC | 5414876 | 5420902 |
| *O. sativa* | Os03g0185500 | Chr3 | 4486362 | 4489500 |
| *O. alta* | g517 | Chr1CC | 4815470 | 4815847 |
| *O. alta* | g17437 | Chr3CC | 5421697 | 5422074 |
| *O. sativa* | Os03g0185600 | Chr3 | 4489695 | 4490510 |
| *O. alta* | g518 | Chr1CC | 4820806 | 4822158 |
| *O. alta* | g17438 | Chr3CC | 5424529 | 5425851 |
| *O. sativa* | Os03g0185700 | Chr3 | 4492084 | 4494048 |
| *O. alta* | g519 | Chr1CC | 4822658 | 4826569 |
| *O. alta* | g17439 | Chr3CC | 5426618 | 5430830 |
| *O. sativa* | Os03g0186100 | Chr3 | 4496734 | 4500824 |
| *O. alta* | g520 | Chr1CC | 4839486 | 4844532 |
| *O. alta* | g17440 | Chr3CC | 5443673 | 5448646 |
| *O. sativa* | Os03g0186500 | Chr3 | 4513398 | 4518521 |
| *O. alta* | g521 | Chr1CC | 4845856 | 4846687 |
| *O. alta* | g17441 | Chr3CC | 5449928 | 5456249 |
| *O. sativa* | Os03g0186600 | Chr3 | 4519405 | 4525692 |
| *O. alta* | g522 | Chr1CC | 4869519 | 4872557 |
| *O. alta* | g17442 | Chr3CC | 5462567 | 5464736 |
| *O. sativa* | Os03g0186800 | Chr3 | 4532886 | 4536584 |
| *O. alta* | g523 | Chr1CC | 4877108 | 4878418 |
| *O. alta* | g17443 | Chr3CC | 5478140 | 5479450 |
| *O. sativa* | Os03g0186900 | Chr3 | 4541351 | 4543092 |

| | | | | |
|---|---|---|---|---|
| *O. alta* | g524 | Chr1CC | 4879004 | 4880218 |
| *O. alta* | g17444 | Chr3CC | 5479981 | 5481319 |
| *O. sativa* | Os03g0186950 | Chr3 | 4543174 | 4545072 |
| *O. alta* | g525 | Chr1CC | 4881251 | 4884109 |
| *O. alta* | g17445 | Chr3CC | 5482169 | 5484629 |
| *O. sativa* | Os03g0187000 | Chr3 | 4545306 | 4547966 |
| *O. alta* | g526 | Chr1CC | 4888164 | 4906703 |
| *O. alta* | g17446 | Chr3CC | 5488864 | 5496842 |
| *O. sativa* | Os03g0187100 | Chr3 | 4555660 | 4559410 |
| *O. alta* | g527 | Chr1CC | 4911241 | 4922530 |
| *O. alta* | g17448 | Chr3CC | 5510883 | 5512963 |
| *O. sativa* | Os03g0187400 | Chr3 | 4573148 | 4575550 |
| *O. alta* | g528 | Chr1CC | 4931074 | 4933122 |
| *O. alta* | g17449 | Chr3CC | 5527034 | 5529088 |
| *O. sativa* | Os03g0187500 | Chr3 | 4584427 | 4586848 |
| *O. alta* | g529 | Chr1CC | 4941411 | 4943612 |
| *O. alta* | g17450 | Chr3CC | 5534806 | 5536763 |
| *O. sativa* | Os03g0187600 | Chr3 | 4592489 | 4594797 |
| *O. alta* | g530 | Chr1CC | 4944857 | 4946690 |
| *O. alta* | g17451 | Chr3CC | 5537884 | 5539703 |
| *O. sativa* | Os03g0187700 | Chr3 | 4596108 | 4598257 |
| *O. alta* | g531 | Chr1CC | 4948020 | 4949198 |
| *O. alta* | g17452 | Chr3CC | 5540871 | 5542049 |
| *O. sativa* | Os03g0187800 | Chr3 | 4598924 | 4600430 |
| *O. alta* | g532 | Chr1CC | 4991106 | 4997987 |
| *O. alta* | g17453 | Chr3CC | 5551380 | 5555357 |
| *O. sativa* | Os03g0188100 | Chr3 | 4606945 | 4616239 |
| *O. alta* | g534 | Chr1CC | 5004724 | 5005761 |
| *O. alta* | g17454 | Chr3CC | 5556304 | 5557374 |
| *O. sativa* | Os03g0188200 | Chr3 | 4616778 | 4618026 |
| *O. alta* | g536 | Chr1CC | 5022053 | 5023306 |
| *O. alta* | g17455 | Chr3CC | 5572276 | 5573506 |
| *O. sativa* | Os03g0188400 | Chr3 | 4628935 | 4630514 |
| *O. alta* | g537 | Chr1CC | 5028799 | 5029164 |
| *O. alta* | g17456 | Chr3CC | 5577340 | 5577723 |
| *O. sativa* | Os03g0188500 | Chr3 | 4634963 | 4635591 |

| | | | | |
|---|---|---|---|---|
| *O. alta* | g539 | Chr1CC | 5061534 | 5062461 |
| *O. alta* | g17457 | Chr3CC | 5592239 | 5593232 |
| *O. sativa* | Os03g0188900 | Chr3 | 4652930 | 4654781 |
| *O. alta* | g540 | Chr1CC | 5068008 | 5068538 |
| *O. alta* | g17459 | Chr3CC | 5603516 | 5604052 |
| *O. sativa* | Os03g0189100 | Chr3 | 4660862 | 4661923 |
| *O. alta* | g541 | Chr1CC | 5071806 | 5073239 |
| *O. alta* | g17460 | Chr3CC | 5608763 | 5610196 |
| *O. sativa* | Os03g0189300 | Chr3 | 4665311 | 4666944 |
| *O. alta* | g542 | Chr1CC | 5073788 | 5077907 |
| *O. alta* | g17461 | Chr3CC | 5610757 | 5612189 |
| *O. sativa* | Os03g0189400 | Chr3 | 4667276 | 4679965 |
| *O. alta* | g544 | Chr1CC | 5088531 | 5090978 |
| *O. alta* | g17464 | Chr3CC | 5633799 | 5635860 |
| *O. sativa* | Os03g0189600 | Chr3 | 4682354 | 4685319 |
| *O. alta* | g545 | Chr1CC | 5099831 | 5100406 |
| *O. alta* | g17465 | Chr3CC | 5651904 | 5652494 |
| *O. sativa* | Os03g0190000 | Chr3 | 4694814 | 4695941 |
| *O. alta* | g546 | Chr1CC | 5102668 | 5104900 |
| *O. alta* | g17466 | Chr3CC | 5654764 | 5656981 |
| *O. sativa* | Os03g0190100 | Chr3 | 4697347 | 4700133 |
| *O. alta* | g547 | Chr1CC | 5108163 | 5111146 |
| *O. alta* | g17468 | Chr3CC | 5661567 | 5665216 |
| *O. sativa* | Os03g0190300 | Chr3 | 4702183 | 4705151 |
| *O. alta* | g548 | Chr1CC | 5118680 | 5122292 |
| *O. alta* | g17469 | Chr3CC | 5670339 | 5674798 |
| *O. sativa* | Os03g0190800 | Chr3 | 4715169 | 4719205 |
| *O. alta* | g549 | Chr1CC | 5124048 | 5127135 |
| *O. alta* | g17470 | Chr3CC | 5677569 | 5680642 |
| *O. sativa* | Os03g0190900 | Chr3 | 4720699 | 4724099 |
| *O. alta* | g551 | Chr1CC | 5140623 | 5147810 |
| *O. alta* | g17474 | Chr3CC | 5698679 | 5705796 |
| *O. sativa* | Os03g0191000 | Chr3 | 4726165 | 4734003 |
| *O. alta* | g552 | Chr1CC | 5151603 | 5154353 |
| *O. alta* | g17475 | Chr3CC | 5709501 | 5712285 |

| | | | | |
|---|---|---|---|---|
| *O. sativa* | Os03g0191100 | Chr3 | 4737753 | 4741311 |
| *O. alta* | g553 | Chr1CC | 5160121 | 5160958 |
| *O. alta* | g17476 | Chr3CC | 5716627 | 5717461 |
| *O. sativa* | Os03g0191200 | Chr3 | 4748704 | 4750253 |
| *O. alta* | g556 | Chr1CC | 5191910 | 5193910 |
| *O. alta* | g17477 | Chr3CC | 5733171 | 5734989 |
| *O. sativa* | Os03g0191400 | Chr3 | 4763790 | 4766335 |
| *O. alta* | g557 | Chr1CC | 5199009 | 5200424 |
| *O. alta* | g17478 | Chr3CC | 5736437 | 5740211 |
| *O. sativa* | Os03g0191700 | Chr3 | 4768568 | 4773074 |
| *O. alta* | g558 | Chr1CC | 5202501 | 5203331 |
| *O. alta* | g17479 | Chr3CC | 5742200 | 5743036 |
| *O. sativa* | Os03g0191800 | Chr3 | 4775506 | 4776380 |
| *O. alta* | g559 | Chr1CC | 5209200 | 5210084 |
| *O. alta* | g17480 | Chr3CC | 5746896 | 5747756 |
| *O. alta* | g560 | Chr1CC | 5214717 | 5215747 |
| *O. alta* | g17481 | Chr3CC | 5752586 | 5753264 |
| *O. sativa* | Os03g0192000 | Chr3 | 4786226 | 4789319 |
| *O. alta* | g561 | Chr1CC | 5218219 | 5220116 |
| *O. alta* | g17482 | Chr3CC | 5757332 | 5759311 |
| *O. sativa* | Os03g0192100 | Chr3 | 4789625 | 4793058 |
| *O. alta* | g562 | Chr1CC | 5266142 | 5268470 |
| *O. alta* | g17484 | Chr3CC | 5773024 | 5775146 |
| *O. sativa* | Os03g0192400 | Chr3 | 4808253 | 4810821 |
| *O. alta* | g563 | Chr1CC | 5284855 | 5289666 |
| *O. alta* | g17485 | Chr3CC | 5779920 | 5784557 |
| *O. sativa* | Os03g0192500 | Chr3 | 4815464 | 4820358 |
| *O. alta* | g565 | Chr1CC | 5291282 | 5293288 |
| *O. alta* | g17486 | Chr3CC | 5785510 | 5786821 |
| *O. sativa* | Os03g0192600 | Chr3 | 4820699 | 4822659 |
| *O. alta* | g566 | Chr1CC | 5295947 | 5301210 |
| *O. alta* | g17487 | Chr3CC | 5803757 | 5807889 |
| *O. sativa* | Os03g0192700 | Chr3 | 4825697 | 4829533 |
| *O. alta* | g568 | Chr1CC | 5317844 | 5325957 |

| | | | | |
|---|---|---|---|---|
| *O. alta* | g17489 | Chr3CC | 5816032 | 5817611 |
| *O. sativa* | Os03g0192900 | Chr3 | 4842276 | 4844355 |
| *O. alta* | g570 | Chr1CC | 5345257 | 5346864 |
| *O. alta* | g17491 | Chr3CC | 5830562 | 5832136 |
| *O. sativa* | Os03g0193000 | Chr3 | 4847241 | 4853828 |
| *O. alta* | g573 | Chr1CC | 5356933 | 5360944 |
| *O. alta* | g17494 | Chr3CC | 5842131 | 5846269 |
| *O. sativa* | Os03g0193225 | Chr3 | 4862250 | 4866734 |
| *O. alta* | g575 | Chr1CC | 5366508 | 5367716 |
| *O. alta* | g17501 | Chr3CC | 5884397 | 5885605 |
| *O. sativa* | Os03g0194100 | Chr3 | 4894537 | 4897016 |
| *O. alta* | g576 | Chr1CC | 5375874 | 5377121 |
| *O. alta* | g17502 | Chr3CC | 5893473 | 5894723 |
| *O. sativa* | Os03g0194300 | Chr3 | 4901969 | 4903749 |
| *O. alta* | g577 | Chr1CC | 5394873 | 5396985 |
| *O. alta* | g17503 | Chr3CC | 5898017 | 5906328 |
| *O. sativa* | Os03g0194400 | Chr3 | 4906251 | 4908791 |
| *O. alta* | g579 | Chr1CC | 5402778 | 5403365 |
| *O. alta* | g17505 | Chr3CC | 5927783 | 5928373 |
| *O. sativa* | Os03g0194600 | Chr3 | 4913314 | 4914063 |
| *O. alta* | g580 | Chr1CC | 5404056 | 5406079 |
| *O. alta* | g17506 | Chr3CC | 5928836 | 5929624 |
| *O. alta* | g581 | Chr1CC | 5415943 | 5417889 |
| *O. alta* | g17508 | Chr3CC | 5932009 | 5932611 |
| *O. alta* | g615 | Chr1CC | 5969599 | 5971400 |
| *O. alta* | g17509 | Chr3CC | 5940807 | 5942614 |
| *O. sativa* | Os03g0194900 | Chr3 | 4923937 | 4926075 |
| *O. alta* | g614 | Chr1CC | 5936526 | 5940334 |
| *O. alta* | g17510 | Chr3CC | 5949233 | 5953658 |
| *O. sativa* | Os03g0195100 | Chr3 | 4936472 | 4941037 |
| *O. alta* | g611 | Chr1CC | 5913505 | 5916679 |
| *O. alta* | g17513 | Chr3CC | 5969916 | 5974332 |
| *O. sativa* | Os03g0195450 | Chr3 | 4954143 | 4954913 |
| *O. alta* | g604 | Chr1CC | 5745773 | 5746714 |
| *O. alta* | g17520 | Chr3CC | 6095541 | 6096482 |

| | | | | |
|---|---|---|---|---|
| *O. sativa* | Os03g0196600 | Chr3 | 5063818 | 5065197 |
| *O. alta* | g609 | Chr1CC | 5880893 | 5884452 |
| *O. alta* | g17515 | Chr3CC | 6007342 | 6011086 |
| *O. sativa* | Os03g0195800 | Chr3 | 4987877 | 4992392 |
| *O. alta* | g607 | Chr1CC | 5832155 | 5832574 |
| *O. alta* | g17517 | Chr3CC | 6033510 | 6033935 |
| *O. sativa* | Os03g0196250 | Chr3 | 5014119 | 5014675 |
| *O. alta* | g606 | Chr1CC | 5793524 | 5798136 |
| *O. alta* | g17518 | Chr3CC | 6056978 | 6062166 |
| *O. sativa* | Os03g0196400 | Chr3 | 5034168 | 5040417 |
| *O. alta* | g605 | Chr1CC | 5767600 | 5768136 |
| *O. alta* | g17519 | Chr3CC | 6068843 | 6069379 |
| *O. sativa* | Os03g0196500 | Chr3 | 5049125 | 5049901 |
| *O. alta* | g603 | Chr1CC | 5737696 | 5740951 |
| *O. alta* | g17521 | Chr3CC | 6112603 | 6116375 |
| *O. sativa* | Os03g0196800 | Chr3 | 5078248 | 5081710 |
| *O. alta* | g602 | Chr1CC | 5733916 | 5736660 |
| *O. alta* | g17522 | Chr3CC | 6117081 | 6119447 |
| *O. sativa* | Os03g0196900 | Chr3 | 5082430 | 5083486 |
| *O. alta* | g601 | Chr1CC | 5674983 | 5676629 |
| *O. alta* | g17524 | Chr3CC | 6134610 | 6136252 |
| *O. sativa* | Os03g0197100 | Chr3 | 5098241 | 5100592 |
| *O. alta* | g599 | Chr1CC | 5650858 | 5652510 |
| *O. alta* | g17525 | Chr3CC | 6155573 | 6157228 |
| *O. sativa* | Os03g0197200 | Chr3 | 5112964 | 5114038 |
| *O. alta* | g598 | Chr1CC | 5644047 | 5647145 |
| *O. alta* | g17526 | Chr3CC | 6159149 | 6161448 |
| *O. sativa* | Os03g0197300 | Chr3 | 5116435 | 5119048 |
| *O. alta* | g596 | Chr1CC | 5623170 | 5625605 |
| *O. alta* | g17528 | Chr3CC | 6177430 | 6179919 |
| *O. sativa* | Os03g0197800 | Chr3 | 5134409 | 5137441 |
| *O. alta* | g594 | Chr1CC | 5598502 | 5599338 |
| *O. alta* | g17529 | Chr3CC | 6224874 | 6225674 |
| *O. sativa* | Os03g0197900 | Chr3 | 5150966 | 5152038 |
| *O. alta* | g593 | Chr1CC | 5589305 | 5593295 |

| | | | | |
|---|---|---|---|---|
| *O. alta* | g17530 | Chr3CC | 6230456 | 6234423 |
| *O. sativa* | Os03g0198300 | Chr3 | 5161642 | 5166608 |
| *O. alta* | g592 | Chr1CC | 5584534 | 5588311 |
| *O. alta* | g17531 | Chr3CC | 6302509 | 6305372 |
| *O. sativa* | Os03g0198400 | Chr3 | 5167307 | 5170694 |
| *O. alta* | g590 | Chr1CC | 5571565 | 5572093 |
| *O. alta* | g17532 | Chr3CC | 6311602 | 6313972 |
| *O. sativa* | Os03g0198500 | Chr3 | 5175138 | 5179221 |
| *O. alta* | g589 | Chr1CC | 5568560 | 5569502 |
| *O. alta* | g17533 | Chr3CC | 6316593 | 6317526 |
| *O. sativa* | Os03g0198600 | Chr3 | 5180779 | 5182185 |
| *O. alta* | g587 | Chr1CC | 5532528 | 5532911 |
| *O. alta* | g17535 | Chr3CC | 6347340 | 6347714 |
| *O. alta* | g586 | Chr1CC | 5522302 | 5523471 |
| *O. alta* | g17536 | Chr3CC | 6356037 | 6357167 |
| *O. sativa* | Os03g0199100 | Chr3 | 5214115 | 5217654 |
| *O. alta* | g584 | Chr1CC | 5501365 | 5502294 |
| *O. alta* | g17537 | Chr3CC | 6365122 | 6366056 |
| *O. sativa* | Os03g0199500 | Chr3 | 5225077 | 5226496 |
| *O. alta* | g732 | Chr1CC | 6891858 | 6893114 |
| *O. alta* | g17538 | Chr3CC | 6387341 | 6388601 |
| *O. sativa* | Os03g0200000 | Chr3 | 5253519 | 5254758 |
| *O. alta* | g731 | Chr1CC | 6884991 | 6886318 |
| *O. alta* | g17539 | Chr3CC | 6401756 | 6403130 |
| *O. sativa* | Os03g0200200 | Chr3 | 5261710 | 5263669 |
| *O. alta* | g730 | Chr1CC | 6873547 | 6873956 |
| *O. alta* | g17540 | Chr3CC | 6406912 | 6407325 |
| *O. sativa* | Os03g0200400 | Chr3 | 5271246 | 5272329 |
| *O. alta* | g728 | Chr1CC | 6857510 | 6859409 |
| *O. alta* | g17541 | Chr3CC | 6411871 | 6414117 |
| *O. sativa* | Os03g0200500 | Chr3 | 5274929 | 5277136 |
| *O. alta* | g727 | Chr1CC | 6853672 | 6856735 |
| *O. alta* | g17542 | Chr3CC | 6414886 | 6417929 |
| *O. alta* | g726 | Chr1CC | 6844226 | 6844930 |
| *O. alta* | g17543 | Chr3CC | 6418598 | 6424274 |

| | | | | |
|---|---|---|---|---|
| *O. sativa* | Os03g0200700 | Chr3 | 5283459 | 5286844 |
| *O. alta* | g725 | Chr1CC | 6840595 | 6842918 |
| *O. alta* | g17544 | Chr3CC | 6425182 | 6427403 |
| *O. sativa* | Os03g0200800 | Chr3 | 5287681 | 5290481 |
| *O. alta* | g723 | Chr1CC | 6824907 | 6827067 |
| *O. alta* | g17545 | Chr3CC | 6430135 | 6432373 |
| *O. sativa* | Os03g0201000 | Chr3 | 5294523 | 5297017 |
| *O. alta* | g722 | Chr1CC | 6820422 | 6823561 |
| *O. alta* | g17546 | Chr3CC | 6433724 | 6436848 |
| *O. sativa* | Os03g0201100 | Chr3 | 5297336 | 5301348 |
| *O. alta* | g721 | Chr1CC | 6818086 | 6820082 |
| *O. alta* | g17547 | Chr3CC | 6437219 | 6439672 |
| *O. sativa* | Os03g0201200 | Chr3 | 5302428 | 5304763 |
| *O. alta* | g720 | Chr1CC | 6813232 | 6815616 |
| *O. alta* | g17548 | Chr3CC | 6440199 | 6442586 |
| *O. sativa* | Os03g0201400 | Chr3 | 5304942 | 5314273 |
| *O. alta* | g719 | Chr1CC | 6805256 | 6808836 |
| *O. alta* | g17549 | Chr3CC | 6447530 | 6450316 |
| *O. sativa* | Os03g0201500 | Chr3 | 5317215 | 5321081 |
| *O. alta* | g718 | Chr1CC | 6798653 | 6801661 |
| *O. alta* | g17551 | Chr3CC | 6453880 | 6456605 |
| *O. sativa* | Os03g0201700 | Chr3 | 5324635 | 5327751 |
| *O. alta* | g717 | Chr1CC | 6792976 | 6797702 |
| *O. alta* | g17555 | Chr3CC | 6463697 | 6467775 |
| *O. sativa* | Os03g0201901 | Chr3 | 5331153 | 5335603 |
| *O. alta* | g715 | Chr1CC | 6769977 | 6773065 |
| *O. alta* | g17558 | Chr3CC | 6478498 | 6481641 |
| *O. sativa* | Os03g0202200 | Chr3 | 5348313 | 5351951 |
| *O. alta* | g714 | Chr1CC | 6760139 | 6768909 |
| *O. alta* | g17559 | Chr3CC | 6482737 | 6487153 |
| *O. sativa* | Os03g0202300 | Chr3 | 5352944 | 5357283 |
| *O. alta* | g713 | Chr1CC | 6738986 | 6740032 |
| *O. alta* | g17562 | Chr3CC | 6524740 | 6525781 |
| *O. sativa* | Os03g0203200 | Chr3 | 5422148 | 5426577 |
| *O. alta* | g710 | Chr1CC | 6725195 | 6730064 |

| | | | | |
|---|---|---|---|---|
| *O. alta* | g17564 | Chr3CC | 6561383 | 6566380 |
| *O. sativa* | Os03g0203700 | Chr3 | 5433914 | 5440324 |
| *O. alta* | g709 | Chr1CC | 6721334 | 6722800 |
| *O. alta* | g17563 | Chr3CC | 6557431 | 6558914 |
| *O. sativa* | Os03g0203800 | Chr3 | 5441845 | 5444188 |
| *O. alta* | g708 | Chr1CC | 6707254 | 6711051 |
| *O. alta* | g17566 | Chr3CC | 6577435 | 6581163 |
| *O. sativa* | Os03g0204100 | Chr3 | 5458099 | 5462322 |
| *O. alta* | g707 | Chr1CC | 6700916 | 6701240 |
| *O. alta* | g17572 | Chr3CC | 6625327 | 6625649 |
| *O. sativa* | Os03g0204900 | Chr3 | 5481504 | 5482358 |
| *O. alta* | g705 | Chr1CC | 6685679 | 6690972 |
| *O. alta* | g17573 | Chr3CC | 6626790 | 6632142 |
| *O. sativa* | Os03g0205000 | Chr3 | 5483405 | 5489241 |
| *O. alta* | g703 | Chr1CC | 6664198 | 6667893 |
| *O. alta* | g17574 | Chr3CC | 6635299 | 6636836 |
| *O. sativa* | Os03g0205150 | Chr3 | 5491325 | 5494409 |
| *O. alta* | g702 | Chr1CC | 6655508 | 6656965 |
| *O. alta* | g17575 | Chr3CC | 6641412 | 6642869 |
| *O. sativa* | Os03g0205300 | Chr3 | 5495851 | 5497269 |
| *O. alta* | g701 | Chr1CC | 6638537 | 6646432 |
| *O. alta* | g17576 | Chr3CC | 6653312 | 6661185 |
| *O. sativa* | Os03g0205400 | Chr3 | 5508753 | 5516754 |
| *O. alta* | g699 | Chr1CC | 6619730 | 6622526 |
| *O. alta* | g17578 | Chr3CC | 6677249 | 6680075 |
| *O. sativa* | Os03g0205700 | Chr3 | 5530670 | 5534319 |
| *O. alta* | g698 | Chr1CC | 6604831 | 6606307 |
| *O. alta* | g17579 | Chr3CC | 6727596 | 6729039 |
| *O. sativa* | Os03g0205800 | Chr3 | 5538810 | 5540247 |
| *O. alta* | g697 | Chr1CC | 6594327 | 6598418 |
| *O. alta* | g17580 | Chr3CC | 6735963 | 6740058 |
| *O. alta* | g696 | Chr1CC | 6575126 | 6577886 |
| *O. alta* | g17581 | Chr3CC | 6763619 | 6766380 |
| *O. sativa* | Os03g0206300 | Chr3 | 5573373 | 5576434 |
| *O. alta* | g695 | Chr1CC | 6568320 | 6568691 |

| | | | | |
|---|---|---|---|---|
| *O. alta* | g17583 | Chr3CC | 6789569 | 6789936 |
| *O. sativa* | Os03g0206400 | Chr3 | 5582059 | 5582713 |
| *O. alta* | g694 | Chr1CC | 6559482 | 6563201 |
| *O. alta* | g17584 | Chr3CC | 6794091 | 6796729 |
| *O. sativa* | Os03g0206600 | Chr3 | 5586002 | 5588859 |
| *O. alta* | g693 | Chr1CC | 6555605 | 6558287 |
| *O. alta* | g17585 | Chr3CC | 6797895 | 6800400 |
| *O. sativa* | Os03g0206700 | Chr3 | 5589846 | 5592643 |
| *O. alta* | g692 | Chr1CC | 6551939 | 6553650 |
| *O. alta* | g17586 | Chr3CC | 6804223 | 6805927 |
| *O. sativa* | Os03g0206900 | Chr3 | 5594051 | 5595994 |
| *O. alta* | g691 | Chr1CC | 6536491 | 6539427 |
| *O. alta* | g17588 | Chr3CC | 6807919 | 6810757 |
| *O. alta* | g688 | Chr1CC | 6515487 | 6517061 |
| *O. alta* | g17591 | Chr3CC | 6824235 | 6825855 |
| *O. sativa* | Os03g0207250 | Chr3 | 5613333 | 5615437 |
| *O. alta* | g690 | Chr1CC | 6523537 | 6526257 |
| *O. alta* | g17589 | Chr3CC | 6816983 | 6819655 |
| *O. alta* | g689 | Chr1CC | 6520771 | 6522366 |
| *O. alta* | g17590 | Chr3CC | 6820284 | 6822566 |
| *O. sativa* | Os03g0207200 | Chr3 | 5609827 | 5612733 |
| *O. alta* | g686 | Chr1CC | 6505301 | 6506668 |
| *O. alta* | g17594 | Chr3CC | 6835813 | 6837211 |
| *O. sativa* | Os03g0207400 | Chr3 | 5625733 | 5627711 |
| *O. alta* | g685 | Chr1CC | 6497695 | 6499899 |
| *O. alta* | g17595 | Chr3CC | 6841902 | 6843893 |
| *O. sativa* | Os03g0207800 | Chr3 | 5641293 | 5643559 |
| *O. alta* | g684 | Chr1CC | 6491408 | 6494863 |
| *O. alta* | g17596 | Chr3CC | 6846555 | 6850061 |
| *O. sativa* | Os03g0207900 | Chr3 | 5646348 | 5650212 |
| *O. alta* | g683 | Chr1CC | 6483279 | 6486844 |
| *O. alta* | g17597 | Chr3CC | 6852780 | 6856464 |
| *O. sativa* | Os03g0208500 | Chr3 | 5655157 | 5659172 |
| *O. alta* | g682 | Chr1CC | 6466347 | 6468083 |
| *O. alta* | g17598 | Chr3CC | 6865635 | 6867224 |

| | | | | |
|---|---|---|---|---|
| *O. sativa* | Os03g0208600 | Chr3 | 5669927 | 5671969 |
| *O. alta* | g679 | Chr1CC | 6453739 | 6454209 |
| *O. alta* | g17604 | Chr3CC | 6904739 | 6905221 |
| *O. sativa* | Os03g0209000 | Chr3 | 5686617 | 5687556 |
| *O. alta* | g680 | Chr1CC | 6455657 | 6460927 |
| *O. alta* | g17603 | Chr3CC | 6899149 | 6903364 |
| *O. sativa* | Os03g0208900 | Chr3 | 5680647 | 5685342 |
| *O. alta* | g675 | Chr1CC | 6419981 | 6421626 |
| *O. alta* | g17605 | Chr3CC | 6918627 | 6920732 |
| *O. sativa* | Os03g0209400 | Chr3 | 5707404 | 5710078 |
| *O. alta* | g674 | Chr1CC | 6414690 | 6418815 |
| *O. alta* | g17606 | Chr3CC | 6921752 | 6925902 |
| *O. sativa* | Os03g0209500 | Chr3 | 5710240 | 5715009 |
| *O. alta* | g673 | Chr1CC | 6402454 | 6409585 |
| *O. alta* | g17607 | Chr3CC | 6932499 | 6939773 |
| *O. sativa* | Os03g0209600 | Chr3 | 5721104 | 5728581 |
| *O. alta* | g670 | Chr1CC | 6377506 | 6381334 |
| *O. alta* | g17610 | Chr3CC | 6957476 | 6958153 |
| *O. alta* | g667 | Chr1CC | 6360475 | 6360831 |
| *O. alta* | g17612 | Chr3CC | 6968258 | 6968629 |
| *O. sativa* | Os03g0210100 | Chr3 | 5753283 | 5753636 |
| *O. alta* | g666 | Chr1CC | 6358297 | 6358644 |
| *O. alta* | g17615 | Chr3CC | 6987106 | 6987456 |
| *O. sativa* | Os03g0210200 | Chr3 | 5755354 | 5755913 |
| *O. alta* | g665 | Chr1CC | 6356533 | 6357675 |
| *O. alta* | g17616 | Chr3CC | 6988192 | 6989504 |
| *O. alta* | g664 | Chr1CC | 6348537 | 6354676 |
| *O. alta* | g17617 | Chr3CC | 6990733 | 6996852 |
| *O. sativa* | Os03g0210400 | Chr3 | 5760848 | 5767440 |
| *O. alta* | g662 | Chr1CC | 6328900 | 6331162 |
| *O. alta* | g17619 | Chr3CC | 7003912 | 7006101 |
| *O. sativa* | Os03g0210500 | Chr3 | 5771944 | 5772406 |
| *O. alta* | g661 | Chr1CC | 6324688 | 6325797 |
| *O. alta* | g17620 | Chr3CC | 7008841 | 7009989 |
| *O. sativa* | Os03g0210600 | Chr3 | 5774735 | 5775940 |

| | | | | |
|---|---|---|---|---|
| *O. alta* | g660 | Chr1CC | 6322313 | 6323245 |
| *O. alta* | g17621 | Chr3CC | 7011256 | 7012510 |
| *O. sativa* | Os03g0210700 | Chr3 | 5777776 | 5779292 |
| *O. alta* | g659 | Chr1CC | 6312241 | 6320884 |
| *O. alta* | g17622 | Chr3CC | 7014903 | 7022332 |
| *O. sativa* | Os03g0210800 | Chr3 | 5779625 | 5784938 |
| *O. alta* | g657 | Chr1CC | 6302405 | 6303187 |
| *O. alta* | g17624 | Chr3CC | 7033067 | 7033849 |
| *O. sativa* | Os03g0211100 | Chr3 | 5798558 | 5801146 |
| *O. alta* | g658 | Chr1CC | 6307480 | 6307933 |
| *O. alta* | g17623 | Chr3CC | 7029595 | 7030056 |
| *O. sativa* | Os03g0210900 | Chr3 | 5796329 | 5797044 |
| *O. alta* | g656 | Chr1CC | 6300657 | 6301778 |
| *O. alta* | g17626 | Chr3CC | 7044934 | 7052291 |
| *O. alta* | g655 | Chr1CC | 6287904 | 6290219 |
| *O. alta* | g17628 | Chr3CC | 7068161 | 7070482 |
| *O. sativa* | Os03g0211400 | Chr3 | 5807868 | 5812643 |
| *O. alta* | g653 | Chr1CC | 6281037 | 6283485 |
| *O. alta* | g17630 | Chr3CC | 7074837 | 7077231 |
| *O. sativa* | Os03g0211600 | Chr3 | 5817877 | 5819374 |
| *O. alta* | g652 | Chr1CC | 6272441 | 6275837 |
| *O. alta* | g17631 | Chr3CC | 7090673 | 7094038 |
| *O. sativa* | Os03g0211700 | Chr3 | 5823655 | 5828110 |
| *O. alta* | g651 | Chr1CC | 6265458 | 6270521 |
| *O. alta* | g17632 | Chr3CC | 7096466 | 7101561 |
| *O. sativa* | Os03g0211800 | Chr3 | 5829016 | 5835122 |
| *O. alta* | g650 | Chr1CC | 6253170 | 6257019 |
| *O. alta* | g17633 | Chr3CC | 7107819 | 7112180 |
| *O. sativa* | Os03g0211900 | Chr3 | 5840714 | 5844831 |
| *O. alta* | g649 | Chr1CC | 6247999 | 6249498 |
| *O. alta* | g17634 | Chr3CC | 7117979 | 7119457 |
| *O. sativa* | Os03g0212000 | Chr3 | 5848833 | 5850383 |
| *O. alta* | g648 | Chr1CC | 6243934 | 6247278 |
| *O. alta* | g17635 | Chr3CC | 7120411 | 7123699 |
| *O. sativa* | Os03g0212200 | Chr3 | 5853962 | 5857465 |

| | | | | |
|---|---|---|---|---|
| *O. alta* | g646 | Chr1CC | 6241731 | 6242024 |
| *O. alta* | g17636 | Chr3CC | 7124822 | 7125939 |
| *O. sativa* | Os03g0212300 | Chr3 | 5858227 | 5859977 |
| *O. alta* | g643 | Chr1CC | 6234855 | 6235115 |
| *O. alta* | g17637 | Chr3CC | 7127537 | 7127794 |
| *O. alta* | g642 | Chr1CC | 6232350 | 6233776 |
| *O. alta* | g17638 | Chr3CC | 7128960 | 7130406 |
| *O. sativa* | Os03g0212400 | Chr3 | 5860717 | 5863891 |
| *O. alta* | g641 | Chr1CC | 6230801 | 6231139 |
| *O. alta* | g17639 | Chr3CC | 7131952 | 7132296 |
| *O. alta* | g640 | Chr1CC | 6226540 | 6229313 |
| *O. alta* | g17640 | Chr3CC | 7133240 | 7136450 |
| *O. sativa* | Os03g0212600 | Chr3 | 5865638 | 5869191 |
| *O. alta* | g639 | Chr1CC | 6220472 | 6224579 |
| *O. alta* | g17641 | Chr3CC | 7137615 | 7141309 |
| *O. sativa* | Os03g0212700 | Chr3 | 5870156 | 5874813 |
| *O. alta* | g638 | Chr1CC | 6211967 | 6218254 |
| *O. alta* | g17642 | Chr3CC | 7144996 | 7150278 |
| *O. sativa* | Os03g0212800 | Chr3 | 5876410 | 5883222 |
| *O. alta* | g637 | Chr1CC | 6193515 | 6195509 |
| *O. alta* | g17643 | Chr3CC | 7156274 | 7158277 |
| *O. sativa* | Os03g0213100 | Chr3 | 5896038 | 5899522 |
| *O. alta* | g636 | Chr1CC | 6190600 | 6192701 |
| *O. alta* | g17644 | Chr3CC | 7168165 | 7168464 |
| *O. sativa* | Os03g0213200 | Chr3 | 5900087 | 5902461 |
| *O. alta* | g635 | Chr1CC | 6176983 | 6183378 |
| *O. alta* | g17645 | Chr3CC | 7177986 | 7184287 |
| *O. sativa* | Os03g0213300 | Chr3 | 5907703 | 5917030 |
| *O. alta* | g634 | Chr1CC | 6154122 | 6173125 |
| *O. alta* | g17646 | Chr3CC | 7190072 | 7208420 |
| *O. sativa* | Os03g0213400 | Chr3 | 5920111 | 5931373 |
| *O. alta* | g632 | Chr1CC | 6147452 | 6147946 |
| *O. alta* | g17647 | Chr3CC | 7211099 | 7211587 |
| *O. sativa* | Os03g0213500 | Chr3 | 5941804 | 5943541 |

| | | | | |
|---|---|---|---|---|
| *O. alta* | g631 | Chr1CC | 6144893 | 6145093 |
| *O. alta* | g17648 | Chr3CC | 7213258 | 7221849 |
| *O. sativa* | Os03g0213600 | Chr3 | 5946380 | 5955447 |
| *O. alta* | g629 | Chr1CC | 6131154 | 6135724 |
| *O. alta* | g17649 | Chr3CC | 7224662 | 7226851 |
| *O. sativa* | Os03g0213700 | Chr3 | 5955407 | 5960532 |
| *O. alta* | g628 | Chr1CC | 6126729 | 6130221 |
| *O. alta* | g17650 | Chr3CC | 7227580 | 7231244 |
| *O. sativa* | Os03g0213800 | Chr3 | 5960793 | 5964717 |
| *O. alta* | g627 | Chr1CC | 6108843 | 6115412 |
| *O. alta* | g17651 | Chr3CC | 7240069 | 7244015 |
| *O. sativa* | Os03g0213900 | Chr3 | 5967215 | 5973703 |
| *O. alta* | g626 | Chr1CC | 6103857 | 6108117 |
| *O. alta* | g17653 | Chr3CC | 7246948 | 7251327 |
| *O. sativa* | Os03g0214000 | Chr3 | 5973919 | 5979076 |
| *O. alta* | g625 | Chr1CC | 6099983 | 6103114 |
| *O. alta* | g17654 | Chr3CC | 7253904 | 7256991 |
| *O. sativa* | Os03g0214100 | Chr3 | 5979594 | 5983020 |
| *O. alta* | g624 | Chr1CC | 6090482 | 6095311 |
| *O. alta* | g17655 | Chr3CC | 7258001 | 7263036 |
| *O. sativa* | Os03g0214200 | Chr3 | 5983462 | 5988740 |
| *O. alta* | g623 | Chr1CC | 6079465 | 6082154 |
| *O. alta* | g17656 | Chr3CC | 7266494 | 7269163 |
| *O. sativa* | Os03g0214400 | Chr3 | 5994179 | 5997434 |
| *O. alta* | g620 | Chr1CC | 6069406 | 6073106 |
| *O. alta* | g17657 | Chr3CC | 7272158 | 7275544 |
| *O. sativa* | Os03g0214600 | Chr3 | 5999908 | 6003768 |
| *O. alta* | g619 | Chr1CC | 6064080 | 6068021 |
| *O. alta* | g17659 | Chr3CC | 7277843 | 7282386 |
| *O. sativa* | Os03g0214900 | Chr3 | 6006751 | 6011719 |
| *O. alta* | g616 | Chr1CC | 6019888 | 6025277 |
| *O. alta* | g17661 | Chr3CC | 7311794 | 7317050 |
| *O. sativa* | Os03g0215200 | Chr3 | 6041245 | 6048687 |
| *O. alta* | g735 | Chr1CC | 6963062 | 6967023 |
| *O. alta* | g17664 | Chr3CC | 7348990 | 7353228 |
| *O. sativa* | Os03g0215700 | Chr3 | 6079161 | 6084477 |

| | | | | |
|---|---|---|---|---|
| *O. alta* | g736 | Chr1CC | 6968732 | 6971703 |
| *O. alta* | g17665 | Chr3CC | 7362176 | 7365114 |
| *O. sativa* | Os03g0215800 | Chr3 | 6085640 | 6089018 |
| *O. alta* | g737 | Chr1CC | 6972849 | 6974873 |
| *O. alta* | g17666 | Chr3CC | 7365655 | 7368412 |
| *O. sativa* | Os03g0215900 | Chr3 | 6089248 | 6091523 |
| *O. alta* | g738 | Chr1CC | 6982205 | 6983074 |
| *O. alta* | g17667 | Chr3CC | 7375154 | 7376053 |
| *O. sativa* | Os03g0216000 | Chr3 | 6098433 | 6100341 |
| *O. alta* | g740 | Chr1CC | 7004155 | 7005444 |
| *O. alta* | g17669 | Chr3CC | 7392408 | 7393250 |
| *O. sativa* | Os03g0216400 | Chr3 | 6118856 | 6120196 |
| *O. alta* | g741 | Chr1CC | 7015242 | 7016288 |
| *O. alta* | g17670 | Chr3CC | 7395218 | 7396107 |
| *O. sativa* | Os03g0216500 | Chr3 | 6121324 | 6122376 |
| *O. alta* | g742 | Chr1CC | 7017442 | 7021153 |
| *O. alta* | g17671 | Chr3CC | 7399664 | 7403974 |
| *O. sativa* | Os03g0216600 | Chr3 | 6127334 | 6131136 |
| *O. alta* | g743 | Chr1CC | 7022423 | 7026370 |
| *O. alta* | g17672 | Chr3CC | 7405238 | 7409168 |
| *O. sativa* | Os03g0216700 | Chr3 | 6131846 | 6142781 |
| *O. alta* | g744 | Chr1CC | 7039176 | 7041811 |
| *O. alta* | g17673 | Chr3CC | 7422482 | 7425056 |
| *O. sativa* | Os03g0216800 | Chr3 | 6152114 | 6155182 |
| *O. alta* | g745 | Chr1CC | 7044880 | 7056330 |
| *O. alta* | g17674 | Chr3CC | 7428023 | 7432319 |
| *O. sativa* | Os03g0216900 | Chr3 | 6159948 | 6163869 |
| *O. alta* | g746 | Chr1CC | 7056885 | 7057280 |
| *O. alta* | g17675 | Chr3CC | 7432877 | 7433281 |
| *O. sativa* | Os03g0217000 | Chr3 | 6163963 | 6164560 |
| *O. alta* | g747 | Chr1CC | 7092386 | 7097484 |
| *O. alta* | g17676 | Chr3CC | 7448334 | 7453452 |
| *O. sativa* | Os03g0218300 | Chr3 | 6227695 | 6229743 |
| *O. alta* | g748 | Chr1CC | 7098108 | 7102405 |
| *O. alta* | g17677 | Chr3CC | 7454317 | 7462292 |

| | | | | |
|---|---|---|---|---|
| *O. sativa* | Os03g0218400 | Chr3 | 6233563 | 6238297 |
| *O. alta* | g751 | Chr1CC | 7133409 | 7136130 |
| *O. alta* | g17678 | Chr3CC | 7478768 | 7481458 |
| *O. sativa* | Os03g0219100 | Chr3 | 6264770 | 6267856 |
| *O. alta* | g752 | Chr1CC | 7139493 | 7142581 |
| *O. alta* | g17679 | Chr3CC | 7484814 | 7488501 |
| *O. sativa* | Os03g0219200 | Chr3 | 6271895 | 6275386 |
| *O. alta* | g754 | Chr1CC | 7167449 | 7169349 |
| *O. alta* | g17681 | Chr3CC | 7495629 | 7497538 |
| *O. sativa* | Os03g0219400 | Chr3 | 6280458 | 6282712 |
| *O. alta* | g755 | Chr1CC | 7169538 | 7170752 |
| *O. alta* | g17682 | Chr3CC | 7498646 | 7498954 |
| *O. sativa* | Os03g0219500 | Chr3 | 6282872 | 6284348 |
| *O. alta* | g756 | Chr1CC | 7172915 | 7174636 |
| *O. alta* | g17683 | Chr3CC | 7500536 | 7502257 |
| *O. sativa* | Os03g0219700 | Chr3 | 6286013 | 6288421 |
| *O. alta* | g757 | Chr1CC | 7203033 | 7206783 |
| *O. alta* | g17684 | Chr3CC | 7512522 | 7516294 |
| *O. sativa* | Os03g0219800 | Chr3 | 6290732 | 6295238 |
| *O. alta* | g758 | Chr1CC | 7207400 | 7209362 |
| *O. alta* | g17685 | Chr3CC | 7516925 | 7518914 |
| *O. sativa* | Os03g0219900 | Chr3 | 6295480 | 6297596 |
| *O. alta* | g760 | Chr1CC | 7213164 | 7214754 |
| *O. alta* | g17686 | Chr3CC | 7523437 | 7525026 |
| *O. sativa* | Os03g0220100 | Chr3 | 6302865 | 6304849 |
| *O. alta* | g762 | Chr1CC | 7292206 | 7294605 |
| *O. alta* | g17688 | Chr3CC | 7566702 | 7569144 |
| *O. sativa* | Os03g0221200 | Chr3 | 6354859 | 6357792 |
| *O. alta* | g763 | Chr1CC | 7295566 | 7307115 |
| *O. alta* | g17689 | Chr3CC | 7570056 | 7577277 |
| *O. sativa* | Os03g0221300 | Chr3 | 6358095 | 6364239 |
| *O. alta* | g764 | Chr1CC | 7325755 | 7329200 |
| *O. alta* | g17690 | Chr3CC | 7583173 | 7586602 |
| *O. sativa* | Os03g0221500 | Chr3 | 6369452 | 6373704 |
| *O. alta* | g765 | Chr1CC | 7333400 | 7335910 |

| | | | | |
|---|---|---|---|---|
| *O. alta* | g17691 | Chr3CC | 7589983 | 7592508 |
| *O. sativa* | Os03g0221700 | Chr3 | 6377414 | 6380243 |
| *O. alta* | g767 | Chr1CC | 7360958 | 7362946 |
| *O. alta* | g17693 | Chr3CC | 7609698 | 7611692 |
| *O. sativa* | Os03g0222100 | Chr3 | 6392811 | 6395478 |
| *O. alta* | g770 | Chr1CC | 7374618 | 7375819 |
| *O. alta* | g17697 | Chr3CC | 7644997 | 7645512 |
| *O. sativa* | Os03g0222500 | Chr3 | 6405347 | 6406081 |
| *O. alta* | g772 | Chr1CC | 7387761 | 7390097 |
| *O. alta* | g17699 | Chr3CC | 7650043 | 7652314 |
| *O. sativa* | Os03g0222600 | Chr3 | 6418114 | 6420549 |
| *O. alta* | g773 | Chr1CC | 7394363 | 7398573 |
| *O. alta* | g17700 | Chr3CC | 7653731 | 7658728 |
| *O. sativa* | Os03g0222800 | Chr3 | 6424393 | 6426901 |
| *O. alta* | g774 | Chr1CC | 7400337 | 7402506 |
| *O. alta* | g17701 | Chr3CC | 7659654 | 7668105 |
| *O. alta* | g775 | Chr1CC | 7405528 | 7408614 |
| *O. alta* | g17703 | Chr3CC | 7674531 | 7677632 |
| *O. sativa* | Os03g0223000 | Chr3 | 6428279 | 6438500 |
| *O. alta* | g776 | Chr1CC | 7431574 | 7433157 |
| *O. alta* | g17705 | Chr3CC | 7695006 | 7696586 |
| *O. sativa* | Os03g0223100 | Chr3 | 6443453 | 6445461 |
| *O. alta* | g777 | Chr1CC | 7440151 | 7442218 |
| *O. alta* | g17706 | Chr3CC | 7707864 | 7710338 |
| *O. sativa* | Os03g0223200 | Chr3 | 6453022 | 6455453 |
| *O. alta* | g778 | Chr1CC | 7445177 | 7448652 |
| *O. alta* | g17707 | Chr3CC | 7711093 | 7715317 |
| *O. sativa* | Os03g0223400 | Chr3 | 6457915 | 6462146 |
| *O. alta* | g779 | Chr1CC | 7457183 | 7464178 |
| *O. alta* | g17708 | Chr3CC | 7720006 | 7727075 |
| *O. sativa* | Os03g0223700 | Chr3 | 6468343 | 6476409 |
| *O. alta* | g782 | Chr1CC | 7500614 | 7501199 |
| *O. alta* | g17711 | Chr3CC | 7765038 | 7769796 |
| *O. sativa* | Os03g0223900 | Chr3 | 6493908 | 6498974 |
| *O. alta* | g783 | Chr1CC | 7513327 | 7517709 |

| | | | | |
|---|---|---|---|---|
| *O. alta* | g17712 | Chr3CC | 7777533 | 7781885 |
| *O. sativa* | Os03g0224200 | Chr3 | 6512771 | 6518511 |
| *O. alta* | g784 | Chr1CC | 7519402 | 7522432 |
| *O. alta* | g17713 | Chr3CC | 7783415 | 7786446 |
| *O. sativa* | Os03g0224300 | Chr3 | 6519054 | 6522132 |
| *O. alta* | g785 | Chr1CC | 7568061 | 7570636 |
| *O. alta* | g17715 | Chr3CC | 7800620 | 7803549 |
| *O. sativa* | Os03g0224700 | Chr3 | 6537573 | 6541381 |
| *O. alta* | g787 | Chr1CC | 7577192 | 7578238 |
| *O. alta* | g17716 | Chr3CC | 7809691 | 7810707 |
| *O. sativa* | Os03g0225100 | Chr3 | 6545153 | 6546478 |
| *O. alta* | g788 | Chr1CC | 7590462 | 7594283 |
| *O. alta* | g17717 | Chr3CC | 7819056 | 7823699 |
| *O. sativa* | Os03g0225200 | Chr3 | 6556389 | 6562025 |
| *O. alta* | g789 | Chr1CC | 7596906 | 7598324 |
| *O. alta* | g17718 | Chr3CC | 7825503 | 7826918 |
| *O. sativa* | Os03g0225300 | Chr3 | 6563392 | 6565219 |
| *O. alta* | g791 | Chr1CC | 7611918 | 7618956 |
| *O. alta* | g17720 | Chr3CC | 7834988 | 7842084 |
| *O. sativa* | Os03g0225500 | Chr3 | 6573239 | 6580610 |
| *O. alta* | g792 | Chr1CC | 7620850 | 7627104 |
| *O. alta* | g17722 | Chr3CC | 7844103 | 7850136 |
| *O. sativa* | Os03g0225600 | Chr3 | 6587240 | 6587714 |
| *O. alta* | g793 | Chr1CC | 7629307 | 7633836 |
| *O. alta* | g17723 | Chr3CC | 7853408 | 7857570 |
| *O. sativa* | Os03g0225700 | Chr3 | 6589784 | 6594037 |
| *O. alta* | g794 | Chr1CC | 7644362 | 7645798 |
| *O. alta* | g17724 | Chr3CC | 7863010 | 7864449 |
| *O. sativa* | Os03g0225900 | Chr3 | 6608577 | 6610549 |
| *O. alta* | g796 | Chr1CC | 7671985 | 7673841 |
| *O. alta* | g17725 | Chr3CC | 7886642 | 7887442 |
| *O. sativa* | Os03g0226300 | Chr3 | 6626961 | 6630244 |
| *O. alta* | g801 | Chr1CC | 7716000 | 7720363 |
| *O. alta* | g17726 | Chr3CC | 7913371 | 7917370 |
| *O. sativa* | Os03g0226400 | Chr3 | 6636675 | 6641550 |

| | | | | |
|---|---|---|---|---|
| *O. alta* | g800 | Chr1CC | 7700011 | 7706154 |
| *O. alta* | g17728 | Chr3CC | 7924454 | 7930519 |
| *O. sativa* | Os03g0226600 | Chr3 | 6647026 | 6653918 |
| *O. alta* | g799 | Chr1CC | 7696146 | 7697904 |
| *O. alta* | g17729 | Chr3CC | 7932232 | 7934157 |
| *O. sativa* | Os03g0226700 | Chr3 | 6654751 | 6656679 |
| *O. alta* | g798 | Chr1CC | 7681829 | 7682200 |
| *O. alta* | g17730 | Chr3CC | 7935383 | 7935853 |
| *O. alta* | g797 | Chr1CC | 7677485 | 7681795 |
| *O. alta* | g17731 | Chr3CC | 7935953 | 7945411 |
| *O. sativa* | Os03g0226800 | Chr3 | 6657601 | 6663830 |
| *O. alta* | g804 | Chr1CC | 7741735 | 7748308 |
| *O. alta* | g17732 | Chr3CC | 7948761 | 7955273 |
| *O. sativa* | Os03g0227000 | Chr3 | 6676410 | 6683840 |
| *O. alta* | g805 | Chr1CC | 7754077 | 7761395 |
| *O. alta* | g17733 | Chr3CC | 7957450 | 7965506 |
| *O. sativa* | Os03g0227300 | Chr3 | 6698297 | 6706333 |
| *O. alta* | g806 | Chr1CC | 7762033 | 7765485 |
| *O. alta* | g17734 | Chr3CC | 7965939 | 7969339 |
| *O. sativa* | Os03g0227400 | Chr3 | 6706278 | 6708143 |
| *O. alta* | g807 | Chr1CC | 7774620 | 7778566 |
| *O. alta* | g17735 | Chr3CC | 7976341 | 7982884 |
| *O. sativa* | Os03g0227500 | Chr3 | 6716409 | 6720052 |
| *O. alta* | g808 | Chr1CC | 7796418 | 7802487 |
| *O. alta* | g17737 | Chr3CC | 8002485 | 8009655 |
| *O. sativa* | Os03g0227700 | Chr3 | 6737549 | 6744486 |
| *O. alta* | g811 | Chr1CC | 7832751 | 7836431 |
| *O. alta* | g17738 | Chr3CC | 8015726 | 8020091 |
| *O. sativa* | Os03g0227900 | Chr3 | 6763115 | 6767691 |
| *O. alta* | g812 | Chr1CC | 7842856 | 7843788 |
| *O. alta* | g17740 | Chr3CC | 8054678 | 8055118 |
| *O. alta* | g813 | Chr1CC | 7849088 | 7850041 |
| *O. alta* | g17741 | Chr3CC | 8061667 | 8062640 |
| *O. sativa* | Os03g0228200 | Chr3 | 6776463 | 6778596 |
| *O. alta* | g814 | Chr1CC | 7860364 | 7862536 |

| | | | | |
|---|---|---|---|---|
| *O. alta* | g17742 | Chr3CC | 8072623 | 8074840 |
| *O. sativa* | Os03g0228400 | Chr3 | 6782702 | 6785978 |
| *O. alta* | g815 | Chr1CC | 7868825 | 7873109 |
| *O. alta* | g17743 | Chr3CC | 8077109 | 8080339 |
| *O. sativa* | Os03g0228500 | Chr3 | 6786712 | 6790137 |
| *O. alta* | g817 | Chr1CC | 7912885 | 7916385 |
| *O. alta* | g17744 | Chr3CC | 8091832 | 8095852 |
| *O. sativa* | Os03g0228800 | Chr3 | 6804619 | 6808346 |
| *O. alta* | g819 | Chr1CC | 7929177 | 7930159 |
| *O. alta* | g17746 | Chr3CC | 8126567 | 8128415 |
| *O. sativa* | Os03g0229100 | Chr3 | 6826703 | 6832274 |
| *O. alta* | g820 | Chr1CC | 7975168 | 7977413 |
| *O. alta* | g17747 | Chr3CC | 8152454 | 8154723 |
| *O. alta* | g821 | Chr1CC | 7995074 | 8003306 |
| *O. alta* | g17750 | Chr3CC | 8180688 | 8184381 |
| *O. sativa* | Os03g0229600 | Chr3 | 6871825 | 6875383 |
| *O. alta* | g822 | Chr1CC | 8032437 | 8034629 |
| *O. alta* | g17751 | Chr3CC | 8208246 | 8210937 |
| *O. sativa* | Os03g0230300 | Chr3 | 6895178 | 6898355 |
| *O. alta* | g824 | Chr1CC | 8081896 | 8083298 |
| *O. alta* | g17754 | Chr3CC | 8242042 | 8243435 |
| *O. sativa* | Os03g0231150 | Chr3 | 6933005 | 6933836 |
| *O. alta* | g825 | Chr1CC | 8115078 | 8117204 |
| *O. alta* | g17755 | Chr3CC | 8259065 | 8261205 |
| *O. sativa* | Os03g0231600 | Chr3 | 6948035 | 6950951 |
| *O. alta* | g827 | Chr1CC | 8123473 | 8128309 |
| *O. alta* | g17758 | Chr3CC | 8268001 | 8272666 |
| *O. sativa* | Os03g0231800 | Chr3 | 6957671 | 6962492 |
| *O. alta* | g829 | Chr1CC | 8156095 | 8165576 |
| *O. alta* | g17760 | Chr3CC | 8292458 | 8301252 |
| *O. sativa* | Os03g0231950 | Chr3 | 6980640 | 6982645 |
| *O. alta* | g831 | Chr1CC | 8173346 | 8176993 |
| *O. alta* | g17761 | Chr3CC | 8304367 | 8308198 |
| *O. sativa* | Os03g0232200 | Chr3 | 6992879 | 6997181 |
| *O. alta* | g832 | Chr1CC | 8215554 | 8215986 |

| | | | | |
|---|---|---|---|---|
| *O. alta* | g17762 | Chr3CC | 8336127 | 8336811 |
| *O. sativa* | Os03g0232400 | Chr3 | 7018347 | 7019179 |
| *O. alta* | g833 | Chr1CC | 8218420 | 8221268 |
| *O. alta* | g17763 | Chr3CC | 8339999 | 8343405 |
| *O. sativa* | Os03g0232500 | Chr3 | 7022844 | 7026172 |
| *O. alta* | g834 | Chr1CC | 8225084 | 8226463 |
| *O. alta* | g17764 | Chr3CC | 8348015 | 8349394 |
| *O. sativa* | Os03g0232600 | Chr3 | 7029174 | 7031611 |
| *O. alta* | g836 | Chr1CC | 8262183 | 8263589 |
| *O. alta* | g17765 | Chr3CC | 8366574 | 8368024 |
| *O. sativa* | Os03g0232800 | Chr3 | 7043271 | 7046128 |
| *O. alta* | g839 | Chr1CC | 8304501 | 8305686 |
| *O. alta* | g17767 | Chr3CC | 8382989 | 8384168 |
| *O. alta* | g840 | Chr1CC | 8306461 | 8307335 |
| *O. alta* | g17768 | Chr3CC | 8385599 | 8386481 |
| *O. sativa* | Os03g0233200 | Chr3 | 7066551 | 7067714 |
| *O. alta* | g841 | Chr1CC | 8313945 | 8315813 |
| *O. alta* | g17769 | Chr3CC | 8390799 | 8392673 |
| *O. sativa* | Os03g0233300 | Chr3 | 7067985 | 7070574 |
| *O. alta* | g842 | Chr1CC | 8321095 | 8324657 |
| *O. alta* | g17770 | Chr3CC | 8410375 | 8413941 |
| *O. sativa* | Os03g0233500 | Chr3 | 7074684 | 7078859 |
| *O. alta* | g843 | Chr1CC | 8328905 | 8332210 |
| *O. alta* | g17771 | Chr3CC | 8418248 | 8421537 |
| *O. sativa* | Os03g0233800 | Chr3 | 7088357 | 7092494 |
| *O. alta* | g844 | Chr1CC | 8332887 | 8333702 |
| *O. alta* | g17772 | Chr3CC | 8422188 | 8423021 |
| *O. sativa* | Os03g0233900 | Chr3 | 7092864 | 7093988 |
| *O. alta* | g847 | Chr1CC | 8348406 | 8350688 |
| *O. alta* | g17776 | Chr3CC | 8438500 | 8441220 |
| *O. alta* | g846 | Chr1CC | 8345148 | 8345918 |
| *O. alta* | g17773 | Chr3CC | 8426184 | 8427073 |
| *O. sativa* | Os03g0234000 | Chr3 | 7096670 | 7097510 |
| *O. alta* | g851 | Chr1CC | 8376407 | 8380271 |
| *O. alta* | g17778 | Chr3CC | 8447054 | 8450918 |

| | | | | |
|---|---|---|---|---|
| *O. sativa* | Os03g0234600 | Chr3 | 7111305 | 7115133 |
| *O. alta* | g852 | Chr1CC | 8386149 | 8387902 |
| *O. alta* | g17779 | Chr3CC | 8456963 | 8458906 |
| *O. sativa* | Os03g0234900 | Chr3 | 7124925 | 7127129 |
| *O. alta* | g853 | Chr1CC | 8390737 | 8392493 |
| *O. alta* | g17780 | Chr3CC | 8460665 | 8462456 |
| *O. sativa* | Os03g0235000 | Chr3 | 7128273 | 7129609 |
| *O. alta* | g854 | Chr1CC | 8397112 | 8400177 |
| *O. alta* | g17781 | Chr3CC | 8473801 | 8476783 |
| *O. sativa* | Os03g0235100 | Chr3 | 7132567 | 7135981 |
| *O. alta* | g855 | Chr1CC | 8400656 | 8402579 |
| *O. alta* | g17782 | Chr3CC | 8477548 | 8479470 |
| *O. sativa* | Os03g0235200 | Chr3 | 7136651 | 7138895 |
| *O. alta* | g856 | Chr1CC | 8403236 | 8430415 |
| *O. alta* | g17784 | Chr3CC | 8484859 | 8488816 |
| *O. sativa* | Os03g0235700 | Chr3 | 7147366 | 7151263 |
| *O. alta* | g858 | Chr1CC | 8450508 | 8452907 |
| *O. alta* | g17787 | Chr3CC | 8511821 | 8514185 |
| *O. sativa* | Os03g0236200 | Chr3 | 7181080 | 7184036 |
| *O. alta* | g859 | Chr1CC | 8453922 | 8454641 |
| *O. alta* | g17788 | Chr3CC | 8514917 | 8515651 |
| *O. sativa* | Os03g0236300 | Chr3 | 7184393 | 7189221 |
| *O. alta* | g861 | Chr1CC | 8477829 | 8479778 |
| *O. alta* | g17791 | Chr3CC | 8530000 | 8538586 |
| *O. alta* | g862 | Chr1CC | 8489297 | 8493462 |
| *O. alta* | g17792 | Chr3CC | 8541844 | 8545641 |
| *O. sativa* | Os03g0237000 | Chr3 | 7223777 | 7228021 |
| *O. alta* | g863 | Chr1CC | 8496408 | 8499016 |
| *O. alta* | g17793 | Chr3CC | 8547181 | 8549401 |
| *O. sativa* | Os03g0237100 | Chr3 | 7229093 | 7231670 |
| *O. alta* | g864 | Chr1CC | 8516975 | 8521296 |
| *O. alta* | g17794 | Chr3CC | 8560505 | 8564607 |
| *O. sativa* | Os03g0237250 | Chr3 | 7237162 | 7241445 |
| *O. alta* | g866 | Chr1CC | 8555429 | 8555971 |
| *O. alta* | g17795 | Chr3CC | 8596003 | 8596545 |

| | | | | |
|---|---|---|---|---|
| *O. sativa* | Os03g0237500 | Chr3 | 7274601 | 7277422 |
| *O. alta* | g867 | Chr1CC | 8559063 | 8564613 |
| *O. alta* | g17796 | Chr3CC | 8599611 | 8604712 |
| *O. sativa* | Os03g0237600 | Chr3 | 7277308 | 7284276 |
| *O. alta* | g868 | Chr1CC | 8571121 | 8573918 |
| *O. alta* | g17798 | Chr3CC | 8614013 | 8616194 |
| *O. sativa* | Os03g0237900 | Chr3 | 7290796 | 7293226 |
| *O. alta* | g869 | Chr1CC | 8577051 | 8577836 |
| *O. alta* | g17799 | Chr3CC | 8616859 | 8617638 |
| *O. sativa* | Os03g0237950 | Chr3 | 7293721 | 7293988 |
| *O. alta* | g870 | Chr1CC | 8581367 | 8584466 |
| *O. alta* | g17801 | Chr3CC | 8623404 | 8624592 |
| *O. sativa* | Os03g0238300 | Chr3 | 7305284 | 7308499 |
| *O. alta* | g872 | Chr1CC | 8610051 | 8612917 |
| *O. alta* | g17802 | Chr3CC | 8654346 | 8656522 |
| *O. sativa* | Os03g0238600 | Chr3 | 7322529 | 7325182 |
| *O. alta* | g873 | Chr1CC | 8613753 | 8616857 |
| *O. alta* | g17803 | Chr3CC | 8657321 | 8667109 |
| *O. sativa* | Os03g0238700 | Chr3 | 7324024 | 7328856 |
| *O. alta* | g875 | Chr1CC | 8626157 | 8630382 |
| *O. alta* | g17804 | Chr3CC | 8672090 | 8675967 |
| *O. sativa* | Os03g0239000 | Chr3 | 7339645 | 7342730 |
| *O. alta* | g876 | Chr1CC | 8647615 | 8651401 |
| *O. alta* | g17805 | Chr3CC | 8680564 | 8684368 |
| *O. sativa* | Os03g0239200 | Chr3 | 7347523 | 7351722 |
| *O. alta* | g883 | Chr1CC | 8689276 | 8689998 |
| *O. alta* | g17806 | Chr3CC | 8686024 | 8686737 |
| *O. sativa* | Os03g0239300 | Chr3 | 7353076 | 7354368 |
| *O. alta* | g884 | Chr1CC | 8696036 | 8704330 |
| *O. alta* | g17807 | Chr3CC | 8692031 | 8697136 |
| *O. sativa* | Os03g0239400 | Chr3 | 7358092 | 7369614 |
| *O. alta* | g886 | Chr1CC | 8710973 | 8714176 |
| *O. alta* | g17809 | Chr3CC | 8713170 | 8715970 |
| *O. sativa* | Os03g0240500 | Chr3 | 7434060 | 7438082 |
| *O. alta* | g888 | Chr1CC | 8718028 | 8719359 |

| | | | | |
|---|---|---|---|---|
| *O. alta* | g17810 | Chr3CC | 8718037 | 8719365 |
| *O. sativa* | Os03g0240600 | Chr3 | 7440402 | 7442007 |
| *O. alta* | g889 | Chr1CC | 8724125 | 8726028 |
| *O. alta* | g17811 | Chr3CC | 8724030 | 8726116 |
| *O. sativa* | Os03g0240700 | Chr3 | 7446000 | 7449581 |
| *O. alta* | g890 | Chr1CC | 8726526 | 8727179 |
| *O. alta* | g17812 | Chr3CC | 8740072 | 8742788 |
| *O. sativa* | Os03g0240800 | Chr3 | 7449797 | 7453038 |
| *O. alta* | g891 | Chr1CC | 8735899 | 8742382 |
| *O. alta* | g17815 | Chr3CC | 8748775 | 8760133 |
| *O. sativa* | Os03g0241100 | Chr3 | 7469617 | 7475442 |
| *O. alta* | g892 | Chr1CC | 8744360 | 8746818 |
| *O. alta* | g17817 | Chr3CC | 8764890 | 8767301 |
| *O. sativa* | Os03g0241300 | Chr3 | 7480379 | 7483234 |
| *O. alta* | g893 | Chr1CC | 8747349 | 8750558 |
| *O. alta* | g17818 | Chr3CC | 8767858 | 8771021 |
| *O. sativa* | Os03g0241600 | Chr3 | 7483344 | 7487296 |
| *O. alta* | g894 | Chr1CC | 8759087 | 8762395 |
| *O. alta* | g17819 | Chr3CC | 8778544 | 8782544 |
| *O. sativa* | Os03g0241800 | Chr3 | 7497368 | 7499362 |
| *O. alta* | g897 | Chr1CC | 8774995 | 8778465 |
| *O. alta* | g17820 | Chr3CC | 8785165 | 8788570 |
| *O. sativa* | Os03g0241900 | Chr3 | 7500164 | 7503768 |
| *O. alta* | g898 | Chr1CC | 8781547 | 8784874 |
| *O. alta* | g17821 | Chr3CC | 8794740 | 8801457 |
| *O. alta* | g899 | Chr1CC | 8792930 | 8796733 |
| *O. alta* | g17822 | Chr3CC | 8802920 | 8806436 |
| *O. sativa* | Os03g0242200 | Chr3 | 7514225 | 7517543 |
| *O. alta* | g900 | Chr1CC | 8800163 | 8800858 |
| *O. alta* | g17823 | Chr3CC | 8809445 | 8810137 |
| *O. sativa* | Os03g0242300 | Chr3 | 7519754 | 7521478 |
| *O. alta* | g902 | Chr1CC | 8829197 | 8831545 |
| *O. alta* | g17824 | Chr3CC | 8822279 | 8824624 |
| *O. sativa* | Os03g0242900 | Chr3 | 7559117 | 7561984 |
| *O. alta* | g903 | Chr1CC | 8866179 | 8868243 |

| | | | | |
|---|---|---|---|---|
| *O. alta* | g17825 | Chr3CC | 8826982 | 8828849 |
| *O. sativa* | Os03g0243100 | Chr3 | 7565328 | 7567774 |
| *O. alta* | g904 | Chr1CC | 8874967 | 8877926 |
| *O. alta* | g17826 | Chr3CC | 8837566 | 8842281 |
| *O. sativa* | Os03g0243200 | Chr3 | 7573271 | 7577555 |
| *O. alta* | g906 | Chr1CC | 8882962 | 8886005 |
| *O. alta* | g17827 | Chr3CC | 8846643 | 8849666 |
| *O. sativa* | Os03g0243300 | Chr3 | 7579448 | 7583738 |
| *O. alta* | g907 | Chr1CC | 8910931 | 8916007 |
| *O. alta* | g17828 | Chr3CC | 8907535 | 8912597 |
| *O. sativa* | Os03g0243600 | Chr3 | 7591870 | 7597405 |
| *O. alta* | g908 | Chr1CC | 8923033 | 8926716 |
| *O. alta* | g17829 | Chr3CC | 8916110 | 8918344 |
| *O. sativa* | Os03g0243700 | Chr3 | 7601448 | 7603037 |
| *O. alta* | g909 | Chr1CC | 8927100 | 8935308 |
| *O. alta* | g17830 | Chr3CC | 8918703 | 8922998 |
| *O. sativa* | Os03g0243800 | Chr3 | 7603131 | 7607537 |
| *O. alta* | g910 | Chr1CC | 8938848 | 8940784 |
| *O. alta* | g17831 | Chr3CC | 8927287 | 8929177 |
| *O. sativa* | Os03g0243900 | Chr3 | 7611453 | 7614083 |
| *O. alta* | g911 | Chr1CC | 8942979 | 8945725 |
| *O. alta* | g17832 | Chr3CC | 8933691 | 8935217 |
| *O. sativa* | Os03g0244000 | Chr3 | 7615918 | 7618902 |
| *O. alta* | g913 | Chr1CC | 8992455 | 8994783 |
| *O. alta* | g17833 | Chr3CC | 8943580 | 8945965 |
| *O. sativa* | Os03g0244200 | Chr3 | 7626455 | 7629565 |
| *O. alta* | g916 | Chr1CC | 9011927 | 9015421 |
| *O. alta* | g17834 | Chr3CC | 8959910 | 8963355 |
| *O. sativa* | Os03g0244600 | Chr3 | 7646649 | 7650520 |
| *O. alta* | g917 | Chr1CC | 9025699 | 9027636 |
| *O. alta* | g17836 | Chr3CC | 8979441 | 8981375 |
| *O. sativa* | Os03g0244700 | Chr3 | 7656198 | 7658476 |
| *O. alta* | g918 | Chr1CC | 9047369 | 9047860 |
| *O. alta* | g17837 | Chr3CC | 8992232 | 8992693 |
| *O. sativa* | Os03g0244950 | Chr3 | 7669855 | 7670370 |

| | | | | |
|---|---|---|---|---|
| *O. alta* | g919 | Chr1CC | 9050567 | 9053751 |
| *O. alta* | g17838 | Chr3CC | 8993970 | 8995746 |
| *O. sativa* | Os03g0245100 | Chr3 | 7671601 | 7676854 |
| *O. alta* | g920 | Chr1CC | 9054639 | 9055684 |
| *O. alta* | g17840 | Chr3CC | 8998059 | 8999131 |
| *O. sativa* | Os03g0245200 | Chr3 | 7675772 | 7677111 |
| *O. alta* | g921 | Chr1CC | 9056430 | 9057582 |
| *O. alta* | g17841 | Chr3CC | 9000050 | 9001245 |
| *O. sativa* | Os03g0245300 | Chr3 | 7677693 | 7679130 |
| *O. alta* | g925 | Chr1CC | 9062057 | 9062947 |
| *O. alta* | g17842 | Chr3CC | 9002959 | 9003903 |
| *O. sativa* | Os03g0245500 | Chr3 | 7681570 | 7682838 |
| *O. alta* | g926 | Chr1CC | 9066027 | 9067625 |
| *O. alta* | g17843 | Chr3CC | 9006877 | 9008475 |
| *O. sativa* | Os03g0245700 | Chr3 | 7686713 | 7688842 |
| *O. alta* | g927 | Chr1CC | 9073398 | 9074120 |
| *O. alta* | g17844 | Chr3CC | 9012943 | 9013665 |
| *O. sativa* | Os03g0245800 | Chr3 | 7697015 | 7698027 |
| *O. alta* | g928 | Chr1CC | 9075132 | 9075782 |
| *O. alta* | g17845 | Chr3CC | 9014911 | 9016049 |
| *O. alta* | g929 | Chr1CC | 9084112 | 9085426 |
| *O. alta* | g17846 | Chr3CC | 9019569 | 9020873 |
| *O. sativa* | Os03g0246100 | Chr3 | 7707769 | 7710526 |
| *O. alta* | g932 | Chr1CC | 9128216 | 9132888 |
| *O. alta* | g17847 | Chr3CC | 9055487 | 9060239 |
| *O. sativa* | Os03g0246500 | Chr3 | 7740991 | 7749358 |
| *O. alta* | g933 | Chr1CC | 9135598 | 9144378 |
| *O. alta* | g17848 | Chr3CC | 9063880 | 9072689 |
| *O. sativa* | Os03g0246800 | Chr3 | 7753164 | 7763728 |
| *O. alta* | g934 | Chr1CC | 9147602 | 9148051 |
| *O. alta* | g17849 | Chr3CC | 9077061 | 9078654 |
| *O. sativa* | Os03g0246900 | Chr3 | 7767062 | 7768701 |
| *O. alta* | g936 | Chr1CC | 9158509 | 9163879 |
| *O. alta* | g17850 | Chr3CC | 9085449 | 9091301 |
| *O. sativa* | Os03g0247000 | Chr3 | 7773866 | 7779865 |

| | | | | |
|---|---|---|---|---|
| *O. alta* | g937 | Chr1CC | 9174614 | 9176563 |
| *O. alta* | g17851 | Chr3CC | 9097049 | 9098998 |
| *O. sativa* | Os03g0247100 | Chr3 | 7782444 | 7784795 |
| *O. alta* | g938 | Chr1CC | 9181299 | 9181727 |
| *O. alta* | g17852 | Chr3CC | 9103441 | 9103848 |
| *O. sativa* | Os03g0247200 | Chr3 | 7788798 | 7789593 |
| *O. alta* | g941 | Chr1CC | 9206783 | 9207046 |
| *O. alta* | g17858 | Chr3CC | 9163760 | 9164005 |
| *O. sativa* | Os03g0247750 | Chr3 | 7802434 | 7802697 |
| *O. alta* | g942 | Chr1CC | 9217875 | 9219969 |
| *O. alta* | g17859 | Chr3CC | 9174181 | 9176298 |
| *O. sativa* | Os03g0247900 | Chr3 | 7815147 | 7817732 |
| *O. alta* | g943 | Chr1CC | 9223400 | 9229806 |
| *O. alta* | g17860 | Chr3CC | 9179047 | 9185664 |
| *O. sativa* | Os03g0248000 | Chr3 | 7819901 | 7827315 |
| *O. alta* | g945 | Chr1CC | 9240870 | 9242652 |
| *O. alta* | g17862 | Chr3CC | 9191730 | 9193498 |
| *O. alta* | g946 | Chr1CC | 9285865 | 9287306 |
| *O. alta* | g17865 | Chr3CC | 9233668 | 9235110 |
| *O. sativa* | Os03g0249100 | Chr3 | 7871494 | 7873366 |
| *O. alta* | g947 | Chr1CC | 9287885 | 9290528 |
| *O. alta* | g17866 | Chr3CC | 9235651 | 9238572 |
| *O. sativa* | Os03g0249200 | Chr3 | 7874537 | 7878425 |
| *O. alta* | g948 | Chr1CC | 9293830 | 9296223 |
| *O. alta* | g17867 | Chr3CC | 9252629 | 9255074 |
| *O. sativa* | Os03g0249300 | Chr3 | 7881242 | 7884214 |
| *O. alta* | g949 | Chr1CC | 9297298 | 9298807 |
| *O. alta* | g17868 | Chr3CC | 9256107 | 9257611 |
| *O. sativa* | Os03g0249400 | Chr3 | 7884815 | 7886786 |
| *O. alta* | g950 | Chr1CC | 9301453 | 9302904 |
| *O. alta* | g17869 | Chr3CC | 9259976 | 9261427 |
| *O. sativa* | Os03g0249500 | Chr3 | 7889338 | 7891292 |
| *O. alta* | g952 | Chr1CC | 9309519 | 9310085 |
| *O. alta* | g17871 | Chr3CC | 9277571 | 9278503 |
| *O. sativa* | Os03g0248300 | Chr3 | 7840812 | 7841454 |

| | | | | |
|---|---|---|---|---|
| *O. alta* | g954 | Chr1CC | 9347169 | 9347561 |
| *O. alta* | g17872 | Chr3CC | 9284222 | 9284614 |
| *O. sativa* | Os03g0249700 | Chr3 | 7907082 | 7907812 |
| *O. alta* | g955 | Chr1CC | 9349101 | 9351427 |
| *O. alta* | g17873 | Chr3CC | 9294878 | 9297222 |
| *O. sativa* | Os03g0249900 | Chr3 | 7911693 | 7914268 |
| *O. alta* | g17874 | Chr3CC | 9298055 | 9300890 |
| *O. alta* | g956 | Chr1CC | 9355209 | 9358471 |
| *O. sativa* | Os03g0250000 | Chr3 | 7914611 | 7918255 |
| *O. alta* | g17875 | Chr3CC | 9302825 | 9307704 |
| *O. alta* | g957 | Chr1CC | 9361572 | 9365076 |
| *O. sativa* | Os03g0250200 | Chr3 | 7929012 | 7933985 |
| *O. alta* | g17879 | Chr3CC | 9399065 | 9399808 |
| *O. alta* | g959 | Chr1CC | 9387096 | 9387839 |
| *O. alta* | g17880 | Chr3CC | 9402780 | 9408945 |
| *O. alta* | g960 | Chr1CC | 9391065 | 9397500 |
| *O. sativa* | Os03g0251500 | Chr3 | 7982396 | 7989433 |
| *O. alta* | g17883 | Chr3CC | 9423551 | 9426991 |
| *O. alta* | g963 | Chr1CC | 9414012 | 9416744 |
| *O. sativa* | Os03g0251800 | Chr3 | 8003074 | 8008237 |
| *O. alta* | g17884 | Chr3CC | 9432948 | 9433937 |
| *O. alta* | g964 | Chr1CC | 9425998 | 9426981 |
| *O. sativa* | Os03g0252100 | Chr3 | 8010885 | 8012566 |
| *O. alta* | g17886 | Chr3CC | 9465767 | 9469960 |
| *O. alta* | g966 | Chr1CC | 9467730 | 9471671 |
| *O. sativa* | Os03g0252800 | Chr3 | 8047380 | 8052536 |
| *O. alta* | g17887 | Chr3CC | 9492686 | 9493071 |
| *O. alta* | g967 | Chr1CC | 9481831 | 9482217 |
| *O. sativa* | Os03g0252900 | Chr3 | 8061598 | 8061988 |
| *O. alta* | g17888 | Chr3CC | 9497176 | 9502312 |
| *O. alta* | g968 | Chr1CC | 9486318 | 9490193 |
| *O. sativa* | Os03g0253100 | Chr3 | 8067630 | 8071826 |
| *O. alta* | g17889 | Chr3CC | 9516994 | 9518520 |
| *O. alta* | g969 | Chr1CC | 9504766 | 9506295 |
| *O. sativa* | Os03g0253200 | Chr3 | 8087161 | 8088952 |

| | | | | |
|---|---|---|---|---|
| *O. alta* | g17890 | Chr3CC | 9534968 | 9536272 |
| *O. alta* | g971 | Chr1CC | 9518823 | 9520133 |
| *O. sativa* | Os03g0253500 | Chr3 | 8101572 | 8102858 |
| *O. alta* | g17891 | Chr3CC | 9541373 | 9542371 |
| *O. alta* | g975 | Chr1CC | 9531565 | 9532563 |
| *O. sativa* | Os03g0253600 | Chr3 | 8106340 | 8108381 |
| *O. alta* | g17892 | Chr3CC | 9546873 | 9551940 |
| *O. alta* | g976 | Chr1CC | 9536781 | 9542076 |
| *O. sativa* | Os03g0253800 | Chr3 | 8112267 | 8117742 |
| *O. alta* | g17893 | Chr3CC | 9559252 | 9562687 |
| *O. alta* | g977 | Chr1CC | 9553444 | 9556911 |
| *O. sativa* | Os03g0254000 | Chr3 | 8122290 | 8126074 |
| *O. alta* | g17894 | Chr3CC | 9566226 | 9570835 |
| *O. alta* | g978 | Chr1CC | 9562388 | 9565603 |
| *O. sativa* | Os03g0254200 | Chr3 | 8130378 | 8132030 |
| *O. alta* | g17897 | Chr3CC | 9611982 | 9619210 |
| *O. alta* | g981 | Chr1CC | 9604036 | 9611214 |
| *O. sativa* | Os03g0254700 | Chr3 | 8165827 | 8174895 |
| *O. alta* | g17898 | Chr3CC | 9624675 | 9628107 |
| *O. alta* | g983 | Chr1CC | 9615203 | 9618645 |
| *O. sativa* | Os03g0254800 | Chr3 | 8177756 | 8181722 |
| *O. alta* | g17899 | Chr3CC | 9629166 | 9632695 |
| *O. alta* | g984 | Chr1CC | 9619720 | 9623002 |
| *O. sativa* | Os03g0254900 | Chr3 | 8182249 | 8186746 |
| *O. alta* | g17900 | Chr3CC | 9643810 | 9647037 |
| *O. alta* | g985 | Chr1CC | 9630476 | 9633478 |
| *O. sativa* | Os03g0255000 | Chr3 | 8191896 | 8195080 |
| *O. alta* | g17901 | Chr3CC | 9648330 | 9653008 |
| *O. alta* | g986 | Chr1CC | 9635028 | 9639891 |
| *O. sativa* | Os03g0255100 | Chr3 | 8196070 | 8201999 |
| *O. alta* | g17903 | Chr3CC | 9659201 | 9662195 |
| *O. alta* | g987 | Chr1CC | 9646659 | 9649561 |
| *O. sativa* | Os03g0255200 | Chr3 | 8207583 | 8211913 |
| *O. alta* | g17905 | Chr3CC | 9670296 | 9674887 |
| *O. alta* | g989 | Chr1CC | 9687360 | 9691984 |
| *O. sativa* | Os03g0255500 | Chr3 | 8218898 | 8224127 |

| | | | | |
|---|---|---|---|---|
| *O. alta* | g17906 | Chr3CC | 9689455 | 9690115 |
| *O. alta* | g990 | Chr1CC | 9705030 | 9705242 |
| *O. sativa* | Os03g0255900 | Chr3 | 8241532 | 8242113 |
| *O. alta* | g17907 | Chr3CC | 9696171 | 9696818 |
| *O. alta* | g992 | Chr1CC | 9712914 | 9713579 |
| *O. alta* | g17909 | Chr3CC | 9711406 | 9721146 |
| *O. alta* | g994 | Chr1CC | 9723623 | 9733383 |
| *O. alta* | g17910 | Chr3CC | 9723822 | 9739459 |
| *O. alta* | g995 | Chr1CC | 9736143 | 9739501 |
| *O. sativa* | Os03g0257000 | Chr3 | 8307296 | 8311472 |
| *O. alta* | g17911 | Chr3CC | 9741329 | 9741610 |
| *O. alta* | g997 | Chr1CC | 9743809 | 9744084 |
| *O. alta* | g17914 | Chr3CC | 9752686 | 9754829 |
| *O. alta* | g998 | Chr1CC | 9745736 | 9749493 |
| *O. sativa* | Os03g0257600 | Chr3 | 8329118 | 8331769 |
| *O. alta* | g17915 | Chr3CC | 9758562 | 9760519 |
| *O. alta* | g999 | Chr1CC | 9758732 | 9760662 |
| *O. sativa* | Os03g0257900 | Chr3 | 8337473 | 8342481 |
| *O. alta* | g1001 | Chr1CC | 9767168 | 9768250 |
| *O. alta* | g17918 | Chr3CC | 9776075 | 9777163 |
| *O. sativa* | Os03g0258200 | Chr3 | 8352195 | 8353718 |
| *O. alta* | g1004 | Chr1CC | 9786123 | 9787802 |
| *O. alta* | g17919 | Chr3CC | 9808742 | 9810424 |
| *O. sativa* | Os03g0258900 | Chr3 | 8384206 | 8386087 |
| *O. alta* | g1005 | Chr1CC | 9803053 | 9803533 |
| *O. alta* | g17920 | Chr3CC | 9821059 | 9821526 |
| *O. sativa* | Os03g0259100 | Chr3 | 8396569 | 8397410 |
| *O. alta* | g1006 | Chr1CC | 9805006 | 9811499 |
| *O. alta* | g17921 | Chr3CC | 9822882 | 9829450 |
| *O. sativa* | Os03g0259300 | Chr3 | 8398817 | 8406410 |
| *O. alta* | g1007 | Chr1CC | 9815321 | 9816809 |
| *O. alta* | g17922 | Chr3CC | 9830568 | 9832026 |
| *O. sativa* | Os03g0259400 | Chr3 | 8406809 | 8408296 |
| *O. alta* | g1008 | Chr1CC | 9818730 | 9819222 |

| | | | | |
|---|---|---|---|---|
| *O. alta* | g17923 | Chr3CC | 9832925 | 9833426 |
| *O. alta* | g1009 | Chr1CC | 9828538 | 9831691 |
| *O. alta* | g17924 | Chr3CC | 9838482 | 9841721 |
| *O. sativa* | Os03g0259700 | Chr3 | 8415745 | 8419093 |
| *O. alta* | g1010 | Chr1CC | 9837308 | 9840301 |
| *O. alta* | g17925 | Chr3CC | 9848992 | 9851958 |
| *O. sativa* | Os03g0259900 | Chr3 | 8429310 | 8433531 |
| *O. alta* | g1012 | Chr1CC | 9844189 | 9845319 |
| *O. alta* | g17926 | Chr3CC | 9856751 | 9857020 |
| *O. sativa* | Os03g0260000 | Chr3 | 8434789 | 8437195 |
| *O. alta* | g1013 | Chr1CC | 9846079 | 9850095 |
| *O. alta* | g17928 | Chr3CC | 9858854 | 9863231 |
| *O. sativa* | Os03g0260100 | Chr3 | 8437361 | 8441906 |
| *O. alta* | g1014 | Chr1CC | 9852957 | 9853214 |
| *O. alta* | g17929 | Chr3CC | 9876820 | 9877077 |
| *O. sativa* | Os03g0260432 | Chr3 | 8445623 | 8446519 |
| *O. alta* | g1016 | Chr1CC | 9872122 | 9874604 |
| *O. alta* | g17930 | Chr3CC | 9887150 | 9889636 |
| *O. sativa* | Os03g0260600 | Chr3 | 8458131 | 8461066 |
| *O. alta* | g1017 | Chr1CC | 9884452 | 9884781 |
| *O. alta* | g17931 | Chr3CC | 9906657 | 9907405 |
| *O. sativa* | Os03g0261100 | Chr3 | 8487523 | 8489366 |
| *O. alta* | g1018 | Chr1CC | 9886664 | 9892241 |
| *O. alta* | g17932 | Chr3CC | 9912606 | 9916306 |
| *O. sativa* | Os03g0261500 | Chr3 | 8501364 | 8505645 |
| *O. alta* | g1019 | Chr1CC | 9903971 | 9905152 |
| *O. alta* | g17933 | Chr3CC | 9928735 | 9929907 |
| *O. sativa* | Os03g0261800 | Chr3 | 8531062 | 8533071 |
| *O. alta* | g1020 | Chr1CC | 9968947 | 9977977 |
| *O. alta* | g17935 | Chr3CC | 9943518 | 9952612 |
| *O. sativa* | Os03g0261900 | Chr3 | 8546948 | 8552114 |
| *O. alta* | g1021 | Chr1CC | 9983647 | 9986671 |
| *O. alta* | g17937 | Chr3CC | 9959421 | 9962595 |
| *O. sativa* | Os03g0262000 | Chr3 | 8558993 | 8562806 |
| *O. alta* | g1022 | Chr1CC | 9994733 | 9997665 |

| | | | | |
|---|---|---|---|---|
| *O. alta* | g17938 | Chr3CC | 9967379 | 9970607 |
| *O. sativa* | Os03g0262100 | Chr3 | 8568445 | 8571825 |
| *O. alta* | g1023 | Chr1CC | 10000279 | 10003760 |
| *O. alta* | g17939 | Chr3CC | 9972418 | 9975855 |
| *O. sativa* | Os03g0262200 | Chr3 | 8573083 | 8577492 |
| *O. alta* | g1024 | Chr1CC | 10020872 | 10024738 |
| *O. alta* | g17940 | Chr3CC | 9989860 | 9993475 |
| *O. sativa* | Os03g0262300 | Chr3 | 8581503 | 8596767 |
| *O. alta* | g1025 | Chr1CC | 10029478 | 10032718 |
| *O. alta* | g17941 | Chr3CC | 9994813 | 9997183 |
| *O. sativa* | Os03g0262400 | Chr3 | 8597173 | 8600595 |
| *O. alta* | g1026 | Chr1CC | 10035883 | 10036695 |
| *O. alta* | g17942 | Chr3CC | 10002559 | 10003372 |
| *O. sativa* | Os03g0262700 | Chr3 | 8623321 | 8624555 |
| *O. alta* | g1027 | Chr1CC | 10042823 | 10048321 |
| *O. alta* | g17944 | Chr3CC | 10012087 | 10016780 |
| *O. sativa* | Os03g0262900 | Chr3 | 8629862 | 8635160 |
| *O. alta* | g1028 | Chr1CC | 10050205 | 10050987 |
| *O. alta* | g17945 | Chr3CC | 10018443 | 10019231 |
| *O. alta* | g1029 | Chr1CC | 10079928 | 10081667 |
| *O. alta* | g17947 | Chr3CC | 10044805 | 10046559 |
| *O. sativa* | Os03g0263300 | Chr3 | 8651497 | 8653224 |
| *O. alta* | g1030 | Chr1CC | 10087621 | 10091252 |
| *O. alta* | g17948 | Chr3CC | 10053386 | 10056232 |
| *O. sativa* | Os03g0263400 | Chr3 | 8656444 | 8660017 |
| *O. alta* | g1031 | Chr1CC | 10092069 | 10094638 |
| *O. alta* | g17949 | Chr3CC | 10057362 | 10059968 |
| *O. sativa* | Os03g0263500 | Chr3 | 8660232 | 8663021 |
| *O. alta* | g1033 | Chr1CC | 10104661 | 10106202 |
| *O. alta* | g17950 | Chr3CC | 10061513 | 10063048 |
| *O. sativa* | Os03g0263600 | Chr3 | 8664268 | 8666146 |
| *O. alta* | g1034 | Chr1CC | 10115215 | 10118136 |
| *O. alta* | g17952 | Chr3CC | 10091300 | 10094182 |
| *O. sativa* | Os03g0263800 | Chr3 | 8673598 | 8676510 |

| | | | | |
|---|---|---|---|---|
| *O. alta* | g1035 | Chr1CC | 10120679 | 10123225 |
| *O. alta* | g17953 | Chr3CC | 10096949 | 10099495 |
| *O. sativa* | Os03g0263900 | Chr3 | 8679164 | 8682334 |
| *O. alta* | g1036 | Chr1CC | 10127498 | 10128650 |
| *O. alta* | g17954 | Chr3CC | 10103914 | 10105024 |
| *O. sativa* | Os03g0264000 | Chr3 | 8683771 | 8687476 |
| *O. alta* | g1038 | Chr1CC | 10139491 | 10143653 |
| *O. alta* | g17956 | Chr3CC | 10118566 | 10123170 |
| *O. sativa* | Os03g0264400 | Chr3 | 8702305 | 8707258 |
| *O. alta* | g1039 | Chr1CC | 10147844 | 10149187 |
| *O. alta* | g17957 | Chr3CC | 10135642 | 10137000 |
| *O. sativa* | Os03g0264600 | Chr3 | 8713068 | 8714646 |
| *O. alta* | g1040 | Chr1CC | 10154961 | 10156400 |
| *O. alta* | g17958 | Chr3CC | 10140369 | 10141820 |
| *O. sativa* | Os03g0264700 | Chr3 | 8718148 | 8719815 |
| *O. alta* | g1041 | Chr1CC | 10162006 | 10165309 |
| *O. alta* | g17959 | Chr3CC | 10144836 | 10149927 |
| *O. sativa* | Os03g0264800 | Chr3 | 8720259 | 8728315 |
| *O. alta* | g1042 | Chr1CC | 10172049 | 10175647 |
| *O. alta* | g17960 | Chr3CC | 10156446 | 10159937 |
| *O. sativa* | Os03g0265100 | Chr3 | 8737129 | 8741179 |
| *O. alta* | g1043 | Chr1CC | 10176797 | 10178161 |
| *O. alta* | g17961 | Chr3CC | 10161312 | 10164540 |
| *O. sativa* | Os03g0265200 | Chr3 | 8741914 | 8744127 |
| *O. alta* | g1044 | Chr1CC | 10179563 | 10182068 |
| *O. alta* | g17962 | Chr3CC | 10174024 | 10176582 |
| *O. sativa* | Os03g0265300 | Chr3 | 8749943 | 8752907 |
| *O. alta* | g1045 | Chr1CC | 10182934 | 10184992 |
| *O. alta* | g17963 | Chr3CC | 10177664 | 10179788 |
| *O. sativa* | Os03g0265400 | Chr3 | 8753551 | 8756010 |
| *O. alta* | g1047 | Chr1CC | 10195127 | 10197345 |
| *O. alta* | g17964 | Chr3CC | 10212391 | 10214646 |
| *O. sativa* | Os03g0265500 | Chr3 | 8761804 | 8766894 |
| *O. alta* | g1048 | Chr1CC | 10200116 | 10200615 |
| *O. alta* | g17965 | Chr3CC | 10247673 | 10250468 |
| *O. sativa* | Os03g0265600 | Chr3 | 8767643 | 8770548 |

| | | | | |
|---|---|---|---|---|
| *O. alta* | g1049 | Chr1CC | 10224915 | 10235335 |
| *O. alta* | g17966 | Chr3CC | 10256405 | 10266071 |
| *O. sativa* | Os03g0265700 | Chr3 | 8776183 | 8786052 |
| *O. alta* | g1051 | Chr1CC | 10247540 | 10248376 |
| *O. alta* | g17968 | Chr3CC | 10282721 | 10283887 |
| *O. sativa* | Os03g0265900 | Chr3 | 8789242 | 8790378 |
| *O. alta* | g1050 | Chr1CC | 10237282 | 10237692 |
| *O. alta* | g17967 | Chr3CC | 10281808 | 10282209 |
| *O. sativa* | Os03g0265800 | Chr3 | 8788679 | 8789277 |
| *O. alta* | g1052 | Chr1CC | 10248845 | 10252565 |
| *O. alta* | g17969 | Chr3CC | 10284396 | 10287607 |
| *O. sativa* | Os03g0266000 | Chr3 | 8790747 | 8795463 |
| *O. alta* | g1054 | Chr1CC | 10257088 | 10261299 |
| *O. alta* | g17971 | Chr3CC | 10292441 | 10296367 |
| *O. sativa* | Os03g0266200 | Chr3 | 8800461 | 8803575 |
| *O. alta* | g1055 | Chr1CC | 10263508 | 10263990 |
| *O. alta* | g17972 | Chr3CC | 10297011 | 10297496 |
| *O. sativa* | Os03g0266300 | Chr3 | 8805567 | 8806556 |
| *O. alta* | g1057 | Chr1CC | 10291420 | 10295882 |
| *O. alta* | g17975 | Chr3CC | 10313804 | 10321642 |
| *O. sativa* | Os03g0266700 | Chr3 | 8819434 | 8824333 |
| *O. alta* | g1058 | Chr1CC | 10301212 | 10305722 |
| *O. alta* | g17976 | Chr3CC | 10321779 | 10326611 |
| *O. sativa* | Os03g0266800 | Chr3 | 8827752 | 8833517 |
| *O. alta* | g1060 | Chr1CC | 10307674 | 10308138 |
| *O. alta* | g17978 | Chr3CC | 10331674 | 10332144 |
| *O. sativa* | Os03g0267200 | Chr3 | 8837821 | 8838527 |
| *O. alta* | g1062 | Chr1CC | 10320598 | 10322139 |
| *O. alta* | g17980 | Chr3CC | 10338532 | 10340065 |
| *O. sativa* | Os03g0267300 | Chr3 | 8841311 | 8843028 |
| *O. alta* | g1063 | Chr1CC | 10332710 | 10335424 |
| *O. alta* | g17989 | Chr3CC | 10569619 | 10572303 |
| *O. sativa* | Os03g0267500 | Chr3 | 8851332 | 8854426 |
| *O. alta* | g1064 | Chr1CC | 10336209 | 10338994 |
| *O. alta* | g17990 | Chr3CC | 10573090 | 10574950 |

| | | | | |
|---|---|---|---|---|
| *O. sativa* | Os03g0267600 | Chr3 | 8854646 | 8859005 |
| *O. alta* | g1065 | Chr1CC | 10346665 | 10349084 |
| *O. alta* | g17991 | Chr3CC | 10588273 | 10590705 |
| *O. sativa* | Os03g0267700 | Chr3 | 8865964 | 8869087 |
| *O. alta* | g1066 | Chr1CC | 10350811 | 10355096 |
| *O. alta* | g17992 | Chr3CC | 10592113 | 10596527 |
| *O. sativa* | Os03g0267800 | Chr3 | 8870192 | 8875401 |
| *O. alta* | g1067 | Chr1CC | 10370100 | 10373949 |
| *O. alta* | g17993 | Chr3CC | 10614419 | 10618391 |
| *O. sativa* | Os03g0268000 | Chr3 | 8884203 | 8889781 |
| *O. alta* | g1068 | Chr1CC | 10378361 | 10381355 |
| *O. alta* | g17994 | Chr3CC | 10624605 | 10628507 |
| *O. sativa* | Os03g0268100 | Chr3 | 8892244 | 8895472 |
| *O. alta* | g1069 | Chr1CC | 10383063 | 10388266 |
| *O. alta* | g17995 | Chr3CC | 10628678 | 10633874 |
| *O. sativa* | Os03g0268200 | Chr3 | 8896186 | 8901979 |
| *O. alta* | g1070 | Chr1CC | 10390137 | 10392009 |
| *O. alta* | g17996 | Chr3CC | 10635009 | 10637570 |
| *O. sativa* | Os03g0268300 | Chr3 | 8902743 | 8906318 |
| *O. alta* | g1071 | Chr1CC | 10402901 | 10404686 |
| *O. alta* | g17997 | Chr3CC | 10640080 | 10641856 |
| *O. sativa* | Os03g0268400 | Chr3 | 8909302 | 8912790 |
| *O. alta* | g1072 | Chr1CC | 10410402 | 10410692 |
| *O. alta* | g17998 | Chr3CC | 10644633 | 10647365 |
| *O. alta* | g1073 | Chr1CC | 10413832 | 10415599 |
| *O. alta* | g17999 | Chr3CC | 10671498 | 10673062 |
| *O. alta* | g1075 | Chr1CC | 10427160 | 10429470 |
| *O. alta* | g18000 | Chr3CC | 10702069 | 10704493 |
| *O. sativa* | Os03g0268900 | Chr3 | 8938481 | 8941311 |
| *O. alta* | g1076 | Chr1CC | 10430346 | 10432785 |
| *O. alta* | g18001 | Chr3CC | 10709784 | 10711961 |
| *O. sativa* | Os03g0269000 | Chr3 | 8945289 | 8947912 |
| *O. alta* | g1077 | Chr1CC | 10437904 | 10440239 |
| *O. alta* | g18004 | Chr3CC | 10720015 | 10722630 |
| *O. sativa* | Os03g0269300 | Chr3 | 8964025 | 8966806 |

| | | | | |
|---|---|---|---|---|
| *O. alta* | g1078 | Chr1CC | 10443945 | 10444598 |
| *O. alta* | g18005 | Chr3CC | 10725307 | 10727572 |
| *O. sativa* | Os03g0269601 | Chr3 | 8981683 | 8982582 |
| *O. alta* | g1080 | Chr1CC | 10461402 | 10464948 |
| *O. alta* | g18006 | Chr3CC | 10747675 | 10751009 |
| *O. sativa* | Os03g0269900 | Chr3 | 9011629 | 9016463 |
| *O. alta* | g1081 | Chr1CC | 10470096 | 10470872 |
| *O. alta* | g18007 | Chr3CC | 10759743 | 10760522 |
| *O. sativa* | Os03g0270000 | Chr3 | 9022593 | 9023708 |
| *O. alta* | g1082 | Chr1CC | 10478387 | 10480190 |
| *O. alta* | g18008 | Chr3CC | 10767502 | 10769403 |
| *O. sativa* | Os03g0270200 | Chr3 | 9028771 | 9035291 |
| *O. alta* | g1086 | Chr1CC | 10505157 | 10506575 |
| *O. alta* | g18012 | Chr3CC | 10810886 | 10812304 |
| *O. sativa* | Os03g0270500 | Chr3 | 9039598 | 9040733 |
| *O. alta* | g1083 | Chr1CC | 10480543 | 10480974 |
| *O. alta* | g18009 | Chr3CC | 10769677 | 10770108 |
| *O. sativa* | Os03g0270300 | Chr3 | 9031969 | 9032400 |
| *O. alta* | g1088 | Chr1CC | 10513791 | 10514234 |
| *O. alta* | g18014 | Chr3CC | 10818995 | 10819865 |
| *O. sativa* | Os03g0270800 | Chr3 | 9051853 | 9052532 |
| *O. alta* | g1090 | Chr1CC | 10534807 | 10537453 |
| *O. alta* | g18015 | Chr3CC | 10821855 | 10824563 |
| *O. sativa* | Os03g0271100 | Chr3 | 9064856 | 9068046 |
| *O. alta* | g1091 | Chr1CC | 10538103 | 10542301 |
| *O. alta* | g18016 | Chr3CC | 10825233 | 10829842 |
| *O. sativa* | Os03g0271200 | Chr3 | 9068392 | 9073951 |
| *O. alta* | g1092 | Chr1CC | 10543040 | 10543993 |
| *O. alta* | g18017 | Chr3CC | 10830542 | 10831951 |
| *O. sativa* | Os03g0271300 | Chr3 | 9074313 | 9076182 |
| *O. alta* | g1094 | Chr1CC | 10544950 | 10551705 |
| *O. alta* | g18018 | Chr3CC | 10832761 | 10835210 |
| *O. sativa* | Os03g0271400 | Chr3 | 9076047 | 9079383 |
| *O. alta* | g1095 | Chr1CC | 10552183 | 10552769 |
| *O. alta* | g18019 | Chr3CC | 10835659 | 10836219 |

| | | | | |
|---|---|---|---|---|
| *O. sativa* | Os03g0271500 | Chr3 | 9079602 | 9081544 |
| *O. alta* | g1097 | Chr1CC | 10566780 | 10567616 |
| *O. alta* | g18020 | Chr3CC | 10843631 | 10844470 |
| *O. sativa* | Os03g0271600 | Chr3 | 9084480 | 9085918 |
| *O. alta* | g1098 | Chr1CC | 10573843 | 10575207 |
| *O. alta* | g18021 | Chr3CC | 10855201 | 10856559 |
| *O. sativa* | Os03g0271900 | Chr3 | 9096403 | 9098430 |
| *O. alta* | g1099 | Chr1CC | 10576148 | 10578995 |
| *O. alta* | g18022 | Chr3CC | 10858241 | 10861084 |
| *O. sativa* | Os03g0272300 | Chr3 | 9132796 | 9136698 |
| *O. alta* | g1100 | Chr1CC | 10581375 | 10585814 |
| *O. alta* | g18023 | Chr3CC | 10865322 | 10868945 |
| *O. sativa* | Os03g0272400 | Chr3 | 9138612 | 9143528 |
| *O. alta* | g1104 | Chr1CC | 10645985 | 10646275 |
| *O. alta* | g18024 | Chr3CC | 10879780 | 10880073 |
| *O. sativa* | Os03g0272900 | Chr3 | 9157102 | 9157955 |
| *O. alta* | g1105 | Chr1CC | 10659381 | 10662608 |
| *O. alta* | g18025 | Chr3CC | 10893945 | 10895985 |
| *O. sativa* | Os03g0273200 | Chr3 | 9167619 | 9170326 |
| *O. alta* | g1107 | Chr1CC | 10691520 | 10693262 |
| *O. alta* | g18026 | Chr3CC | 10934981 | 10936731 |
| *O. sativa* | Os03g0273800 | Chr3 | 9202541 | 9204642 |
| *O. alta* | g1108 | Chr1CC | 10708668 | 10713227 |
| *O. alta* | g18027 | Chr3CC | 10958569 | 10967490 |
| *O. sativa* | Os03g0274000 | Chr3 | 9218819 | 9224101 |
| *O. alta* | g1109 | Chr1CC | 10731281 | 10734719 |
| *O. alta* | g18028 | Chr3CC | 10989401 | 10992847 |
| *O. sativa* | Os03g0274300 | Chr3 | 9247336 | 9252590 |
| *O. alta* | g1110 | Chr1CC | 10744433 | 10747032 |
| *O. alta* | g18029 | Chr3CC | 11002179 | 11005043 |
| *O. sativa* | Os03g0274800 | Chr3 | 9270240 | 9273393 |
| *O. alta* | g1111 | Chr1CC | 10758472 | 10761362 |
| *O. alta* | g18030 | Chr3CC | 11026098 | 11029000 |
| *O. sativa* | Os03g0275100 | Chr3 | 9285328 | 9288857 |
| *O. alta* | g1112 | Chr1CC | 10771397 | 10774033 |

| | | | | |
|---|---|---|---|---|
| *O. alta* | g18031 | Chr3CC | 11044907 | 11047631 |
| *O. sativa* | Os03g0275300 | Chr3 | 9296962 | 9300002 |
| *O. alta* | g1113 | Chr1CC | 10781126 | 10784971 |
| *O. alta* | g18032 | Chr3CC | 11054914 | 11058680 |
| *O. sativa* | Os03g0275400 | Chr3 | 9307510 | 9312238 |
| *O. alta* | g1114 | Chr1CC | 10786993 | 10788945 |
| *O. alta* | g18033 | Chr3CC | 11061160 | 11063115 |
| *O. sativa* | Os03g0275500 | Chr3 | 9313767 | 9316626 |
| *O. alta* | g1115 | Chr1CC | 10804424 | 10809384 |
| *O. alta* | g18035 | Chr3CC | 11124239 | 11129665 |
| *O. sativa* | Os03g0275900 | Chr3 | 9329407 | 9335088 |
| *O. alta* | g1117 | Chr1CC | 10831094 | 10838227 |
| *O. alta* | g18037 | Chr3CC | 11186875 | 11188534 |
| *O. sativa* | Os03g0276300 | Chr3 | 9359575 | 9359994 |
| *O. alta* | g1119 | Chr1CC | 10861045 | 10863581 |
| *O. alta* | g18038 | Chr3CC | 11214401 | 11217073 |
| *O. sativa* | Os03g0276500 | Chr3 | 9370160 | 9373067 |
| *O. alta* | g1120 | Chr1CC | 10864391 | 10866484 |
| *O. alta* | g18039 | Chr3CC | 11217864 | 11219726 |
| *O. sativa* | Os03g0276600 | Chr3 | 9373380 | 9375758 |
| *O. alta* | g1121 | Chr1CC | 10868312 | 10868930 |
| *O. alta* | g18040 | Chr3CC | 11221579 | 11221922 |
| *O. sativa* | Os03g0276700 | Chr3 | 9375984 | 9377688 |
| *O. alta* | g1123 | Chr1CC | 10871636 | 10875482 |
| *O. alta* | g18043 | Chr3CC | 11287841 | 11291473 |
| *O. sativa* | Os03g0276900 | Chr3 | 9383048 | 9387390 |
| *O. alta* | g1124 | Chr1CC | 10883729 | 10887162 |
| *O. alta* | g18044 | Chr3CC | 11295307 | 11298506 |
| *O. sativa* | Os03g0277000 | Chr3 | 9392349 | 9395747 |
| *O. alta* | g1125 | Chr1CC | 10890212 | 10891825 |
| *O. alta* | g18045 | Chr3CC | 11301456 | 11303075 |
| *O. sativa* | Os03g0277100 | Chr3 | 9404128 | 9405129 |
| *O. alta* | g1127 | Chr1CC | 10918670 | 10920990 |
| *O. alta* | g18046 | Chr3CC | 11315650 | 11317934 |
| *O. sativa* | Os03g0277300 | Chr3 | 9411494 | 9416082 |

| | | | | |
|---|---|---|---|---|
| *O. alta* | g1128 | Chr1CC | 10924507 | 10925008 |
| *O. alta* | g18047 | Chr3CC | 11318690 | 11319565 |
| *O. sativa* | Os03g0277500 | Chr3 | 9418945 | 9419670 |
| *O. alta* | g1129 | Chr1CC | 10928112 | 10929003 |
| *O. alta* | g18048 | Chr3CC | 11323658 | 11324554 |
| *O. sativa* | Os03g0277600 | Chr3 | 9421699 | 9422925 |
| *O. alta* | g1131 | Chr1CC | 10961703 | 10964823 |
| *O. alta* | g18050 | Chr3CC | 11346309 | 11349421 |
| *O. sativa* | Os03g0278000 | Chr3 | 9436483 | 9440576 |
| *O. alta* | g1132 | Chr1CC | 10969485 | 10971609 |
| *O. alta* | g18051 | Chr3CC | 11353981 | 11356103 |
| *O. sativa* | Os03g0278200 | Chr3 | 9445434 | 9448696 |
| *O. alta* | g1133 | Chr1CC | 10974852 | 10976867 |
| *O. alta* | g18052 | Chr3CC | 11358729 | 11360714 |
| *O. sativa* | Os03g0278300 | Chr3 | 9449358 | 9453031 |
| *O. alta* | g1134 | Chr1CC | 10982056 | 10985009 |
| *O. alta* | g18054 | Chr3CC | 11370995 | 11373784 |
| *O. sativa* | Os03g0278400 | Chr3 | 9455580 | 9458843 |
| *O. alta* | g1135 | Chr1CC | 10985675 | 10989793 |
| *O. alta* | g18055 | Chr3CC | 11374479 | 11378451 |
| *O. sativa* | Os03g0278500 | Chr3 | 9461167 | 9465395 |
| *O. alta* | g1136 | Chr1CC | 11002718 | 11011278 |
| *O. alta* | g18057 | Chr3CC | 11384094 | 11387444 |
| *O. sativa* | Os03g0278700 | Chr3 | 9480447 | 9488909 |
| *O. alta* | g1137 | Chr1CC | 11012322 | 11013424 |
| *O. alta* | g18058 | Chr3CC | 11393460 | 11395249 |
| *O. sativa* | Os03g0278800 | Chr3 | 9489475 | 9490784 |
| *O. alta* | g1138 | Chr1CC | 11014114 | 11014743 |
| *O. alta* | g18059 | Chr3CC | 11403157 | 11403801 |
| *O. sativa* | Os03g0278900 | Chr3 | 9491526 | 9492494 |
| *O. alta* | g1139 | Chr1CC | 11015341 | 11015799 |
| *O. alta* | g18060 | Chr3CC | 11404301 | 11404747 |
| *O. sativa* | Os03g0279000 | Chr3 | 9492827 | 9493512 |
| *O. alta* | g1140 | Chr1CC | 11025633 | 11026217 |
| *O. alta* | g18061 | Chr3CC | 11408946 | 11409528 |
| *O. sativa* | Os03g0279200 | Chr3 | 9498973 | 9499826 |

| | | | | |
|---|---|---|---|---|
| *O. sativa* | Os01g0502700 | Chr1 | 17420279 | 17421266 |
| *O. alta* | g1141 | Chr1CC | 11034448 | 11037857 |
| *O. alta* | g18062 | Chr3CC | 11415046 | 11418410 |
| *O. sativa* | Os03g0279400 | Chr3 | 9506763 | 9510932 |
| *O. alta* | g1143 | Chr1CC | 11046672 | 11047412 |
| *O. alta* | g18064 | Chr3CC | 11431077 | 11431820 |
| *O. sativa* | Os03g0279600 | Chr3 | 9520046 | 9521110 |
| *O. alta* | g1144 | Chr1CC | 11049452 | 11050012 |
| *O. alta* | g18067 | Chr3CC | 11461241 | 11465850 |
| *O. sativa* | Os03g0279700 | Chr3 | 9524919 | 9525850 |
| *O. alta* | g1145 | Chr1CC | 11062308 | 11070374 |
| *O. alta* | g18068 | Chr3CC | 11485307 | 11493121 |
| *O. sativa* | Os03g0279832 | Chr3 | 9551782 | 9552498 |
| *O. alta* | g1146 | Chr1CC | 11071536 | 11073141 |
| *O. alta* | g18072 | Chr3CC | 11507333 | 11508990 |
| *O. sativa* | Os03g0279900 | Chr3 | 9553187 | 9555112 |
| *O. alta* | g1148 | Chr1CC | 11076039 | 11083678 |
| *O. alta* | g18074 | Chr3CC | 11511958 | 11520290 |
| *O. sativa* | Os03g0280000 | Chr3 | 9557535 | 9566491 |
| *O. alta* | g1150 | Chr1CC | 11108126 | 11108507 |
| *O. alta* | g18076 | Chr3CC | 11533319 | 11534061 |
| *O. alta* | g1153 | Chr1CC | 11135459 | 11136418 |
| *O. alta* | g18080 | Chr3CC | 11561905 | 11562855 |
| *O. sativa* | Os03g0280750 | Chr3 | 9599365 | 9600675 |
| *O. alta* | g1154 | Chr1CC | 11136897 | 11139370 |
| *O. alta* | g18081 | Chr3CC | 11563384 | 11565937 |
| *O. alta* | g1155 | Chr1CC | 11142417 | 11145018 |
| *O. alta* | g18082 | Chr3CC | 11567847 | 11570426 |
| *O. sativa* | Os03g0281000 | Chr3 | 9605184 | 9608335 |
| *O. alta* | g1156 | Chr1CC | 11149875 | 11151662 |
| *O. alta* | g18083 | Chr3CC | 11573612 | 11575416 |
| *O. sativa* | Os03g0281100 | Chr3 | 9611073 | 9612943 |
| *O. alta* | g1158 | Chr1CC | 11168868 | 11169791 |
| *O. alta* | g18084 | Chr3CC | 11582035 | 11582958 |
| *O. sativa* | Os03g0281201 | Chr3 | 9617509 | 9618641 |

| | | | | |
|---|---|---|---|---|
| *O. alta* | g1159 | Chr1CC | 11171459 | 11178026 |
| *O. alta* | g18085 | Chr3CC | 11584240 | 11589836 |
| *O. sativa* | Os03g0281300 | Chr3 | 9619836 | 9628261 |
| *O. alta* | g1160 | Chr1CC | 11178528 | 11180206 |
| *O. alta* | g18086 | Chr3CC | 11590424 | 11590729 |
| *O. alta* | g1161 | Chr1CC | 11190517 | 11193033 |
| *O. alta* | g18087 | Chr3CC | 11598211 | 11600739 |
| *O. sativa* | Os03g0281500 | Chr3 | 9632806 | 9635793 |
| *O. alta* | g1162 | Chr1CC | 11193698 | 11199253 |
| *O. alta* | g18088 | Chr3CC | 11601389 | 11606953 |
| *O. sativa* | Os03g0281600 | Chr3 | 9635957 | 9642093 |
| *O. alta* | g1163 | Chr1CC | 11235508 | 11236476 |
| *O. alta* | g18089 | Chr3CC | 11613036 | 11614016 |
| *O. alta* | g1164 | Chr1CC | 11237142 | 11240461 |
| *O. alta* | g18090 | Chr3CC | 11614478 | 11621949 |
| *O. sativa* | Os03g0281800 | Chr3 | 9646913 | 9651687 |
| *O. alta* | g1166 | Chr1CC | 11243905 | 11246280 |
| *O. alta* | g18091 | Chr3CC | 11627309 | 11629681 |
| *O. sativa* | Os03g0281900 | Chr3 | 9654333 | 9657024 |
| *O. alta* | g1167 | Chr1CC | 11251980 | 11256697 |
| *O. alta* | g18092 | Chr3CC | 11640760 | 11643030 |
| *O. sativa* | Os03g0282100 | Chr3 | 9664896 | 9667255 |
| *O. alta* | g1168 | Chr1CC | 11282527 | 11286228 |
| *O. alta* | g18095 | Chr3CC | 11663683 | 11667385 |
| *O. sativa* | Os03g0282300 | Chr3 | 9683383 | 9684918 |
| *O. alta* | g1170 | Chr1CC | 11306641 | 11309830 |
| *O. alta* | g18097 | Chr3CC | 11679610 | 11683603 |
| *O. sativa* | Os03g0282700 | Chr3 | 9699177 | 9700391 |
| *O. alta* | g1171 | Chr1CC | 11310194 | 11312891 |
| *O. alta* | g18098 | Chr3CC | 11683996 | 11686741 |
| *O. sativa* | Os03g0282800 | Chr3 | 9700521 | 9703425 |
| *O. alta* | g1177 | Chr1CC | 11331141 | 11333006 |
| *O. alta* | g18101 | Chr3CC | 11710554 | 11713182 |
| *O. sativa* | Os03g0283100 | Chr3 | 9712224 | 9714920 |

| | | | | |
|---|---|---|---|---|
| *O. alta* | g1178 | Chr1CC | 11334198 | 11336631 |
| *O. alta* | g18103 | Chr3CC | 11730302 | 11738394 |
| *O. sativa* | Os03g0283300 | Chr3 | 9721925 | 9725003 |
| *O. alta* | g1179 | Chr1CC | 11337291 | 11344478 |
| *O. alta* | g18105 | Chr3CC | 11744939 | 11746735 |
| *O. sativa* | Os03g0283500 | Chr3 | 9730441 | 9732958 |
| *O. alta* | g1180 | Chr1CC | 11345208 | 11349212 |
| *O. alta* | g18106 | Chr3CC | 11747458 | 11752042 |
| *O. sativa* | Os03g0283600 | Chr3 | 9733147 | 9737559 |
| *O. alta* | g1181 | Chr1CC | 11354452 | 11359562 |
| *O. alta* | g18107 | Chr3CC | 11768244 | 11773095 |
| *O. sativa* | Os03g0283800 | Chr3 | 9744220 | 9751248 |
| *O. alta* | g1182 | Chr1CC | 11363549 | 11368527 |
| *O. alta* | g18108 | Chr3CC | 11775116 | 11778431 |
| *O. sativa* | Os03g0283900 | Chr3 | 9751802 | 9755757 |
| *O. alta* | g1183 | Chr1CC | 11368583 | 11370198 |
| *O. alta* | g18109 | Chr3CC | 11779153 | 11781157 |
| *O. sativa* | Os03g0284000 | Chr3 | 9756908 | 9759147 |
| *O. alta* | g1184 | Chr1CC | 11371457 | 11380243 |
| *O. alta* | g18110 | Chr3CC | 11782084 | 11790042 |
| *O. sativa* | Os03g0284100 | Chr3 | 9759666 | 9768689 |
| *O. alta* | g1185 | Chr1CC | 11390650 | 11391318 |
| *O. alta* | g18112 | Chr3CC | 11810550 | 11811203 |
| *O. sativa* | Os03g0284400 | Chr3 | 9781436 | 9782524 |
| *O. alta* | g1187 | Chr1CC | 11397750 | 11401018 |
| *O. alta* | g18114 | Chr3CC | 11818529 | 11825403 |
| *O. sativa* | Os03g0284600 | Chr3 | 9788873 | 9791847 |
| *O. alta* | g1189 | Chr1CC | 11411968 | 11413281 |
| *O. alta* | g18115 | Chr3CC | 11826496 | 11827818 |
| *O. sativa* | Os03g0284800 | Chr3 | 9797238 | 9799120 |
| *O. alta* | g1190 | Chr1CC | 11415276 | 11421686 |
| *O. alta* | g18116 | Chr3CC | 11829037 | 11835845 |
| *O. sativa* | Os03g0284900 | Chr3 | 9799120 | 9822067 |
| *O. alta* | g1191 | Chr1CC | 11422248 | 11425319 |
| *O. alta* | g18117 | Chr3CC | 11836433 | 11838614 |
| *O. sativa* | Os03g0285100 | Chr3 | 9822842 | 9826639 |

| | | | | |
|---|---|---|---|---|
| *O. alta* | g1193 | Chr1CC | 11431126 | 11431335 |
| *O. alta* | g18121 | Chr3CC | 11856729 | 11856995 |
| *O. sativa* | Os03g0285300 | Chr3 | 9832684 | 9833183 |
| *O. alta* | g1195 | Chr1CC | 11441571 | 11445241 |
| *O. alta* | g18120 | Chr3CC | 11846679 | 11850235 |
| *O. sativa* | Os03g0285700 | Chr3 | 9843336 | 9846670 |
| *O. alta* | g1196 | Chr1CC | 11446876 | 11449133 |
| *O. alta* | g18119 | Chr3CC | 11842773 | 11845022 |
| *O. sativa* | Os03g0285800 | Chr3 | 9847723 | 9850384 |
| *O. alta* | g1197 | Chr1CC | 11451785 | 11454020 |
| *O. alta* | g18123 | Chr3CC | 11864745 | 11868656 |
| *O. sativa* | Os03g0285900 | Chr3 | 9852813 | 9856006 |
| *O. alta* | g1198 | Chr1CC | 11470691 | 11473518 |
| *O. alta* | g18124 | Chr3CC | 11870489 | 11873540 |
| *O. sativa* | Os03g0286200 | Chr3 | 9863844 | 9868646 |
| *O. alta* | g1199 | Chr1CC | 11477458 | 11481065 |
| *O. alta* | g18125 | Chr3CC | 11879836 | 11884025 |
| *O. sativa* | Os03g0286300 | Chr3 | 9872301 | 9876576 |
| *O. alta* | g1200 | Chr1CC | 11483778 | 11486059 |
| *O. alta* | g18126 | Chr3CC | 11886598 | 11888479 |
| *O. sativa* | Os03g0286500 | Chr3 | 9878106 | 9880793 |
| *O. alta* | g1201 | Chr1CC | 11507689 | 11510009 |
| *O. alta* | g18128 | Chr3CC | 11902353 | 11905080 |
| *O. sativa* | Os03g0286800 | Chr3 | 9893421 | 9896598 |
| *O. alta* | g1203 | Chr1CC | 11515849 | 11517836 |
| *O. alta* | g18131 | Chr3CC | 11912705 | 11915711 |
| *O. sativa* | Os03g0287100 | Chr3 | 9904631 | 9907008 |
| *O. alta* | g1204 | Chr1CC | 11525521 | 11526314 |
| *O. alta* | g18132 | Chr3CC | 11926460 | 11927252 |
| *O. sativa* | Os03g0287400 | Chr3 | 9916095 | 9917286 |
| *O. alta* | g1205 | Chr1CC | 11553904 | 11557425 |
| *O. alta* | g18133 | Chr3CC | 11957782 | 11961310 |
| *O. sativa* | Os03g0287600 | Chr3 | 9935309 | 9937411 |
| *O. alta* | g1206 | Chr1CC | 11564657 | 11568294 |
| *O. alta* | g18134 | Chr3CC | 11967470 | 11972147 |

| | | | | |
|---|---|---|---|---|
| *O. sativa* | Os03g0287800 | Chr3 | 9946578 | 9949629 |
| *O. alta* | g1207 | Chr1CC | 11578831 | 11579871 |
| *O. alta* | g18135 | Chr3CC | 11973738 | 11976433 |
| *O. sativa* | Os03g0287900 | Chr3 | 9952865 | 9955129 |
| *O. alta* | g1209 | Chr1CC | 11590528 | 11595229 |
| *O. alta* | g18137 | Chr3CC | 11991913 | 11997720 |
| *O. sativa* | Os03g0288300 | Chr3 | 9972495 | 9977763 |
| *O. alta* | g1210 | Chr1CC | 11599460 | 11600042 |
| *O. alta* | g18139 | Chr3CC | 12013554 | 12014135 |
| *O. sativa* | Os03g0288400 | Chr3 | 9978906 | 9980978 |
| *O. alta* | g1211 | Chr1CC | 11600700 | 11603964 |
| *O. alta* | g18140 | Chr3CC | 12014843 | 12016458 |
| *O. sativa* | Os03g0288500 | Chr3 | 9981700 | 9986561 |
| *O. alta* | g1212 | Chr1CC | 11606081 | 11608899 |
| *O. alta* | g18141 | Chr3CC | 12020564 | 12023382 |
| *O. sativa* | Os03g0288600 | Chr3 | 9987257 | 9990388 |
| *O. alta* | g1213 | Chr1CC | 11610090 | 11612711 |
| *O. alta* | g18142 | Chr3CC | 12024588 | 12026457 |
| *O. sativa* | Os03g0288700 | Chr3 | 9990513 | 9993307 |
| *O. alta* | g1214 | Chr1CC | 11620224 | 11623745 |
| *O. alta* | g18143 | Chr3CC | 12028446 | 12031881 |
| *O. sativa* | Os03g0288800 | Chr3 | 9994917 | 9998844 |
| *O. alta* | g1215 | Chr1CC | 11626010 | 11628542 |
| *O. alta* | g18144 | Chr3CC | 12037177 | 12038161 |
| *O. sativa* | Os03g0288900 | Chr3 | 9999638 | 10002736 |
| *O. alta* | g1216 | Chr1CC | 11647327 | 11650849 |
| *O. alta* | g18146 | Chr3CC | 12044853 | 12049235 |
| *O. sativa* | Os03g0289100 | Chr3 | 10006390 | 10012024 |
| *O. alta* | g1217 | Chr1CC | 11656797 | 11657567 |
| *O. alta* | g18147 | Chr3CC | 12062326 | 12063096 |
| *O. sativa* | Os03g0289200 | Chr3 | 10016501 | 10019508 |
| *O. alta* | g1218 | Chr1CC | 11677492 | 11679466 |
| *O. alta* | g18148 | Chr3CC | 12067482 | 12070648 |
| *O. sativa* | Os03g0289300 | Chr3 | 10024083 | 10027655 |
| *O. alta* | g1219 | Chr1CC | 11680274 | 11681018 |

| | | | | |
|---|---|---|---|---|
| *O. alta* | g18149 | Chr3CC | 12071473 | 12072188 |
| *O. sativa* | Os03g0289400 | Chr3 | 10028091 | 10029033 |
| *O. alta* | g1220 | Chr1CC | 11686966 | 11691319 |
| *O. alta* | g18150 | Chr3CC | 12089153 | 12093614 |
| *O. sativa* | Os03g0289800 | Chr3 | 10039387 | 10044161 |
| *O. alta* | g1221 | Chr1CC | 11746418 | 11749326 |
| *O. alta* | g18151 | Chr3CC | 12118448 | 12121385 |
| *O. sativa* | Os03g0290300 | Chr3 | 10080334 | 10083451 |
| *O. alta* | g1222 | Chr1CC | 11756932 | 11767002 |
| *O. alta* | g18152 | Chr3CC | 12126344 | 12135554 |
| *O. sativa* | Os03g0290500 | Chr3 | 10090151 | 10097850 |
| *O. alta* | g1223 | Chr1CC | 11773918 | 11777261 |
| *O. alta* | g18154 | Chr3CC | 12149499 | 12151775 |
| *O. sativa* | Os03g0291200 | Chr3 | 10116219 | 10120372 |
| *O. alta* | g1224 | Chr1CC | 11787608 | 11789265 |
| *O. alta* | g18153 | Chr3CC | 12142064 | 12143715 |
| *O. sativa* | Os03g0290900 | Chr3 | 10105989 | 10107964 |
| *O. alta* | g1227 | Chr1CC | 11807768 | 11811601 |
| *O. alta* | g18155 | Chr3CC | 12152089 | 12155918 |
| *O. sativa* | Os03g0291500 | Chr3 | 10120289 | 10124384 |
| *O. alta* | g1228 | Chr1CC | 11835196 | 11839172 |
| *O. alta* | g18158 | Chr3CC | 12194835 | 12198843 |
| *O. sativa* | Os03g0291800 | Chr3 | 10148550 | 10153087 |
| *O. alta* | g1229 | Chr1CC | 11856889 | 11858345 |
| *O. alta* | g18160 | Chr3CC | 12219305 | 12220776 |
| *O. sativa* | Os03g0292100 | Chr3 | 10163441 | 10165442 |
| *O. alta* | g1230 | Chr1CC | 11864045 | 11866492 |
| *O. alta* | g18161 | Chr3CC | 12226516 | 12228791 |
| *O. sativa* | Os03g0292200 | Chr3 | 10171907 | 10174512 |
| *O. alta* | g1232 | Chr1CC | 11878382 | 11880526 |
| *O. alta* | g18163 | Chr3CC | 12249310 | 12252032 |
| *O. sativa* | Os03g0292800 | Chr3 | 10187429 | 10190243 |
| *O. alta* | g1234 | Chr1CC | 11895292 | 11900473 |
| *O. alta* | g18164 | Chr3CC | 12255925 | 12261145 |
| *O. sativa* | Os03g0293000 | Chr3 | 10206161 | 10208503 |

| | | | | |
|---|---|---|---|---|
| *O. alta* | g1235 | Chr1CC | 11901290 | 11904443 |
| *O. alta* | g18165 | Chr3CC | 12262503 | 12264147 |
| *O. sativa* | Os03g0293400 | Chr3 | 10214020 | 10215765 |
| *O. alta* | g1236 | Chr1CC | 11905899 | 11908455 |
| *O. alta* | g18167 | Chr3CC | 12267890 | 12270386 |
| *O. sativa* | Os03g0293500 | Chr3 | 10217810 | 10220748 |
| *O. alta* | g1238 | Chr1CC | 11923100 | 11925179 |
| *O. alta* | g18168 | Chr3CC | 12278843 | 12280925 |
| *O. sativa* | Os03g0294100 | Chr3 | 10255888 | 10257612 |
| *O. alta* | g1239 | Chr1CC | 11926456 | 11950027 |
| *O. alta* | g18169 | Chr3CC | 12282177 | 12296363 |
| *O. sativa* | Os03g0294200 | Chr3 | 10258670 | 10270774 |
| *O. alta* | g1241 | Chr1CC | 11958691 | 11959479 |
| *O. alta* | g18170 | Chr3CC | 12298221 | 12298979 |
| *O. alta* | g1242 | Chr1CC | 11961890 | 11964733 |
| *O. alta* | g18171 | Chr3CC | 12301321 | 12304193 |
| *O. sativa* | Os03g0294600 | Chr3 | 10279758 | 10281523 |
| *O. alta* | g1246 | Chr1CC | 12042690 | 12046698 |
| *O. alta* | g18172 | Chr3CC | 12312858 | 12316862 |
| *O. sativa* | Os03g0294700 | Chr3 | 10287901 | 10291530 |
| *O. alta* | g1247 | Chr1CC | 12048324 | 12051214 |
| *O. alta* | g18173 | Chr3CC | 12325180 | 12328349 |
| *O. sativa* | Os03g0294800 | Chr3 | 10292291 | 10295620 |
| *O. alta* | g1249 | Chr1CC | 12064469 | 12068896 |
| *O. alta* | g18175 | Chr3CC | 12339036 | 12343115 |
| *O. sativa* | Os03g0295400 | Chr3 | 10320222 | 10324794 |
| *O. alta* | g1250 | Chr1CC | 12075997 | 12079316 |
| *O. alta* | g18176 | Chr3CC | 12343549 | 12346873 |
| *O. sativa* | Os03g0295500 | Chr3 | 10325025 | 10330450 |
| *O. alta* | g1251 | Chr1CC | 12088727 | 12091395 |
| *O. alta* | g18177 | Chr3CC | 12352159 | 12354842 |
| *O. sativa* | Os03g0295600 | Chr3 | 10333323 | 10336943 |
| *O. alta* | g1252 | Chr1CC | 12105837 | 12119138 |
| *O. alta* | g18181 | Chr3CC | 12393225 | 12395014 |
| *O. sativa* | Os03g0295700 | Chr3 | 10344987 | 10355461 |

| | | | | |
|---|---|---|---|---|
| *O. alta* | g1254 | Chr1CC | 12123207 | 12123500 |
| *O. alta* | g18184 | Chr3CC | 12406841 | 12407116 |
| *O. sativa* | Os03g0295800 | Chr3 | 10355674 | 10359296 |
| *O. alta* | g1256 | Chr1CC | 12143194 | 12144915 |
| *O. alta* | g18186 | Chr3CC | 12413838 | 12415703 |
| *O. alta* | g1255 | Chr1CC | 12124127 | 12131025 |
| *O. alta* | g18185 | Chr3CC | 12409734 | 12410876 |
| *O. sativa* | Os03g0295866 | Chr3 | 10359847 | 10361451 |
| *O. alta* | g1257 | Chr1CC | 12146484 | 12147502 |
| *O. alta* | g18187 | Chr3CC | 12416897 | 12418037 |
| *O. sativa* | Os03g0296200 | Chr3 | 10366901 | 10368154 |
| *O. alta* | g1258 | Chr1CC | 12148351 | 12151243 |
| *O. alta* | g18188 | Chr3CC | 12421730 | 12421948 |
| *O. sativa* | Os03g0296300 | Chr3 | 10368893 | 10372593 |
| *O. alta* | g1259 | Chr1CC | 12151828 | 12154645 |
| *O. alta* | g18189 | Chr3CC | 12422542 | 12425657 |
| *O. sativa* | Os03g0296400 | Chr3 | 10372812 | 10376020 |
| *O. alta* | g1261 | Chr1CC | 12157594 | 12158361 |
| *O. alta* | g18190 | Chr3CC | 12426758 | 12429368 |
| *O. sativa* | Os03g0296500 | Chr3 | 10376613 | 10379598 |
| *O. alta* | g1262 | Chr1CC | 12160576 | 12161043 |
| *O. alta* | g18191 | Chr3CC | 12430591 | 12430932 |
| *O. sativa* | Os03g0296600 | Chr3 | 10380090 | 10380764 |
| *O. alta* | g1263 | Chr1CC | 12169050 | 12171484 |
| *O. alta* | g18192 | Chr3CC | 12439953 | 12442377 |
| *O. sativa* | Os03g0296800 | Chr3 | 10388030 | 10390923 |
| *O. alta* | g1264 | Chr1CC | 12172664 | 12173263 |
| *O. alta* | g18193 | Chr3CC | 12443649 | 12444255 |
| *O. sativa* | Os03g0297000 | Chr3 | 10391789 | 10392766 |
| *O. alta* | g1265 | Chr1CC | 12174819 | 12176469 |
| *O. alta* | g18194 | Chr3CC | 12445384 | 12447037 |
| *O. sativa* | Os03g0297100 | Chr3 | 10393657 | 10395829 |
| *O. alta* | g1266 | Chr1CC | 12177627 | 12180737 |
| *O. alta* | g18195 | Chr3CC | 12448121 | 12451213 |
| *O. sativa* | Os03g0297400 | Chr3 | 10400794 | 10404233 |

| | | | | |
|---|---|---|---|---|
| *O. alta* | g1268 | Chr1CC | 12209423 | 12210109 |
| *O. alta* | g18196 | Chr3CC | 12463861 | 12464547 |
| *O. sativa* | Os03g0297600 | Chr3 | 10418661 | 10419833 |
| *O. alta* | g1271 | Chr1CC | 12240579 | 12243199 |
| *O. alta* | g18197 | Chr3CC | 12476969 | 12480618 |
| *O. sativa* | Os03g0297700 | Chr3 | 10428927 | 10433729 |
| *O. alta* | g1272 | Chr1CC | 12244942 | 12247688 |
| *O. alta* | g18198 | Chr3CC | 12482191 | 12484860 |
| *O. sativa* | Os03g0297800 | Chr3 | 10438693 | 10440452 |
| *O. alta* | g1273 | Chr1CC | 12249843 | 12252664 |
| *O. alta* | g18199 | Chr3CC | 12487549 | 12490472 |
| *O. sativa* | Os03g0297900 | Chr3 | 10443193 | 10445991 |
| *O. alta* | g1274 | Chr1CC | 12256248 | 12256796 |
| *O. alta* | g18200 | Chr3CC | 12494782 | 12495321 |
| *O. sativa* | Os03g0298100 | Chr3 | 10452240 | 10453026 |
| *O. alta* | g1275 | Chr1CC | 12257715 | 12258389 |
| *O. alta* | g18201 | Chr3CC | 12496421 | 12496837 |
| *O. alta* | g1276 | Chr1CC | 12262202 | 12264249 |
| *O. alta* | g18204 | Chr3CC | 12512843 | 12514905 |
| *O. sativa* | Os03g0298300 | Chr3 | 10456984 | 10459641 |
| *O. alta* | g1277 | Chr1CC | 12269579 | 12276485 |
| *O. alta* | g18205 | Chr3CC | 12518513 | 12522142 |
| *O. sativa* | Os03g0298400 | Chr3 | 10462452 | 10466149 |
| *O. alta* | g1278 | Chr1CC | 12278398 | 12280010 |
| *O. alta* | g18206 | Chr3CC | 12523439 | 12525793 |
| *O. sativa* | Os03g0298600 | Chr3 | 10467372 | 10469878 |
| *O. alta* | g1279 | Chr1CC | 12281289 | 12286938 |
| *O. alta* | g18207 | Chr3CC | 12527259 | 12532968 |
| *O. sativa* | Os03g0298700 | Chr3 | 10470676 | 10477048 |
| *O. alta* | g1280 | Chr1CC | 12289204 | 12293311 |
| *O. alta* | g18208 | Chr3CC | 12534780 | 12538835 |
| *O. sativa* | Os03g0298800 | Chr3 | 10478466 | 10482571 |
| *O. alta* | g1281 | Chr1CC | 12304841 | 12305939 |
| *O. alta* | g18209 | Chr3CC | 12596140 | 12597231 |
| *O. sativa* | Os03g0299200 | Chr3 | 10502015 | 10503291 |

| | | | | |
|---|---|---|---|---|
| *O. alta* | g1283 | Chr1CC | 12352518 | 12355089 |
| *O. alta* | g18212 | Chr3CC | 12615856 | 12617757 |
| *O. sativa* | Os03g0299600 | Chr3 | 10517166 | 10518361 |
| *O. alta* | g1284 | Chr1CC | 12360376 | 12362091 |
| *O. alta* | g18213 | Chr3CC | 12628588 | 12630306 |
| *O. sativa* | Os03g0299800 | Chr3 | 10525760 | 10531373 |
| *O. alta* | g1285 | Chr1CC | 12375165 | 12378175 |
| *O. alta* | g18214 | Chr3CC | 12641090 | 12644068 |
| *O. sativa* | Os03g0299900 | Chr3 | 10542880 | 10546261 |
| *O. alta* | g1286 | Chr1CC | 12379898 | 12381247 |
| *O. alta* | g18215 | Chr3CC | 12649053 | 12650396 |
| *O. sativa* | Os03g0300000 | Chr3 | 10547471 | 10549503 |
| *O. alta* | g1287 | Chr1CC | 12383790 | 12387703 |
| *O. alta* | g18216 | Chr3CC | 12657416 | 12660550 |
| *O. sativa* | Os03g0300200 | Chr3 | 10551935 | 10556107 |
| *O. alta* | g1288 | Chr1CC | 12387961 | 12393292 |
| *O. alta* | g18217 | Chr3CC | 12660832 | 12666128 |
| *O. sativa* | Os03g0300300 | Chr3 | 10556232 | 10561713 |
| *O. alta* | g1289 | Chr1CC | 12394077 | 12394637 |
| *O. alta* | g18218 | Chr3CC | 12666945 | 12667515 |
| *O. sativa* | Os03g0300400 | Chr3 | 10562231 | 10563277 |
| *O. alta* | g1290 | Chr1CC | 12396948 | 12398117 |
| *O. alta* | g18219 | Chr3CC | 12685363 | 12687060 |
| *O. sativa* | Os03g0300500 | Chr3 | 10564710 | 10566806 |
| *O. alta* | g1292 | Chr1CC | 12401739 | 12403216 |
| *O. alta* | g18220 | Chr3CC | 12692484 | 12693404 |
| *O. sativa* | Os03g0300600 | Chr3 | 10571694 | 10573269 |
| *O. alta* | g1293 | Chr1CC | 12425108 | 12425377 |
| *O. alta* | g18221 | Chr3CC | 12696160 | 12697131 |
| *O. sativa* | Os03g0300700 | Chr3 | 10575398 | 10576700 |
| *O. alta* | g1295 | Chr1CC | 12434501 | 12435607 |
| *O. alta* | g18222 | Chr3CC | 12704149 | 12705255 |
| *O. sativa* | Os03g0300900 | Chr3 | 10583988 | 10586011 |
| *O. alta* | g1296 | Chr1CC | 12467832 | 12469853 |
| *O. alta* | g18223 | Chr3CC | 12726439 | 12728472 |
| *O. sativa* | Os03g0301200 | Chr3 | 10599142 | 10601525 |

| | | | | |
|---|---|---|---|---|
| *O. alta* | g1297 | Chr1CC | 12476305 | 12482760 |
| *O. alta* | g18224 | Chr3CC | 12735722 | 12739551 |
| *O. alta* | g1298 | Chr1CC | 12486717 | 12489358 |
| *O. alta* | g18226 | Chr3CC | 12744802 | 12747407 |
| *O. sativa* | Os03g0301600 | Chr3 | 10620065 | 10620827 |
| *O. alta* | g1299 | Chr1CC | 12491089 | 12494005 |
| *O. alta* | g18227 | Chr3CC | 12749077 | 12752174 |
| *O. sativa* | Os03g0301700 | Chr3 | 10621704 | 10625395 |
| *O. alta* | g1300 | Chr1CC | 12500135 | 12505782 |
| *O. alta* | g18228 | Chr3CC | 12756418 | 12761298 |
| *O. sativa* | Os03g0301800 | Chr3 | 10629886 | 10632864 |
| *O. alta* | g1301 | Chr1CC | 12524434 | 12525183 |
| *O. alta* | g18229 | Chr3CC | 12795187 | 12795939 |
| *O. sativa* | Os03g0301950 | Chr3 | 10645184 | 10646734 |
| *O. alta* | g1302 | Chr1CC | 12533279 | 12534007 |
| *O. alta* | g18230 | Chr3CC | 12797773 | 12798501 |
| *O. sativa* | Os03g0302000 | Chr3 | 10647683 | 10648574 |
| *O. alta* | g1303 | Chr1CC | 12537466 | 12539821 |
| *O. alta* | g18231 | Chr3CC | 12802081 | 12805526 |
| *O. sativa* | Os03g0302200 | Chr3 | 10651199 | 10655589 |
| *O. alta* | g1306 | Chr1CC | 12556986 | 12558019 |
| *O. alta* | g18233 | Chr3CC | 12814489 | 12814743 |
| *O. sativa* | Os03g0302700 | Chr3 | 10672591 | 10674055 |
| *O. alta* | g1307 | Chr1CC | 12561484 | 12562149 |
| *O. alta* | g18234 | Chr3CC | 12818250 | 12818900 |
| *O. sativa* | Os03g0302800 | Chr3 | 10677797 | 10678946 |
| *O. alta* | g1309 | Chr1CC | 12572020 | 12580693 |
| *O. alta* | g18235 | Chr3CC | 12826966 | 12830952 |
| *O. sativa* | Os03g0302900 | Chr3 | 10684315 | 10688768 |
| *O. alta* | g76 | Chr1CC | 1075834 | 1081919 |
| *O. alta* | g16988 | Chr3CC | 1549934 | 1555429 |
| *O. sativa* | Os03g0122000 | Chr3 | 1211681 | 1217860 |
| *O. alta* | g26255 | Chr4CC | 33527897 | 33529264 |
| *O. alta* | g43472 | Chr7CC | 34535491 | 34536858 |
| *O. sativa* | Os04g0619300 | Chr4 | 31458951 | 31462013 |

| | | | | |
|---|---|---|---|---|
| *O. alta* | g26256 | Chr4CC | 33530852 | 33532485 |
| *O. alta* | g43473 | Chr7CC | 34538450 | 34540084 |
| *O. sativa* | Os04g0619400 | Chr4 | 31463105 | 31466291 |
| *O. alta* | g26257 | Chr4CC | 33540974 | 33543545 |
| *O. alta* | g43474 | Chr7CC | 34549230 | 34551811 |
| *O. sativa* | Os04g0619500 | Chr4 | 31474399 | 31477315 |
| *O. alta* | g26258 | Chr4CC | 33545252 | 33547786 |
| *O. alta* | g43476 | Chr7CC | 34555655 | 34557277 |
| *O. sativa* | Os04g0619600 | Chr4 | 31478250 | 31481997 |
| *O. alta* | g26260 | Chr4CC | 33552319 | 33554001 |
| *O. alta* | g43478 | Chr7CC | 34562500 | 34564185 |
| *O. sativa* | Os04g0619700 | Chr4 | 31485367 | 31488482 |
| *O. alta* | g26261 | Chr4CC | 33564821 | 33566141 |
| *O. alta* | g43481 | Chr7CC | 34570254 | 34571565 |
| *O. sativa* | Os04g0619800 | Chr4 | 31492252 | 31493474 |
| *O. alta* | g26262 | Chr4CC | 33583165 | 33587859 |
| *O. alta* | g43482 | Chr7CC | 34591255 | 34596104 |
| *O. sativa* | Os04g0619900 | Chr4 | 31497058 | 31500179 |
| *O. alta* | g26263 | Chr4CC | 33588797 | 33599796 |
| *O. alta* | g43483 | Chr7CC | 34597024 | 34608429 |
| *O. sativa* | Os04g0620000 | Chr4 | 31502967 | 31514308 |
| *O. alta* | g26264 | Chr4CC | 33608354 | 33611218 |
| *O. alta* | g43484 | Chr7CC | 34621760 | 34623960 |
| *O. sativa* | Os04g0620200 | Chr4 | 31524614 | 31527861 |
| *O. alta* | g26266 | Chr4CC | 33620351 | 33626072 |
| *O. alta* | g43488 | Chr7CC | 34644245 | 34651971 |
| *O. sativa* | Os04g0620400 | Chr4 | 31531978 | 31541045 |
| *O. alta* | g26267 | Chr4CC | 33626780 | 33627220 |
| *O. alta* | g43489 | Chr7CC | 34652771 | 34653123 |
| *O. sativa* | Os04g0620600 | Chr4 | 31541994 | 31542752 |
| *O. alta* | g26269 | Chr4CC | 33638144 | 33638491 |
| *O. alta* | g43490 | Chr7CC | 34655754 | 34656098 |
| *O. alta* | g26270 | Chr4CC | 33652843 | 33658129 |
| *O. alta* | g43491 | Chr7CC | 34682945 | 34688081 |
| *O. sativa* | Os04g0620700 | Chr4 | 31544124 | 31551203 |

| | | | | |
|---|---|---|---|---|
| *O. alta* | g26283 | Chr4CC | 33779179 | 33779616 |
| *O. alta* | g43494 | Chr7CC | 34709988 | 34710404 |
| *O. sativa* | Os04g0623200 | Chr4 | 31666207 | 31667028 |
| *O. alta* | g26284 | Chr4CC | 33782468 | 33786618 |
| *O. alta* | g43496 | Chr7CC | 34728072 | 34732227 |
| *O. sativa* | Os04g0623300 | Chr4 | 31669220 | 31674595 |
| *O. alta* | g26286 | Chr4CC | 33806491 | 33808757 |
| *O. alta* | g43498 | Chr7CC | 34755644 | 34758791 |
| *O. sativa* | Os04g0623500 | Chr4 | 31688721 | 31692502 |
| *O. alta* | g26287 | Chr4CC | 33809636 | 33812197 |
| *O. alta* | g43500 | Chr7CC | 34763568 | 34766032 |
| *O. sativa* | Os04g0623700 | Chr4 | 31697288 | 31700055 |
| *O. alta* | g26288 | Chr4CC | 33813311 | 33815779 |
| *O. alta* | g43501 | Chr7CC | 34767132 | 34769608 |
| *O. sativa* | Os04g0623800 | Chr4 | 31700976 | 31703589 |
| *O. alta* | g26292 | Chr4CC | 33844289 | 33846332 |
| *O. alta* | g43507 | Chr7CC | 34793805 | 34795081 |
| *O. sativa* | Os04g0624450 | Chr4 | 31740856 | 31741449 |
| *O. alta* | g26295 | Chr4CC | 33868702 | 33875876 |
| *O. alta* | g43508 | Chr7CC | 34795713 | 34803169 |
| *O. sativa* | Os04g0624600 | Chr4 | 31751600 | 31759420 |
| *O. alta* | g26296 | Chr4CC | 33881194 | 33883861 |
| *O. alta* | g43509 | Chr7CC | 34808656 | 34811743 |
| *O. sativa* | Os04g0624800 | Chr4 | 31764133 | 31766854 |
| *O. alta* | g26297 | Chr4CC | 33884023 | 33884292 |
| *O. alta* | g43510 | Chr7CC | 34813079 | 34813348 |
| *O. sativa* | Os04g0624900 | Chr4 | 31767073 | 31767689 |
| *O. alta* | g26298 | Chr4CC | 33886530 | 33890163 |
| *O. alta* | g43511 | Chr7CC | 34821693 | 34825307 |
| *O. sativa* | Os04g0625000 | Chr4 | 31769056 | 31773154 |
| *O. alta* | g26299 | Chr4CC | 33896307 | 33899303 |
| *O. alta* | g43512 | Chr7CC | 34849049 | 34852150 |
| *O. sativa* | Os04g0625100 | Chr4 | 31777314 | 31780978 |
| *O. alta* | g26300 | Chr4CC | 33906686 | 33907932 |
| *O. alta* | g43513 | Chr7CC | 34864469 | 34867199 |

| | | | | |
|---|---|---|---|---|
| *O. sativa* | Os04g0625200 | Chr4 | 31786589 | 31788246 |
| *O. alta* | g26301 | Chr4CC | 33909058 | 33920945 |
| *O. alta* | g43514 | Chr7CC | 34867642 | 34880194 |
| *O. sativa* | Os04g0625300 | Chr4 | 31788464 | 31790490 |
| *O. alta* | g26303 | Chr4CC | 33948602 | 33949723 |
| *O. alta* | g43516 | Chr7CC | 34882375 | 34883507 |
| *O. sativa* | Os04g0625500 | Chr4 | 31809708 | 31810835 |
| *O. alta* | g26304 | Chr4CC | 33950813 | 33951913 |
| *O. alta* | g43518 | Chr7CC | 34887726 | 34888850 |
| *O. sativa* | Os04g0625600 | Chr4 | 31811661 | 31813399 |
| *O. alta* | g26306 | Chr4CC | 33955204 | 33958884 |
| *O. alta* | g43519 | Chr7CC | 34890143 | 34893792 |
| *O. alta* | g26307 | Chr4CC | 33959906 | 33963981 |
| *O. alta* | g43520 | Chr7CC | 34895415 | 34899720 |
| *O. sativa* | Os04g0625900 | Chr4 | 31824102 | 31828850 |
| *O. alta* | g26308 | Chr4CC | 33968459 | 33970664 |
| *O. alta* | g43521 | Chr7CC | 34918455 | 34919456 |
| *O. sativa* | Os04g0626000 | Chr4 | 31831245 | 31836454 |
| *O. alta* | g26309 | Chr4CC | 33978589 | 33979218 |
| *O. alta* | g43522 | Chr7CC | 34923218 | 34923829 |
| *O. sativa* | Os04g0626100 | Chr4 | 31838166 | 31842050 |
| *O. alta* | g26313 | Chr4CC | 34056517 | 34065665 |
| *O. alta* | g43528 | Chr7CC | 34961871 | 34966340 |
| *O. sativa* | Os04g0626900 | Chr4 | 31895293 | 31900168 |
| *O. alta* | g26314 | Chr4CC | 34066734 | 34072175 |
| *O. alta* | g43529 | Chr7CC | 34967376 | 34972793 |
| *O. sativa* | Os04g0627000 | Chr4 | 31900217 | 31907077 |
| *O. alta* | g26315 | Chr4CC | 34092322 | 34092996 |
| *O. alta* | g43530 | Chr7CC | 34984829 | 34987912 |
| *O. sativa* | Os04g0627200 | Chr4 | 31927789 | 31928733 |
| *O. alta* | g26316 | Chr4CC | 34095986 | 34096731 |
| *O. alta* | g43531 | Chr7CC | 34997393 | 34998125 |
| *O. sativa* | Os04g0627300 | Chr4 | 31932864 | 31933709 |
| *O. alta* | g26324 | Chr4CC | 34133941 | 34134422 |
| *O. alta* | g43532 | Chr7CC | 35005691 | 35007639 |

| | | | | |
|---|---|---|---|---|
| *O. sativa* | Os04g0627400 | Chr4 | 31937356 | 31939121 |
| *O. alta* | g26326 | Chr4CC | 34140372 | 34141717 |
| *O. alta* | g43535 | Chr7CC | 35020764 | 35024052 |
| *O. sativa* | Os04g0628000 | Chr4 | 31959627 | 31961695 |
| *O. alta* | g26328 | Chr4CC | 34145466 | 34147306 |
| *O. alta* | g43537 | Chr7CC | 35039329 | 35041028 |
| *O. sativa* | Os04g0628200 | Chr4 | 31965780 | 31973005 |
| *O. alta* | g26329 | Chr4CC | 34150094 | 34151865 |
| *O. alta* | g43538 | Chr7CC | 35043942 | 35045751 |
| *O. alta* | g26330 | Chr4CC | 34153433 | 34154089 |
| *O. alta* | g43539 | Chr7CC | 35048216 | 35048854 |
| *O. sativa* | Os04g0628300 | Chr4 | 31975720 | 31976388 |
| *O. alta* | g26331 | Chr4CC | 34156296 | 34158897 |
| *O. alta* | g43540 | Chr7CC | 35050428 | 35053023 |
| *O. sativa* | Os04g0628400 | Chr4 | 31977268 | 31981158 |
| *O. alta* | g26332 | Chr4CC | 34168099 | 34175083 |
| *O. alta* | g43541 | Chr7CC | 35106965 | 35113924 |
| *O. sativa* | Os04g0628600 | Chr4 | 31982348 | 31992462 |
| *O. alta* | g26333 | Chr4CC | 34182114 | 34183950 |
| *O. alta* | g43542 | Chr7CC | 35116591 | 35118363 |
| *O. sativa* | Os04g0628900 | Chr4 | 31995412 | 31998252 |
| *O. alta* | g26334 | Chr4CC | 34185281 | 34187743 |
| *O. alta* | g43543 | Chr7CC | 35119893 | 35122423 |
| *O. sativa* | Os04g0629000 | Chr4 | 31999368 | 32002716 |
| *O. alta* | g26335 | Chr4CC | 34188383 | 34192674 |
| *O. alta* | g43544 | Chr7CC | 35123179 | 35127510 |
| *O. sativa* | Os04g0629100 | Chr4 | 32002722 | 32007327 |
| *O. alta* | g26338 | Chr4CC | 34205654 | 34206244 |
| *O. alta* | g43546 | Chr7CC | 35136278 | 35136849 |
| *O. sativa* | Os04g0629200 | Chr4 | 32008449 | 32009263 |
| *O. alta* | g26339 | Chr4CC | 34207041 | 34212494 |
| *O. alta* | g43547 | Chr7CC | 35137640 | 35143059 |
| *O. sativa* | Os04g0629300 | Chr4 | 32009536 | 32016407 |
| *O. alta* | g26340 | Chr4CC | 34214440 | 34214787 |
| *O. alta* | g43548 | Chr7CC | 35144970 | 35145347 |

| | | | | |
|---|---|---|---|---|
| *O. sativa* | Os04g0629400 | Chr4 | 32017656 | 32018383 |
| *O. alta* | g26341 | Chr4CC | 34220528 | 34221659 |
| *O. alta* | g43549 | Chr7CC | 35147075 | 35148652 |
| *O. sativa* | Os04g0629500 | Chr4 | 32019411 | 32022508 |
| *O. alta* | g26342 | Chr4CC | 34224627 | 34229052 |
| *O. alta* | g43550 | Chr7CC | 35152712 | 35157188 |
| *O. sativa* | Os04g0629700 | Chr4 | 32033904 | 32039009 |
| *O. alta* | g26343 | Chr4CC | 34232991 | 34237890 |
| *O. alta* | g43551 | Chr7CC | 35160219 | 35164852 |
| *O. sativa* | Os04g0630000 | Chr4 | 32044764 | 32048497 |
| *O. alta* | g26346 | Chr4CC | 34270288 | 34271894 |
| *O. alta* | g43553 | Chr7CC | 35173117 | 35175709 |
| *O. sativa* | Os04g0630300 | Chr4 | 32059560 | 32062390 |
| *O. alta* | g26351 | Chr4CC | 34302684 | 34305164 |
| *O. alta* | g43555 | Chr7CC | 35193433 | 35198966 |
| *O. sativa* | Os04g0630800 | Chr4 | 32085612 | 32088272 |
| *O. alta* | g26352 | Chr4CC | 34310875 | 34314171 |
| *O. alta* | g43557 | Chr7CC | 35235539 | 35238831 |
| *O. sativa* | Os04g0631100 | Chr4 | 32139084 | 32140993 |
| *O. alta* | g26353 | Chr4CC | 34322703 | 34324155 |
| *O. alta* | g43558 | Chr7CC | 35245082 | 35246532 |
| *O. alta* | g26354 | Chr4CC | 34335186 | 34340085 |
| *O. alta* | g43559 | Chr7CC | 35262993 | 35267906 |
| *O. sativa* | Os04g0631600 | Chr4 | 32166724 | 32169575 |
| *O. alta* | g26355 | Chr4CC | 34341240 | 34344289 |
| *O. alta* | g43560 | Chr7CC | 35270925 | 35272268 |
| *O. sativa* | Os04g0631800 | Chr4 | 32173571 | 32177352 |
| *O. alta* | g26359 | Chr4CC | 34373177 | 34373659 |
| *O. alta* | g43561 | Chr7CC | 35277509 | 35285406 |
| *O. sativa* | Os04g0632300 | Chr4 | 32183880 | 32187311 |
| *O. alta* | g26363 | Chr4CC | 34389580 | 34392645 |
| *O. alta* | g43563 | Chr7CC | 35297379 | 35300969 |
| *O. sativa* | Os04g0633300 | Chr4 | 32229267 | 32232359 |
| *O. alta* | g26374 | Chr4CC | 34461804 | 34465635 |
| *O. alta* | g43566 | Chr7CC | 35314522 | 35317700 |

| | | | | |
|---|---|---|---|---|
| *O. sativa* | Os04g0633900 | Chr4 | 32248949 | 32253028 |
| *O. alta* | g26376 | Chr4CC | 34520205 | 34522325 |
| *O. alta* | g43607 | Chr7CC | 35608229 | 35612448 |
| *O. sativa* | Os04g0634500 | Chr4 | 32278194 | 32281408 |
| *O. alta* | g26375 | Chr4CC | 34515980 | 34519440 |
| *O. alta* | g43608 | Chr7CC | 35613750 | 35617159 |
| *O. sativa* | Os04g0634400 | Chr4 | 32273357 | 32276841 |
| *O. alta* | g26377 | Chr4CC | 34526388 | 34529798 |
| *O. alta* | g43606 | Chr7CC | 35601322 | 35604645 |
| *O. sativa* | Os04g0634700 | Chr4 | 32286661 | 32291433 |
| *O. alta* | g26378 | Chr4CC | 34531549 | 34531812 |
| *O. alta* | g43605 | Chr7CC | 35599890 | 35600159 |
| *O. sativa* | Os04g0634800 | Chr4 | 32294993 | 32295517 |
| *O. alta* | g26379 | Chr4CC | 34532634 | 34532900 |
| *O. alta* | g43604 | Chr7CC | 35598681 | 35598968 |
| *O. sativa* | Os04g0634900 | Chr4 | 32296094 | 32296600 |
| *O. alta* | g26380 | Chr4CC | 34534067 | 34534366 |
| *O. alta* | g43603 | Chr7CC | 35596900 | 35597196 |
| *O. sativa* | Os04g0635000 | Chr4 | 32297571 | 32298126 |
| *O. alta* | g26381 | Chr4CC | 34537293 | 34537565 |
| *O. alta* | g43602 | Chr7CC | 35593475 | 35593744 |
| *O. alta* | g26383 | Chr4CC | 34543906 | 34544166 |
| *O. alta* | g43599 | Chr7CC | 35573716 | 35573976 |
| *O. sativa* | Os04g0635500 | Chr4 | 32333452 | 32333991 |
| *O. alta* | g26386 | Chr4CC | 34553390 | 34553989 |
| *O. alta* | g43598 | Chr7CC | 35569603 | 35570202 |
| *O. sativa* | Os04g0635800 | Chr4 | 32339714 | 32342396 |
| *O. alta* | g26387 | Chr4CC | 34554285 | 34559798 |
| *O. alta* | g43597 | Chr7CC | 35562376 | 35568674 |
| *O. sativa* | Os04g0635900 | Chr4 | 32342580 | 32348393 |
| *O. alta* | g26388 | Chr4CC | 34560381 | 34563364 |
| *O. alta* | g43596 | Chr7CC | 35560461 | 35562021 |
| *O. sativa* | Os04g0636000 | Chr4 | 32348410 | 32351851 |
| *O. alta* | g26389 | Chr4CC | 34565009 | 34566829 |
| *O. alta* | g43595 | Chr7CC | 35556873 | 35558707 |

| | | | | |
|---|---|---|---|---|
| *O. sativa* | Os04g0636100 | Chr4 | 32353487 | 32355998 |
| *O. alta* | g26394 | Chr4CC | 34578801 | 34581894 |
| *O. alta* | g43591 | Chr7CC | 35542836 | 35545835 |
| *O. sativa* | Os04g0636500 | Chr4 | 32365686 | 32369961 |
| *O. alta* | g26390 | Chr4CC | 34567803 | 34569759 |
| *O. alta* | g43594 | Chr7CC | 35553976 | 35555891 |
| *O. sativa* | Os04g0636200 | Chr4 | 32356297 | 32358756 |
| *O. alta* | g26392 | Chr4CC | 34574450 | 34576314 |
| *O. alta* | g43592 | Chr7CC | 35547072 | 35549026 |
| *O. sativa* | Os04g0636400 | Chr4 | 32363010 | 32365037 |
| *O. alta* | g26395 | Chr4CC | 34582788 | 34589728 |
| *O. alta* | g43590 | Chr7CC | 35535026 | 35541993 |
| *O. sativa* | Os04g0636600 | Chr4 | 32370472 | 32378265 |
| *O. alta* | g26397 | Chr4CC | 34598905 | 34604363 |
| *O. alta* | g43587 | Chr7CC | 35514270 | 35516345 |
| *O. sativa* | Os04g0636800 | Chr4 | 32384028 | 32387076 |
| *O. alta* | g26398 | Chr4CC | 34605777 | 34607321 |
| *O. alta* | g43586 | Chr7CC | 35507859 | 35509397 |
| *O. sativa* | Os04g0636900 | Chr4 | 32390377 | 32392487 |
| *O. alta* | g26400 | Chr4CC | 34624321 | 34626271 |
| *O. alta* | g43585 | Chr7CC | 35493336 | 35495285 |
| *O. sativa* | Os04g0637000 | Chr4 | 32403357 | 32412059 |
| *O. alta* | g26401 | Chr4CC | 34628735 | 34629370 |
| *O. alta* | g43584 | Chr7CC | 35481535 | 35482061 |
| *O. alta* | g26402 | Chr4CC | 34631012 | 34637446 |
| *O. alta* | g43583 | Chr7CC | 35473795 | 35480296 |
| *O. sativa* | Os04g0637400 | Chr4 | 32419840 | 32423901 |
| *O. alta* | g26403 | Chr4CC | 34638447 | 34639622 |
| *O. alta* | g43582 | Chr7CC | 35471753 | 35472989 |
| *O. sativa* | Os04g0637500 | Chr4 | 32424262 | 32425566 |
| *O. alta* | g26404 | Chr4CC | 34681760 | 34683200 |
| *O. alta* | g43581 | Chr7CC | 35469625 | 35471707 |
| *O. sativa* | Os04g0638100 | Chr4 | 32448789 | 32450307 |
| *O. alta* | g26407 | Chr4CC | 34725019 | 34725561 |
| *O. alta* | g43580 | Chr7CC | 35447389 | 35447951 |

| | | | | |
|---|---|---|---|---|
| *O. sativa* | Os04g0638700 | Chr4 | 32459623 | 32460498 |
| *O. alta* | g26411 | Chr4CC | 34771735 | 34772601 |
| *O. alta* | g43579 | Chr7CC | 35441844 | 35442731 |
| *O. sativa* | Os04g0638800 | Chr4 | 32464933 | 32466032 |
| *O. alta* | g26413 | Chr4CC | 34787452 | 34787906 |
| *O. alta* | g43578 | Chr7CC | 35420864 | 35421318 |
| *O. sativa* | Os04g0639100 | Chr4 | 32478683 | 32479514 |
| *O. alta* | g26414 | Chr4CC | 34789490 | 34790171 |
| *O. alta* | g43576 | Chr7CC | 35418508 | 35419186 |
| *O. sativa* | Os04g0639200 | Chr4 | 32481008 | 32482342 |
| *O. alta* | g26415 | Chr4CC | 34793243 | 34795579 |
| *O. alta* | g43575 | Chr7CC | 35412398 | 35414727 |
| *O. sativa* | Os04g0639300 | Chr4 | 32483088 | 32484417 |
| *O. alta* | g26416 | Chr4CC | 34796003 | 34802641 |
| *O. alta* | g43574 | Chr7CC | 35404923 | 35411992 |
| *O. sativa* | Os04g0640500 | Chr4 | 32574405 | 32581634 |
| *O. alta* | g26418 | Chr4CC | 34817486 | 34823774 |
| *O. alta* | g43572 | Chr7CC | 35388693 | 35396999 |
| *O. sativa* | Os04g0640700 | Chr4 | 32589116 | 32595893 |
| *O. alta* | g26419 | Chr4CC | 34828587 | 34830929 |
| *O. alta* | g43571 | Chr7CC | 35379782 | 35381817 |
| *O. sativa* | Os04g0640800 | Chr4 | 32600395 | 32603306 |
| *O. alta* | g26420 | Chr4CC | 34834670 | 34836583 |
| *O. alta* | g43570 | Chr7CC | 35373670 | 35375538 |
| *O. sativa* | Os04g0640900 | Chr4 | 32606375 | 32608648 |
| *O. alta* | g26421 | Chr4CC | 34849360 | 34858469 |
| *O. alta* | g43569 | Chr7CC | 35355071 | 35362075 |
| *O. sativa* | Os04g0641000 | Chr4 | 32609882 | 32618925 |
| *O. alta* | g26422 | Chr4CC | 34864372 | 34866615 |
| *O. alta* | g43609 | Chr7CC | 35626391 | 35629332 |
| *O. sativa* | Os04g0641200 | Chr4 | 32625806 | 32629133 |
| *O. alta* | g26424 | Chr4CC | 34870362 | 34871573 |
| *O. alta* | g43610 | Chr7CC | 35644213 | 35645430 |
| *O. sativa* | Os04g0641300 | Chr4 | 32632279 | 32633954 |
| *O. alta* | g26425 | Chr4CC | 34872221 | 34873905 |

| | | | | |
|---|---|---|---|---|
| *O. alta* | g43611 | Chr7CC | 35646079 | 35647821 |
| *O. sativa* | Os04g0641400 | Chr4 | 32634024 | 32637824 |
| *O. alta* | g26426 | Chr4CC | 34892170 | 34892579 |
| *O. alta* | g43613 | Chr7CC | 35667428 | 35667826 |
| *O. alta* | g26428 | Chr4CC | 34917126 | 34922965 |
| *O. alta* | g43615 | Chr7CC | 35683495 | 35689622 |
| *O. sativa* | Os04g0642000 | Chr4 | 32671919 | 32678320 |
| *O. alta* | g26434 | Chr4CC | 34961944 | 34962753 |
| *O. alta* | g43625 | Chr7CC | 35753827 | 35754636 |
| *O. sativa* | Os04g0643100 | Chr4 | 32732642 | 32735542 |
| *O. alta* | g26430 | Chr4CC | 34940084 | 34940821 |
| *O. alta* | g43620 | Chr7CC | 35735404 | 35735757 |
| *O. alta* | g26432 | Chr4CC | 34946060 | 34947636 |
| *O. alta* | g43622 | Chr7CC | 35740468 | 35742054 |
| *O. alta* | g26433 | Chr4CC | 34953490 | 34957936 |
| *O. alta* | g43624 | Chr7CC | 35747795 | 35752172 |
| *O. sativa* | Os04g0643000 | Chr4 | 32726344 | 32730766 |
| *O. alta* | g26436 | Chr4CC | 34972680 | 34977150 |
| *O. alta* | g43627 | Chr7CC | 35768463 | 35772843 |
| *O. sativa* | Os04g0643300 | Chr4 | 32741436 | 32746458 |
| *O. alta* | g26437 | Chr4CC | 34978381 | 34978626 |
| *O. alta* | g43628 | Chr7CC | 35774282 | 35774527 |
| *O. sativa* | Os04g0643500 | Chr4 | 32747192 | 32750831 |
| *O. alta* | g26440 | Chr4CC | 34989864 | 34991207 |
| *O. alta* | g43631 | Chr7CC | 35799937 | 35801268 |
| *O. sativa* | Os04g0643700 | Chr4 | 32758880 | 32762453 |
| *O. alta* | g26442 | Chr4CC | 34999589 | 35001025 |
| *O. alta* | g43632 | Chr7CC | 35811406 | 35812860 |
| *O. sativa* | Os04g0643800 | Chr4 | 32764848 | 32765688 |
| *O. alta* | g26446 | Chr4CC | 35027687 | 35031244 |
| *O. alta* | g43633 | Chr7CC | 35824321 | 35828121 |
| *O. sativa* | Os04g0644300 | Chr4 | 32791897 | 32792697 |
| *O. alta* | g26443 | Chr4CC | 35004126 | 35006466 |
| *O. alta* | g43634 | Chr7CC | 35829915 | 35832252 |
| *O. sativa* | Os04g0644000 | Chr4 | 32769559 | 32772259 |

| | | | | |
|---|---|---|---|---|
| *O. alta* | g26444 | Chr4CC | 35011405 | 35011971 |
| *O. alta* | g43635 | Chr7CC | 35833289 | 35833809 |
| *O. sativa* | Os04g0644100 | Chr4 | 32773316 | 32774144 |
| *O. alta* | g26445 | Chr4CC | 35017915 | 35019231 |
| *O. alta* | g43636 | Chr7CC | 35844025 | 35845452 |
| *O. sativa* | Os04g0644200 | Chr4 | 32777808 | 32779467 |
| *O. alta* | g26447 | Chr4CC | 35031976 | 35032797 |
| *O. alta* | g43638 | Chr7CC | 35857763 | 35858869 |
| *O. sativa* | Os04g0644400 | Chr4 | 32792893 | 32794163 |
| *O. alta* | g26448 | Chr4CC | 35034811 | 35035203 |
| *O. alta* | g43639 | Chr7CC | 35860133 | 35860519 |
| *O. alta* | g26449 | Chr4CC | 35042839 | 35047583 |
| *O. alta* | g43640 | Chr7CC | 35866461 | 35871090 |
| *O. sativa* | Os04g0644600 | Chr4 | 32801074 | 32807528 |
| *O. alta* | g26451 | Chr4CC | 35058498 | 35062315 |
| *O. alta* | g43642 | Chr7CC | 35900532 | 35902274 |
| *O. sativa* | Os04g0644700 | Chr4 | 32817477 | 32819752 |
| *O. alta* | g26452 | Chr4CC | 35063278 | 35067650 |
| *O. alta* | g43643 | Chr7CC | 35903268 | 35907669 |
| *O. alta* | g26453 | Chr4CC | 35069245 | 35073572 |
| *O. alta* | g43644 | Chr7CC | 35912522 | 35917894 |
| *O. sativa* | Os04g0644900 | Chr4 | 32830756 | 32831401 |
| *O. alta* | g26454 | Chr4CC | 35077156 | 35089800 |
| *O. alta* | g43645 | Chr7CC | 35920768 | 35933897 |
| *O. sativa* | Os04g0645100 | Chr4 | 32835280 | 32848285 |
| *O. alta* | g26455 | Chr4CC | 35092610 | 35093215 |
| *O. alta* | g43646 | Chr7CC | 35940092 | 35940709 |
| *O. sativa* | Os04g0645200 | Chr4 | 32852142 | 32853025 |
| *O. alta* | g26456 | Chr4CC | 35097028 | 35097795 |
| *O. alta* | g43647 | Chr7CC | 35943941 | 35944696 |
| *O. sativa* | Os04g0645500 | Chr4 | 32855657 | 32856649 |
| *O. alta* | g26457 | Chr4CC | 35098218 | 35101332 |
| *O. alta* | g43648 | Chr7CC | 35945128 | 35948265 |
| *O. sativa* | Os04g0645600 | Chr4 | 32856597 | 32860277 |

| | | | | |
|---|---|---|---|---|
| *O. alta* | g26458 | Chr4CC | 35106141 | 35109346 |
| *O. alta* | g43649 | Chr7CC | 35954673 | 35957876 |
| *O. sativa* | Os04g0647300 | Chr4 | 32933242 | 32937209 |
| *O. alta* | g26462 | Chr4CC | 35175173 | 35179413 |
| *O. alta* | g43650 | Chr7CC | 35981310 | 35985342 |
| *O. sativa* | Os04g0647800 | Chr4 | 32956000 | 32959941 |
| *O. alta* | g26465 | Chr4CC | 35193721 | 35204491 |
| *O. alta* | g43651 | Chr7CC | 35993542 | 35994881 |
| *O. sativa* | Os04g0647900 | Chr4 | 32963581 | 32967002 |
| *O. alta* | g26466 | Chr4CC | 35209733 | 35215948 |
| *O. alta* | g43653 | Chr7CC | 36014742 | 36019238 |
| *O. sativa* | Os04g0648500 | Chr4 | 32994714 | 32999780 |
| *O. alta* | g26467 | Chr4CC | 35218033 | 35218833 |
| *O. alta* | g43654 | Chr7CC | 36021527 | 36022327 |
| *O. sativa* | Os04g0648600 | Chr4 | 33001948 | 33003003 |
| *O. alta* | g26469 | Chr4CC | 35243474 | 35247203 |
| *O. alta* | g43656 | Chr7CC | 36047708 | 36051509 |
| *O. sativa* | Os04g0648800 | Chr4 | 33022835 | 33028358 |
| *O. alta* | g26471 | Chr4CC | 35282088 | 35284707 |
| *O. alta* | g43658 | Chr7CC | 36080871 | 36083443 |
| *O. sativa* | Os04g0649100 | Chr4 | 33071622 | 33075371 |
| *O. alta* | g26472 | Chr4CC | 35293497 | 35297048 |
| *O. alta* | g43659 | Chr7CC | 36092356 | 36096458 |
| *O. sativa* | Os04g0649200 | Chr4 | 33080927 | 33086419 |
| *O. alta* | g26475 | Chr4CC | 35313876 | 35314741 |
| *O. alta* | g43661 | Chr7CC | 36120972 | 36121552 |
| *O. sativa* | Os04g0649400 | Chr4 | 33095273 | 33096554 |
| *O. alta* | g26476 | Chr4CC | 35322898 | 35323239 |
| *O. alta* | g43663 | Chr7CC | 36146978 | 36147310 |
| *O. sativa* | Os04g0649500 | Chr4 | 33101621 | 33102439 |
| *O. alta* | g26477 | Chr4CC | 35329787 | 35330071 |
| *O. alta* | g43664 | Chr7CC | 36151792 | 36152079 |
| *O. sativa* | Os04g0649600 | Chr4 | 33107761 | 33108336 |
| *O. alta* | g26478 | Chr4CC | 35331719 | 35334833 |
| *O. alta* | g43665 | Chr7CC | 36153806 | 36156896 |
| *O. sativa* | Os04g0649700 | Chr4 | 33109718 | 33113326 |

| | | | | |
|---|---|---|---|---|
| *O. alta* | g26479 | Chr4CC | 35350272 | 35351228 |
| *O. alta* | g43667 | Chr7CC | 36163703 | 36164659 |
| *O. sativa* | Os04g0649900 | Chr4 | 33118368 | 33119696 |
| *O. alta* | g26480 | Chr4CC | 35357436 | 35360733 |
| *O. alta* | g43668 | Chr7CC | 36171224 | 36174511 |
| *O. sativa* | Os04g0650000 | Chr4 | 33123327 | 33126987 |
| *O. alta* | g26482 | Chr4CC | 35372602 | 35375081 |
| *O. alta* | g43669 | Chr7CC | 36191018 | 36196210 |
| *O. sativa* | Os04g0650366 | Chr4 | 33138559 | 33140012 |
| *O. alta* | g26484 | Chr4CC | 35382950 | 35384288 |
| *O. alta* | g43671 | Chr7CC | 36208240 | 36209197 |
| *O. sativa* | Os04g0650500 | Chr4 | 33147439 | 33150039 |
| *O. alta* | g26483 | Chr4CC | 35380650 | 35382067 |
| *O. alta* | g43670 | Chr7CC | 36206323 | 36207700 |
| *O. alta* | g26485 | Chr4CC | 35385635 | 35388990 |
| *O. alta* | g43672 | Chr7CC | 36211031 | 36213833 |
| *O. sativa* | Os04g0650600 | Chr4 | 33150022 | 33153292 |
| *O. alta* | g26487 | Chr4CC | 35398899 | 35401961 |
| *O. alta* | g43674 | Chr7CC | 36229522 | 36232530 |
| *O. sativa* | Os04g0650700 | Chr4 | 33161539 | 33164512 |
| *O. alta* | g26488 | Chr4CC | 35404950 | 35407546 |
| *O. alta* | g43675 | Chr7CC | 36234908 | 36237481 |
| *O. sativa* | Os04g0650800 | Chr4 | 33167054 | 33169881 |
| *O. alta* | g26490 | Chr4CC | 35417928 | 35419376 |
| *O. alta* | g43677 | Chr7CC | 36248462 | 36249869 |
| *O. sativa* | Os04g0651000 | Chr4 | 33179647 | 33181417 |
| *O. alta* | g26492 | Chr4CC | 35431862 | 35432968 |
| *O. alta* | g43678 | Chr7CC | 36266992 | 36267471 |
| *O. alta* | g26495 | Chr4CC | 35462147 | 35466706 |
| *O. alta* | g43680 | Chr7CC | 36302874 | 36307930 |
| *O. sativa* | Os04g0652400 | Chr4 | 33215040 | 33220582 |
| *O. alta* | g26497 | Chr4CC | 35472925 | 35479778 |
| *O. alta* | g43681 | Chr7CC | 36323611 | 36330694 |
| *O. sativa* | Os04g0652500 | Chr4 | 33225809 | 33230258 |

| | | | | |
|---|---|---|---|---|
| *O. alta* | g26499 | Chr4CC | 35491856 | 35494336 |
| *O. alta* | g43682 | Chr7CC | 36345159 | 36347494 |
| *O. sativa* | Os04g0652700 | Chr4 | 33253920 | 33256572 |
| *O. alta* | g26500 | Chr4CC | 35495321 | 35497774 |
| *O. alta* | g43683 | Chr7CC | 36348875 | 36350821 |
| *O. sativa* | Os04g0652900 | Chr4 | 33258681 | 33261192 |
| *O. alta* | g26501 | Chr4CC | 35513212 | 35516957 |
| *O. alta* | g43685 | Chr7CC | 36383873 | 36384845 |
| *O. sativa* | Os04g0653000 | Chr4 | 33306468 | 33310169 |
| *O. alta* | g26504 | Chr4CC | 35521339 | 35524727 |
| *O. alta* | g43687 | Chr7CC | 36389257 | 36392721 |
| *O. sativa* | Os04g0653200 | Chr4 | 33314700 | 33319036 |
| *O. alta* | g26505 | Chr4CC | 35529120 | 35529554 |
| *O. alta* | g43688 | Chr7CC | 36403001 | 36403432 |
| *O. sativa* | Os04g0653300 | Chr4 | 33325306 | 33325943 |
| *O. alta* | g26506 | Chr4CC | 35533185 | 35538808 |
| *O. alta* | g43689 | Chr7CC | 36406943 | 36412675 |
| *O. sativa* | Os04g0653400 | Chr4 | 33329793 | 33335864 |
| *O. alta* | g26507 | Chr4CC | 35548289 | 35552075 |
| *O. alta* | g43690 | Chr7CC | 36422477 | 36426230 |
| *O. sativa* | Os04g0653600 | Chr4 | 33341978 | 33346528 |
| *O. alta* | g26509 | Chr4CC | 35558155 | 35558732 |
| *O. alta* | g43692 | Chr7CC | 36435808 | 36436110 |
| *O. alta* | g26512 | Chr4CC | 35634339 | 35640024 |
| *O. alta* | g43695 | Chr7CC | 36459009 | 36466882 |
| *O. sativa* | Os04g0654600 | Chr4 | 33386622 | 33391644 |
| *O. alta* | g26513 | Chr4CC | 35644352 | 35647246 |
| *O. alta* | g43696 | Chr7CC | 36471656 | 36474420 |
| *O. sativa* | Os04g0654700 | Chr4 | 33394485 | 33397972 |
| *O. alta* | g26515 | Chr4CC | 35662390 | 35664858 |
| *O. alta* | g43697 | Chr7CC | 36477320 | 36479767 |
| *O. sativa* | Os04g0654800 | Chr4 | 33399564 | 33402242 |
| *O. alta* | g26516 | Chr4CC | 35670551 | 35672986 |
| *O. alta* | g43698 | Chr7CC | 36494751 | 36497183 |
| *O. sativa* | Os04g0655000 | Chr4 | 33407358 | 33409844 |

| | | | | |
|---|---|---|---|---|
| *O. alta* | g26517 | Chr4CC | 35711182 | 35711631 |
| *O. alta* | g43699 | Chr7CC | 36503588 | 36504052 |
| *O. alta* | g26518 | Chr4CC | 35733343 | 35734757 |
| *O. alta* | g43700 | Chr7CC | 36515373 | 36517887 |
| *O. sativa* | Os04g0655300 | Chr4 | 33418637 | 33421199 |
| *O. alta* | g26519 | Chr4CC | 35735332 | 35736486 |
| *O. alta* | g43701 | Chr7CC | 36518385 | 36519792 |
| *O. alta* | g26521 | Chr4CC | 35752446 | 35755036 |
| *O. alta* | g43702 | Chr7CC | 36530142 | 36531344 |
| *O. alta* | g26522 | Chr4CC | 35755978 | 35762605 |
| *O. alta* | g43703 | Chr7CC | 36532412 | 36538787 |
| *O. sativa* | Os04g0655600 | Chr4 | 33429129 | 33436181 |
| *O. alta* | g26526 | Chr4CC | 35788725 | 35794933 |
| *O. alta* | g43705 | Chr7CC | 36563801 | 36570006 |
| *O. sativa* | Os04g0656100 | Chr4 | 33457745 | 33464852 |
| *O. alta* | g26527 | Chr4CC | 35821945 | 35825347 |
| *O. alta* | g43706 | Chr7CC | 36594556 | 36597867 |
| *O. sativa* | Os04g0656500 | Chr4 | 33488722 | 33492700 |
| *O. alta* | g26528 | Chr4CC | 35837067 | 35841646 |
| *O. alta* | g43707 | Chr7CC | 36603486 | 36607001 |
| *O. sativa* | Os04g0656800 | Chr4 | 33500008 | 33506112 |
| *O. alta* | g26529 | Chr4CC | 35851370 | 35854000 |
| *O. alta* | g43709 | Chr7CC | 36615510 | 36618212 |
| *O. sativa* | Os04g0657100 | Chr4 | 33517229 | 33520052 |
| *O. alta* | g26530 | Chr4CC | 35856880 | 35860346 |
| *O. alta* | g43710 | Chr7CC | 36621186 | 36627326 |
| *O. alta* | g26531 | Chr4CC | 35861882 | 35864534 |
| *O. alta* | g43711 | Chr7CC | 36629079 | 36631661 |
| *O. sativa* | Os04g0657500 | Chr4 | 33530289 | 33533734 |
| *O. alta* | g26533 | Chr4CC | 35887545 | 35887849 |
| *O. alta* | g43712 | Chr7CC | 36659265 | 36659555 |
| *O. sativa* | Os04g0657900 | Chr4 | 33554837 | 33555369 |
| *O. alta* | g26534 | Chr4CC | 35891006 | 35895605 |
| *O. alta* | g43713 | Chr7CC | 36663501 | 36668082 |
| *O. sativa* | Os04g0658000 | Chr4 | 33557486 | 33563976 |

| | | | | |
|---|---|---|---|---|
| *O. alta* | g26535 | Chr4CC | 35899662 | 35900948 |
| *O. alta* | g43714 | Chr7CC | 36680023 | 36681303 |
| *O. sativa* | Os04g0658100 | Chr4 | 33565812 | 33568359 |
| *O. alta* | g26537 | Chr4CC | 35918857 | 35920355 |
| *O. alta* | g43715 | Chr7CC | 36683474 | 36696190 |
| *O. sativa* | Os04g0658300 | Chr4 | 33575152 | 33579653 |
| *O. alta* | g26541 | Chr4CC | 35936859 | 35937645 |
| *O. alta* | g43718 | Chr7CC | 36707169 | 36708020 |
| *O. alta* | g26542 | Chr4CC | 35940636 | 35942309 |
| *O. alta* | g43719 | Chr7CC | 36709388 | 36711055 |
| *O. sativa* | Os04g0658600 | Chr4 | 33589945 | 33591841 |
| *O. alta* | g26544 | Chr4CC | 35944531 | 35946320 |
| *O. alta* | g43720 | Chr7CC | 36713836 | 36715596 |
| *O. sativa* | Os04g0658700 | Chr4 | 33594301 | 33596091 |
| *O. alta* | g26545 | Chr4CC | 35959456 | 35959734 |
| *O. alta* | g43722 | Chr7CC | 36733447 | 36733719 |
| *O. sativa* | Os04g0658800 | Chr4 | 33604376 | 33605201 |
| *O. alta* | g26546 | Chr4CC | 35967070 | 35971549 |
| *O. alta* | g43723 | Chr7CC | 36745663 | 36746246 |
| *O. sativa* | Os04g0659000 | Chr4 | 33623137 | 33623920 |
| *O. alta* | g26547 | Chr4CC | 35972425 | 35976037 |
| *O. alta* | g43724 | Chr7CC | 36747115 | 36750702 |
| *O. sativa* | Os04g0659100 | Chr4 | 33624428 | 33631025 |
| *O. alta* | g26548 | Chr4CC | 35977181 | 35979009 |
| *O. alta* | g43725 | Chr7CC | 36751770 | 36754566 |
| *O. sativa* | Os04g0659150 | Chr4 | 33631897 | 33634738 |
| *O. alta* | g26551 | Chr4CC | 35988224 | 35988997 |
| *O. alta* | g43729 | Chr7CC | 36776862 | 36777638 |
| *O. sativa* | Os04g0659300 | Chr4 | 33641298 | 33642277 |
| *O. alta* | g26552 | Chr4CC | 35990953 | 35994572 |
| *O. alta* | g43730 | Chr7CC | 36781702 | 36785493 |
| *O. sativa* | Os04g0659500 | Chr4 | 33649693 | 33653854 |
| *O. alta* | g26553 | Chr4CC | 35996951 | 35998010 |
| *O. alta* | g43731 | Chr7CC | 36788496 | 36789433 |

| | | | | |
|---|---|---|---|---|
| *O. alta* | g26554 | Chr4CC | 36002272 | 36004333 |
| *O. alta* | g43732 | Chr7CC | 36791775 | 36793824 |
| *O. sativa* | Os04g0659800 | Chr4 | 33658425 | 33664069 |
| *O. alta* | g26555 | Chr4CC | 36007084 | 36014132 |
| *O. alta* | g43734 | Chr7CC | 36796107 | 36802210 |
| *O. sativa* | Os04g0659900 | Chr4 | 33665023 | 33672967 |
| *O. alta* | g26556 | Chr4CC | 36017920 | 36018234 |
| *O. alta* | g43735 | Chr7CC | 36806698 | 36807006 |
| *O. sativa* | Os04g0660000 | Chr4 | 33676329 | 33676925 |
| *O. alta* | g26557 | Chr4CC | 36024925 | 36025527 |
| *O. alta* | g43736 | Chr7CC | 36813468 | 36814091 |
| *O. sativa* | Os04g0660100 | Chr4 | 33676818 | 33683540 |
| *O. alta* | g26573 | Chr4CC | 36200720 | 36201899 |
| *O. alta* | g43753 | Chr7CC | 36966548 | 36967601 |
| *O. sativa* | Os04g0661800 | Chr4 | 33774455 | 33777293 |
| *O. alta* | g26558 | Chr4CC | 36065049 | 36069718 |
| *O. alta* | g43738 | Chr7CC | 36835272 | 36839995 |
| *O. sativa* | Os04g0660200 | Chr4 | 33697886 | 33703610 |
| *O. alta* | g26559 | Chr4CC | 36072441 | 36073340 |
| *O. alta* | g43739 | Chr7CC | 36846589 | 36847791 |
| *O. alta* | g26560 | Chr4CC | 36073374 | 36074417 |
| *O. alta* | g43740 | Chr7CC | 36847831 | 36848874 |
| *O. sativa* | Os04g0660400 | Chr4 | 33704890 | 33707242 |
| *O. alta* | g26561 | Chr4CC | 36075203 | 36086262 |
| *O. alta* | g43741 | Chr7CC | 36849639 | 36860782 |
| *O. sativa* | Os04g0660500 | Chr4 | 33707455 | 33719236 |
| *O. alta* | g26562 | Chr4CC | 36113389 | 36116265 |
| *O. alta* | g43744 | Chr7CC | 36912376 | 36919913 |
| *O. sativa* | Os04g0660900 | Chr4 | 33729812 | 33732506 |
| *O. alta* | g26565 | Chr4CC | 36132968 | 36133488 |
| *O. alta* | g43746 | Chr7CC | 36921221 | 36922939 |
| *O. sativa* | Os04g0661100 | Chr4 | 33733561 | 33734332 |
| *O. alta* | g26567 | Chr4CC | 36139949 | 36145997 |
| *O. alta* | g43747 | Chr7CC | 36937582 | 36943597 |
| *O. sativa* | Os04g0661300 | Chr4 | 33740412 | 33746973 |

| | | | | |
|---|---|---|---|---|
| *O. alta* | g26568 | Chr4CC | 36149192 | 36149486 |
| *O. alta* | g43749 | Chr7CC | 36947014 | 36947314 |
| *O. alta* | g26570 | Chr4CC | 36180445 | 36184253 |
| *O. alta* | g43750 | Chr7CC | 36952485 | 36956332 |
| *O. sativa* | Os04g0661600 | Chr4 | 33755461 | 33764473 |
| *O. alta* | g26572 | Chr4CC | 36187784 | 36194893 |
| *O. alta* | g43752 | Chr7CC | 36959826 | 36965899 |
| *O. sativa* | Os04g0661700 | Chr4 | 33765000 | 33772389 |
| *O. alta* | g26574 | Chr4CC | 36203670 | 36207406 |
| *O. alta* | g43754 | Chr7CC | 36968755 | 36971894 |
| *O. sativa* | Os04g0661900 | Chr4 | 33777470 | 33781125 |
| *O. alta* | g26576 | Chr4CC | 36211482 | 36213220 |
| *O. alta* | g43756 | Chr7CC | 36975384 | 36977082 |
| *O. sativa* | Os04g0662100 | Chr4 | 33784400 | 33786414 |
| *O. alta* | g26577 | Chr4CC | 36216545 | 36216988 |
| *O. alta* | g43759 | Chr7CC | 36981453 | 36981896 |
| *O. sativa* | Os04g0662200 | Chr4 | 33789435 | 33790386 |
| *O. alta* | g26578 | Chr4CC | 36230158 | 36230634 |
| *O. alta* | g43760 | Chr7CC | 37027944 | 37028420 |
| *O. sativa* | Os04g0662400 | Chr4 | 33804424 | 33805473 |
| *O. alta* | g26579 | Chr4CC | 36238114 | 36240745 |
| *O. alta* | g43761 | Chr7CC | 37035829 | 37037961 |
| *O. alta* | g26580 | Chr4CC | 36243800 | 36245826 |
| *O. alta* | g43762 | Chr7CC | 37040027 | 37043860 |
| *O. sativa* | Os04g0662700 | Chr4 | 33814158 | 33818739 |
| *O. alta* | g26582 | Chr4CC | 36264993 | 36270716 |
| *O. alta* | g43763 | Chr7CC | 37050551 | 37056303 |
| *O. sativa* | Os04g0662800 | Chr4 | 33822434 | 33828560 |
| *O. alta* | g26583 | Chr4CC | 36271644 | 36273625 |
| *O. alta* | g43764 | Chr7CC | 37058070 | 37060250 |
| *O. sativa* | Os04g0662900 | Chr4 | 33828951 | 33832503 |
| *O. alta* | g26585 | Chr4CC | 36285998 | 36290820 |
| *O. alta* | g43765 | Chr7CC | 37074448 | 37079380 |
| *O. sativa* | Os04g0663100 | Chr4 | 33840011 | 33845901 |
| *O. alta* | g26586 | Chr4CC | 36294531 | 36299273 |

| | | | | |
|---|---|---|---|---|
| *O. alta* | g43766 | Chr7CC | 37094691 | 37099341 |
| *O. sativa* | Os04g0663200 | Chr4 | 33848655 | 33850175 |
| *O. alta* | g26587 | Chr4CC | 36301125 | 36301457 |
| *O. alta* | g43767 | Chr7CC | 37100901 | 37101233 |
| *O. sativa* | Os04g0663300 | Chr4 | 33854490 | 33856500 |
| *O. alta* | g26588 | Chr4CC | 36302715 | 36303032 |
| *O. alta* | g43768 | Chr7CC | 37102620 | 37103021 |
| *O. alta* | g26589 | Chr4CC | 36303133 | 36304989 |
| *O. alta* | g43769 | Chr7CC | 37103092 | 37103409 |
| *O. sativa* | Os04g0663500 | Chr4 | 33856586 | 33859127 |
| *O. alta* | g26590 | Chr4CC | 36306994 | 36308052 |
| *O. alta* | g43770 | Chr7CC | 37110001 | 37111028 |
| *O. sativa* | Os04g0663600 | Chr4 | 33860374 | 33861424 |
| *O. alta* | g26591 | Chr4CC | 36320475 | 36323647 |
| *O. alta* | g43771 | Chr7CC | 37121918 | 37125005 |
| *O. sativa* | Os04g0663700 | Chr4 | 33869085 | 33872898 |
| *O. alta* | g26592 | Chr4CC | 36324390 | 36328506 |
| *O. alta* | g43772 | Chr7CC | 37125772 | 37129236 |
| *O. sativa* | Os04g0663800 | Chr4 | 33873137 | 33875586 |
| *O. alta* | g26593 | Chr4CC | 36328749 | 36329000 |
| *O. alta* | g43773 | Chr7CC | 37129309 | 37129644 |
| *O. alta* | g26598 | Chr4CC | 36375904 | 36377247 |
| *O. alta* | g43777 | Chr7CC | 37156169 | 37157650 |
| *O. alta* | g26602 | Chr4CC | 36397446 | 36399783 |
| *O. alta* | g43780 | Chr7CC | 37167812 | 37173103 |
| *O. sativa* | Os04g0664900 | Chr4 | 33943991 | 33946513 |
| *O. alta* | g26603 | Chr4CC | 36401124 | 36403017 |
| *O. alta* | g43781 | Chr7CC | 37174639 | 37175835 |
| *O. sativa* | Os04g0665000 | Chr4 | 33947318 | 33949741 |
| *O. alta* | g26604 | Chr4CC | 36403647 | 36407128 |
| *O. alta* | g43782 | Chr7CC | 37177172 | 37180557 |
| *O. sativa* | Os04g0665200 | Chr4 | 33950246 | 33952563 |
| *O. alta* | g26605 | Chr4CC | 36415315 | 36417961 |
| *O. alta* | g43783 | Chr7CC | 37202479 | 37205249 |
| *O. sativa* | Os04g0665400 | Chr4 | 33963664 | 33966703 |

| | | | | |
|---|---|---|---|---|
| *O. alta* | g26606 | Chr4CC | 36419805 | 36423138 |
| *O. alta* | g43784 | Chr7CC | 37206923 | 37210219 |
| *O. sativa* | Os04g0665500 | Chr4 | 33968605 | 33972525 |
| *O. alta* | g26607 | Chr4CC | 36424338 | 36426667 |
| *O. alta* | g43785 | Chr7CC | 37211401 | 37214082 |
| *O. sativa* | Os04g0665600 | Chr4 | 33972619 | 33976252 |
| *O. alta* | g26608 | Chr4CC | 36431937 | 36432572 |
| *O. alta* | g43786 | Chr7CC | 37233422 | 37234048 |
| *O. sativa* | Os04g0665666 | Chr4 | 33981899 | 33982687 |
| *O. alta* | g26609 | Chr4CC | 36436149 | 36438987 |
| *O. alta* | g43787 | Chr7CC | 37234811 | 37237757 |
| *O. sativa* | Os04g0665700 | Chr4 | 33982975 | 33986219 |
| *O. alta* | g26610 | Chr4CC | 36439606 | 36442442 |
| *O. alta* | g43788 | Chr7CC | 37238326 | 37241151 |
| *O. sativa* | Os04g0665800 | Chr4 | 33986389 | 33989930 |
| *O. alta* | g26611 | Chr4CC | 36443371 | 36443796 |
| *O. alta* | g43789 | Chr7CC | 37242060 | 37242500 |
| *O. sativa* | Os04g0665900 | Chr4 | 33990056 | 33990556 |
| *O. alta* | g26612 | Chr4CC | 36480035 | 36485437 |
| *O. alta* | g43792 | Chr7CC | 37265781 | 37266344 |
| *O. alta* | g26614 | Chr4CC | 36497062 | 36498166 |
| *O. alta* | g43795 | Chr7CC | 37289808 | 37291204 |
| *O. alta* | g26615 | Chr4CC | 36500920 | 36501530 |
| *O. alta* | g43796 | Chr7CC | 37295280 | 37295885 |
| *O. sativa* | Os04g0666800 | Chr4 | 34038375 | 34039430 |
| *O. alta* | g26616 | Chr4CC | 36502354 | 36511333 |
| *O. alta* | g43797 | Chr7CC | 37297816 | 37306917 |
| *O. sativa* | Os04g0666900 | Chr4 | 34040123 | 34048308 |
| *O. alta* | g26617 | Chr4CC | 36517001 | 36520883 |
| *O. alta* | g43798 | Chr7CC | 37318187 | 37322510 |
| *O. sativa* | Os04g0667000 | Chr4 | 34054318 | 34058149 |
| *O. alta* | g26619 | Chr4CC | 36535961 | 36538263 |
| *O. alta* | g43799 | Chr7CC | 37329888 | 37332309 |
| *O. sativa* | Os04g0667200 | Chr4 | 34060040 | 34063474 |

| | | | | |
|---|---|---|---|---|
| *O. alta* | g26620 | Chr4CC | 36544141 | 36545504 |
| *O. alta* | g43801 | Chr7CC | 37339476 | 37340827 |
| *O. sativa* | Os04g0667400 | Chr4 | 34067654 | 34070800 |
| *O. alta* | g26621 | Chr4CC | 36558241 | 36560236 |
| *O. alta* | g43802 | Chr7CC | 37345746 | 37348343 |
| *O. sativa* | Os04g0667500 | Chr4 | 34073167 | 34077291 |
| *O. alta* | g26622 | Chr4CC | 36567928 | 36568660 |
| *O. alta* | g43803 | Chr7CC | 37354567 | 37355319 |
| *O. sativa* | Os04g0667600 | Chr4 | 34081143 | 34081905 |
| *O. alta* | g26623 | Chr4CC | 36569935 | 36573634 |
| *O. alta* | g43804 | Chr7CC | 37357385 | 37361384 |
| *O. sativa* | Os04g0667700 | Chr4 | 34083436 | 34087516 |
| *O. alta* | g26624 | Chr4CC | 36574766 | 36576649 |
| *O. alta* | g43805 | Chr7CC | 37362581 | 37364516 |
| *O. sativa* | Os04g0667800 | Chr4 | 34088171 | 34090534 |
| *O. alta* | g26626 | Chr4CC | 36592766 | 36592975 |
| *O. alta* | g43807 | Chr7CC | 37394940 | 37395149 |
| *O. alta* | g26627 | Chr4CC | 36594487 | 36595734 |
| *O. alta* | g43808 | Chr7CC | 37398086 | 37399333 |
| *O. sativa* | Os04g0668600 | Chr4 | 34110319 | 34112130 |
| *O. alta* | g26628 | Chr4CC | 36598454 | 36600334 |
| *O. alta* | g43809 | Chr7CC | 37402200 | 37404074 |
| *O. sativa* | Os04g0668700 | Chr4 | 34114249 | 34116898 |
| *O. alta* | g26629 | Chr4CC | 36620376 | 36623650 |
| *O. alta* | g43810 | Chr7CC | 37409033 | 37412606 |
| *O. sativa* | Os04g0668800 | Chr4 | 34119501 | 34122972 |
| *O. alta* | g26630 | Chr4CC | 36624300 | 36626853 |
| *O. alta* | g43811 | Chr7CC | 37413224 | 37415699 |
| *O. sativa* | Os04g0668900 | Chr4 | 34122909 | 34125915 |
| *O. alta* | g26631 | Chr4CC | 36629597 | 36630782 |
| *O. alta* | g43812 | Chr7CC | 37418690 | 37419571 |
| *O. sativa* | Os04g0669100 | Chr4 | 34128105 | 34131134 |
| *O. alta* | g26632 | Chr4CC | 36634221 | 36634766 |
| *O. alta* | g43813 | Chr7CC | 37427720 | 37428250 |

| | | | | |
|---|---|---|---|---|
| *O. alta* | g26633 | Chr4CC | 36637362 | 36640624 |
| *O. alta* | g43814 | Chr7CC | 37431445 | 37434908 |
| *O. sativa* | Os04g0669300 | Chr4 | 34138228 | 34142138 |
| *O. alta* | g26634 | Chr4CC | 36641552 | 36642220 |
| *O. alta* | g43815 | Chr7CC | 37435852 | 37436532 |
| *O. sativa* | Os04g0669475 | Chr4 | 34142529 | 34143723 |
| *O. alta* | g26635 | Chr4CC | 36646444 | 36648317 |
| *O. alta* | g43816 | Chr7CC | 37441452 | 37443425 |
| *O. sativa* | Os04g0669500 | Chr4 | 34148182 | 34150717 |
| *O. alta* | g26636 | Chr4CC | 36648678 | 36650096 |
| *O. alta* | g43817 | Chr7CC | 37445241 | 37446575 |
| *O. sativa* | Os04g0669600 | Chr4 | 34150913 | 34152431 |
| *O. alta* | g26638 | Chr4CC | 36654192 | 36656975 |
| *O. alta* | g43818 | Chr7CC | 37448649 | 37452237 |
| *O. sativa* | Os04g0669800 | Chr4 | 34155536 | 34158649 |
| *O. alta* | g26639 | Chr4CC | 36703876 | 36706325 |
| *O. alta* | g43819 | Chr7CC | 37454337 | 37456835 |
| *O. sativa* | Os04g0670000 | Chr4 | 34166490 | 34169404 |
| *O. alta* | g26640 | Chr4CC | 36708386 | 36709756 |
| *O. alta* | g43820 | Chr7CC | 37459001 | 37460362 |
| *O. alta* | g26641 | Chr4CC | 36710286 | 36713343 |
| *O. alta* | g43821 | Chr7CC | 37460886 | 37463940 |
| *O. sativa* | Os04g0670200 | Chr4 | 34173356 | 34176732 |
| *O. alta* | g26642 | Chr4CC | 36717910 | 36724247 |
| *O. alta* | g43822 | Chr7CC | 37501555 | 37509975 |
| *O. sativa* | Os04g0670400 | Chr4 | 34201068 | 34205682 |
| *O. alta* | g26644 | Chr4CC | 36730274 | 36731665 |
| *O. alta* | g43823 | Chr7CC | 37511965 | 37513000 |
| *O. sativa* | Os04g0670500 | Chr4 | 34205994 | 34207906 |
| *O. alta* | g26646 | Chr4CC | 36735228 | 36737873 |
| *O. alta* | g43827 | Chr7CC | 37524537 | 37527214 |
| *O. sativa* | Os04g0670600 | Chr4 | 34209652 | 34214800 |
| *O. alta* | g26647 | Chr4CC | 36738299 | 36744148 |
| *O. alta* | g43830 | Chr7CC | 37544961 | 37550593 |
| *O. sativa* | Os04g0670800 | Chr4 | 34215822 | 34222368 |

| | | | | |
|---|---|---|---|---|
| *O. alta* | g26648 | Chr4CC | 36754725 | 36756525 |
| *O. alta* | g43831 | Chr7CC | 37559171 | 37560980 |
| *O. sativa* | Os04g0670900 | Chr4 | 34231186 | 34233221 |
| *O. alta* | g26649 | Chr4CC | 36786775 | 36788732 |
| *O. alta* | g43832 | Chr7CC | 37593932 | 37596421 |
| *O. sativa* | Os04g0671100 | Chr4 | 34236448 | 34237973 |
| *O. alta* | g26650 | Chr4CC | 36789800 | 36792091 |
| *O. alta* | g43833 | Chr7CC | 37597292 | 37599726 |
| *O. sativa* | Os04g0671200 | Chr4 | 34239804 | 34243087 |
| *O. alta* | g26651 | Chr4CC | 36794153 | 36797619 |
| *O. alta* | g43834 | Chr7CC | 37601924 | 37605422 |
| *O. sativa* | Os04g0671300 | Chr4 | 34244273 | 34248323 |
| *O. alta* | g26652 | Chr4CC | 36812007 | 36814865 |
| *O. alta* | g43835 | Chr7CC | 37614839 | 37617446 |
| *O. sativa* | Os04g0671700 | Chr4 | 34267955 | 34270846 |
| *O. alta* | g26653 | Chr4CC | 36816678 | 36820051 |
| *O. alta* | g43836 | Chr7CC | 37619008 | 37622165 |
| *O. sativa* | Os04g0671800 | Chr4 | 34272659 | 34276664 |
| *O. alta* | g26654 | Chr4CC | 36829658 | 36835809 |
| *O. alta* | g43837 | Chr7CC | 37624125 | 37630078 |
| *O. sativa* | Os04g0671900 | Chr4 | 34278477 | 34285845 |
| *O. alta* | g26656 | Chr4CC | 36870822 | 36874944 |
| *O. alta* | g43839 | Chr7CC | 37657743 | 37661630 |
| *O. sativa* | Os04g0672200 | Chr4 | 34314400 | 34320079 |
| *O. alta* | g26657 | Chr4CC | 36878153 | 36882022 |
| *O. alta* | g43840 | Chr7CC | 37667562 | 37669133 |
| *O. sativa* | Os04g0672300 | Chr4 | 34321350 | 34324747 |
| *O. alta* | g26655 | Chr4CC | 36856426 | 36859461 |
| *O. alta* | g43838 | Chr7CC | 37640364 | 37643399 |
| *O. sativa* | Os04g0672600 | Chr4 | 34329739 | 34332205 |
| *O. alta* | g26658 | Chr4CC | 36958229 | 36960403 |
| *O. alta* | g43842 | Chr7CC | 37689437 | 37690195 |
| *O. sativa* | Os04g0672700 | Chr4 | 34333498 | 34336668 |
| *O. alta* | g26659 | Chr4CC | 36976641 | 36979786 |
| *O. alta* | g43843 | Chr7CC | 37692460 | 37695604 |
| *O. sativa* | Os04g0672800 | Chr4 | 34338474 | 34341851 |

| | | | | |
|---|---|---|---|---|
| *O. alta* | g26660 | Chr4CC | 36981429 | 36983608 |
| *O. alta* | g43844 | Chr7CC | 37697137 | 37699369 |
| *O. sativa* | Os04g0672900 | Chr4 | 34342607 | 34345315 |
| *O. alta* | g26661 | Chr4CC | 36985768 | 36989627 |
| *O. alta* | g43845 | Chr7CC | 37701061 | 37705178 |
| *O. sativa* | Os04g0673000 | Chr4 | 34346555 | 34352326 |
| *O. alta* | g26663 | Chr4CC | 37014327 | 37014919 |
| *O. alta* | g43846 | Chr7CC | 37729425 | 37730016 |
| *O. sativa* | Os04g0673300 | Chr4 | 34375978 | 34377092 |
| *O. alta* | g26665 | Chr4CC | 37029798 | 37030436 |
| *O. alta* | g43848 | Chr7CC | 37758327 | 37758964 |
| *O. sativa* | Os04g0673800 | Chr4 | 34397149 | 34398174 |
| *O. alta* | g26667 | Chr4CC | 37032149 | 37033111 |
| *O. alta* | g43849 | Chr7CC | 37760791 | 37761753 |
| *O. sativa* | Os04g0674000 | Chr4 | 34400083 | 34401448 |
| *O. alta* | g26674 | Chr4CC | 37083520 | 37085058 |
| *O. alta* | g43850 | Chr7CC | 37778092 | 37780175 |
| *O. sativa* | Os04g0674100 | Chr4 | 34410429 | 34413185 |
| *O. alta* | g26675 | Chr4CC | 37085967 | 37087231 |
| *O. alta* | g43851 | Chr7CC | 37789911 | 37791156 |
| *O. sativa* | Os04g0674200 | Chr4 | 34413441 | 34415641 |
| *O. alta* | g26676 | Chr4CC | 37088485 | 37092721 |
| *O. alta* | g43852 | Chr7CC | 37793523 | 37797354 |
| *O. sativa* | Os04g0674300 | Chr4 | 34416120 | 34420309 |
| *O. alta* | g26677 | Chr4CC | 37098044 | 37099912 |
| *O. alta* | g43853 | Chr7CC | 37800921 | 37802870 |
| *O. sativa* | Os04g0674400 | Chr4 | 34423647 | 34426231 |
| *O. alta* | g26678 | Chr4CC | 37101679 | 37104029 |
| *O. alta* | g43854 | Chr7CC | 37804546 | 37806679 |
| *O. sativa* | Os04g0674450 | Chr4 | 34427913 | 34431452 |
| *O. alta* | g26679 | Chr4CC | 37113062 | 37116291 |
| *O. alta* | g43855 | Chr7CC | 37841744 | 37845195 |
| *O. sativa* | Os04g0674600 | Chr4 | 34440331 | 34444073 |
| *O. alta* | g26680 | Chr4CC | 37125810 | 37127569 |
| *O. alta* | g43856 | Chr7CC | 37854972 | 37856765 |

| | | | | |
|---|---|---|---|---|
| *O. sativa* | Os04g0674700 | Chr4 | 34444832 | 34446814 |
| *O. alta* | g26681 | Chr4CC | 37143024 | 37145320 |
| *O. alta* | g43857 | Chr7CC | 37864477 | 37866771 |
| *O. sativa* | Os04g0674800 | Chr4 | 34450562 | 34454341 |
| *O. alta* | g26682 | Chr4CC | 37150348 | 37154702 |
| *O. alta* | g43858 | Chr7CC | 37869590 | 37874665 |
| *O. sativa* | Os04g0675000 | Chr4 | 34457067 | 34463729 |
| *O. alta* | g26683 | Chr4CC | 37160811 | 37179605 |
| *O. alta* | g43859 | Chr7CC | 37880777 | 37899726 |
| *O. sativa* | Os04g0675101 | Chr4 | 34468893 | 34476991 |
| *O. alta* | g26684 | Chr4CC | 37180286 | 37182059 |
| *O. alta* | g43860 | Chr7CC | 37900423 | 37902205 |
| *O. sativa* | Os04g0675400 | Chr4 | 34489930 | 34492036 |
| *O. alta* | g26685 | Chr4CC | 37182757 | 37186948 |
| *O. alta* | g43861 | Chr7CC | 37902913 | 37907133 |
| *O. sativa* | Os04g0675500 | Chr4 | 34492078 | 34496501 |
| *O. alta* | g26686 | Chr4CC | 37190009 | 37191622 |
| *O. alta* | g43862 | Chr7CC | 37910179 | 37911801 |
| *O. sativa* | Os04g0675600 | Chr4 | 34498550 | 34499627 |
| *O. alta* | g26691 | Chr4CC | 37206206 | 37210160 |
| *O. alta* | g43863 | Chr7CC | 37919013 | 37922963 |
| *O. sativa* | Os04g0675800 | Chr4 | 34505859 | 34513140 |
| *O. alta* | g26692 | Chr4CC | 37210664 | 37211950 |
| *O. alta* | g43864 | Chr7CC | 37923465 | 37924762 |
| *O. sativa* | Os04g0676100 | Chr4 | 34513602 | 34515261 |
| *O. alta* | g26693 | Chr4CC | 37212766 | 37214132 |
| *O. alta* | g43865 | Chr7CC | 37925550 | 37927230 |
| *O. sativa* | Os04g0676200 | Chr4 | 34515411 | 34517165 |
| *O. alta* | g26694 | Chr4CC | 37215724 | 37218861 |
| *O. alta* | g43866 | Chr7CC | 37929405 | 37932778 |
| *O. sativa* | Os04g0676300 | Chr4 | 34519564 | 34523544 |
| *O. alta* | g26695 | Chr4CC | 37233511 | 37234329 |
| *O. alta* | g43867 | Chr7CC | 37942600 | 37943421 |
| *O. sativa* | Os04g0676400 | Chr4 | 34533897 | 34536098 |
| *O. alta* | g26698 | Chr4CC | 37246223 | 37249861 |

| | | | | |
|---|---|---|---|---|
| *O. alta* | g43869 | Chr7CC | 37962349 | 37967169 |
| *O. sativa* | Os04g0676650 | Chr4 | 34548346 | 34553030 |
| *O. alta* | g26700 | Chr4CC | 37255902 | 37259598 |
| *O. alta* | g43870 | Chr7CC | 37971640 | 37974940 |
| *O. sativa* | Os04g0676700 | Chr4 | 34555255 | 34559555 |
| *O. alta* | g26701 | Chr4CC | 37262463 | 37266606 |
| *O. alta* | g43871 | Chr7CC | 37982421 | 37985940 |
| *O. alta* | g26702 | Chr4CC | 37268319 | 37269722 |
| *O. alta* | g43872 | Chr7CC | 37987967 | 37989382 |
| *O. sativa* | Os04g0677100 | Chr4 | 34586733 | 34588232 |
| *O. alta* | g26705 | Chr4CC | 37287285 | 37288052 |
| *O. alta* | g43877 | Chr7CC | 38001845 | 38002615 |
| *O. sativa* | Os04g0677300 | Chr4 | 34597648 | 34598755 |
| *O. alta* | g26706 | Chr4CC | 37291345 | 37294585 |
| *O. alta* | g43878 | Chr7CC | 38005147 | 38008757 |
| *O. sativa* | Os04g0677400 | Chr4 | 34601247 | 34604930 |
| *O. alta* | g26707 | Chr4CC | 37295613 | 37298654 |
| *O. alta* | g43879 | Chr7CC | 38009767 | 38012789 |
| *O. sativa* | Os04g0677500 | Chr4 | 34605435 | 34609072 |
| *O. alta* | g26708 | Chr4CC | 37311383 | 37311940 |
| *O. alta* | g43880 | Chr7CC | 38016408 | 38016953 |
| *O. sativa* | Os04g0677600 | Chr4 | 34612684 | 34613440 |
| *O. alta* | g26709 | Chr4CC | 37313032 | 37317670 |
| *O. alta* | g43881 | Chr7CC | 38018044 | 38022878 |
| *O. sativa* | Os04g0677700 | Chr4 | 34613664 | 34623117 |
| *O. alta* | g26710 | Chr4CC | 37325652 | 37330989 |
| *O. alta* | g43883 | Chr7CC | 38030593 | 38034645 |
| *O. sativa* | Os04g0677800 | Chr4 | 34626374 | 34631340 |
| *O. alta* | g26711 | Chr4CC | 37331907 | 37332325 |
| *O. alta* | g43885 | Chr7CC | 38037716 | 38038112 |
| *O. alta* | g26712 | Chr4CC | 37337291 | 37338164 |
| *O. alta* | g43886 | Chr7CC | 38042980 | 38044122 |
| *O. sativa* | Os04g0678200 | Chr4 | 34642807 | 34644512 |
| *O. alta* | g26713 | Chr4CC | 37343961 | 37349156 |
| *O. alta* | g43887 | Chr7CC | 38049421 | 38054405 |

| | | | | |
|---|---|---|---|---|
| *O. sativa* | Os04g0678300 | Chr4 | 34649606 | 34655208 |
| *O. alta* | g26714 | Chr4CC | 37349849 | 37350583 |
| *O. alta* | g43888 | Chr7CC | 38055102 | 38055848 |
| *O. sativa* | Os04g0678400 | Chr4 | 34655316 | 34656735 |
| *O. alta* | g26715 | Chr4CC | 37360277 | 37361600 |
| *O. alta* | g43889 | Chr7CC | 38063294 | 38064604 |
| *O. sativa* | Os04g0678700 | Chr4 | 34663055 | 34664559 |
| *O. alta* | g26717 | Chr4CC | 37366890 | 37369462 |
| *O. alta* | g43891 | Chr7CC | 38069573 | 38071816 |
| *O. sativa* | Os04g0679000 | Chr4 | 34669394 | 34671829 |
| *O. alta* | g26718 | Chr4CC | 37370313 | 37372549 |
| *O. alta* | g43892 | Chr7CC | 38072676 | 38074656 |
| *O. sativa* | Os04g0679050 | Chr4 | 34672842 | 34676866 |
| *O. alta* | g26719 | Chr4CC | 37413942 | 37417404 |
| *O. alta* | g43893 | Chr7CC | 38102596 | 38105481 |
| *O. sativa* | Os04g0679100 | Chr4 | 34677521 | 34681308 |
| *O. alta* | g26720 | Chr4CC | 37418050 | 37419876 |
| *O. alta* | g43894 | Chr7CC | 38106117 | 38108134 |
| *O. sativa* | Os04g0679200 | Chr4 | 34681372 | 34685068 |
| *O. alta* | g26724 | Chr4CC | 37445084 | 37447268 |
| *O. alta* | g43896 | Chr7CC | 38130081 | 38132131 |
| *O. alta* | g26725 | Chr4CC | 37448350 | 37451092 |
| *O. alta* | g43897 | Chr7CC | 38133525 | 38136259 |
| *O. sativa* | Os04g0679900 | Chr4 | 34717887 | 34721250 |
| *O. alta* | g26726 | Chr4CC | 37455389 | 37460579 |
| *O. alta* | g43898 | Chr7CC | 38144681 | 38149616 |
| *O. sativa* | Os04g0680000 | Chr4 | 34725156 | 34730395 |
| *O. alta* | g26727 | Chr4CC | 37461167 | 37463237 |
| *O. alta* | g43899 | Chr7CC | 38167139 | 38169234 |
| *O. sativa* | Os04g0680300 | Chr4 | 34733807 | 34736093 |
| *O. alta* | g26728 | Chr4CC | 37464869 | 37468812 |
| *O. alta* | g43900 | Chr7CC | 38171740 | 38175751 |
| *O. sativa* | Os04g0680400 | Chr4 | 34738022 | 34742715 |
| *O. alta* | g26731 | Chr4CC | 37511192 | 37515353 |
| *O. alta* | g43902 | Chr7CC | 38183952 | 38188144 |

| | | | | |
|---|---|---|---|---|
| *O. sativa* | Os04g0680700 | Chr4 | 34751498 | 34756030 |
| *O. alta* | g26732 | Chr4CC | 37518164 | 37523128 |
| *O. alta* | g43903 | Chr7CC | 38191676 | 38196512 |
| *O. sativa* | Os04g0680800 | Chr4 | 34761220 | 34763488 |
| *O. alta* | g26734 | Chr4CC | 37543840 | 37544229 |
| *O. alta* | g43907 | Chr7CC | 38205120 | 38205521 |
| *O. alta* | g26735 | Chr4CC | 37550266 | 37554460 |
| *O. alta* | g43911 | Chr7CC | 38214693 | 38218904 |
| *O. sativa* | Os04g0681600 | Chr4 | 34791847 | 34800725 |
| *O. alta* | g43913 | Chr7CC | 38225735 | 38226058 |
| *O. alta* | g47791 | Chr8CC | 20184525 | 20184857 |
| *O. alta* | g26736 | Chr4CC | 37556044 | 37558099 |
| *O. alta* | g43912 | Chr7CC | 38223313 | 38225090 |
| *O. sativa* | Os04g0681500 | Chr4 | 34787366 | 34790215 |
| *O. alta* | g26740 | Chr4CC | 37572867 | 37575530 |
| *O. alta* | g43915 | Chr7CC | 38237373 | 38240291 |
| *O. sativa* | Os04g0682000 | Chr4 | 34816665 | 34820086 |
| *O. alta* | g26741 | Chr4CC | 37577880 | 37579364 |
| *O. alta* | g43916 | Chr7CC | 38242179 | 38243657 |
| *O. sativa* | Os04g0682100 | Chr4 | 34824492 | 34826306 |
| *O. alta* | g26743 | Chr4CC | 37580403 | 37582806 |
| *O. alta* | g43917 | Chr7CC | 38244437 | 38247030 |
| *O. sativa* | Os04g0682300 | Chr4 | 34828250 | 34831200 |
| *O. alta* | g26744 | Chr4CC | 37583915 | 37588373 |
| *O. alta* | g43918 | Chr7CC | 38248416 | 38252093 |
| *O. sativa* | Os04g0682400 | Chr4 | 34832052 | 34837342 |
| *O. alta* | g26746 | Chr4CC | 37589488 | 37593089 |
| *O. alta* | g43919 | Chr7CC | 38254123 | 38256556 |
| *O. sativa* | Os04g0682500 | Chr4 | 34839497 | 34843336 |
| *O. alta* | g26747 | Chr4CC | 37597821 | 37614542 |
| *O. alta* | g43920 | Chr7CC | 38263113 | 38280714 |
| *O. sativa* | Os04g0682800 | Chr4 | 34860402 | 34869589 |
| *O. alta* | g26748 | Chr4CC | 37615517 | 37619631 |
| *O. alta* | g43921 | Chr7CC | 38281779 | 38284842 |
| *O. sativa* | Os04g0683100 | Chr4 | 34877803 | 34881555 |

| | | | | |
|---|---|---|---|---|
| *O. alta* | g26749 | Chr4CC | 37624796 | 37625170 |
| *O. alta* | g43922 | Chr7CC | 38305366 | 38305746 |
| *O. sativa* | Os04g0683400 | Chr4 | 34891819 | 34892196 |
| *O. alta* | g26750 | Chr4CC | 37625660 | 37627084 |
| *O. alta* | g43923 | Chr7CC | 38307413 | 38310287 |
| *O. sativa* | Os04g0683500 | Chr4 | 34892965 | 34898794 |
| *O. alta* | g26751 | Chr4CC | 37634547 | 37639473 |
| *O. alta* | g43924 | Chr7CC | 38316873 | 38321374 |
| *O. sativa* | Os04g0683600 | Chr4 | 34901628 | 34907296 |
| *O. alta* | g26752 | Chr4CC | 37642112 | 37643756 |
| *O. alta* | g43925 | Chr7CC | 38323529 | 38325177 |
| *O. sativa* | Os04g0683700 | Chr4 | 34908578 | 34910477 |
| *O. alta* | g26753 | Chr4CC | 37650409 | 37653438 |
| *O. alta* | g43926 | Chr7CC | 38329284 | 38332313 |
| *O. sativa* | Os04g0683800 | Chr4 | 34915815 | 34919982 |
| *O. alta* | g26755 | Chr4CC | 37656281 | 37658257 |
| *O. alta* | g43928 | Chr7CC | 38335741 | 38337813 |
| *O. sativa* | Os04g0683900 | Chr4 | 34922245 | 34926223 |
| *O. alta* | g26756 | Chr4CC | 37681550 | 37686972 |
| *O. alta* | g43930 | Chr7CC | 38386827 | 38392352 |
| *O. alta* | g26757 | Chr4CC | 37693261 | 37694135 |
| *O. alta* | g43931 | Chr7CC | 38394247 | 38395110 |
| *O. alta* | g26758 | Chr4CC | 37695690 | 37702666 |
| *O. alta* | g43932 | Chr7CC | 38395993 | 38403275 |
| *O. sativa* | Os04g0684500 | Chr4 | 34958612 | 34971379 |
| *O. alta* | g26759 | Chr4CC | 37706548 | 37706781 |
| *O. alta* | g43933 | Chr7CC | 38408464 | 38408703 |
| *O. sativa* | Os04g0684600 | Chr4 | 34972015 | 34972557 |
| *O. alta* | g26760 | Chr4CC | 37710881 | 37712318 |
| *O. alta* | g43935 | Chr7CC | 38414211 | 38415942 |
| *O. sativa* | Os04g0684800 | Chr4 | 34975665 | 34977774 |
| *O. alta* | g26761 | Chr4CC | 37714440 | 37715423 |
| *O. alta* | g43936 | Chr7CC | 38418065 | 38419003 |
| *O. sativa* | Os04g0684900 | Chr4 | 34980601 | 34981819 |

| | | | | |
|---|---|---|---|---|
| *O. alta* | g26762 | Chr4CC | 37721269 | 37722048 |
| *O. alta* | g43937 | Chr7CC | 38432518 | 38433291 |
| *O. sativa* | Os04g0685000 | Chr4 | 34988985 | 34989978 |
| *O. alta* | g26763 | Chr4CC | 37724445 | 37726822 |
| *O. alta* | g43938 | Chr7CC | 38437566 | 38439912 |
| *O. sativa* | Os04g0685100 | Chr4 | 34992137 | 34994799 |
| *O. alta* | g26764 | Chr4CC | 37729340 | 37730815 |
| *O. alta* | g43939 | Chr7CC | 38441459 | 38442925 |
| *O. sativa* | Os04g0685200 | Chr4 | 34997670 | 34998861 |
| *O. alta* | g26766 | Chr4CC | 37737136 | 37737753 |
| *O. alta* | g43940 | Chr7CC | 38447236 | 38447856 |
| *O. sativa* | Os04g0685300 | Chr4 | 35001018 | 35002142 |
| *O. alta* | g26768 | Chr4CC | 37748741 | 37752679 |
| *O. alta* | g43942 | Chr7CC | 38461822 | 38465833 |
| *O. sativa* | Os04g0685500 | Chr4 | 35013422 | 35020132 |
| *O. alta* | g26769 | Chr4CC | 37754113 | 37759541 |
| *O. alta* | g43943 | Chr7CC | 38469141 | 38474548 |
| *O. sativa* | Os04g0685600 | Chr4 | 35022041 | 35028128 |
| *O. alta* | g26770 | Chr4CC | 37763137 | 37763961 |
| *O. alta* | g43945 | Chr7CC | 38515124 | 38515954 |
| *O. sativa* | Os04g0685700 | Chr4 | 35031210 | 35032643 |
| *O. alta* | g26771 | Chr4CC | 37770358 | 37774214 |
| *O. alta* | g43946 | Chr7CC | 38521245 | 38522315 |
| *O. sativa* | Os04g0685800 | Chr4 | 35034889 | 35037788 |
| *O. alta* | g26772 | Chr4CC | 37776867 | 37780176 |
| *O. alta* | g43947 | Chr7CC | 38530754 | 38534043 |
| *O. sativa* | Os04g0685900 | Chr4 | 35039986 | 35043790 |
| *O. alta* | g26773 | Chr4CC | 37786414 | 37787085 |
| *O. alta* | g43948 | Chr7CC | 38540824 | 38543826 |
| *O. alta* | g26775 | Chr4CC | 37822612 | 37823844 |
| *O. alta* | g43949 | Chr7CC | 38553775 | 38555007 |
| *O. sativa* | Os04g0686000 | Chr4 | 35054571 | 35056161 |
| *O. alta* | g26776 | Chr4CC | 37831787 | 37832044 |
| *O. alta* | g43950 | Chr7CC | 38562337 | 38562603 |
| *O. sativa* | Os04g0686100 | Chr4 | 35064007 | 35064276 |

| | | | | |
|---|---|---|---|---|
| *O. alta* | g26777 | Chr4CC | 37835459 | 37842088 |
| *O. alta* | g43952 | Chr7CC | 38567425 | 38574001 |
| *O. sativa* | Os04g0686200 | Chr4 | 35067307 | 35074531 |
| *O. alta* | g26778 | Chr4CC | 37854579 | 37856342 |
| *O. alta* | g43953 | Chr7CC | 38583667 | 38585552 |
| *O. sativa* | Os04g0686300 | Chr4 | 35084318 | 35086242 |
| *O. alta* | g26779 | Chr4CC | 37857081 | 37859141 |
| *O. alta* | g43954 | Chr7CC | 38593988 | 38596057 |
| *O. sativa* | Os04g0686500 | Chr4 | 35086778 | 35089294 |
| *O. alta* | g26780 | Chr4CC | 37860316 | 37868535 |
| *O. alta* | g43956 | Chr7CC | 38598960 | 38604825 |
| *O. sativa* | Os04g0686650 | Chr4 | 35091588 | 35097920 |
| *O. alta* | g26781 | Chr4CC | 37875746 | 37876990 |
| *O. alta* | g43957 | Chr7CC | 38615747 | 38616985 |
| *O. sativa* | Os04g0686700 | Chr4 | 35105256 | 35106781 |
| *O. alta* | g26782 | Chr4CC | 37891757 | 37892371 |
| *O. alta* | g43958 | Chr7CC | 38646651 | 38647274 |
| *O. sativa* | Os04g0686800 | Chr4 | 35111460 | 35112512 |
| *O. alta* | g26783 | Chr4CC | 37896185 | 37898770 |
| *O. alta* | g43961 | Chr7CC | 38710474 | 38713025 |
| *O. sativa* | Os04g0687100 | Chr4 | 35120757 | 35124119 |
| *O. alta* | g26784 | Chr4CC | 37900490 | 37901183 |
| *O. alta* | g43962 | Chr7CC | 38714718 | 38715385 |
| *O. sativa* | Os04g0687200 | Chr4 | 35127153 | 35128399 |
| *O. alta* | g26785 | Chr4CC | 37916139 | 37919215 |
| *O. alta* | g43963 | Chr7CC | 38726508 | 38729246 |
| *O. sativa* | Os04g0687800 | Chr4 | 35167839 | 35170996 |
| *O. alta* | g26786 | Chr4CC | 37926927 | 37929764 |
| *O. alta* | g43964 | Chr7CC | 38743312 | 38746162 |
| *O. sativa* | Os04g0687900 | Chr4 | 35176157 | 35179785 |
| *O. alta* | g26787 | Chr4CC | 37933003 | 37934271 |
| *O. alta* | g43965 | Chr7CC | 38748412 | 38749689 |
| *O. sativa* | Os04g0688000 | Chr4 | 35182099 | 35183634 |
| *O. alta* | g26788 | Chr4CC | 37938159 | 37939440 |
| *O. alta* | g43966 | Chr7CC | 38756538 | 38757834 |
| *O. sativa* | Os04g0688100 | Chr4 | 35187443 | 35189187 |

| | | | | |
|---|---|---|---|---|
| *O. alta* | g26790 | Chr4CC | 37980273 | 37981391 |
| *O. alta* | g43970 | Chr7CC | 38776439 | 38777233 |
| *O. sativa* | Os04g0689000 | Chr4 | 35245424 | 35246736 |
| *O. alta* | g26793 | Chr4CC | 38004402 | 38007031 |
| *O. alta* | g43972 | Chr7CC | 38785113 | 38789758 |
| *O. sativa* | Os04g0689300 | Chr4 | 35267299 | 35270253 |
| *O. alta* | g26794 | Chr4CC | 38007860 | 38013484 |
| *O. alta* | g43973 | Chr7CC | 38790552 | 38795973 |
| *O. sativa* | Os04g0689400 | Chr4 | 35271050 | 35276776 |
| *O. alta* | g26795 | Chr4CC | 38014965 | 38016012 |
| *O. alta* | g43974 | Chr7CC | 38797719 | 38798774 |
| *O. sativa* | Os04g0689500 | Chr4 | 35277706 | 35279151 |
| *O. alta* | g26796 | Chr4CC | 38019944 | 38021127 |
| *O. alta* | g43975 | Chr7CC | 38800997 | 38803802 |
| *O. sativa* | Os04g0689700 | Chr4 | 35280703 | 35282692 |
| *O. alta* | g26799 | Chr4CC | 38057198 | 38058145 |
| *O. alta* | g43981 | Chr7CC | 38852936 | 38853761 |
| *O. sativa* | Os04g0689900 | Chr4 | 35287781 | 35289153 |
| *O. alta* | g26800 | Chr4CC | 38068590 | 38070884 |
| *O. alta* | g43982 | Chr7CC | 38913457 | 38916038 |
| *O. sativa* | Os04g0690100 | Chr4 | 35297676 | 35300310 |
| *O. alta* | g26801 | Chr4CC | 38072968 | 38078081 |
| *O. alta* | g43983 | Chr7CC | 38917393 | 38922486 |
| *O. sativa* | Os04g0690300 | Chr4 | 35310316 | 35315659 |
| *O. alta* | g26802 | Chr4CC | 38080014 | 38082113 |
| *O. alta* | g43984 | Chr7CC | 38924582 | 38926768 |
| *O. sativa* | Os04g0690400 | Chr4 | 35317098 | 35323984 |
| *O. alta* | g26803 | Chr4CC | 38088966 | 38089759 |
| *O. alta* | g43985 | Chr7CC | 38934291 | 38935014 |
| *O. sativa* | Os04g0690500 | Chr4 | 35327019 | 35328615 |
| *O. alta* | g26804 | Chr4CC | 38092092 | 38094103 |
| *O. alta* | g43986 | Chr7CC | 38938315 | 38940185 |
| *O. sativa* | Os04g0690600 | Chr4 | 35329565 | 35333095 |
| *O. alta* | g26805 | Chr4CC | 38099772 | 38100539 |
| *O. alta* | g43987 | Chr7CC | 38946947 | 38947693 |

| | | | | |
|---|---|---|---|---|
| *O. sativa* | Os04g0690800 | Chr4 | 35337067 | 35338179 |
| *O. alta* | g26806 | Chr4CC | 38114099 | 38115491 |
| *O. alta* | g43989 | Chr7CC | 38957453 | 38958846 |
| *O. sativa* | Os04g0691100 | Chr4 | 35344338 | 35346212 |
| *O. alta* | g26807 | Chr4CC | 38116517 | 38125005 |
| *O. alta* | g43990 | Chr7CC | 38959797 | 38968379 |
| *O. sativa* | Os04g0691200 | Chr4 | 35347095 | 35350031 |
| *O. alta* | g26808 | Chr4CC | 38132586 | 38133482 |
| *O. alta* | g43991 | Chr7CC | 38976185 | 38977135 |
| *O. sativa* | Os04g0691300 | Chr4 | 35358602 | 35362522 |
| *O. alta* | g26809 | Chr4CC | 38135227 | 38137407 |
| *O. alta* | g43992 | Chr7CC | 38982536 | 38990758 |
| *O. sativa* | Os04g0691400 | Chr4 | 35362185 | 35363363 |
| *O. alta* | g26810 | Chr4CC | 38148032 | 38159876 |
| *O. alta* | g43993 | Chr7CC | 39006861 | 39020456 |
| *O. sativa* | Os04g0691500 | Chr4 | 35369085 | 35381225 |
| *O. alta* | g26812 | Chr4CC | 38163818 | 38172714 |
| *O. alta* | g43994 | Chr7CC | 39024857 | 39031999 |
| *O. sativa* | Os04g0691700 | Chr4 | 35383000 | 35390423 |
| *O. alta* | g26813 | Chr4CC | 38200451 | 38203549 |
| *O. alta* | g43995 | Chr7CC | 39040136 | 39043237 |
| *O. sativa* | Os04g0691800 | Chr4 | 35398774 | 35402551 |
| *O. alta* | g26814 | Chr4CC | 38254763 | 38259625 |
| *O. alta* | g43999 | Chr7CC | 39097983 | 39102925 |
| *O. sativa* | Os04g0691900 | Chr4 | 35413301 | 35418842 |
| *O. alta* | g26815 | Chr4CC | 38260211 | 38262367 |
| *O. alta* | g44000 | Chr7CC | 39103532 | 39107591 |
| *O. sativa* | Os04g0692000 | Chr4 | 35418974 | 35421543 |
| *O. alta* | g26816 | Chr4CC | 38264008 | 38267795 |
| *O. alta* | g44001 | Chr7CC | 39111835 | 39112236 |
| *O. sativa* | Os04g0692100 | Chr4 | 35423055 | 35426811 |
| *O. alta* | g26817 | Chr4CC | 38267941 | 38270333 |
| *O. alta* | g44002 | Chr7CC | 39112465 | 39114794 |
| *O. sativa* | Os04g0692200 | Chr4 | 35426979 | 35429567 |
| *O. alta* | g26818 | Chr4CC | 38271680 | 38273686 |

| | | | | |
|---|---|---|---|---|
| *O. alta* | g44003 | Chr7CC | 39116527 | 39118484 |
| *O. sativa* | Os04g0692300 | Chr4 | 35430341 | 35432796 |
| *O. alta* | g26820 | Chr4CC | 38282688 | 38286553 |
| *O. alta* | g44004 | Chr7CC | 39139899 | 39149449 |
| *O. sativa* | Os04g0692500 | Chr4 | 35439339 | 35443769 |
| *O. alta* | g26823 | Chr4CC | 38298783 | 38306098 |
| *O. alta* | g44005 | Chr7CC | 39149927 | 39155168 |
| *O. sativa* | Os04g0692750 | Chr4 | 35452104 | 35459864 |
| *O. alta* | g26825 | Chr4CC | 38318115 | 38318330 |
| *O. alta* | g44009 | Chr7CC | 39194091 | 39194946 |
| *O. alta* | g26824 | Chr4CC | 38309929 | 38314780 |
| *O. alta* | g44007 | Chr7CC | 39184852 | 39189107 |
| *O. sativa* | Os04g0692800 | Chr4 | 35462597 | 35463100 |
| *O. alta* | g26827 | Chr4CC | 38319849 | 38321633 |
| *O. alta* | g44011 | Chr7CC | 39196602 | 39198409 |
| *O. sativa* | Os04g0693250 | Chr4 | 35493998 | 35497285 |
| *O. alta* | g26826 | Chr4CC | 38318999 | 38319677 |
| *O. alta* | g44010 | Chr7CC | 39195775 | 39196414 |
| *O. alta* | g26828 | Chr4CC | 38339869 | 38341743 |
| *O. alta* | g44013 | Chr7CC | 39200864 | 39202924 |
| *O. sativa* | Os04g0693300 | Chr4 | 35499332 | 35499760 |
| *O. alta* | g26829 | Chr4CC | 38352448 | 38356609 |
| *O. alta* | g44015 | Chr7CC | 39214101 | 39217063 |
| *O. alta* | g26830 | Chr4CC | 38357667 | 38363928 |
| *O. alta* | g44017 | Chr7CC | 39232010 | 39238333 |

**Supplementary Table 7.** GenBank accessions of chloroplast genomes used in the chloroplast-based phylogenetic

| | Species name | Genome type | GenBank Accession number | Reference (DOI or GenBank Link) |
|---|---|---|---|---|
| 1 | *O. barthii* | AA | NC_027460.1 | 10.1038/srep13957 |
| 2 | *O. glaberrima* | AA | NC_024175.1 | 10.1111/1755-0998.12258 |
| 3 | *O. nivara* | AA | OL912836 | 10.1080/23802359.2021.1977197 |
| 4 | *O. sativa japonica* | AA | NC_001320.1 | 10.1007/BF00336789 |
| 5 | *O. rufipogon* | AA | NC_017835.1 | 10.1002/ece3.66 |
| 6 | *O. sativa indica* | AA | NC_008155.1 | 10.1104/pp.103.031245 |
| 7 | *O. meridionalis* | AA | NC_016927.1 | 10.1002/ece3.66 |
| 8 | *O. glumipatula* | AA | NC_027461.1 | 10.1038/srep13957 |
| 9 | *O. longistaminata* | AA | NC_027462.1 | 10.1038/srep13957 |
| 10 | *O. punctata* | BB | KF359908 | KF359908 |
| **11** | ***O. malampuzhaensis*** | **BBCC** | **This study** | **-** |
| **12** | ***O. minuta*** | **BBCC** | **This study** | **-** |
| 13 | *O. eichingeri* | CC | NC_034759.1 | NC_034759.1 |
| 14 | *O. officinalis* | CC | KF359910 | KF359910 |
| 15 | *O. rhizomatis* | CC | NC_034758.1 | NC_034758.1 |
| **16** | ***O. alta*** | **CCDD** | **This study** | **-** |
| **17** | ***O. grandiglumis*** | **CCDD** | **This study** | **-** |
| **18** | ***O. latifolia*** | **CCDD** | **This study** | **-** |
| 19 | *O. australiensis* | EE | KF359916 | KF359916 |
| **20** | ***O. coarctata*** | **KKLL** | **This study** | **-** |
| **21** | ***O. schlecteri*** | **HHKK** | **This study** | **-** |
| **23** | ***O. ridleyi*** | **HHJJ** | **This study** | **-** |
| **24** | ***O. longiglumis*** | **HHJJ** | **This study** | **-** |
| 22 | *O. brachyantha* | FF | KF359917 | KF359917 |
| **25** | ***O. meyeriana*** | **GG** | **This study** | **-** |
| 26 | *O. neocaledonica* | GG | NC_053276.1 | 10.1007/s11103-020-01054-3 |
| 27 | *L. japonica* | Outgroup | KF359922.1 | KF359922.1 |

**Supplementary Table 8.** Average percentage of gene retention at the chromosome, subgenome, and genome level.

| Average % gene retention in chromosome | *O. malampuzhaensis* BB | *O. malampuzhaensis* CC | *O. minuta* BB | *O. minuta* CC | *O. alta* CC | *O. alta* DD | *O. grandiglumis* CC | *O. grandiglumis* DD | *O. latifolia* CC | *O. latifolia* DD | *O. coarctata* KK | *O. coarctata* LL | *O. schlechteri* HH | *O. schlechteri* KK | *O. longiglumis* HH | *O. longiglumis* JJ | *O. ridleyi* HH | *O. ridleyi* JJ |
|---|---|---|---|---|---|---|---|---|---|---|---|---|---|---|---|---|---|---|
| Chr1 | 54.32 | 56.07 | 53.93 | 57.15 | 61.12 | 61.59 | 62.23 | 62.19 | 57.54 | 57.80 | 56.68 | 59.24 | 50.68 | 54.37 | 57.66 | 52.87 | 53.99 | 48.94 |
| Chr2 | 56.14 | 58.01 | 56.46 | 59.18 | 62.90 | 60.99 | 62.66 | 60.94 | 57.56 | 58.15 | 60.14 | 61.72 | 51.48 | 54.83 | 60.03 | 55.05 | 58.73 | 52.20 |
| Chr3 | 58.57 | 63.47 | 59.24 | 63.10 | 63.45 | 61.07 | 62.30 | 61.20 | 63.63 | 60.53 | 62.69 | 64.08 | 53.30 | 57.79 | 65.23 | 55.94 | 61.62 | 53.37 |
| Chr4 | 52.42 | 54.43 | 53.56 | 55.99 | 56.19 | 56.80 | 55.76 | 56.22 | 56.21 | 53.34 | 55.64 | 58.12 | 46.73 | 49.97 | 57.25 | 49.26 | 54.50 | 48.19 |
| Chr5 | 55.31 | 56.88 | 55.42 | 57.51 | 59.24 | 60.43 | 58.92 | 59.91 | 54.24 | 52.96 | 54.83 | 56.93 | 48.10 | 52.29 | 59.35 | 52.32 | 58.41 | 51.11 |
| Chr6 | 50.80 | 52.75 | 51.42 | 54.41 | 54.50 | 54.97 | 54.19 | 53.74 | 51.61 | 51.04 | 54.05 | 55.74 | 45.14 | 45.55 | 54.67 | 48.37 | 53.27 | 47.41 |
| Chr7 | 45.73 | 47.94 | 47.00 | 48.30 | 50.65 | 49.97 | 49.63 | 49.70 | 47.31 | 45.23 | 50.43 | 52.32 | 40.65 | 41.40 | 51.31 | 46.78 | 48.60 | 44.53 |
| Chr8 | 47.61 | 49.82 | 48.43 | 50.82 | 56.33 | 52.00 | 55.64 | 51.85 | 51.83 | 46.84 | 53.23 | 54.62 | 44.83 | 43.14 | 52.92 | 47.19 | 50.45 | 45.17 |
| Chr9 | 49.23 | 48.98 | 52.36 | 51.88 | 57.42 | 54.93 | 56.13 | 56.27 | 52.28 | 49.45 | 53.71 | 55.47 | 44.06 | 46.28 | 48.77 | 46.60 | 48.43 | 45.01 |
| Chr10 | 40.53 | 46.90 | 41.74 | 48.41 | 47.67 | 47.90 | 46.69 | 47.45 | 44.55 | 42.57 | 49.67 | 49.15 | 36.03 | 39.83 | 44.39 | 38.30 | 45.04 | 38.33 |
| Chr11 | 37.72 | 41.90 | 38.17 | 43.57 | 47.50 | 41.88 | 44.62 | 41.27 | 42.10 | 35.86 | 44.64 | 44.99 | 31.64 | 35.28 | 42.78 | 35.85 | 40.99 | 35.29 |
| Chr12 | 45.81 | 48.50 | 46.54 | 48.89 | 55.24 | 50.59 | 51.64 | 49.15 | 46.98 | 45.01 | 48.55 | 53.62 | 39.39 | 40.42 | 49.94 | 44.38 | 49.95 | 44.05 |
| **Average % gene retention in subgenome** | **50.82** | **53.49** | **51.55** | **54.59** | **56.87** | **55.28** | **56.12** | **55.23** | **53.63** | **51.49** | **55.21** | **57.07** | **45.76** | **48.35** | **55.35** | **49.19** | **53.34** | **47.34** |
| **Average % gene retention in genome** | **52.12** | | **53.03** | | **56.07** | | **55.67** | | **52.54** | | **56.12** | | **47.02** | | **52.09** | | **50.15** | |

**Supplementary Table 9.** Number of genes and median transcript abundance as $\log_2$(TPM + 1) in the four categories of homoeologous genes of *O. coarctata* and the two categories of homologous genes of *O. sativa*, in the leaf and in the root.

| Sample | Category | Number of genes | Median $\log_2$(TPM + 1) Leaf | Wilcoxon rank sum test (Leaf) | Median $\log_2$(TPM + 1) Root | Wilcoxon rank sum test (Root) |
|---|---|---|---|---|---|---|
| All replicates | KK(hom) | 20,821 | 2.170 | 7.50E-148 | 2.910 | 2.50E-107 |
| All replicates | KK(lost) | 6,856 | 0.981 | | 1.883 | |
| All replicates | LL(hom) | 21,408 | 2.111 | 8.10E-123 | 3.203 | 7.20E-239 |
| All replicates | LL(lost) | 6,624 | 1.012 | | 1.660 | |
| All replicates | Os(hom) | 22,962 | 2.283 | 1.50E-42 | 2.668 | 1.70E-77 |
| All replicates | Os(lost) | 14,865 | 1.758 | | 2.015 | |

**Supplementary Table 10.** Number of genes expressed at least two-fold higher compared to their homoeologs in the other subgenome, and genes not dominantly expressed, in the leaf and in the root. Binomial test p-value is provided for each tissue.

| Log$_2$-fold change | Number of Genes in Leaf | | | | Number of Genes in Root | | | |
|---|---|---|---|---|---|---|---|---|
| | **KK is dominantly expressed** | **LL is dominantly expressed** | **not dominantly expressed** | **Grand Total** | **KK is dominantly expressed** | **LL is dominantly expressed** | **not dominantly expressed** | **Grand Total** |
| -13; -12 | 4 | | | 4 | 2 | | | 2 |
| -12; -11 | 1 | | | 1 | 2 | | | 2 |
| -11; -10 | 5 | | | 5 | 2 | | | 2 |
| -10; -9 | 6 | | | 6 | 2 | | | 2 |
| -9; -8 | 37 | | | 37 | 11 | | | 11 |
| -8; -7 | 53 | | | 53 | 41 | | | 41 |
| -7; -6 | 107 | | | 107 | 68 | | | 68 |
| -6; -5 | 205 | | | 205 | 146 | | | 146 |
| -5; -4 | 438 | | | 438 | 342 | | | 342 |
| -4; -3 | 802 | | | 802 | 635 | | | 635 |
| -3; -2 | 1,489 | | | 1,489 | 1,248 | | | 1,248 |
| -2; -1 | 2,764 | | | 2,764 | 2,357 | | | 2,357 |
| -1; 0 | | | 5,207 | 5,207 | | | 4,498 | 4,498 |
| 0; 1 | | | 4,780 | 4,780 | | | 5,156 | 5,156 |
| 1; 2 | | 2,526 | | 2,526 | | 3,300 | | 3,300 |
| 2; 3 | | 1,361 | | 1,361 | | 1,691 | | 1,691 |
| 3; 4 | | 738 | | 738 | | 856 | | 856 |
| 4; 5 | | 372 | | 372 | | 475 | | 475 |
| 5; 6 | | 171 | | 171 | | 234 | | 234 |
| 6; 7 | | 108 | | 108 | | 121 | | 121 |
| 7; 8 | | 52 | | 52 | | 40 | | 40 |
| 8; 9 | | 19 | | 19 | | 21 | | 21 |
| 9; 10 | | 12 | | 12 | | 9 | | 9 |
| 10; 11 | | 3 | | 3 | | 1 | | 1 |
| 11; 12 | | 3 | | 3 | | 4 | | 4 |
| 12; 13 | | | | | | 1 | | 1 |
| **Grand Total** | **5,911** | **5,365** | **9,987** | **21,263** | **4,856** | **6,753** | **9,654** | **21,263** |
| **Percentage** | 27.80% | 25.23% | 46.97% | 100.00% | 22.84% | 31.76% | 45.40% | 100.00% |
| **Binomial Test (p-value)** | 2.846e-07 | | | | | < 2.2e-16 | | |

**Data Table.** BioSample, BioProject, and SRA identification numbers. TBD: to be determined.

| Species | Genome Accession | BioSample | BioProject | SRA, PacBio data (CLR) | SRA, PacBio data (CCS) | SRA, PacBio data (IsoSeq) | SRA, Illumina data (DNA) | SRA, Illumina data (RNA) | SRA, Optical Genome Map |
|---|---|---|---|---|---|---|---|---|---|
| *O. alta* | IRGC 105143 | SAMN38217704 | PRJNA1039467 | SRR26965618<br>SRR26965613<br>SRR26965616<br>SRR26965617<br>SRR26965611<br>SRR26965615<br>SRR26965614<br>SRR26965612<br>SRR26965610 | - | SRR26994967 (fasta) | SRR26976184<br>SRR26976183<br>SRR26976182 | - | SUPPF_0000005541 |
| *O. australiensis* | IRGC 100882 | SAMN13386652 | PRJNA591699 | SRR10604178<br>SRR10604179<br>SRR10604180<br>SRR10604181<br>SRR10604182<br>SRR10604183<br>SRR10604184<br>SRR10604185<br>SRR10604186<br>SRR10604187<br>SRR10604188<br>SRR10604189<br>SRR10604190<br>SRR10604191<br>SRR10604192<br>SRR10604177 | - | - | SRR10604302<br>SRR10604303 | - | SUPPF_0000005562 |
| *O. coarctata* | IRGC 104502 | SAMN08744577 | PRJNA439330 | SRR29014502 | SRR28024371<br>SRR28024370<br>SRR28024369<br>SRR28024368 | - | - | SRR29007052<br>SRR29020228<br>SRR29020227 | SUPPF_0000005609 |
| *O. grandiglumis* | IRGC 105669 | SAMN19687255 | PRJNA737282 | SRR14876898<br>SRR14876899 | - | SRR26967108 | SRR14878280 | - | SUPPF_0000003996 |
| *O. latifolia* | IRGC 100890 | SAMN19696891 | PRJNA737486 | SRR14887296<br>SRR14887297 | - | SRR26967215 | SRR14878697 | - | SUPPF_0000003995 |
| *O. longiglumis* | IRGC 106525 | SAMN37370020 | PRJNA1016142 | - | SRR26070823<br>SRR26070823 | SRR26976672 (fasta) | - | - | - |
| *O. malampuzhaensis* | IRGC 80765 | SAMN20924811 | PRJNA757598 | SRR15989943<br>SRR15989944 | - | SRR26953381 | SRR15989942 | SRR26951331 | SUPPF_0000005539 |
| *O. meyeriana* | IRGC 106473 | SAMN38217705 | PRJNA1039468 | - | SRR27033731<br>SRR27033730 | SRR27033726 | - | - | - |
| *O. minuta* | IRGC 101141 | SAMN20924774 | PRJNA757599 | SRR15990419<br>SRR15990420<br>SRR15990421 | - | SRR26953380 | SRR15989941 | SRR26951414 | SUPPF_0000005540 |
| *O. ridleyi* | IRGC 100821 | SAMN17151037 | PRJNA687623 | SRR13358904<br>SRR13358910<br>SRR13358909<br>SRR13358913<br>SRR13358912<br>SRR13358907<br>SRR13358906<br>SRR13358905<br>SRR13358911<br>SRR13358908<br>SRR13358903 | - | SRR26977006 | SRR15096900<br>SRR15096901 | SRR26994926 | SUPPF_0000003862 |
| *O. schlechteri* | IRGC 82047 | SAMN19312591 | PRJNA732115 | TBD | - | TBD | TBD | TBD | SUPPF_0000003977 |
